# Alterations to the broad-spectrum formin inhibitor SMIFH2 improve potency

**DOI:** 10.1101/2022.03.10.483826

**Authors:** Marina Orman, Maya Landis, Aisha Oza, Deepika Nambiar, Joana Gjeci, Kristen Song, Vivian Huang, Amanda Klestzick, Carla Hachicho, Su Qing Liu, Judith M. Kamm, Francesca Bartolini, Jean J. Vadakkan, Christian M. Rojas, Christina L. Vizcarra

## Abstract

SMIFH2 is a small molecule inhibitor of the formin family of cytoskeletal regulators that was originally identified in a screen for suppression of actin polymerization induced by the mouse formin Diaphanous 1 (mDia1). Despite widespread use of this compound, it is unknown whether SMIFH2 inhibits all human formins. Additionally, the nature of protein/inhibitor interactions remains elusive. We assayed SMIFH2 against human formins representing six of the seven mammalian classes and found inhibitory activity against all formins tested. We synthesized a panel of SMIFH2 derivatives and found that, while many alterations disrupt SMIFH2 activity, substitution of an electron-donating methoxy group in place of the bromine along with halogenation of the furan ring increases potency by approximately five-fold. Similar to SMIFH2, the active derivatives are also pan-inhibitors for the formins tested. This result suggests that while potency can be improved, the goal of distinguishing between highly conserved FH2 domains may not be achievable using the SMIFH2 scaffold.

## INTRODUCTION

Members of the formin family of cytoskeletal regulators are associated with a wide variety of essential eukaryotic cellular processes.^1,2^ Formins vary in their biochemical functions, which include actin filament nucleation, processive association with growing filament ends, actin filament bundling, microtubule binding/stabilization, and actin filament severing (reviewed in by Chesarone et al.).^3^ These functions are usually executed by the formin homology (FH) 1 and 2 domains in the C-terminal part of the protein (Fig. 1B). The FH2 domain is conserved from yeast to humans^4,5^ and is a mostly α-helical homodimer with a donut-like architecture that encircles the dynamic “barbed” end of the actin filament.^6–9^ The FH1 domain is an unstructured proline-rich region that binds profilin, an abundant actin monomer-binding protein.^10,11^ Together these domains can modulate the growth of actin filaments^12^ and also displace other barbed end binding proteins.^13,14^ In many formin isoforms, the N-terminal region contains regulatory domains that suppress the activity of the FH1 and FH2 domains.^15–17^

SMIFH2 (1-(3-bromophenyl)-5-(furan-2-ylmethylene)-2-thioxodihydropyrimidine-4,6(1*H*,5*H*)-dione) was discovered in an *in vitro* screen for compounds that inhibit the mDia FH2 domain, hence its name <u>S</u>mall <u>M</u>olecule <u>I</u>nhibitor of <u>FH2</u>.^18^ SMIFH2 is assumed to be a pan-formin inhibitor based on biochemical data showing that it can inhibit yeast formins Bni1, Fus1 and Cdc12, the nematode formin CYK-1, and the mouse formin DIAPH1.^18^ More recent data show SMIFH2 interactions with plant formins including *Arabidopsis* formin-1,^19^ among others. However, the assumption that SMIFH2 is a pan-inhibitor has never been tested *in vitro* for diverse mammalian FH2 domains. To probe how formins contribute to cellular processes, both “pan” inhibitors as well as specific inhibitors are useful. An isoform-specific inhibitor of mouse Dia1 and Dia2 but not Dia3 has been reported, but low solubility in cell culture media has hampered its use.^20^

**Figure 1.**
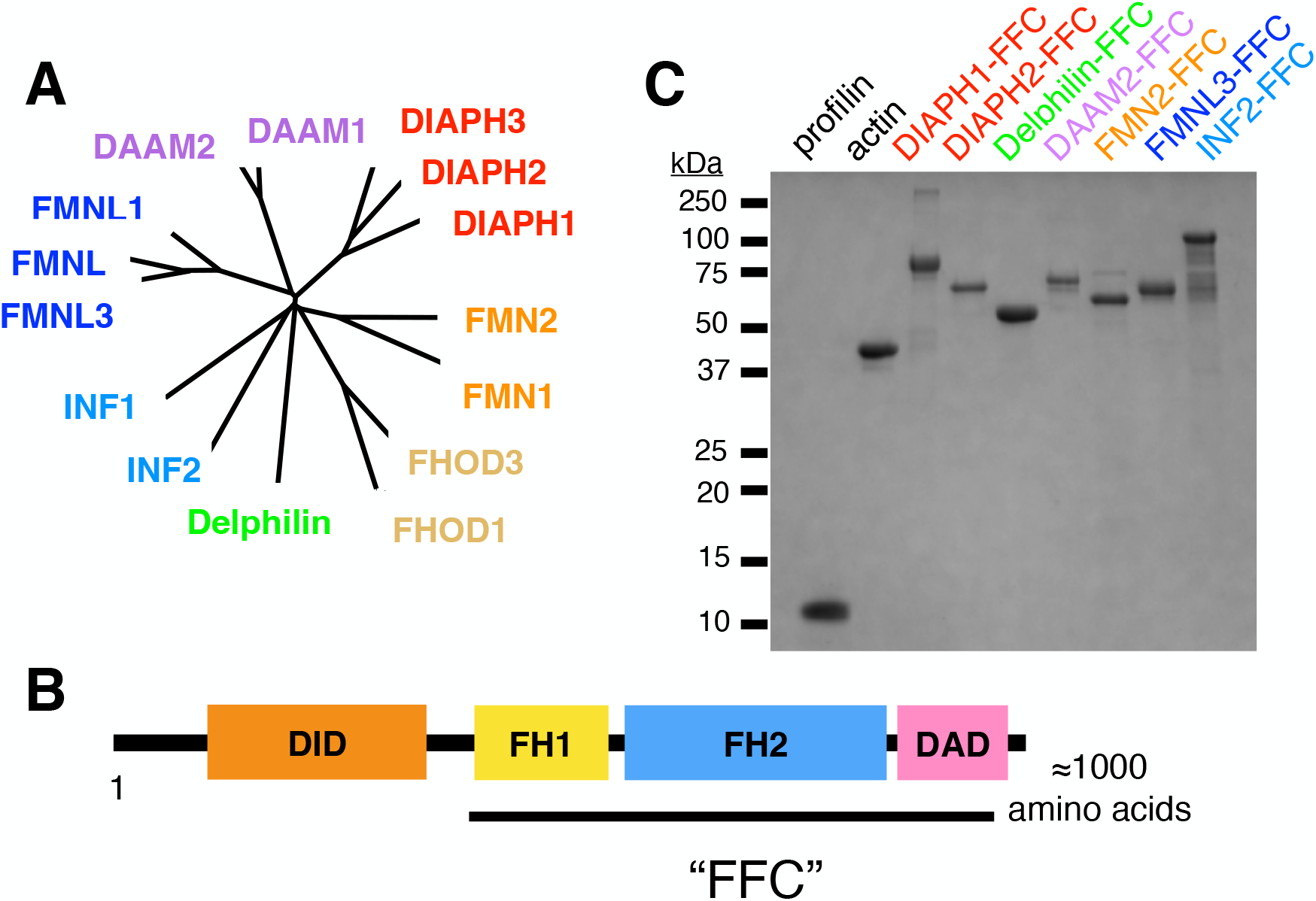
Formin constructs and purification. (A) Phylogenetic tree of formins based on (DeWard, et al.). To assay formin proteins from diverse human formins, we selected of representatives of the six mammalian formin subclasses. (B,C) ‘FFC’ constructs, which include the FH1 and FH2 domains and all C-terminal sequences, were isolated for each formin.

Small molecule inhibitors have been invaluable in studying cytoskeletal function. These include natural product compounds that act directly on cytoskeletal filaments,^21^ as well as the mostly synthetic compounds that target cytoskeletal binding proteins like SMIFH2, the Arp2/3 inhibitor CK666,^22^ and the myosin inhibitor blebbistatin and its analogs.^23^ Due to the importance of formin-mediated cytoskeletal regulation in eukaryotes, SMIFH2 has been used in at least 308 studies of the cellular roles of formins in building both the actin and microtubule cytoskeletons (original manuscripts in a Google Scholar search for ‘SMIFH2’, December 2021).

In the context of this widespread usage, several off-target SMIFH2 interactions have been reported: one with the tumor suppressor p53^24^ and another with several members of the myosin superfamily.^25^ The latter is particularly concerning, given that formins and myosins are both cytoskeletal regulators, and SMIFH2-induced effects on the cytoskeleton may be difficult to interpret in certain settings. Surprisingly, SMIFH2 more potently inhibits the actin-induced ATPase activity of *Drosophila* myosin 5 (IC_50_ ≈ 2 μM) than formin/mediated actin interactions (IC_50_ ≈ 10-20 μM).^25^ These off-target interactions may be related to the fact that SMIFH2 is classified as a <u>P</u>an <u>A</u>ssay <u>IN</u>terference compound (or PAIN) due to the electrophilicity of its α/β-unsaturated dicarbonyl alkylidene moiety.^26^ Both the widespread use and potential drawbacks of SMIFH2 have motivated a search for either optimized analogs of SMIFH2 or even a new generation of formin inhibitors.

In this study, we report a structure-activity study of SMIFH2 against a panel of diverse human formins, using the *in vitro* pyrene-actin polymerization assay.^27^ We find that SMIFH2 is a pan-inhibitor of these human formins, and that perturbations to its structure modulate activity but not specificity.

## RESULTS & DISCUSSION

SMIFH2 was selected from a library of drug-like molecules for its ability to inhibit the mouse formin mDia1 in a pyrene-actin polymerization assay.^18^ We used the pyrene-actin polymerization assay to characterize the inhibitory activity against a panel of human formins representing six of the seven classes of mammalian formins (Fig. 1A). These formins were expressed recombinantly as constitutively active ‘FFC’ fragments (containing FH1, FH2, and C-terminal regions; Fig. 1B) and purified using affinity chromatography (Fig. 1C).

SMIFH2 and its analogs were synthesized as described in the Experimental Procedures. In all NMR characterization experiments, it was observed that SMIFH2 was a mix of *E* and *Z* isomers, with a doubling of each proton peak due to the different environments in each isomer. In some cases, an uneven distribution of isomers was observed immediately after dissolving SMIFH2, as shown for the solvent deuterated tetrahydrofuran (THF-d_8_) (Fig. 2). After incubating at room temperature for 20 hrs, this sample equilibrated to a 1:1 mixture of isomers, as judged by the equivalent peak integrations for each pair. The shift from unequal to equal peak integrations indicates that the equilibrium constant for the isomerization process is close to 1 and that the molecule exchanges between these isomers on an experimentally relevant timescale. We cannot say which peak of each pair corresponds to which isomer. Furthermore, it is unknown whether both isomers or just one isomer inhibits the FH2 domain.

**Figure 2.**
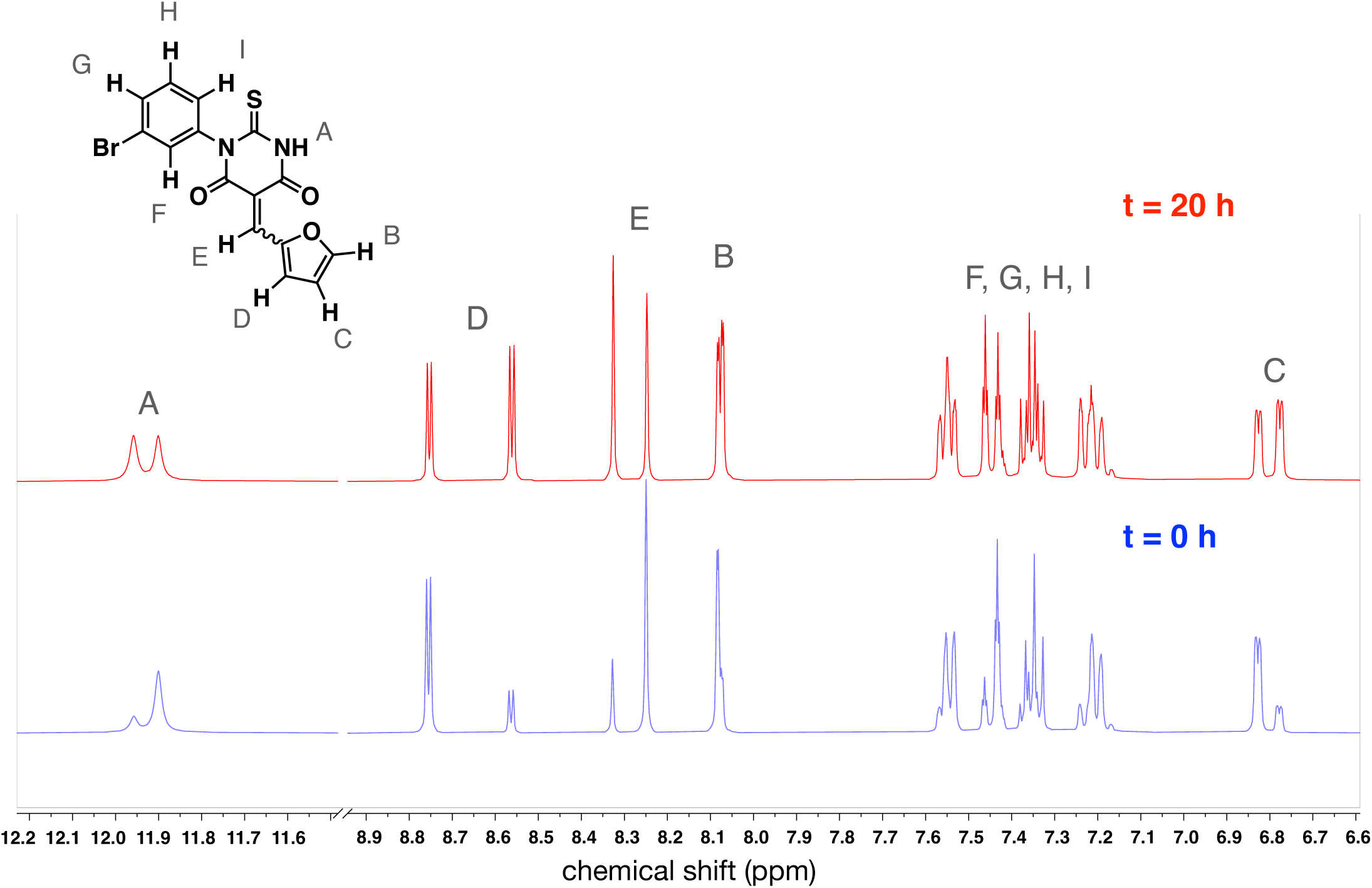
SMIFH2 is a mix of isomers that equilibrate over time. ^1^H NMR spectra of the same sample were collected at two time points. The blue spectrum (bottom) corresponds to the SMIFH2 sample immediately after preparation in THF-*d*_8_. The uneven peak integrations are evident in the signals for protons A, B, C, D and E. The initial isomeric ratio was calculated as 3.5:1 by integration of major resonance at 8.25 ppm and minor resonance at 8.33 ppm for proton E. After 20 h at 25 °C (red spectrum; top), the peak integrations were similar for each isomer, reflecting equilibration to a ≈1:1 *E*:*Z* mixture (isomeric ratio 1.1:1 by integration of resonances at 8.33 ppm and 8.25 ppm).

All FFC fragments were active in pyrene-actin polymerization assays, as judged by their ability to stimulate assembly over the baseline rate of actin/profilin alone (Fig. 3, comparing red and gray curves). As reported previously, diaphanous related formins (DIAPH1 and DIAPH2) and inverted formin 2 (INF2) stimulated assembly at low nanomolar concentrations (5 nM). FMNL3, FMN2, and DAAM2 had intermediate nucleation efficiency (20–25 nM formin induced actin polymerization such that steady state was reached within 2000 s), and Delphilin (25 nM) had quite weak nucleation activity, as already reported for rabbit skeletal muscle actin.^28^

**Figure 3.**
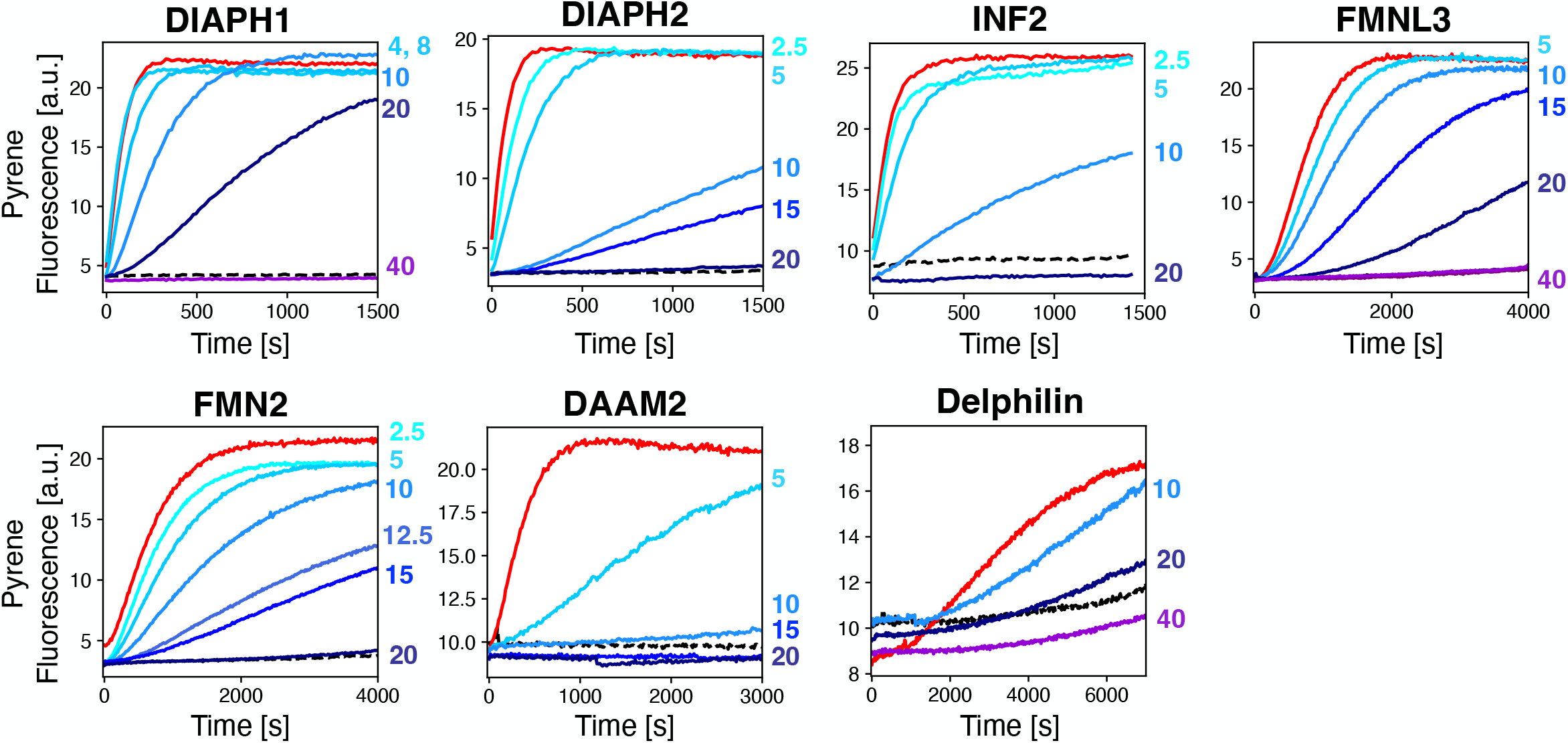
SMIFH2 has broad activity across the human formins. For each titration, pyrene fluorescence was monitored for 2 μM actin (5% pyrene-labeled)/ 4 μM *S. pombe* profilin. The black dashed line shows profilin/actin alone, and the red trace corresponds to addition of formin FFC to profilin/actin (5 nM DIAPH1, DIAPH2, INF2; 20 nM FMNL3; 20 nM FMN2; 25 nM DAAM2; 25 Nm Delphilin). Increasing concentration of SMIFH2 is indicated by blue traces, with the μM concentration labeled to the right of each plot. The volume of DMSO added was the same for all traces, regardless of SMIFH2 concentration.

To characterize the specificity of SMIFH2, we titrated the inhibitor against each purified formin, controlling for the volume of DMSO for all polymerization reactions (Fig. 3, as indicated by blue traces). In all cases, SMIFH2 had a concentration for 50% inhibition (IC_50_) in the 10–20 μM range. This is comparable to the reported value of 15 μM for mouse Dia1.^18^ Among the diaphanous formins, SMIFH2 has a more potent effect on DIAPH2 (Dia3) than on DIAPH1 (Dia1), a notable reversal of preference compared to the quinoxaline-based inhibitor reported by Higgs and coworkers (compound 2 in Gauvin et al.).^20^

Several SMIFH2 analogs are known to retain or lose inhibitory activity against mDia1.^18^ We sought to expand this dataset to more extensively characterize structure-activity relationships in the SMIFH2/formin complex. All 17 analogs shown in Figure 4 were synthesized according to the scheme described in the Experimental Procedures and characterized by proton NMR (Fig. S5). Each inhibitor was tested for potential interaction with actin in the absence of formin (Figs. S1, S2). At the highest concentrations tested, we found that some inhibitors had modest effects on the polymerization of 2 μM actin (Fig. S1). These effects were less apparent in the presence of 4 μM profilin (Fig. S2), indicating that the profilin/actin interaction may mask the inhibitor interaction or suppress polymerization of a potential actin/SMIFH2 complex. This panel of 18 inhibitors was tested against five formin FFC fragments (DIAPH1, DIAPH2, INF2, FMNL3, and FMN2), and the IC_50_ values were measured (Fig. 4 and Table 1). One notable trend in this dataset is that the lack of specificity, noted above for SMIFH2 (Fig. 3), holds for its analogs: no modifications made to SMIFH2 led to a selective inhibitor, at least among the five formins that were tested. In other words, active inhibitors inhibited all formins and inactive inhibitors lacked activity against all formins tested.

**Figure 4.**
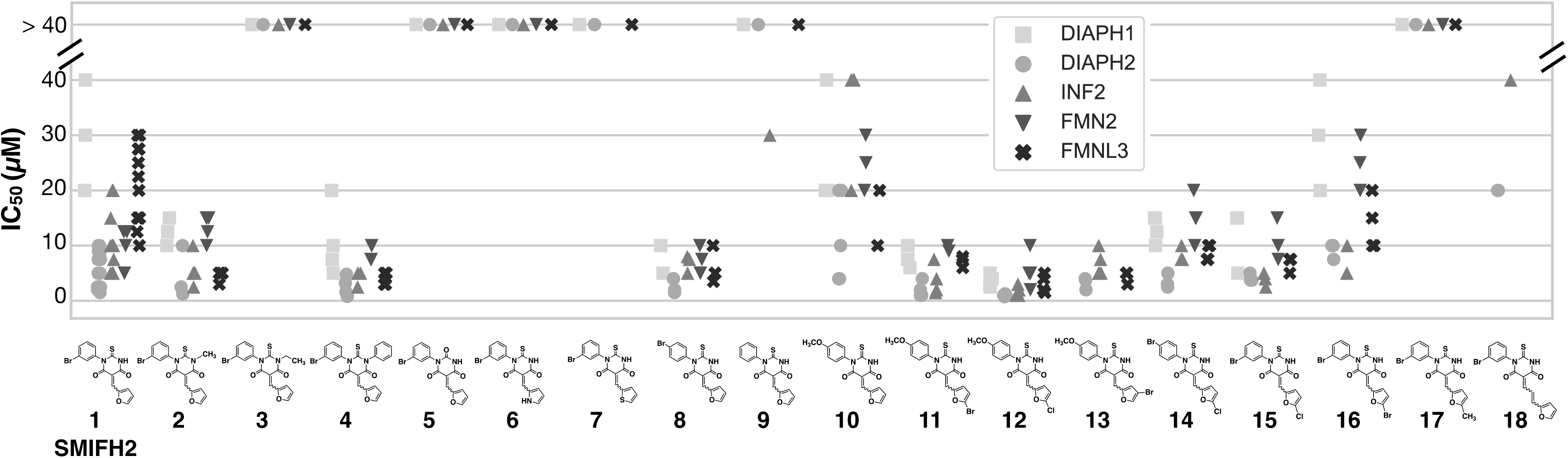
Structure activity data for SMIFH2 and 17 analogs tested against five human formins. Each individual data point corresponds to an IC_50_ value from a titration of inhibitor in the pyrene-actin assembly assay. The average and standard deviation for each formin/analog pair is reported in Table 1. Conditions: 2 μM actin (5% pyrene-labeled), 4 μM *S. pombe* profilin, formin (5 nM DIAPH1-FFC, 5 nM DIAPH2-FFC, 5 nM INF2-FFC, 20-37.5 nM FMN2-FFC, 20 nM FMNL3-FFC), inhibitor with a fixed volume of DMSO. Due to low solubility, compound 9 was dissolved in DMF. Because we cannot reliably measure IC_50_ values above 40 μM, those points are indicated for at the > 40 μM line. Individual IC_50_ values are listed in Table S1.

Modifications to the thiobarbiturate core were of two types: alkylation or arylation of N3 and replacement of the thiocarbonyl with an oxygen carbonyl (atom numbering in gray Table 1 structure). The latter substitution (**5**) led to a complete loss of inhibitory activity (Fig. 4 and Table 1), consistent with data reported by Kovar and coworkers (compound 3 in Rizvi, et al.).^18^ Both methylation (**2**) and arylation (**4**) of the SMIFH2 N3 were well tolerated, and the N3 phenyl analog (**4**) had 3–4 fold more potent inhibitory activity than SMIFH2. Surprisingly, the ethyl-substituted analog (**3**) had no detectable activity. This was unexpected since the high activity of compound **4** showed that the binding pocket on the FH2 domain could accommodate substituents as large as a phenyl ring. One defining feature of PAINs is non-sensical SAR data,^26^ and it is possible that activity data for compounds **2, 3** and **4** are confounded by the other factors apart from the free energy change associated with protein/inhibitor binding. The tolerance of SMIFH2 to methylation (**2**) and arylation (**4**) of N3 further supports a model where a carbonyl tautomer (as drawn in Table 1), not a thio enol or oxygen enol,^29,30^ is the active form of SMIFH2. The activity of compounds **2** and **4** also indicates that the –NH group of SMIFH2 most likely does not act as a hydrogen bond donor in the protein binding pocket.

Meanwhile, several alterations to the furan-ring portion of SMIFH2 had a dramatic effect on activity. Both the pyrrole (**6**) and thiophene (**7**) analogs had no detectable inhibitory effect on any formin tested. On the other hand, halogenation of the furan ring at the C5’ position, either with Cl (**15**) or Br (**16**), was well tolerated, while methylation at that same position (**17**) abolished activity. This is consistent with the high activity of an analog with another halogen (I) at the 5’ position, reported by Kovar and coworkers (compound 2 in Rizvi, et al.).^18^ We synthesized an analog with an extended linker between the furan and central rings. This compound (**18**) had modest activity.

**Table 1.** IC_50_ values for SMIFH2 and 16 analogs tested against five human formins.

|  | X | Y | R<sub>1</sub> | R<sub>2</sub> | R<sub>3</sub> | R<sub>4</sub> | R<sub>5</sub> | DIAPH1 | DIAPH2 | INF2 | FMNL3 | FMN2 |
| --- | --- | --- | --- | --- | --- | --- | --- | --- | --- | --- | --- | --- |
| 1 | S | O | H | Br | H | H | H | 30 ± 10 | 6 ± 3 | 10 ± 5 | 21 ± 7 | 10 ± 3 |
| 2 | S | O | H | Br | CH<sub>3</sub> | H | H | 13 ± 3 | 5 ± 5 | 5 ± 2 | 4 ± 1 | 14 ± 2 |
| 3 | S | O | H | Br | CH<sub>2</sub>CH<sub>3</sub> | H | H | > 40 | > 40 | > 40 | 40 | > 40 |
| 4 | S | O | H | Br | Ph | H | H | 11 ± 7 | 2.3 ± 1.7 | 4 ± 1 | 4 ± 1 | 9 ± 1 |
| 5 | O | O | H | Br | H | H | H | > 40 | > 40 | > 40 | > 40 | > 40 |
| 6 | S | NH | H | Br | H | H | H | > 40 | > 40 | > 40 | > 40 | > 40 |
| 7 | S | S | H | Br | H | H | H | > 40 | > 40 | > 40 | > 40 | n.m.* |
| 8 | S | O | Br | H | H | H | H | 8 ± 3 | 3 ± 1 | 7 ± 2 | 6 ± 3 | 8 ± 4 |
| 9 | S | O | H | H | H | H | H | > 40 | 40 | 30 | > 40 | n.m. |
| 10 | S | O | OCH<sub>3</sub> | H | H | H | H | 27 ± 12 | 12 ± 8 | 37 ± 16 | 17 ± 6 | 23 ± 4 |
| 11 | S | O | OCH<sub>3</sub> | H | H | H | Br | 8 ± 2 | 2.0 ± 1.4 | 3 ± 2 | 7 ± 1 | 10 ± 1 |
| 12 | S | O | OCH<sub>3</sub> | H | H | H | Cl | 4 ± 1 | 1.0 ± 0.2 | 2 ± 1 | 3 ± 2 | 5 ± 3 |
| 13 | S | O | OCH<sub>3</sub> | H | H | Br | H | n.m. | 3.3 ± 0.2 | 6 ± 2 | 4 ± 1 | n.m. |
| 14 | S | O | Br | H | H | H | Cl | 14 ± 2 | 4 ± 1 | 8 ± 1 | 10 | 15 |
| 15 | S | O | H | Br | H | H | Cl | 12 ± 6 | 4 ± 1 | 4 ± 1 | 7 ± 1 | 11 ± 4 |
| 16 | S | O | H | Br | H | H | Br | 30 ± 10 | 9 ± 1 | 7 ± 3 | 14 ± 5 | 25 ± 5 |
| 17 | S | O | H | Br | H | H | CH<sub>3</sub> | > 40 | > 40 | > 40 | > 40 | > 40 |
| 18 | See Figure 4 for structure of 18 |  |  |  |  |  |  | n.m. | 20 | > 40 | n.m. | n.m. |

\*n.m. = not measured;
\*\* cyan: strong inhibition, orange: weak inhibition, gray: no inhibition

SMIFH2 has a Br in the meta position of the N1 ring (R_2_ position in Table 1). The effect of removing this Br was explored with compound **9**, which showed measurable but weak to minimal activity. Compound **9** was poorly soluble in DMSO, and had to be dissolved in DMF for assays. Aside from removing the Br altogether, we found that moving this Br to the para position (**8**) maintained or slightly enhanced inhibitory activity. We were interested in the effects of a methoxy group on the N1 ring, and we successfully synthesized the para-methoxy analog (**10**). While compound **10** had reduced activity compared to meta-Br (**1**) or para-Br (**8**) SMIFH2, substitution of the furan with either Br or Cl at the C5’ position (**11** or **12**, respectively) or Br at the C4’ position (**13**) in addition to the N1 para-methoxy phenyl group appeared to have synergistic effects. This was most striking with compound **12**, the most potent inhibitor in our series of compounds. Including these two substitutions together decreased IC_50_ values by 2–9 fold (comparing compounds **1** and **12**), depending on the formin. For instance, the IC_50_ against INF2 was decreased from 10 ± 5 μM for SMIFH2 (**1**) to 2 ± 1 μM for compound **12**. Unlike incorporation of a halogen at the furan C4’ or C5’ positions in the para-methoxy series (**11, 12, 13**), addition of a Cl at furan C5’ (**14**) did not have increased potency compared with para-Br alone (**8**).

Taken together, the activity data for SMIFH2 and the panel of 17 analogs highlight two essential components of the SMIFH2 structure: the thiocarbonyl and the furan ring. Our results also indicate that SMIFH2 derivatives maintain activity when the structure is modified at the thiobarbiturate N3 position and also at the meta and para positions on the N1-phenyl ring and the 4’ and 5’ positions of the furan ring. Despite the chemical diversity explored in this study, we did not identify an isoform-specific formin inhibitor. Such an inhibitor may exist in another region of SMIFH2 chemical space not explored here, such as ortho and/or multiple substitutions on the N1-phenyl ring or furans attached to the beta carbon at their 3’ carbon, among many other potential modifications. Alternatively, the lack of specificity among the panel of molecules in this study may suggest that a more specific inhibitor may require a scaffold completely different from the thiobarbiturate core of SMIFH2 in order to distinguish among highly conserved FH2 domains. Higgs and coworkers identified a quinoxalinone-based inhibitor with specificity among the diaphanous formins,^20^ suggesting that specific targeting among FH2 domains is achievable.

To date, no atomic-resolution structural data is available for SMIFH2 bound to the FH2 domain, hindering the rational design of structural perturbations to the inhibitor. This is in contrast to the Arp2/3 inhibitor CK666,^22^ whose x-ray crystal structure was used as a starting point for free energy perturbation calculations to optimize inhibitor/protein interactions.^31^ The more potent analogs reported here may be promising candidates for structural studies of a protein/SMIFH2 complex, particularly with the human formin DIAPH2 (ortholog of mDia3). Based on IC_50_ values, SMIFH2 analog **12** may have a higher affinity to DIAPH2 than any other formin/inhibitor pairs measured, although this would need to be confirmed by equilibrium experiments.

To our knowledge, it is uncertain where SMIFH2 lies in the taxonomy of covalent/noncovalent or reversible/irreversible inhibitors. Its α/β unsaturated dicarbonyl motif is likely a Michael acceptor, which would react with nucleophilic amino acids, placing SMIFH2 in the covalent category.^32–34^ The furan substitution at the β carbon of SMIFH2 suggests that addition of a thiol group could be reversible.^35^ Similar compounds to SMIFH2 with a modified thiobarbiturate core, have been found to inhibit mushroom tyrosinase irreversibly,^36^ either through covalent modification of an active site residue or induction of protein misfolding. Cellular studies with SMIFH2 have shown reversal of SMIFH2’s effects after a washout.^37–39^ However, whether this is due to new protein synthesis or dissociation of the SMIFH2/formin complex is unknown.

Motivated by the wide range of inhibitory activities observed in our structure-activity study (Fig. 4 and Table 1) and the potential for covalent targeting by SMIFH2, we used computational modeling to explore differences in molecular structure and reactivity among this group of molecules. We used Gaussian density functional theory calculations at the B3LYP/6311G++(d,p) level of theory and basis set to optimize the geometry of a subset of ten molecules, selected because of their variable activities. Figure 5A shows their optimized geometries in water for both the *E* and *Z* isomers. The optimized geometry was not sensitive to basis set or solvent (Fig. S3). Overall, the molecular shape was similar for all molecules, regardless of their inhibitory activity. Torsion angles for ‘peripheral’ rings with respect to the central thiobarbiturate ring were between 89.9° and 91.5° for the bond connecting N1 of the central ring to the equivalent of the Br-phenyl ring of SMIFH2 and between 179.5° and 180° for the bond connecting C_β_ of the α/β-unsaturated thiobarbiturate to C2’ of the furan ring.

**Figure 5.**
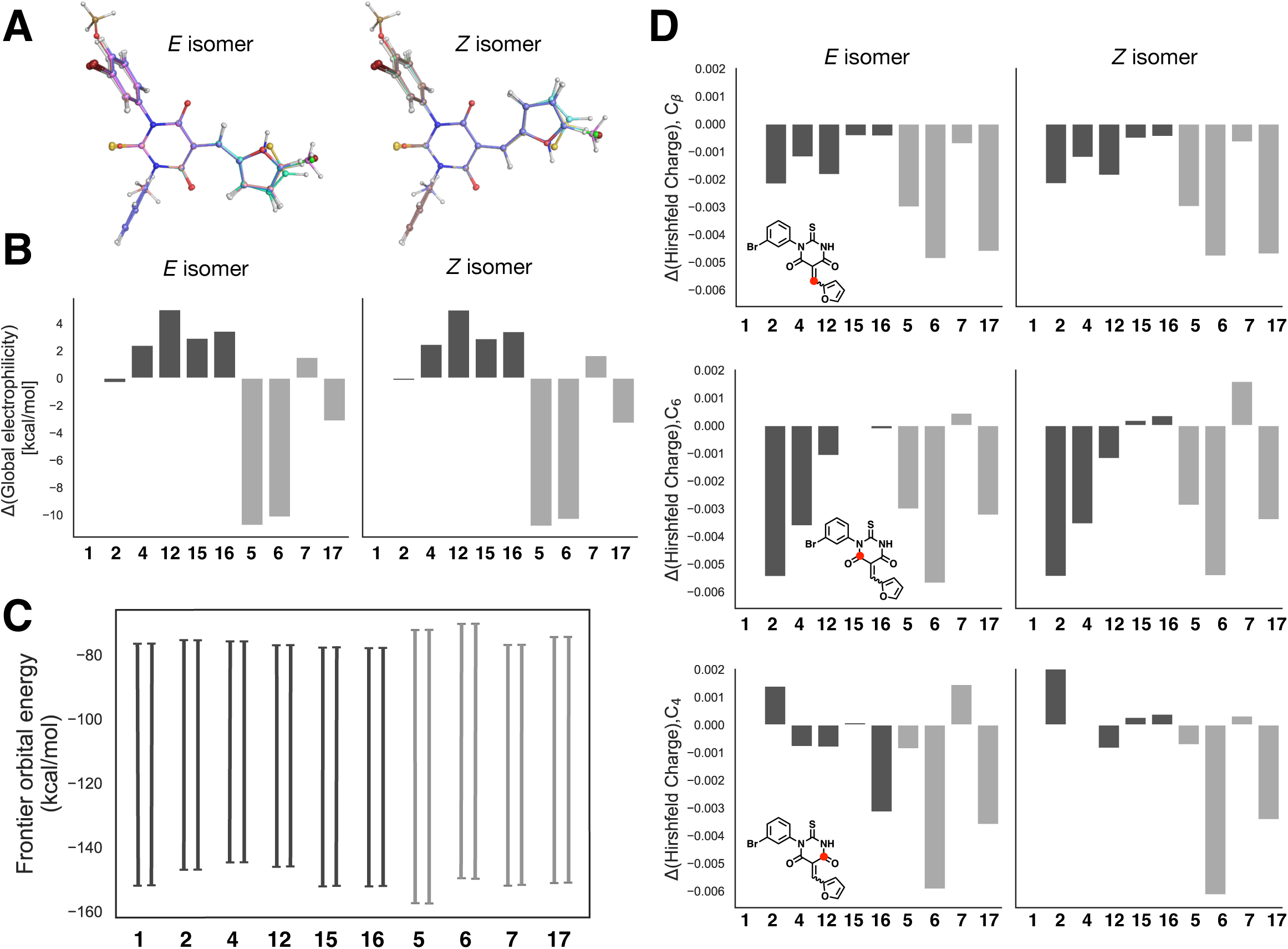
Computational analysis of SMIFH2 analogs. (A) Optimized geometries of SMIFH2 (**1**) and nine analogs aligned in PyMOL based on the thiourea portion (N1-C2-N3) of each molecule. (B) Global electrophilicity plotted as a difference from SMIFH2. (C) Frontier orbital energy plotted showing the gap between the lower energy HOMO and higher energy LUMO. (D) Difference in Hirshfeld charges calculated for SMIFH2 carbons that are potential sites for nucleophilic addition. The specific carbons are indicated by a red dot in the structure exemplified for SMIFH2 (**1**). For panels B, C, and D, less active molecules are denoted with light gray bars and more active molecules are denoted with dark gray bars. All structures, energies, and charges were calculated with Gaussian (see Experimental Procedures for levels of theory and basis sets). See Figure S4 for raw electrophilicity and charge values.

We probed the degree of reactivity of SMIFH2 and these nine derivatives using a global electrophilicity index,^40^ which we calculated from single point energies of water-solvated neutral, anionic and cationic species. We found a trend of higher electrophilicity for active inhibitors (dark bars in Fig. 5B). However, at least one inactive molecule (**7**), which has a thiophene in place of the furan, has an electrophilicity value slightly larger than SMIFH2 (Fig. 5B). This indicates that electrophilicity may be a determinant, but not the sole determinant, of activity. Consistent with higher electrophilicity, active inhibitors also had slightly smaller HOMO/LUMO energy gaps (Fig. 5C). Finally, we calculated Hirshfeld charges for carbons that were potential sites of nucleophilic attack, namely the carbonyl carbons and the carbon in the β position relative to the carbonyls. These charges were not correlated with activity (Fig. 5D). Overall, our computational results indicate these SMIFH2 derivatives have similar geometries and variation in electrophilicity.

The widespread use of SMIFH2 in the decade since the foundational work of Kovar and coworkers^18^ underscores the demand for small molecule formin inhibitors. In addition to the search for formin-specific inhibitors, recent data showing activity of the formin inhibitor SMIFH2 on myosins ^25^ have motivated a search for formin inhibitors with minimal off-target interactions. The structure-activity panel from this study, particularly the SMIFH2 analogs (**2, 4, 11, 12, 13**) with the greatest activity against formins, present a promising starting point for identifying formin inhibitors with minimal off-target interactions. Moving forward, a deeper understanding of mechanisms and specificity of formin-targeting small-molecule treatments is important for understanding the role of formins in cells. We found that while many alterations disrupt SMIFH2 activity, we could increase potency by approximately five-fold. However, we failed to increase specificity, suggesting that while potency can be improved, achieving specificity among closely-related FH2 domains may be hard to achieve with the SMIFH2 scaffold.

## EXPERIMENTAL PROCEDURES

### Plasmid preparation

All formin FFC gene fragments were sub-cloned into a modified pET15b vector with an N-terminal 6-histidine tag, with the exception of DIAPH1, which had a C-terminal 6-histidine tag. Several human genes were codon-optimized for *E. coli* and obtained as g-blocks from Integrated DNA Technologies (Coralville, IA). FFC fragments were defined with amino acid numbers: DIAPH1 (Transomic BC117257, aa 549–1262), DIAPH2 (NP_009293, aa 553–1096), FMNL3 (NP_001354764.1, aa 481-1028), FMN2 (NP_064450.3, aa 1192-1722), Delphilin (NP_001138590, aa 744-1211), INF2 (NP_071934.3, aa 469–1249), and DAAM2 (NP_001188356.1, aa 486-1068). Complete DNA and amino acid sequences are included in the Supporting Information.

### Protein expression and purification

Formin FFC proteins were expressed in ROSETTA-DE3 *E. coli* cells (Novagen) by growing the cells at 37 °C in Terrific Broth to an optical density of 0.5 –1.0 at 600 nm, reducing the temperature to 18 °C for 1 hr, adding 250 μM isopropyl-beta-D-thiogalactoside (IPTG), and growing overnight (≈12–17 hours) at 18°C with shaking at 250 rpm. Cells were harvested, washed with 1× phosphate-buffered saline (PBS), and flash frozen before storing at –80 °C.

INF2-FFC-expressing cells were lysed in Ni-NTA lysis buffer [500 mM NaCl, 50 mM NaPi pH 7.5, 1 mM ethylenediamine tetraacetic acid (EDTA), 1 mM dithiothreitol (DTT)] supplemented with 1 mM phenylmethylsulfonyl fluoride (PMSF), 4 μg/mL DNaseI (Sigma #DN25), 2000-fold diluted protease inhibitor cocktail (Sigma P8849). Cells were lysed by French Press and then centrifugd at 40,000 ×*g* for 30 min. The clarified lysate was passed over a 2-mL HisTrap column (Cytiva) and eluted with Ni-NTA lysis buffer supplemented with 500 mM imidazole. Eluted protein was gel filtered on a Superdex75 16/600 column (Cytiva) equilibrated with SEC buffer [150 mM NaCl, 10 mM HEPES pH 7, 1 mM EDTA, 0.5 mM tris(2-carboxyethyl)phosphine (TCEP)]. Purity was assessed by SDS-PAGE, and the protein was flash frozen in liquid nitrogen and stored at –80 °C.

DAAM2-FFC-expressing cells were lysed, centrifuged, and passed over a HisTrap column as described above for INF2. Pure fractions were pooled and buffer exchanged by PD10 column (Cytiva) into SEC buffer before dialyzing with 1:1 glycerol:SEC buffer overnight. Aliquots were flash frozen and stored at –80 °C.

DIAPH2-FFC expressing cells were lysed by French Press in Ni-NTA lysis buffer supplemented with PMSF, DNaseI and protease inhibitor cocktail and centrifuged, as described for INF2. Clarified lysates were nutated for 1 hr at 4 °C with Ni-NTA resin (Thermo Sci). The resin was washed with 25 column volumes (CV) of Ni-NTA lysis buffer and 25 CV of Ni-NTA lysis buffer supplemented with 10 mM imidazole, before eluting with lysis buffer supplemented with 500 mM imidazole. Eluted protein was buffer exchanged by PD10 column into S-buffer (10 mM NaCl, 10 mM PIPES pH 6.5, 1 mM EDTA, 1mM DTT). The protein was loaded onto a 1-mL SP-FF cation exchange column (Cytiva) and eluted with a gradient from 10 mM to 500 mM NaCl over 30 CV. Selected fractions were exchanged into SEC buffer by PD10 column and then dialyzed into 1:1 glycerol:SEC buffer overnight. Aliquots were flash frozen and stored at –80 °C. FMN2-FFC-expressing cells were lysed in a modified pH 6.5 Ni-NTA lysis buffer. The rest of the purification protocol was identical to that described for DIAPH2-FFC.

FMNL3-FFC-expressing cells were lysed in TALON extraction buffer (300 mM NaCl, 50 mM NaPi pH 8.0, 1 mM β-mercaptoethanol (βME)) supplemented with 1 mM PMSF and 4 μg/mL DNaseI. After lysis and centrifugation as described above for INF2-FFC, the supernatant was nutated with 1 mL TALON resin (Takara) for 30 min at 4 °C. The resin was washed with 25 CV TALON extraction buffer and then 25 CV TALON wash buffer (300 mM NaCl, 50 mM NaPi pH 7, 1 mM βME) before eluting with TALON wash buffer supplemented with 200 mM imidazole. The eluted protein was dialyzed against 50 mM NaPi pH 7, 50 mM NaCl, 1 mM DTT and loaded onto an SP-FF cation exchange column and eluted with a gradient from 50 mM to 500 mM NaCl over 30 CV. Fractions were dialyzed against S-buffer, loaded on an SP-FF column, and eluted with a step gradient from 10 mM NaCl to 500 mM NaCl. Eluted fractions were loaded onto a Superdex200 10/300 column (Cytiva) equilibrated with SEC buffer. Pure fractions were flash frozen in liquid nitrogen and stored at –80 °C. FMNL3 was also purified in a protocol that skipped the first SP-FF column, and there was no difference in activity or purity.

DIAPH1-FFC-expressing cells were lysed by probe sonicator in TALON extraction buffer supplemented with PMSF and DNaseI. The formin was purified using TALON resin as described above for FMNL3-FFC. Eluted fractions were dialyzed against Q-buffer (10 mM Tris pH 8, 1 mM DTT) overnight with 100 units of thrombin to cleave the C-terminal histidine tag. The cleaved protein was loaded onto a monoQ anion exchange column (Cytiva) and eluted with a gradient of 0 to 500 mM KCl over 30 CV. Semi-pure fractions were buffer exchanged into TALON extraction buffer using a PD10 column and incubated with TALON resin to remove uncleaved protein. The fraction that did not bind TALON was collected and dialyzed against Q-buffer and then 1:1 glycerol:Q-buffer. This results in a version of DIAPH1-FFC that is truncated by ∼10-20 residues on the C-terminus as assessed by MALDI mass spectrometry (data not shown).

Delphilin-FFC was purified as described for the human FFC isoform.^28^ Profilin from *Schizosaccharomyces pombe* was expressed in a pET plasmid in BL21-DE3* cells and was purified as described.^41^ Actin was purified from rabbit skeletal muscle and labeled with pyrene-iodoacetamide as described.^42^ For pyrene-labeled actin, the pyrene concentration was calculated using an extinction coefficient of 21,978 M^−1^cm^−1^ at 344 nm, and the actin concentration was calculated using the correction factor of 0.127 at 290 nm ([actin] = (A_290_−0.127 × A_344_) × 38 μM). Extinction coefficients for formins at 280 nm were calculated using Expasy ProtParam^43^: DIAPH1 (22,920 M^−1^cm^−1^), DIAPH2 (25,900 M^−1^cm^−1^), FMNL3 (28,420 M^−1^cm^−1^), FMN2 (36,900 M^−1^cm^−1^), Delphilin (25,440 M^−1^cm^−1^), INF2 (47,440 M^−1^cm^−1^), and DAAM2 (29,910 M^−1^cm^−1^).

### SMIFH2 synthesis and characterization

For the preparation of SMIFH2 and its analogs (Scheme 1), *N*-aryl-substituted thioureas (**A**, X = S, R’ = H) were prepared by treatment of the corresponding anilines with ammonium thiocyanate in aqueous acid. Potassium cyanate gave the urea intermediate (**A**, X = O, R’ = H) en route to analog **5**.^44^ Use of the corresponding alkyl- or phenyl isothiocyanate provided N3-substituted derivatives (**A**, X = S, R’ = Me, Et, Ph). Reaction of the (thio)ureas **A** with diethyl malonate and sodium ethoxide in ethanol solution^29^ formed the thiobarbituric acid core of intermediates **B**. The synthesis of SMIFH2 and analogs (**C**) was completed by Knoevenagel condensation with a selection of aldehydes,^45,46^ either under standard reflux conditions or using a microwave reactor. SMIFH2 and its analogs formed as mixtures of *E*/*Z* isomers and were characterized by ^1^H NMR and, in selected cases, ^13^C NMR and/or HRMS. Further experimental details and characterization data are included in the Supporting Information.

**Scheme 1.**
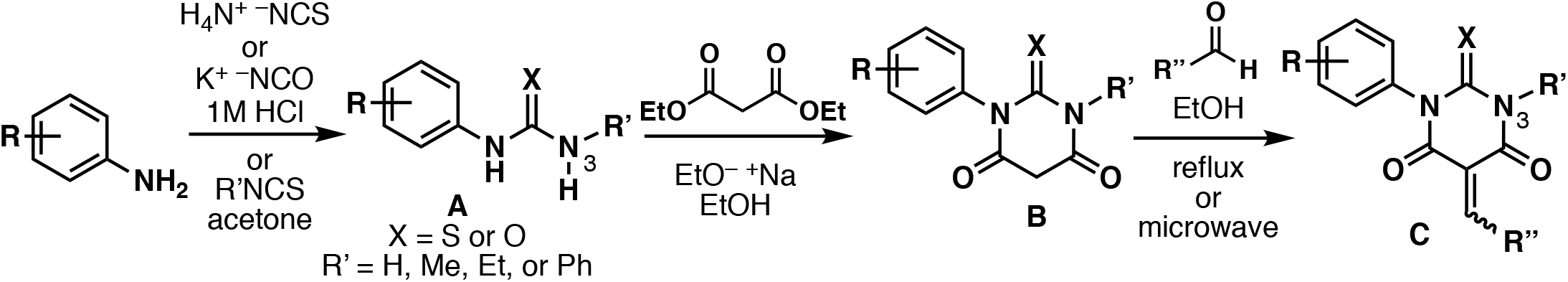

### Pyrene-actin assays

Assays were carried out as described.^47^ Rabbit skeletal muscle actin (5% pyrene labeled) was incubated for 2 min at 25 °C with 200 μM EGTA and 50 μM MgCl_2_ to convert Ca-actin to Mg-actin. When included in the experiment, a 2:1 molar ratio of profilin was incubated with actin for 2 min at 25 °C before conversion to Mg-actin. Polymerization was initiated by adding polymerization buffer (KMEH, final concentration: 10 mM HEPES, pH 7.0, 1 mM EGTA, 50 mM KCl, 1 mM MgCl_2_). Prior to mixing KMEH with actin, formins and then the inhibitor in DMSO or DMF was added at ≤1% of the total volume. Finally, this mixture was rapidly mixed with Mg-actin, with approximately 10-20 s dead time prior to measurement. Fluorescence was monitored every 15 s using a TECAN F200 Pro with λ_ex_ = 360 ± 17 nm and λ_em_ = 415 ± 10 nm. The same volume of organic solvent (DMSO or DMF) was added to each well regardless of inhibitor concentration.

### Computational analysis

SMIFH2 isomers and their chemical analogs were constructed using the program Avogadro.^48^ All quantum mechanical calculations were carried out using Gaussian 16 (versions A-2016 and C-2019).^49^ Geometry optimizations, single point energy, and Hirshfeld population calculations were performed with the unrestricted (U) B3LYP functional with the 6-311+G(d,p) basis set.^50–62^ For geometry optimizations, the opt keyword was set to the ‘tight’ option to ensure adequate convergence. The initial optimization for all compounds was performed in vacuum and used further for solvated calculations reported in this study. Single point energy calculations, for a global electrophilicity analysis, at the neutral, anionic, and cationic states were performed with the unrestricted (U) PW6B95 functional supplemented with Grimme *et al*.’s D3 empirical diffuse function and the aug-cc-pVTZ basis set.^63–67^ SCRF polarizable continuum model (PCM) was used to model an implicit water as the solvent. Optimized geometries were visualized using PyMOL.^68^

To calculate global electrophilicity (ω), we used the convention of Parr et al, ω = μ^2^/(2*η*)where μ and *η* represent the electronic chemical potential and the chemical hardness, respectively.^40,69–72^ Ionization energy and electron affinity were used to calculate μ and *η*, as defined by De Vleeschouwer et al.^73^ We used the (U) PW6B95-D3 functional and aug-cc-pVTZ basis set as described above to determine the global electrophilicity of SMIFH2 and its analogs more accurately.^74^

## Supporting information

Gaussian input files

Coordinates for optimized geometries

## SUPPORTING INFORMATION

Supplementary Experimental Procedures and Compound Characterization Table S1. IC_50_ values for individual trials (related to Table 1)

Figure S1. Effects of SMIFH2 and its derivatives on actin polymerization in the absence of formin.

Figure S2. Effects of SMIFH2 and its derivatives on profilin/actin polymerization in the absence of formin.

Figure S3. Optimized geometries of SMIFH2 using different basis sets and solvents.

Figure S4. Electrophilicity and Hirshfeld charges for SMIFH2 analogs (related to Figure 5)

Figure S5. NMR spectra for SMIFH2 and its derivatives.

Figure S6. Figure S6. ^1^H NMR spectrum of **18** showing second-order, virtual coupling behavior

Figure S7. MestReNova simulation of virtual coupling behavior in **18**

Text file 1. Formin protein sequences. Text file 2. Gaussian input files.

Text file 3. Coordinates for optimized geometries for (U)B3LYP/6-311+G(d,p) (related to Figure 5A).

## ACKNOWLEDGEMENTS

The authors acknowledge the work of the Advanced Chemical Synthesis Laboratory students (Amber Scharnow, Anna Yusov, Jacqueline Chou, Karen Montero, Karol Francisco, Kelsey Lynch, Kiana Harris, Lina Davidson, Maria Paley, Tahrima Jalil, Aisha Hasan, Amanda Sosnowski, Andromeda Urquilla, Choi Mak, Emeline Nguyenduy, Isabel Klein, Jenny Lam, Rachel Dziatko, Ataa Amponsah, Elizabeth Witta, Erica Christensen, Hannah Wentz, Irene Golden, Isabelle Rocroi, Rafaela Brinn, Alaina Hartnett, Allison Forsberg, Anna Hurdle, Emily Latif, Genevieve Nemeth, Janine Sempel, Jessica Glynn, Lucy Zorzano, Shoshana Williams, Tasneem Elkoush, Wendy Xie, Dominique Macaluso, Emily Miura-Stempel, Erika Amemiya, Isabel Hernandez-Rodriguez, Jennifer Niola, Juliet Lee, Kalina Ko, Maya Hoffman, Michelle Lin, Natasha Reich). Dina Merrer, Marisa Buzzeo, and Dalibor Sames for helpful discussion, Rabina Lakha for actin purification, Grace Nickel for help with molecular biology.

