## Supporting Information Table S1, Figures S1-S7 for "Alterations to the broad-spectrum formin inhibitor SMIFH2 improve potency"

**1**....18
**2**....37
**3**....43
**4**....50
**5**....55
**6**....61
**7**....67
**8**....73
**9**....78
**10**....83
**11**....88
**12**....93
**13**....98
**14**....109
**15**....113
**16**....117
**17**....123
**18**....131

Figure S6. $^{1}H$ NMR spectrum of **18** showing second-order, virtual coupling behavior....138

Figure S7. MestReNova simulation of virtual coupling behavior in **18** ....139

*In separate files:*
Text file 1. Formin protein sequences.
Text file 2. Gaussian input files.
Text file 3. Coordinates for optimized geometries for (U)B3LYP/6-311+G(d,p) (related to Figure 5A).

**Experimental Procedures and Compound Characterization Data**

#### General

NMR spectra were recorded in $CDCl_3$, DMSO-$d_6$, or THF-$d_8$ solvent on a Bruker Avance 400 or Bruker Avance 300 spectrometer at 400 MHz or 300 MHz for $^1H$, respectively. The $^1H$ chemical shifts are reported in parts per million ($\delta$), relative to tetramethylsilane (TMS, $\delta$ 0.00) added as an internal standard or the residual solvent proton signals ($CDCl_3$, $\delta$ 7.26; DMSO-$d_6$, $\delta$ 2.50; THF-$d_8$, $\delta$ 3.58). The $^1H$ resonance multiplicities are abbreviated as s (singlet), br s (broad singlet), d (doublet), dd (doublet of doublets), ddd (doublet of doublets of doublets), t (triplet), and m (multiplet). $^{13}C$ NMR chemical shifts in $CDCl_3$ solvent are reported in parts per million relative to the solvent signal ($\delta$ 77.0). The microwave procedure for compound **13** was carried out using a CEM Corporation Discover SP microwave reactor with the, with a ramp temperature to 90 °C at a power of 150 W for 5 min.

#### Preparation of SMIFH2 (1)

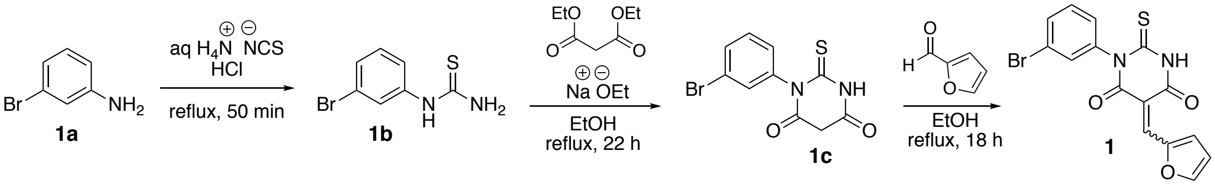

***Synthesis of 1b:*** 3-Bromoaniline (**1a**) (6.1 mL, 56 mmol) was added to a 250 mL round-bottom flask equipped with a magnetic stir bar. Concentrated HCl (6.1 mL) was added dropwise. Clumps formed and were broken up using a glass rod. Ammonium thiocyanate solution (30 wt% in water, 15.2 mL) was added and the reaction mixture was heated to reflux for 50 min. The mixture was then cooled, and a yellow precipitate formed. The contents of the flask were added to 30 mL of cold water, and the precipitate was collected by vacuum filtration. The crude product was recrystallized from hot 4/1 ethanol/water, giving thiourea **1b** as a light-brown crystalline solid (6.2121 g, 48%).

***Synthesis of 1c:*** Anhydrous ethanol (160 mL) was added to a 500 mL round-bottom flask equipped with a magnetic stir bar under nitrogen. Sodium metal (2.020 g, 87.90 mmol) was added to the ethanol, and gas evolution started immediately. After all of the metal had reacted, diethyl malonate (9.7 mL, 64 mmol) was added, and the mixture turned cloudy. The thiourea **1b** (3.6942 g, 15.98 mmol) was added and the mixture was refluxed for 22 h. The reaction mixture was cooled to room temperature. Cold water (160 mL) was added and the mixture turned clear yellow. Ethanol was removed on the rotary evaporator and the remaining mixture was transferred to a separatory funnel and washed with diethyl ether (2 x 160 mL). The aqueous layer was acidified to pH 1 using 2M HCl and a yellow precipitate formed. The light yellow, powdery solid **1c** was collected by vacuum filtration, washed with cold water, and dried (3.5592 g, 74%).

***Synthesis of 1*****:** The thiobarbituric acid **1c** (0.5014 g, 1.676 mmol) was added to a 50 mL round-bottom flask equipped with a magnetic stir bar and a reflux apparatus. Ethanol (15 mL) and furfural (0.14 mL, 1.7 mmol) were added. The mixture was cloudy and yellow as the mixture was heated to reflux for 18 h. A dark-green powder **1d** was collected by vacuum filtration and washed four times with pentane (0.5286 g, 84%).

**Preparation of *p*-methoxy-4'-bromo SMIFH2 analog 13**

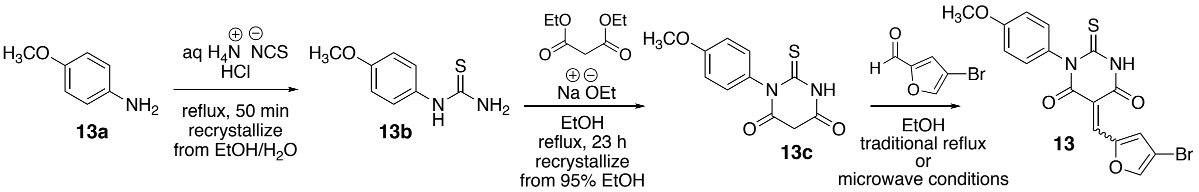

***Synthesis of 13b:*** To 4-anisidine (**13a)** (6.0189 g, 48.87 mmol) in a 250 mL round-bottom flask equipped with a magnetic stir bar was added conc. HCl (6.1 mL) dropwise. Ammonium thiocyanate solution (30 wt % in water, 15.2 mL) was then added and the reaction mixture was heated at reflux for 50 min. The solution was cooled to room temperature and 30 mL of cold water was added. The crude product was collected by vacuum filtration as a green solid (2.0094 g). A batch (0.9931 g) from the crude product was recrystallized from 120 mL of 4/1 ethanol/water, giving *N*-*p*-methoxythiourea **13b** (0.5547 g, 13% based on the proportion of crude product used for recrystallization).

***Synthesis of 13c:*** A 500 mL round-bottom flask equipped with a magnetic stir bar was charged with sodium ethoxide in ethanol (6.24 mL of a 21 wt % solution, 16.7 mmol) under nitrogen, and thiourea **13b** (0.5547 g, 3.04 mmol), additional ethanol (50 mL), and diethyl malonate (1.85 mL, 12.1 mmol) were added. The clear yellow solution was refluxed 23 h then cooled to room temperature and the ethanol was removed by rotary evaporation. The remaining yellow solid was dissolved in room temperature distilled water (20 mL) giving a clear brown solution having pH 11. This solution was acidified to pH 2 by dropwise addition of HCl. A gray precipitate of thiobarbituric acid derivative **13c** formed and was collected by vacuum filtration (0.6916 g, 91%).

***Synthesis of 13 (Traditional):***.When attempting to condense thiobarbituric acid core **13c** with 4-bromo-2-furaldehyde under standard Knoevenagel conditions in refluxing ethanol (cf. the final step in the SMIFH2 (**1**) synthesis), we noted concurrent decomposition of the desired product **13**. Thus, $^1$H NMR analysis of aliquots from the reaction mixture indicated that some conversion to **13** occurred over about 1 h reaction time, but the product signals diminished before all starting material was consumed. The Knoevenagel product **13** could be isolated in very low yield (approx. 3%) by limiting the reflux time to 15 min, followed by purification of **13** by chromatography on silica gel. As a more robust synthetic alternative, we found that microwave heating over 5 min, as described below, gave better results for the preparation of SMIFH2 analog **13**.

***Synthesis of 13 (Microwave):*** Thiobarbituric acid **13c** (0.2194 g, 0.877 mmol) and ethanol (15 mL) were added to a 100 mL round-bottom flask equipped with a magnetic stir bar. To the mixture was added 4-bromo-2-furaldehyde (0.1552 g, 0.887 mmol). The resulting yellow suspension was heated in a microwave reactor at 90 °C with the flask open to the atmosphere for 5 min. During this time, almost all the ethanol evaporated, leaving a pitch-black solid. The small amount of remaining ethanol (<1 mL) was removed in vacuo, and the remaining solid was well stirred in chloroform (50 mL). Insoluble material was removed by filtration and the chloroform filtrate was concentrated to give the product as a black powder, also containg a minor amount of the starting 4-bromo-2-furaldehyde as determined by $^1$H NMR analysis. The residual aldehyde could be removed under reduced pressure (high-vacuum line, overnight), affording the Knoevenagel condensation product **13** (0.2828 g, 0.6944 mmol, 79%).

**Characterization Data for SMIFH2 (1) and Analogs (2–18)**

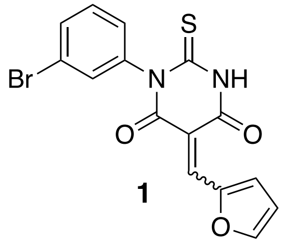

**1**

**1. SMIFH2.** Data for a mixture of *E* and *Z* isomers (isomeric ratio 1.2 : 1 by integration of major resonance at 8.40 ppm and minor resonance at 8.61 ppm; identity of isomers not assigned). *Resonances for the major isomer. **Due to chemical shift overlap, we could not distinguish the major and minor isomer signals. $^1$H NMR (400 MHz, DMSO-$d_6$) $\delta$ 12.83* (s, 1H), 12.78 (s, 1H), 8.61 (d, $J$ = 3.8 Hz, 1H), 8.40* (d, $J$ = 3.9 Hz, 1H), 8.36 (d, $J$ = 1.9 Hz, 1H), 8.34* (d, $J$ = 1.8 Hz, 1H), 8.16* (s, 1H), 8.06 (s, 1H), 7.65-7.58 (m, 3H), 7.56 (dd, $J$ = 2.0, 2.0 Hz, 1H), 7.44* (dd, $J$ = 8.0, 8.0 Hz, 1H), 7.43 (dd, $J$ = 8.0, 8.0 Hz, 1H), 7.37-7.29 (m, 2H), 7.02-6.97 (m, 1H), 6.93-6.88* (m, 1H).

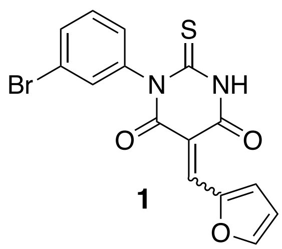

**1**

**1. SMIFH2 in THF-*d*$_8$ analyzed both immediately following NMR sample preparation and also 20 hours later.** Data for a mixture of *E* and *Z* isomers (isomeric ratio 3.5 : 1 by integration of major resonance at 8.25 ppm and minor resonance at 8.33 ppm; identity of isomers not assigned). SMIFH2 t = 20 h (THF) (isomeric ratio 1.1 : 1 by integration of major resonance at 8.33 ppm and minor resonance at 8.25 ppm; identity of isomers not assigned). *Resonances for the major isomer. **Due to chemical shift overlap, we could not distinguish the major and minor isomer signals. $^1$H NMR (400 MHz, THF-$d_8$) $\delta$ 11.96 (s, 1H), 11.90* (s, 1H), 8.76* (d, $J$ = 3.8 Hz, 1H), 8.56 (d, $J$ = 3.8 Hz, 1H), 8.33 (s, 1H), 8.25* (s, 1H), 8.08* (d, $J$ = 1.1 Hz, 1H), 8.07 (d, $J$ = 1.5 Hz, 1H), 7.57 (ddd, $J$ = 5.6, 1.5, 0.9 Hz, 1H), 7.54* (ddd, $J$ = 7.7, 1.2, 0.5 Hz, 1H), 7.46 (dd, $J$ = 1.9, 1.9 Hz, 1H), 7.43* (dd, $J$ = 2.0, 2.0 Hz, 1H), 7.36 (dd, $J$ = 8.0, 8.0 Hz, 1H), 7.35* (dd, $J$ = 8.0, 8.0 Hz, 1H), 7.23 (ddd, $J$ = 8.0, 1.8, 0.9 Hz, 1H), 7.20* (ddd, $J$ = 7.9, 1.7, 0.9 Hz, 1H) 6.83* (ddd, $J$ = 3.7, 1.4, 0.7 Hz, 1H), 6.78 (ddd, $J$ = 4.1, 1.5, 0.8 Hz, 1H).

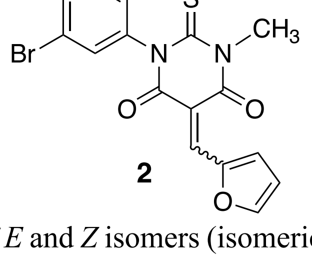

**2**

**2. N3-Methyl.** Data for a mixture of *E* and *Z* isomers (isomeric ratio 1.4 : 1 by integration of major resonance at 6.79 ppm and minor resonance at 6.87 ppm; identity of isomers not assigned). *Resonances for the major isomer. **Due to chemical shift overlap, we could not distinguish the major and minor isomer signals. $^1$H NMR (400 MHz, THF-$d_8$) $\delta$ 8.76 (d, $J$ = 3.9 Hz, 1H), 8.59* (d, $J$ = 3.9 Hz, 1H), 8.40* (apparent s, 1H), 8.31 (apparent s, 1H), 8.12 (d, $J$ = 1.1 Hz, 1H), 8.10* (d, $J$ = 1.1 Hz, 1H), 7.55* (ddd, $J$ = 8.2, 2.0, 1.0 Hz, 1H), 7.53 (ddd, $J$ = 6.3, 1.9, 0.9 Hz, 1H), 7.43* (dd, $J$ = 2.0, 2.0 Hz, 1H), 7.40 (dd, $J$ = 1.9, 1.9 Hz, 1H), 7.36* (dd, $J$ = 8.0, 8.0 Hz, 1H), 7.35 (dd, $J$ = 8.0, 8.0 Hz, 1H), 7.20* (ddd, $J$ = 8.1, 1.9, 1.1 Hz, 1H), 7.17 (ddd, $J$ = 8.0, 1.9, 1.0 Hz, 1H), 6.87 (ddd, $J$ = 2.3, 1.7, 0.8 Hz, 1H), 6.79* (ddd, $J$ = 2.2, 1.5, 0.7 Hz, 1H), 3.77 (s, 3H), 3.75* (s, 3H).

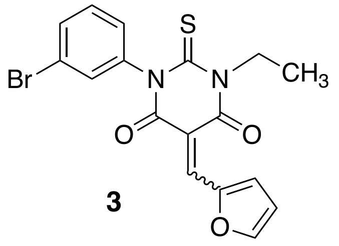

**3**

**3. N3-Ethyl.** Data for a mixture of $E$ and $Z$ isomers (isomeric ratio 1.5 : 1 by integration of major resonance at 6.71 ppm and minor resonance at 6.80 ppm; identity of isomers not assigned). *Resonances for the major isomer. **Due to chemical shift overlap, we could not distinguish the major and minor isomer signals. $^1$H NMR (400 MHz, $CDCl_3$) $\delta$ 8.79 (d, $J$ = 3.9 Hz, 1H), 8.63* (d, $J$ = 3.9 Hz, 1H), 8.55* (s, 1H), 8.45 (s, 1H), 7.92 (d, $J$ = 1.9 Hz, 1H), 7.90* (d, $J$ = 1.8 Hz, 1H), 7.60* (ddd, $J$ = 8.7, 2.0, 1.1 Hz, 1H), 7.58 (ddd, $J$ = 8.7, 1.9, 1.0 Hz, 1H), 7.39* (t, $J$ = 8.0 Hz, 1H), 7.39* (t, $J$ = 2.2 Hz, 1H), 7.37 (t, $J$ = 7.6 Hz, 1H), 7.37 (t, $J$ = 1.9 Hz, 1H), 7.17* (ddd, $J$ = 7.9, 2.1, 1.0 Hz, 1H), 7.15 (ddd, $J$ = 8.1, 2.0, 1.0 Hz, 1H), 6.80 (ddd, $J$ = 4.0, 1.2, 0.8 Hz, 1H), 6.71* (ddd, $J$ = 3.7, 1.4, 0.7 Hz, 1H), 4.59 (q, $J$ = 7.0 Hz, 2H), 4.59* (q, $J$ = 7.0 Hz, 2H), 1.36 (t, $J$ = 7.0 Hz, 3H), 1.35* (t, $J$ = 7.0 Hz, 3H).

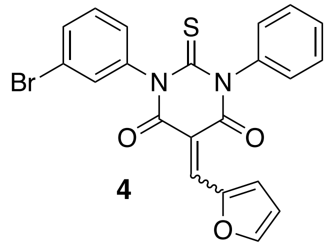

**4**

**4. N3-Phenyl.** Data for a mixture of $E$ and $Z$ isomers (exact isomeric ratio not clear due to overlap of peaks but approximately the ratio is 1 : 1; identity of isomers not assigned). $^1$H NMR (400 MHz, DMSO-$d_6$) $\delta$ 8.49 (d, $J$ = 4.0 Hz, 1H), 8.48 (d, $J$ = 4.0 Hz, 1H), 8.38 (d, $J$ = 1.6 Hz, 1H), 8.38 (d, $J$ = 1.6 Hz, 1H), 8.22 (s, 1H), 8.22 (s, 1H), 7.64 - 7.25 (m, 18H), 6.97 - 6.92 (m, 2H).

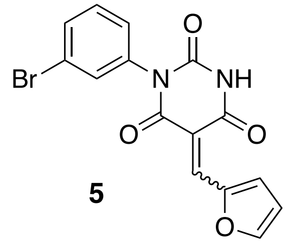

**5**

**5. Pyrimidine trione.** Data for a mixture of $E$ and $Z$ isomers (isomeric ratio 2.3 : 1 by integration of major resonance at 11.76 ppm and minor resonance at 11.69 ppm; identity of isomers not assigned). *Resonances for the major isomer. **Due to chemical shift overlap, we could not distinguish the major and minor isomer signals. $^{1}$H NMR (400 MHz, DMSO-$d_6$) $\delta$ 11.76* (s, 1H), 11.69 (s, 1H), 8.54 (d, $J$ = 3.8 Hz, 1H), 8.36* (d, $J$ = 3.8 Hz, 1H), 8.30 (d, $J$ = 1.7 Hz, 1H), 8.29* (d, $J$ = 1.7 Hz, 1H), 8.15* (s, 1H), 8.08 (s, 1H), 7.67 - 7.62** (m, 3H), 7.61 (dd, J = 1.9, 1.9 Hz, 1H), 7.46* (dd, $J$ = 7.8, 7.8 Hz, 1H), 7.45 (dd, $J$ = 7.9, 7.9 Hz, 1H), 7.38* (ddd, $J$ = 8.1, 1.4, 1.4 Hz, 1H), 7.36 (m, 1H), 6.96 (ddd, $J$ = 3.9, 2.3, 0.8 Hz, 1H), 6.88* (ddd, $J$ = 3.9, 1.7, 0.7 Hz, 1H).

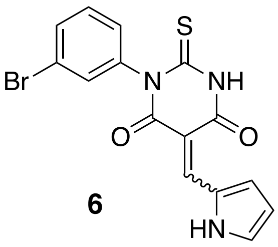

**6**

**6. Pyrrole.** Data for a mixture of $E$ and $Z$ isomers (isomeric ratio 1.2 : 1 by integration of major resonance at 8.23 ppm and minor resonance at 8.12 ppm; identity of isomers not assigned). *Resonances for the major isomer. **Due to chemical shift overlap, we could not distinguish the major and minor isomer signals. $^{1}$H NMR (400 MHz, DMSO-$d_6$) $\delta$ 13.19 (br s, 1H), 12.75* (br s, 1H), 12.69 (br s, 1H), 12.65* (br s, 1H), 8.23* (s, 1H), 8.13 (s, 1H), 7.78 (br s, 1H), 7.65** (dd, $J$ = 2.0, 2.0 Hz, 1H), 7.63 (br s, 1H), 7.61 (ddd, $J$ = 7.9, 1.9, 1.2 Hz, 1H), 7.59* (ddd, $J$ = 7.9, 1.9, 1.2 Hz, 1H), 7.58** (dd, $J$ = 1.9, 1.9 Hz, 1H), 7.44* (dd, $J$ = 7.9, 7.9 Hz, 1H), 7.44 (br, 1H), 7.43 (dd, $J$ = 8.0, 8.0 Hz, 1H), 7.38* (ddd, $J$ = 8.0, 1.5, 1.5 Hz, 1H), 7.32 (ddd, $J$ = 8.0, 1.9, 1.2 Hz, 1H), 6.61 (ddd, $J$ = 4.1, 2.1, 2.1Hz, 1H), 6.56* (ddd, $J$ = 4.1, 2.1, 2.1 Hz, 1H).

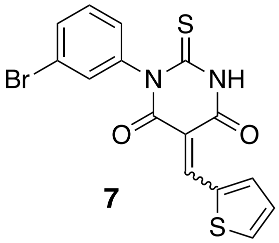

**7**

**7. Thiophene.** Data for a mixture of $E$ and $Z$ isomers (isomeric ratio 1.3 : 1 by integration of major resonance at 8.73 ppm and minor resonance at 8.61 ppm; identity of isomers not assigned). *Resonances for the major isomer. **Due to chemical shift overlap, we could not distinguish the major and minor isomer signals. $^{1}$H NMR (400 MHz, DMSO-$d_6$) $\delta$ 12.77* (s, 1H), 12.76 (s, 1H), 8.73* (s, 1H), 8.61 (s, 1H), 8.40 (ddd, $J$ = 5.1, 1.2, 1.2 Hz, 1H), 8.34* (ddd, $J$ = 1.2, 1.2, 5.1 Hz, 1H), 8.29* (dd, $J$ = 4.0, 1.1 Hz, 1H), 8.26 (dd, $J$ = 4.0, 1.1 Hz, 1H), 7.65-7.58 (m, 6H), 7.47-7.32 (m, 4H); HRMS (ASAP+) $m/z$ calcd for $C_{15}H_{10}N_2O_2S_2{}^{81}Br$ $(M+H)^+$ 394.9346, found 394.9344.

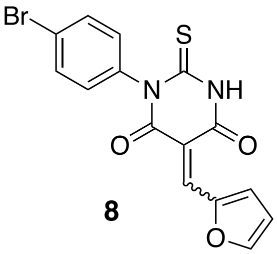

**8**

**8. *p*-Bromo.** Data for a mixture of *E* and *Z* isomers (isomeric ratio 1.4 : 1 by integration of major resonance at 8.39 ppm and minor resonance at 8.61 ppm; identity of isomers not assigned). *Resonances for the major isomer. **Due to chemical shift overlap, we could not distinguish the major and minor isomer signals. $^1$H NMR (400 MHz, DMSO-$d_6$) $\delta$ 12.82* (s, 1H), 12.77 (s, 1H), 8.61 (d, $J$ = 3.9 Hz, 1H), 8.39* (d, $J$ = 3.9 Hz, 1H), 8.35** (d, $J$ = 5.7 Hz, 2H), 8.15* (s, 1H), 8.06 (s, 1H), 7.67 (apparent d, $J$ = 8.4 Hz, 1H), 7.66* (apparent d, $J$ = 8.5 Hz, 1H), 7.27* (apparent d, $J$ = 8.8 Hz, 1H), 7.25 (apparent d, $J$ = 8.9 Hz, 1H), 7.01-6.97 (m, 1H), 6.93-6.89* (m, 1H).

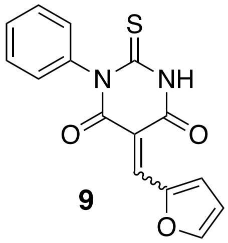

**9**

**9. Debromo.** Data for a mixture of *E* and *Z* isomers (isomeric ratio 1.1 : 1 by integration of major resonance at 8.16 ppm and minor resonance at 8.07 ppm; identity of isomers not assigned). *Resonances for the major isomer. **Due to chemical shift overlap, we could not distinguish the major and minor isomer signals. $^1$H NMR (300 MHz, DMSO-$d_6$) $\delta$ 12.78* (s, 1H), 12.73 (s, 1H), 8.61 (d, $J$ = 3.8 Hz, 1H), 8.40* (d, $J$ = 3.9 Hz, 1H), 8.36 (apparent d, $J$ = 1.6 Hz, 1H), 8.34* (apparent d, $J$ = 1.5 Hz, 1H), 8.16* (s, 1H), 8.07 (s, 1H), 7.49-7.39** (m, 6H), 7.31-7.25** (m, 4H), 6.99 (apparent ddd, $J$ = 2.3, 1.5, 0.8 Hz, 1H), 6.90* (apparent ddd, $J$ = 2.4, 1.6, 0.8 Hz, 1H).

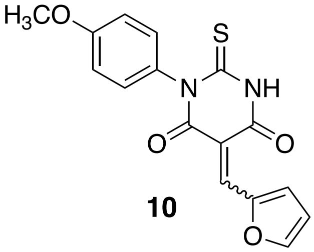

**10**

**10. *p*-Methoxy.** Data for a mixture of *E* and *Z* isomers (isomeric ratio 1.1 : 1 by integration of major resonance at 8.40 ppm and minor resonance at 8.60 ppm; identity of isomers not assigned). *Resonances for major isomer. **Due to chemical shift overlap, we could not distinguish the major and minor isomer signals. $^1$H NMR (300 MHz, DMSO-$d_6$) $\delta$ 12.74* (s, 1H), 12.68 (s, 1H), 8.60 (d, $J$ = 3.9 Hz, 1H), 8.40* (d, $J$ = 3.8 Hz, 1H), 8.34 (d, $J$ = 1.3 Hz, 1H), 8.33* (d, $J$ = 1.3 Hz, 1H), 8.14* (s, 1H), 8.06 (s, 1H), 7.21-7.13** (m, 2H), 7.21-7.13** (m, 2H), 7.03-6.95** (m, 5H), 6.90** (ddd, $J$ = 3.9, 1.5, 0.7 Hz, 1H), 3.80* (s, 3H), 3.79 (s, 3H).

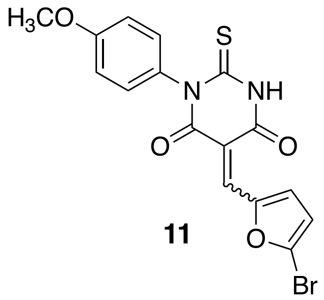

**11**

**11.** ***p*-Methoxy-5'-bromo.** Data for a mixture of *E* and *Z* isomers (isomeric ratio 1.1 : 1 by integration of major resonance at 8.31 ppm and minor resonance at 8.52 ppm; identity of isomers not assigned). *Resonances for major isomer. **Due to chemical shift overlap, we could not distinguish the major and minor isomer signals. $^1$H NMR (400 MHz, DMSO-$d_6$) $\delta$ 12.76* (s, 1H), 12.71 (s, 1H), 8.52 (d, $J$ = 3.9 Hz, 1H), 8.32* (d, $J$ = 3.9 Hz, 1H), 8.03* (s, 1H), 7.95 (s, 1H), 7.18** (apparent d, $J$ = 7.9 Hz, 2H), 7.16** (apparent d, $J$ = 7.8 Hz, 2H), 7.11 (d, $J$ = 3.7 Hz, 1H), 7.03* (d, $J$ = 3.8 Hz, 1H), 7.02-6.95** (m, 2H), 7.02-6.95** (m, 2H), 3.80* (s, 3H), 3.79 (s, 3H).

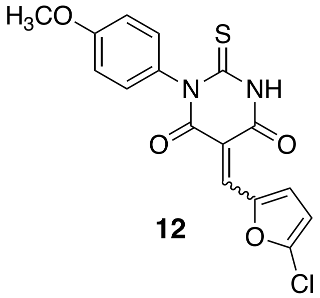

**12**

**12.** ***p*-Methoxy-5'-chloro.** Data for a mixture of *E* and *Z* isomers (isomeric ratio 1.2 : 1 by integration of major resonance at 8.58 ppm and minor resonance at 8.38 ppm; identity of isomers not assigned). *Resonances for the major isomer. **Due to chemical shift overlap, we could not distinguish the major and minor isomer signals. $^1$H NMR (400 MHz, DMSO-$d_6$) $\delta$ 12.77* (s, 1H), 12.73 (s, 1H), 8.58 (d, $J$ = 3.9 Hz, 1H), 8.38* (d, $J$ = 4.0 Hz, 1H), 8.01* (s, 1H), 7.93 (s, 1H), 7.18** (apparent d, $J$ = 8.6 Hz, 2H), 7.16** (apparent d, $J$ = 8.3 Hz, 2H), 7.02 (d, $J$ = 3.9 Hz, 1H), 6.99* (apparent d, $J$ = 9.0 Hz, 2H), 6.98 (d, $J$ = 9.0 Hz, 2H), 6.94* (d, $J$ = 3.9 Hz, 1H), 3.80* (s, 3H), 3.79 (s, 3H).

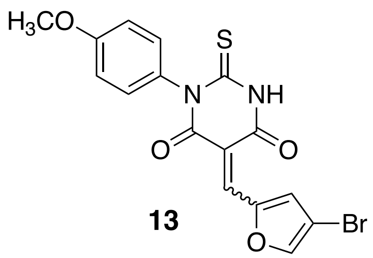

**13**

**13.** ***p*-Methoxy-4'-bromo.** Data for a mixture of *E* and *Z* isomers (isomeric ratio 1.1 : 1 by integration of major resonance at 8.71 ppm and minor resonance at 8.85 ppm; identity of isomers not assigned). *Resonances for the major isomer. **Due to chemical shift overlap, we could not distinguish the major and minor isomer signals. $^1$H NMR (400 MHz, $CDCl_3$) $\delta$ 9.82 (s, 1H), 9.55* (d, $J$ = 2.9 Hz, 1H), 8.85 (s, 1H), 8.71* (s, 1H), 8.41* (s, 1H), 8.33 (s, 1H), 7.87 (d, $J$ = 0.4 Hz, 1H), 7.86* (d, $J$ = 0.4 Hz, 1H), 7.16** (apparent d, $J$ = 8.9 Hz, 2H), 7.14** (apparent d, $J$ = 8.9 Hz, 2H), 7.05** (apparent d, $J$ = 8.9 Hz, 2H), 7.02** (apparent d, $J$ = 8.9 Hz, 2H), 3.87* (s, 3H), 3.86 (s, 3H); $^{13}$C NMR (100 MHz, $CDCl_3$) $\delta$ 178.4, 178.3, 161.7, 160,1, 160.0, 159.9, 159.2, 158.0, 151.3, 151.2, 149.3, 149.1, 141.7, 141.1, 130.8, 130.8, 130.7, 130.4, 129.5, 129.4, 114.9, 114.8, 112.7, 112.5, 105.2, 105.1, 55.4 (2C); HRMS (ESI+) *m/z* calcd for $C_{16}H_{12}N_2O_4S^{81}Br$ $(M+H)^+$ 408.9682, found 408.9713.

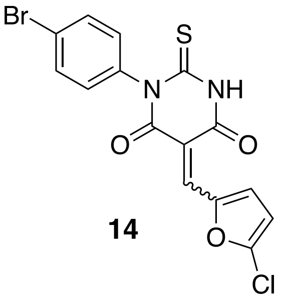

**14**

**14. *p*-Bromo-5'-chloro.** Data for a mixture of *E* and *Z* isomers (isomeric ratio 1.1:1.0 by integration of major resonance at 12.85 ppm and minor resonance at 12.81 ppm; identity of isomers not assigned). *Resonances for major isomer. **Due to chemical shift overlap, we could not distinguish the major and minor isomer signals. $^1$H NMR (400 MHz, DMSO-$d_6$) $\delta$ 12.85* (br s, 1H), 12.81 (apparent d, $J$ = 2.9 Hz, 1H), 8.57 (d, $J$ = 3.9 Hz, 1H), 8.37* (d, $J$ = 4.0 Hz, 1H), 8.01* (s, 1H), 7.92 (s, 1H), 7.72-7.63** (m, 2H), 7.72-7.63** (m, 2H), 7.29-7.22** (m, 2H), 7.29-7.22** (m, 2H), 7.24** (m, 1H), 7.17** (dd, $J$ = 4.0, 1.2 Hz, 1H).

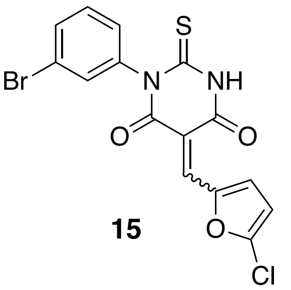

**15**

**15. 5'-Chloro.** Data for a mixture of *E* and *Z* isomers (isomeric ratio 2.9 : 1 by integration of major resonance at 6.63 ppm and minor resonance at 6.55 ppm; identity of isomers not assigned). *Resonances for the major isomer. **Due to chemical shift overlap, we could not distinguish the major and minor isomer signals. $^1$H NMR (300 MHz, $CDCl_3$) $\delta$ 9.39** (s, 1H), 9.39** (s, 1H), 8.83* (d, $J$ = 4.0 Hz, 1H), 8.67 (d, $J$ = 4.0 Hz, 1H), 8.40 (s, 1H), 8.31* (s, 1H), 7.62** (m, 1H), 7.62** (m, 1H), 7.45-7.35** (m, 3H), 7.45-7.35** (m, 3H), 6.63* (d, $J$ = 4.1 Hz, 2H), 6.55 (d, $J$ = 4.1 Hz, 2H).

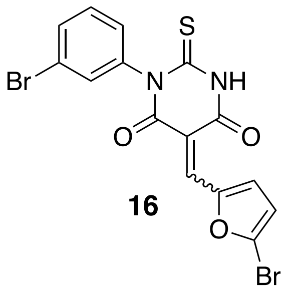

**16**

**16. 5’-Bromo.** Data for a mixture of *E* and *Z* isomers (isomeric ratio 1.2 : 1 by integration of major resonance at 8.33 ppm and minor resonance at 8.54 ppm; identity of isomers not assigned). *Resonances for major isomer. **Due to chemical shift overlap, we could not distinguish the major and minor isomer signals. $^{1}$H NMR (300 MHz, DMSO-$d_6$) $\delta$ 12.89* (s, 1H), 12.84 (s, 1H), 8.54 (d, $J$ = 3.8 Hz, 1H), 8.33* (d, $J$ = 3.9 Hz, 1H), 8.05* (s, 1H), 7.96 (s, 1H), 7.62** (ddd, $J$ = 7.9, 2.0, 1.1 Hz, 1H), 7.62** (ddd, $J$ = 7.9, 2.0, 1.0 Hz, 1H), 7.59** (dd, $J$ = 1.7, 1.7 Hz, 1H), 7.56** (dd, $J$ = 1.8, 1.8 Hz, 1H), 7.44** (dd, $J$ = 7.9 Hz, 1H), 7.44** (dd, $J$ = 8.0 Hz, 1H), 7.34* (ddd, $J$ = 7.9, 1.7, 1.1 Hz, 1H), 7.31 (ddd, $J$ = 7.9, 1.7, 1.0 Hz, 1H), 7.13 (dd, $J$ = 4.0, 0.6 Hz, 1H), 7.04* (dd, $J$ = 3.9, 0.6 Hz, 1H).

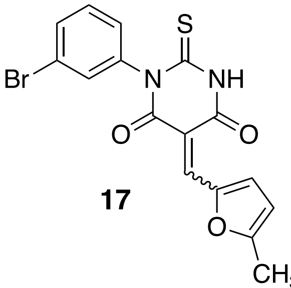

**17**

**17. 5’-Methyl.** Data for a mixture of *E* and *Z* isomers (isomeric ratio 1.2 : 1 by integration of major resonance at 8.85 ppm and minor resonance at 8.68 ppm; identity of isomers not assigned). *Resonances for major isomer. **Due to chemical shift overlap, we could not distinguish the major and minor isomer signals. $^{1}$H NMR (300 MHz, $CDCl_3$) $\delta$ 9.58* (s, 1H), 9.50 (s, 1H), 8.85* (d, $J$ = 3.9 Hz, 1H), 8.58 (d, $J$ = 3.9 Hz, 1H), 8.45 (s, 1H), 8.35* (s, 1H), 7.65-7.56** (m, 1H), 7.65-7.56** (m, 1H), 7.45-7.35** (m, 2H), 7.45-7.35** (m, 2H), 7.24-7.17** (m, 1H), 7.24-7.17** (m, 1H), 6.52* (d, $J$ = 3.9 Hz, 1H), 6.44 (d, $J$ = 3.9 Hz, 1H), 2.52* (s, 3H), 2.50 (s, 3H).

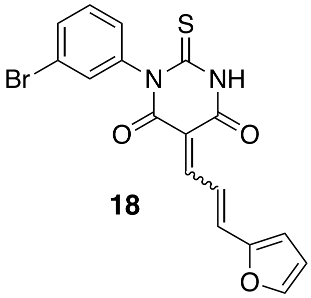

**18**

**18. Vinylene.** Data for a mixture of *E* and *Z* isomers (isomeric ratio 1.0 : 1 by integration of resonances at 6.77 ppm and 6.74 ppm; identity of isomers not assigned). $^{1}$H NMR (400 MHz, DMSO-$d_6$) $\delta$ 12.68 (s, 1H), 12.65 (s, 1H), 8.29 (dd, $J$ = 15.1, 12.4 Hz, 1H), 8.19-8.09 (m, 2H)$^{\ddagger}$, 8.07 (d, $J$ = 1.7 Hz, 1H), 8.05 (d, $J$ = 12.6 Hz, 1H), 8.00 (d, $J$ = 1.6 Hz, 1H), 7.69 (d, $J$ = 14.9 Hz,

1H), 7.74-7.65 (m, 1H)‡, 7.61 (ddd, *J* = 7.9, 1.9, 1.0 Hz, 1H), 7.60 (ddd, *J* = 8.0, 2.0, 1.0 Hz, 1H), 7.58 (dd, *J* = 1.9, 1.9 Hz, 1H), 7.56 (dd, *J* = 1.9, 1.9 Hz, 1H), 7.43 (dd, *J* = 8.0, 8.0 Hz, 1H), 7.42 (dd, *J* = 8.0, 8.0 Hz, 1H), 7.33 (ddd, *J* = 8.0, 1.7, 1.0 Hz, 1H), 7.31 (ddd, *J* = 8.0, 1.8, 1.0 Hz, 1H). 7.13 (d, *J* = 3.3 Hz, 1H), 7.12 (d, *J* = 3.2 Hz, 1H), 6.77 (dd, *J* = 3.5, 1.8 Hz, 1H), 6.74 (dd, *J* = 3.5, 1.8 Hz, 1H); HRMS (ASAP+) *m/z* calcd for $C_{17}H_{12}N_2O_3S^{81}Br$ $(M+H)^+$ 404.9732, found 404.9725.

‡The $^{1}$H NMR resonances for **18** at $\delta$ 8.19-8.09 (2H) and $\delta$ 7.74-7.65 (1H) arise from a second-order 3-spin system that exhibits virtual coupling. As an example of the phenomenon of virtual coupling in $^{1}$H NMR, analysis of this spin system is provided below, including comparison to a simulated spectrum (see also Figure S6 and Figure S7).

**$^{1}$H NMR Analysis of a Mixture of Double Bond Geometrical Isomers in Compound 18, One of Which Has a Virtually Coupled 3-Spin System**

**18**

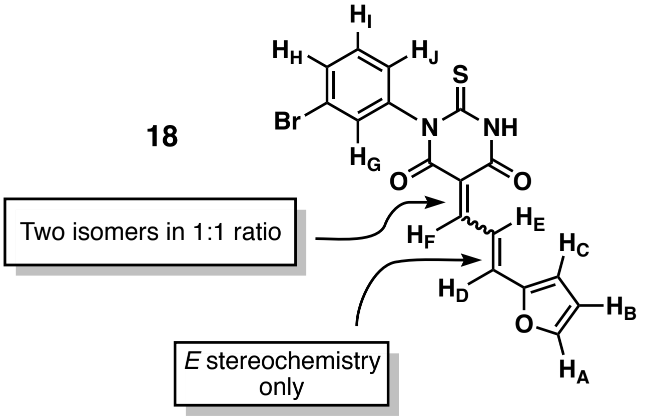

As is typical of SMIFH2 and its analogs, the sample of vinylene SMIFH2 analog **18** for $^{1}$H NMR analysis consisted of a 1:1 mixture of stereoisomers at the double bond indicated above.

The signals for each of the two stereoisomers were assigned by *J*-value and COSY analysis, although which set of resonances corresponds to the *E* versus *Z* isomers at the trisubstituted double bond was not established. Also, there are three isolated spin systems in the $^{1}$H NMR:

1. $H_A$, $H_B$, $H_C$;
2. $H_D$, $H_E$, $H_F$; and
3. $H_G$, $H_H$, $H_I$, $H_J$.

Correlations of nuclei within each spin system were determined, but not further among the different spin systems.

The $^{1}$H NMR (DMSO-$d_6$) spectrum of the one of the double bond isomers (labeled as isomer 1' in Figure S6) has resonances for $H_D$, $H_E$, and $H_F$ with widely spaced chemical shifts. The coupling constants measured for that first-order system are $J_{DE}$ = 15.1 Hz and $J_{EF}$ = 12.4 Hz.

The other stereoisomer (labeled as isomer 2'), however, has $H_E$ and $H_F$ strongly coupled in the 400 MHz NMR spectrum (i.e., $J_{FE} > \Delta\nu_{FE}$), leading to second-order, virtual coupling behavior in the 3-spin system consisting of $H_D$, $H_E$, and $H_F$.

For the second-order, virtually coupled 3-spin system ($H_D$, $H_E$, and $H_F$) of isomer 2', spin simulation using the $J$ values from the other stereoisomer ($J_{DE}$ = 15.1 Hz and $J_{EF}$ = 12.4 Hz) and chemical shifts estimated from the observed spectrum, reproduced the observed $^1$H NMR spectrum (compare the actual spectrum in Figure S6 with the simulation in Figure S7).

Initial analysis was made more challenging by the fact that both of the two lines of the first-order $H_D$ doublet for isomer 1' at $\delta$ 7.69 overlap with two of the six lines observed for the virtually coupled $H_D$ signal of the other stereoisomer, isomer 2'.

Benzene-$d_6$ was titrated into the DMSO-$d_6$ NMR sample, and $^1$H NMR spectra were obtained at DMSO-$d_6$/ $C_6D_6$ ratios of 16:1, 8:1, and 4:1 (at which point there was the beginning of line deformation, presumably because of solubility/miscibility issues). While the solvent change did remove some signal overlap between the two geometrical isomers' resonances, it did not remove the chemical shift coincidence (close coupling) of $H_F$ and $H_E$ in isomer 1' and the 3-spin system ($H_D$, $H_E$, and $H_F$) of that geometrical isomer remained second-order in all the solvent mixtures examined.

### Table S1. IC$_{50}$ values for individual trials *(related to Table 1)*

| | SMIFH2 1 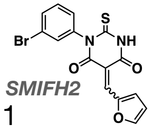 | 2 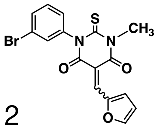 | 3 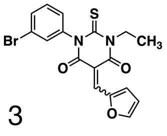 | 4 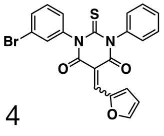 | 5 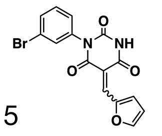 | 6 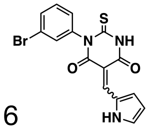 | 7 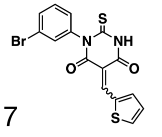 | 8 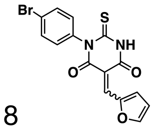 | 9 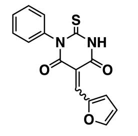 |
|---|---|---|---|---|---|---|---|---|---|
| **DIAPH1** | 20, 30, 40 | 15, 12.5, 10 | > 40 | 10, 20, 5, 7.5 | > 40 | > 40 | > 40 | 10, 10, 5 | > 40 |
| **DIAPH2** | 2.5, 1.5, 2, 2, 2.5, 10, 10, 10, 7.5, 5, 2.5, 8.75, 9, 9, 7.5, 7.5, 10, 7.5, 8 | 10, 1.25, 2.5 | > 40 | 1, 0.8, 1.6, 4.8, 3.2 | > 40 | > 40 | > 40 | 2, 4, 1.5 | 40 |
| **INF2** | 5, 7.5, 5, 5, 5, 10, 10, 10, 20, 15, 10, 20, 10 | 10, 5, 5, 5, 5, 2.5, 5 | > 40 | 5, 5, 5, 2.5 | > 40 | > 40 | > 40 | 8, 5, 7.5 | 30 |
| **FMNL3** | 10, 12.5, 20, 30, 25, 27.5, 22.5, 27.5, 15, 15, 15, [illegible] | 15, 15, 15, 10, 12.5 | 40, 40 | 5, 3, 3, 5, 4 | > 40 | > 40 | > 40 | 5, 5, 10, 3.5 | > 40 |
| **FMN2** | 12.5, 12.5, 10, 5 | 5, 5, 3, 5 | > 40 | 10, 10, 7.5, 7.5 | > 40 | > 40 | n.m. | 5, 10, 5, 7.5 | n.m. |

| | 10 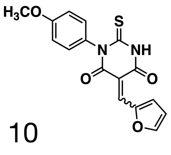 | 11 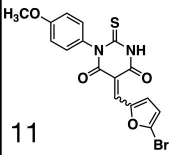 | 12 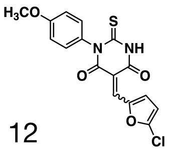 | 13 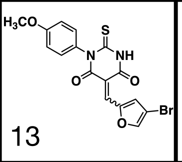 | 14 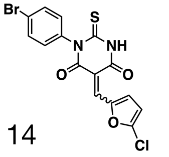 | 15 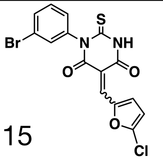 | 16 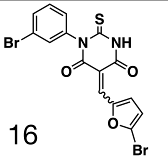 | 17 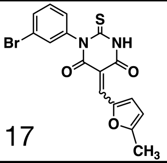 | 18 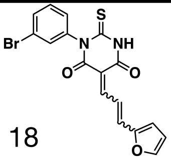 |
|---|---|---|---|---|---|---|---|---|---|
| **DIAPH1** | 20, 20, 40 | 6, 10, 7.5, 10 | 4, 2.5 5, 4, 3 | n.m. | 15, 15, 10, 15, 12.5 | 15, 15, 5 | 20, 40, 30 | > 40 | n.m. |
| **DIAPH2** | 20, 4, 4, 20, 10 | 2, 1, 1, 4 | 1, 0.75, 1.25, 1, 1 | 3.75, 4. 2 | 5, 2.5, 3 | 5, 3.75, 3.75 | 10, 10, 10, 7.5 | > 40 | 20, 20, 20 |
| **INF2** | 40, 40, 20, 40, 40, 40 | 4, 2, 2, 1.5, 7.5 | 1, 1, 2, 3 | 5,10,5, 5, 7.5 | 7.5, 7.5, 7.5, 10 | 2.5, 5, 4 | 5, 5, 10 | > 40 | > 40 |
| **FMNL3** | 10, 20, 20 | 8, 7.5, 6 | 2, 1.5, 5, 4 | 5, 3, 5 | 10, 10, 7.5, 7.5 | 7.5, 7.5, 7.5, 5 | 10, 10, 15, 20 | > 40 | n.m. |
| **FMN2** | 25, 20, 30 | 10, 8, 10 | 10, 5, 5, 5, 2 | n.m. | 15, 15, 10, 20 | 10, 15, 7.5 | 30, 20, 25 | > 40 | n.m. |

** n.m. = not measured, all values are in µM*

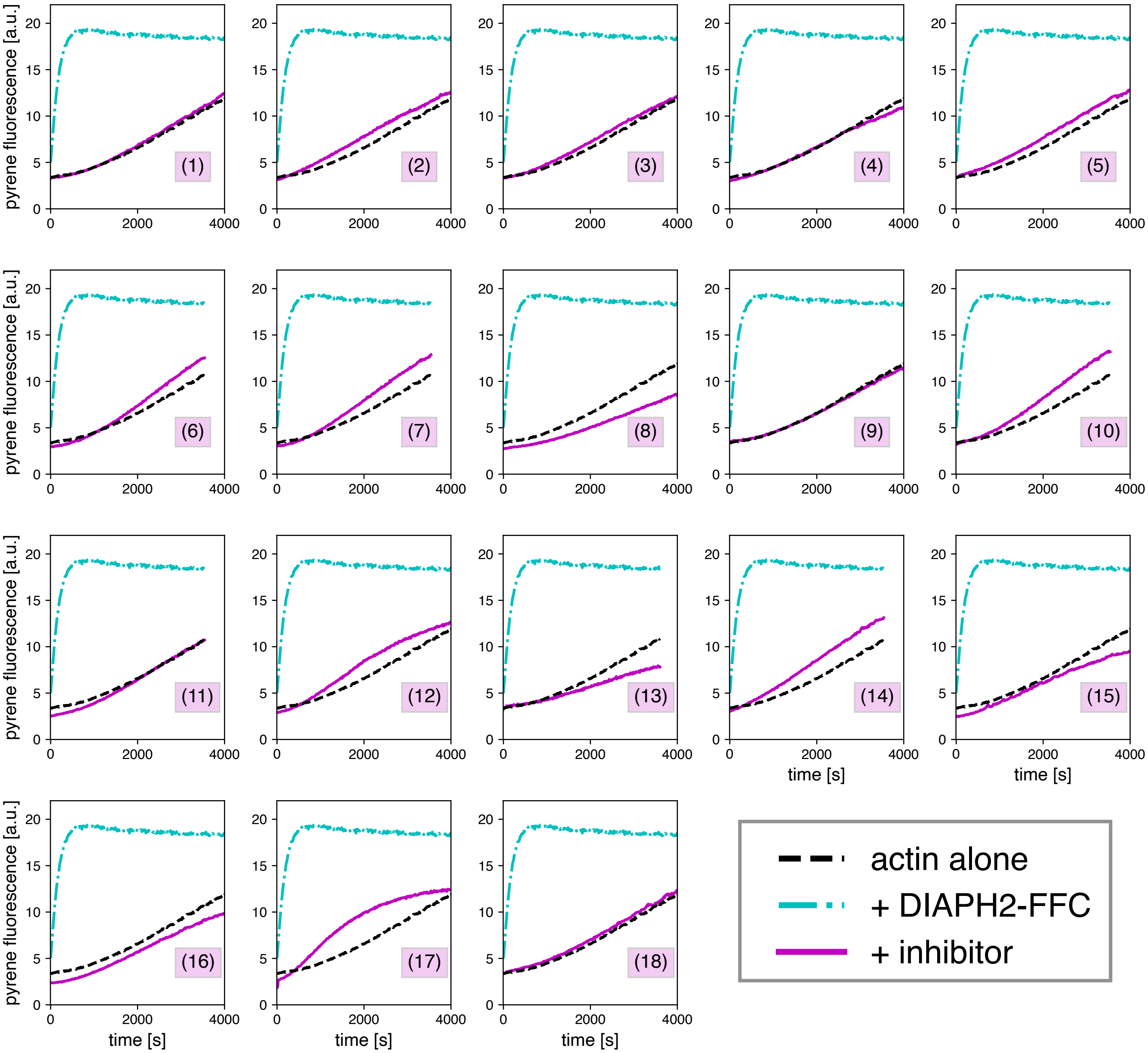

**Figure S1. Effects of SMIFH2 and its derivatives on actin polymerization in the absence of formin.** Pyrene-actin polymerization assays with SMIFH2 analogs in the absence of formin (magenta traces). Polymerization induced by the formin DIAPH2 is shown for reference. *Conditions:* 2 µM actin (5% labeled), 5 nM DIAPH2, 40 µM SMIFH2 analog.

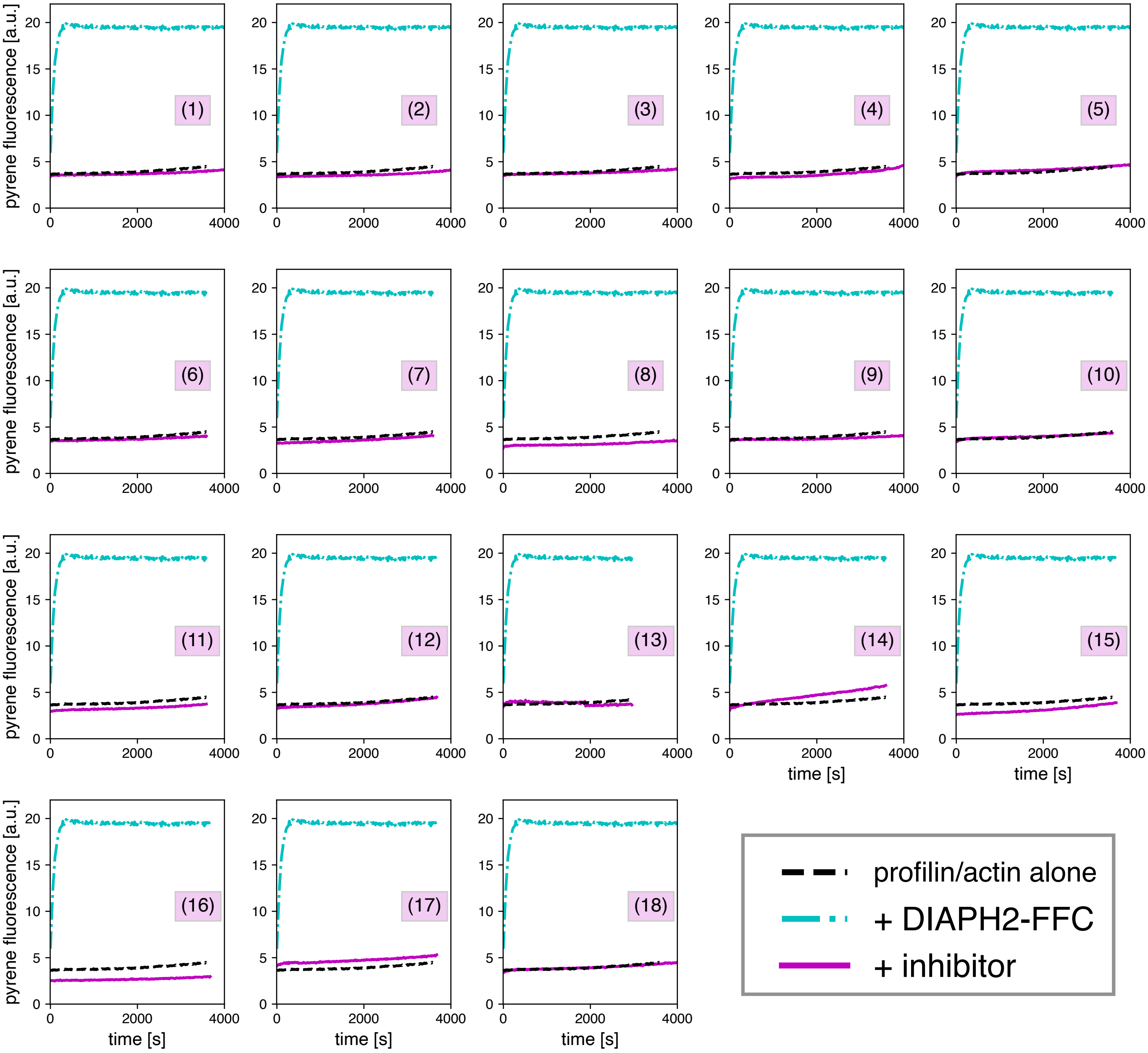

**Figure S2. Effects of SMIFH2 and its derivatives on profilin/actin polymerization in the absence of formin.** Pyrene-actin polymerization assays with SMIFH2 analogs in the absence of formin (magenta traces). Polymerization induced by the formin DIAPH2 is shown for reference. *Conditions:* 2 µM actin (5% labeled)/ 4 µM *S. pombe* profilin, 5 nM DIAPH2, 40 µM SMIFH2 analog.

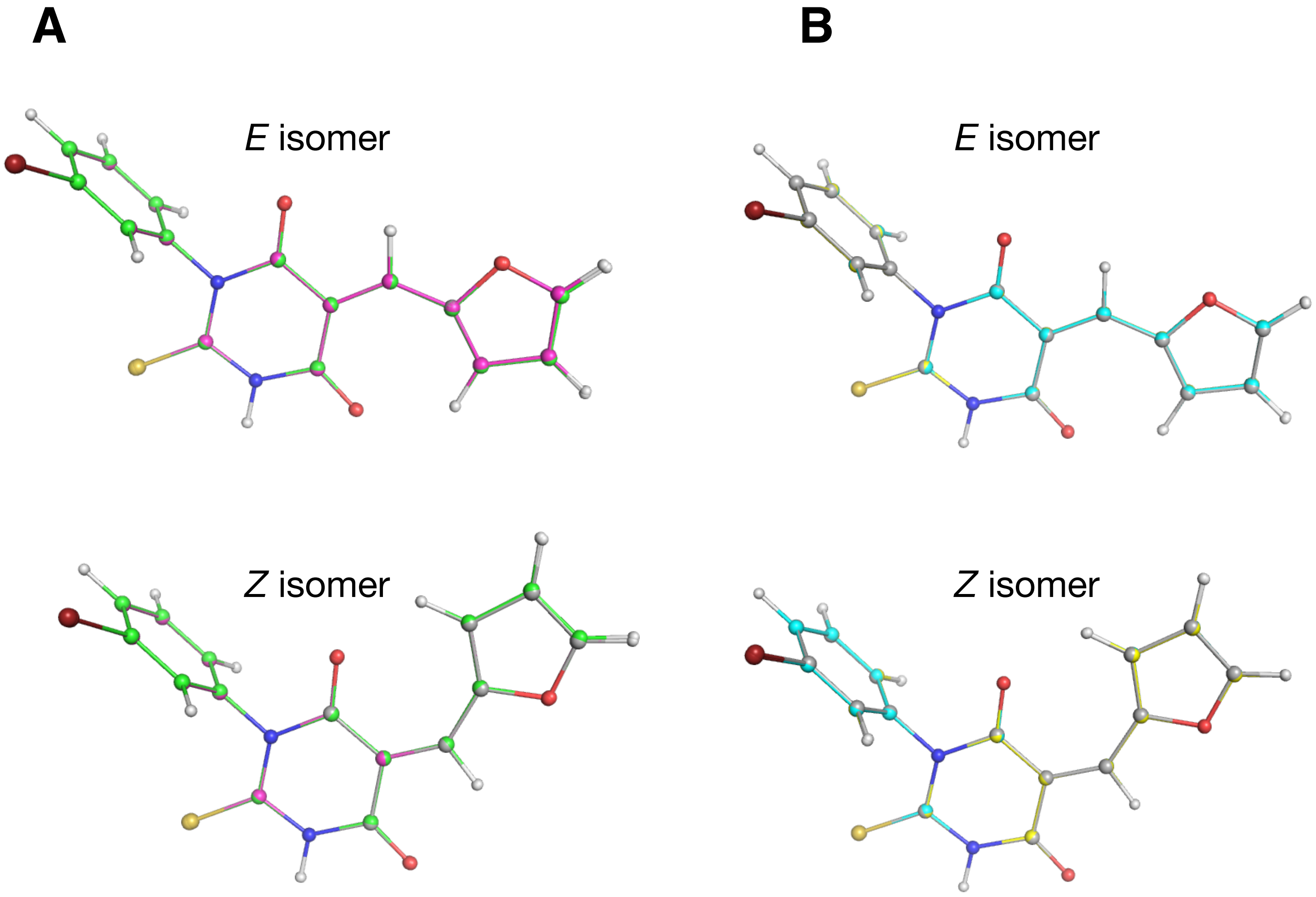

**Figure S3. Optimized geometries of SMIFH2 using different basis sets and solvents.** (A) SMIFH2 was geometry-optimized in vacuum using (U)B3LYP and three different basis sets: 6-311G (green carbons), 6-311+G(d,p) (gray carbons), or aug-cc-pVTZ (magenta carbons). (B) SMIFH2 was geometry-optimized using (U)B3LYP/6-311+G(d,p) in vacuum (gray carbons), DMSO (yellow carbons), or water (cyan carbons). Structures were overlayed in PyMOL using the thiourea portion (N1-C2-N3) as in Figure 5A.

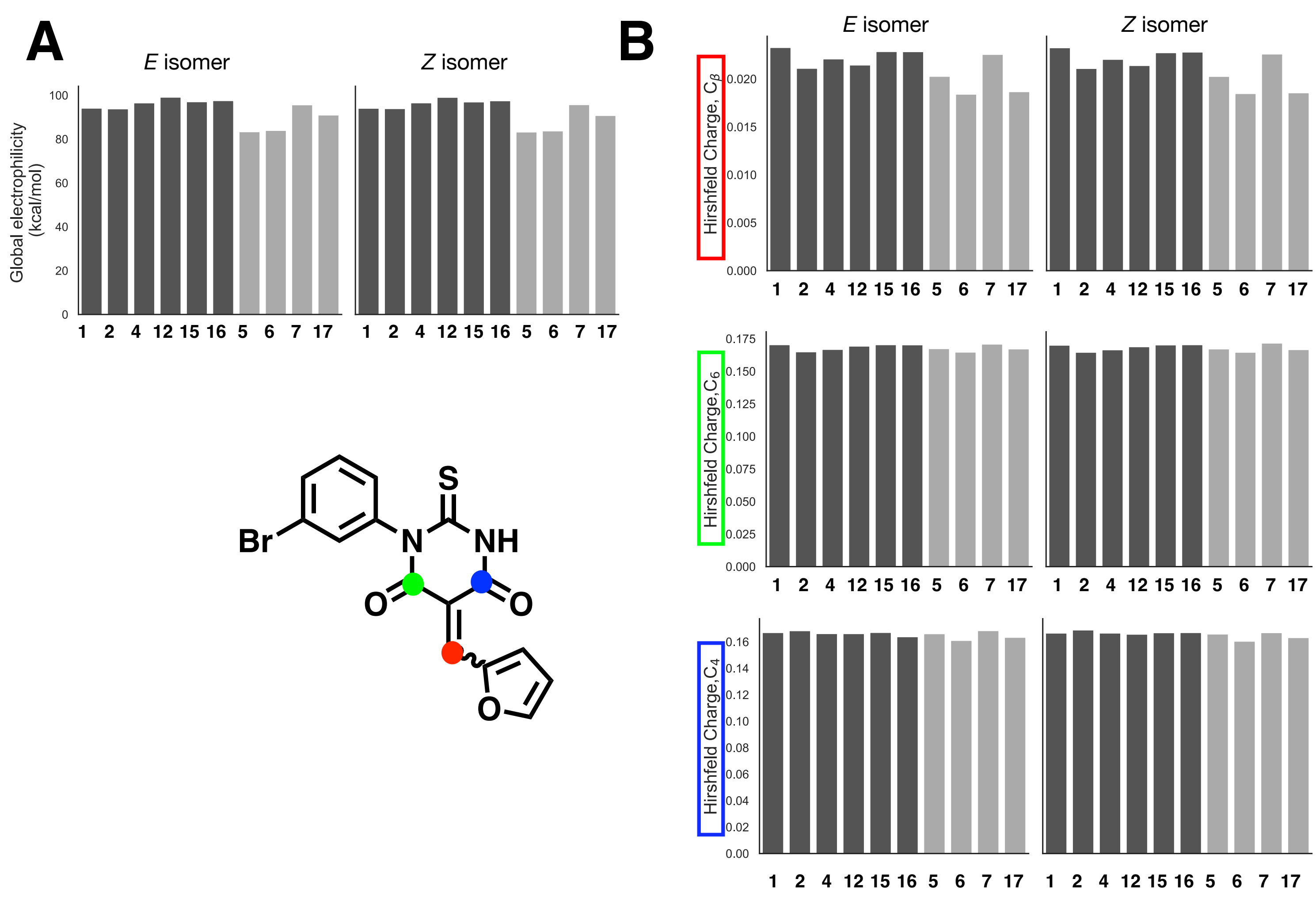

**Figure S4. Electrophilicity and charges for SMIFH2 analogs (related to Figure 5).** (A) Global electrophilicity values corresponding to Figure 5B. (B) Hirshfeld charges for differences plotted in Figure 5D. The specific carbons are indicated by the dots in the structure. All energies and charges were calculated with Gaussian (see Methods for levels of theory and basis sets).

**Figure S5. NMR spectra for SMIFH2 and its derivatives**

Current Data Parameters
NAME I-MAL-78
EXPNO 1
PROCNO 1
Date_ 20200211
Time 15.31 h
INSTRUM Avance Neo Nanobay
PROBHD Z163739_0044 (
PULPROG zg30
TD 65536
SOLVENT DMSO
NS 32
DS 2
SWH 8196.722 Hz
FIDRES 0.250144 Hz
AQ 3.9976959 sec
RG 101
DW 61.000 usec
DE 13.54 usec
TE 296.7 K
D1 1.00000000 sec
TD0 1
SFO1 400.2724717 MHz
NUC1 1H
P0 3.33 usec
P1 10.00 usec
PLW1 14.49800014 W
SI 65536
SF 400.2700030 MHz
WDW EM
SSB 0
LB 0.30 Hz
GB 0
PC 1.00

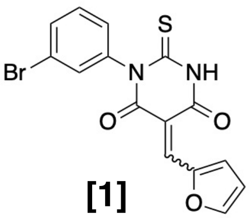

**[1]**

$^1$H NMR (400 MHz, DMSO-$d_6$)

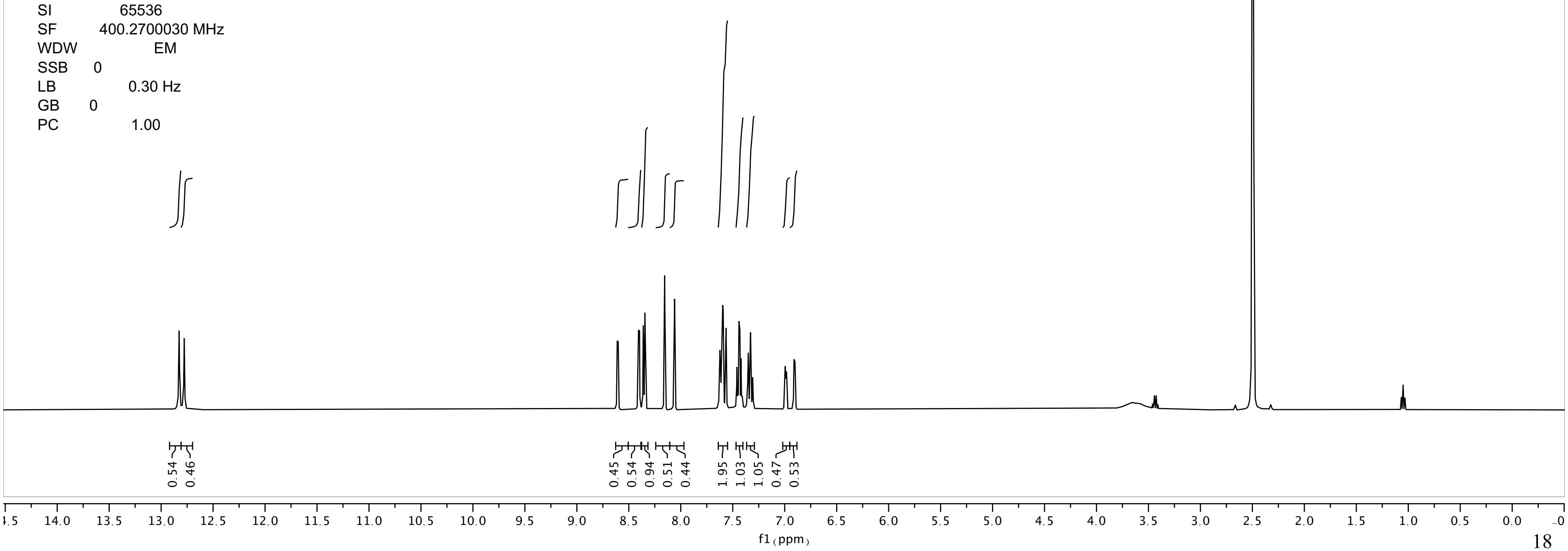

I-MAL-78

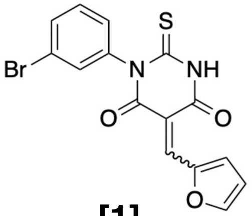

**[1]**

$^{1}$H NMR (400 MHz, DMSO-$d_6$)

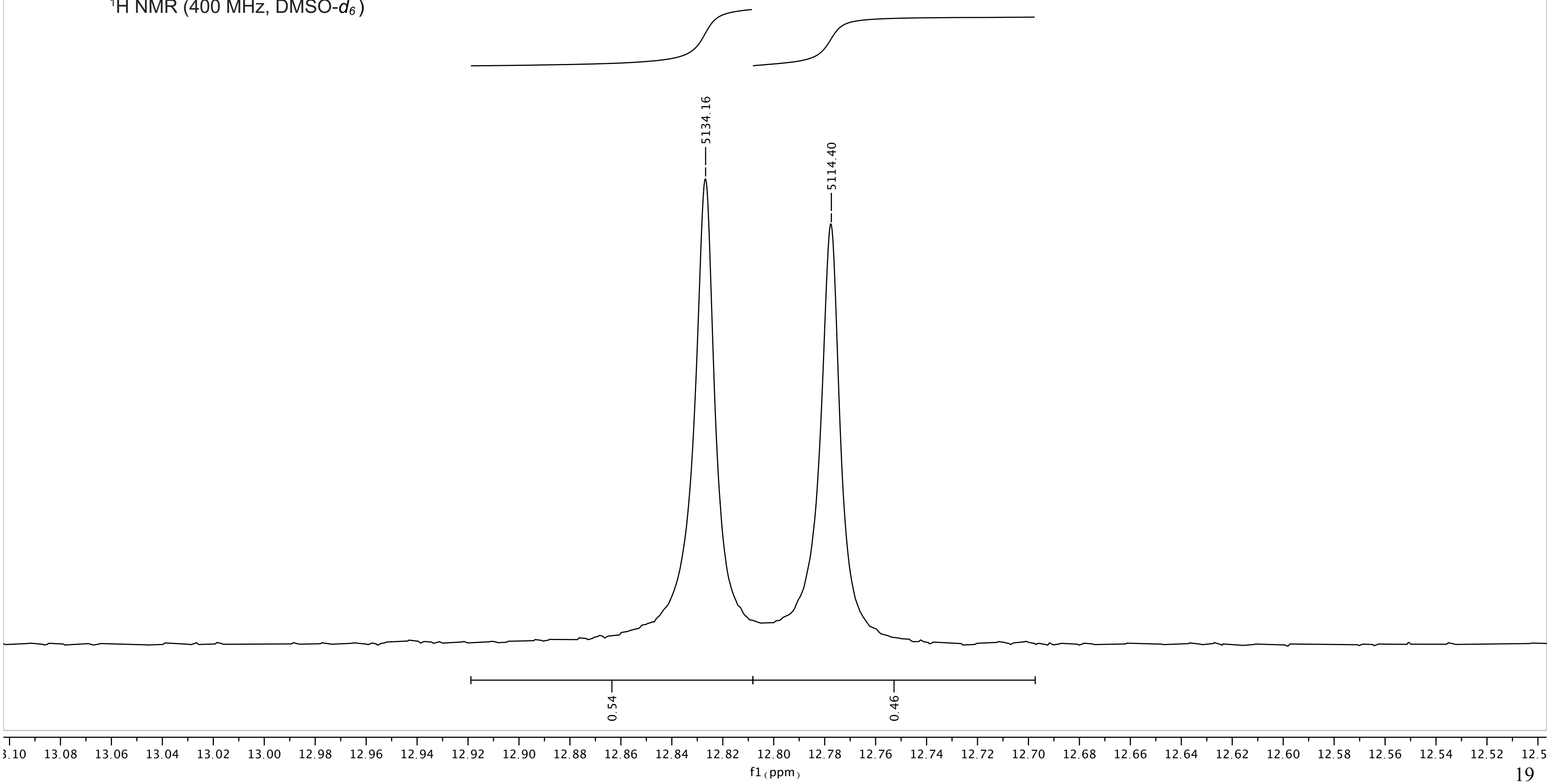

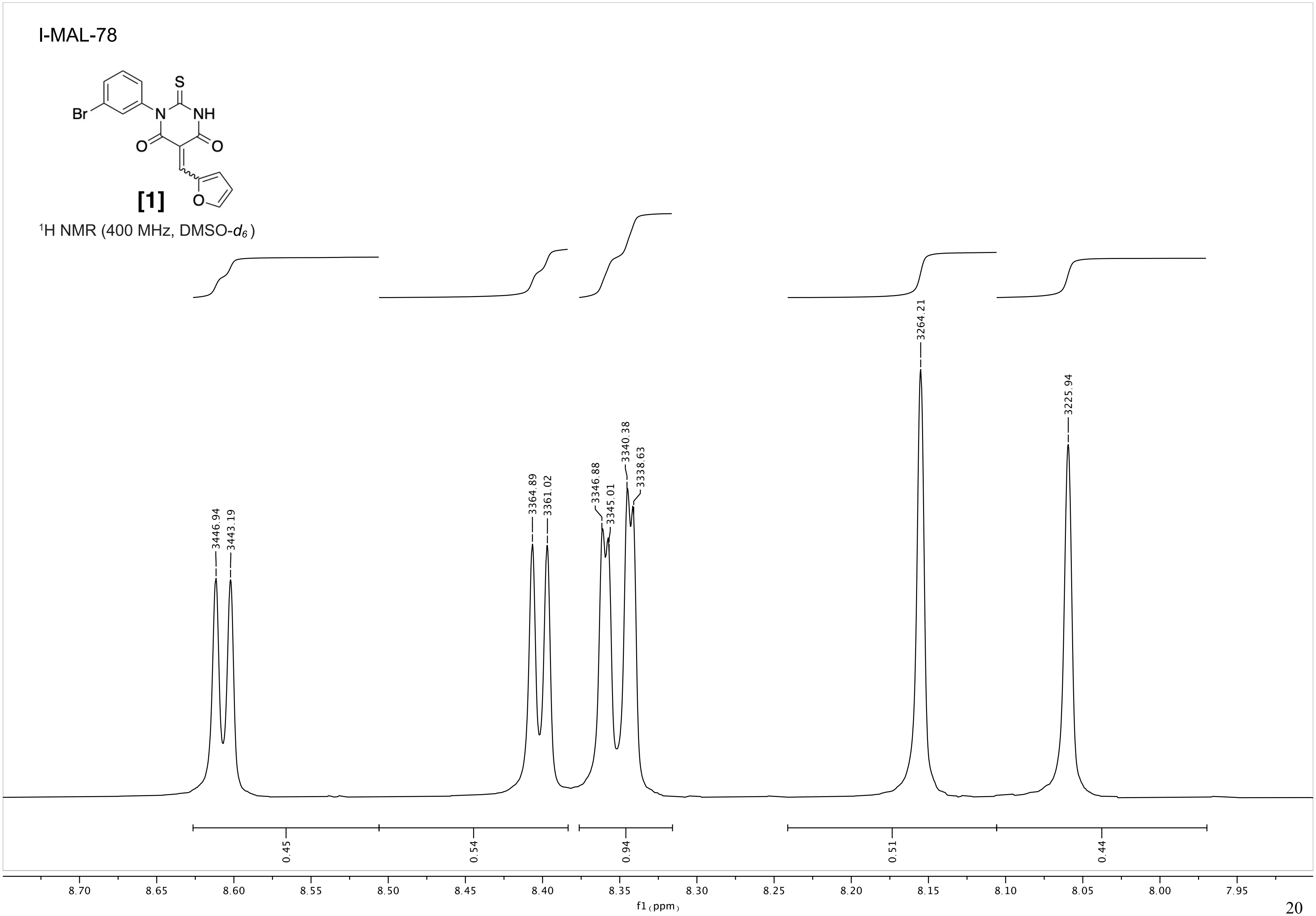
I-MAL-78
Br
N
S
NH
O
O
O
[1]
$^1$H NMR (400 MHz, DMSO-$d_6$)
3446.94
3443.19
3364.89
3361.02
3346.88
3345.01
3340.38
3338.63
3264.21
3225.94
0.45
0.54
0.94
0.51
0.44
8.70
8.65
8.60
8.55
8.50
8.45
8.40
8.35
8.30
8.25
8.20
8.15
8.10
8.05
8.00
7.95
f1 (ppm)

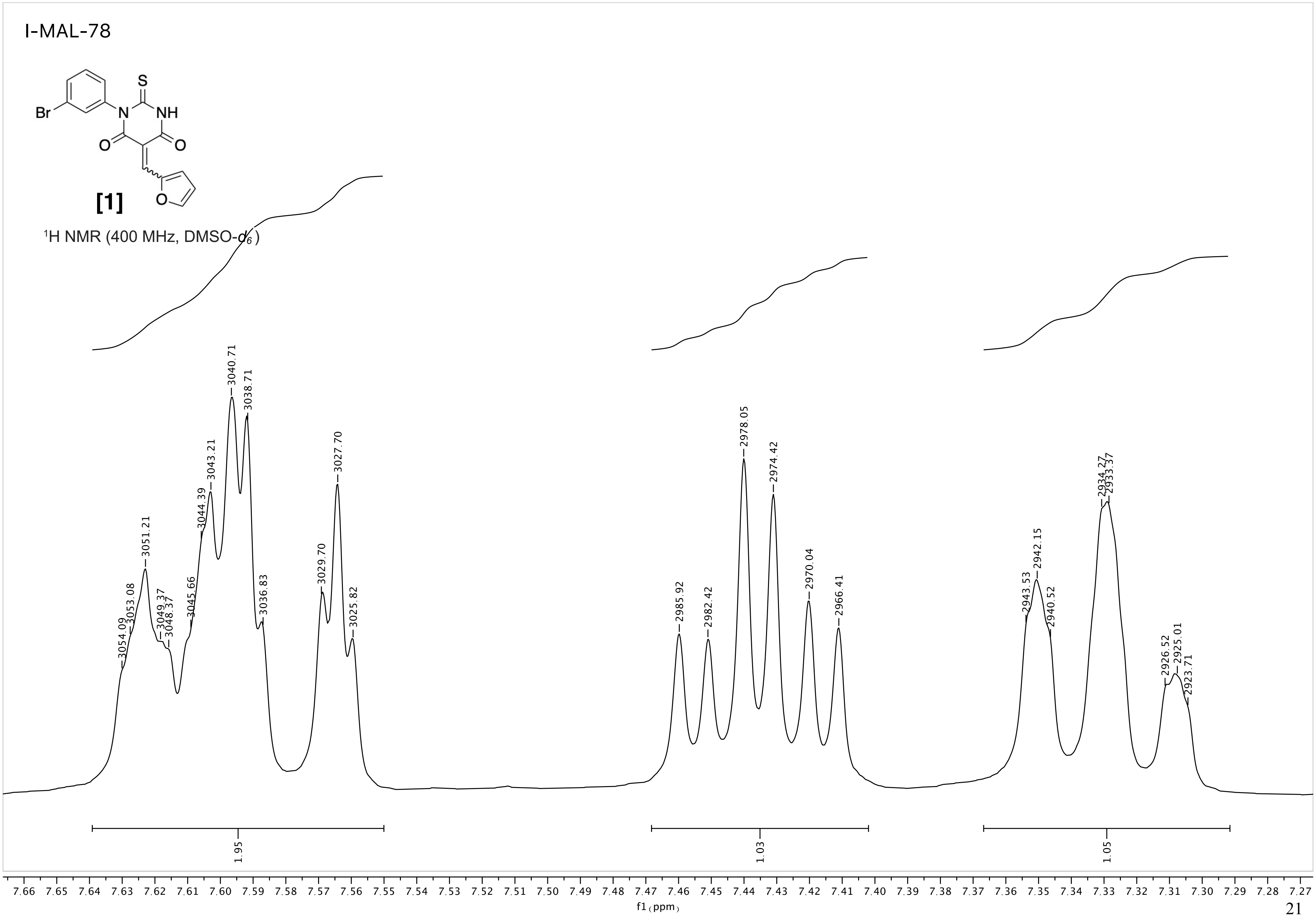
I-MAL-78
Br
N
NH
S
O
O
O
[1]
$^{1}$H NMR (400 MHz, DMSO-$d_6$)
3054.09
3053.08
3051.21
3049.37
3048.37
3045.66
3044.39
3043.21
3040.71
3038.71
3036.83
3029.70
3027.70
3025.82
2985.92
2982.42
2978.05
2974.42
2970.04
2966.41
2943.53
2942.15
2940.52
2934.27
2933.37
2926.52
2925.01
2923.71
1.95
1.03
1.05
7.66 7.65 7.64 7.63 7.62 7.61 7.60 7.59 7.58 7.57 7.56 7.55 7.54 7.53 7.52 7.51 7.50 7.49 7.48 7.47 7.46 7.45 7.44 7.43 7.42 7.41 7.40 7.39 7.38 7.37 7.36 7.35 7.34 7.33 7.32 7.31 7.30 7.29 7.28 7.27
f1 (ppm)

I-MAL-78

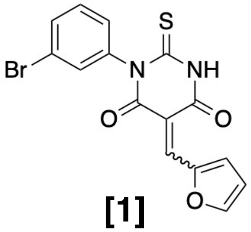

[1]

$^{1}$H NMR (400 MHz, DMSO-$d_6$)

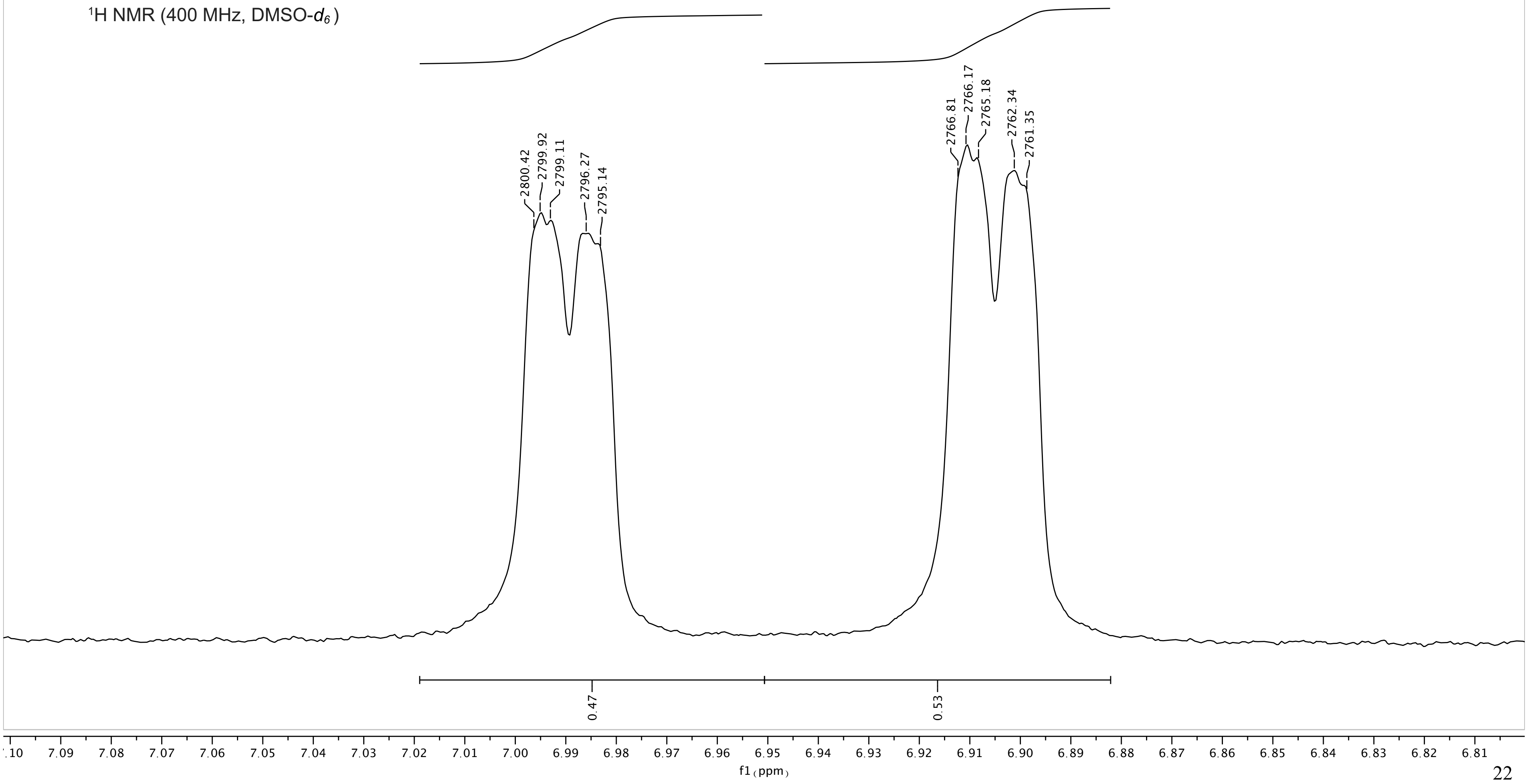

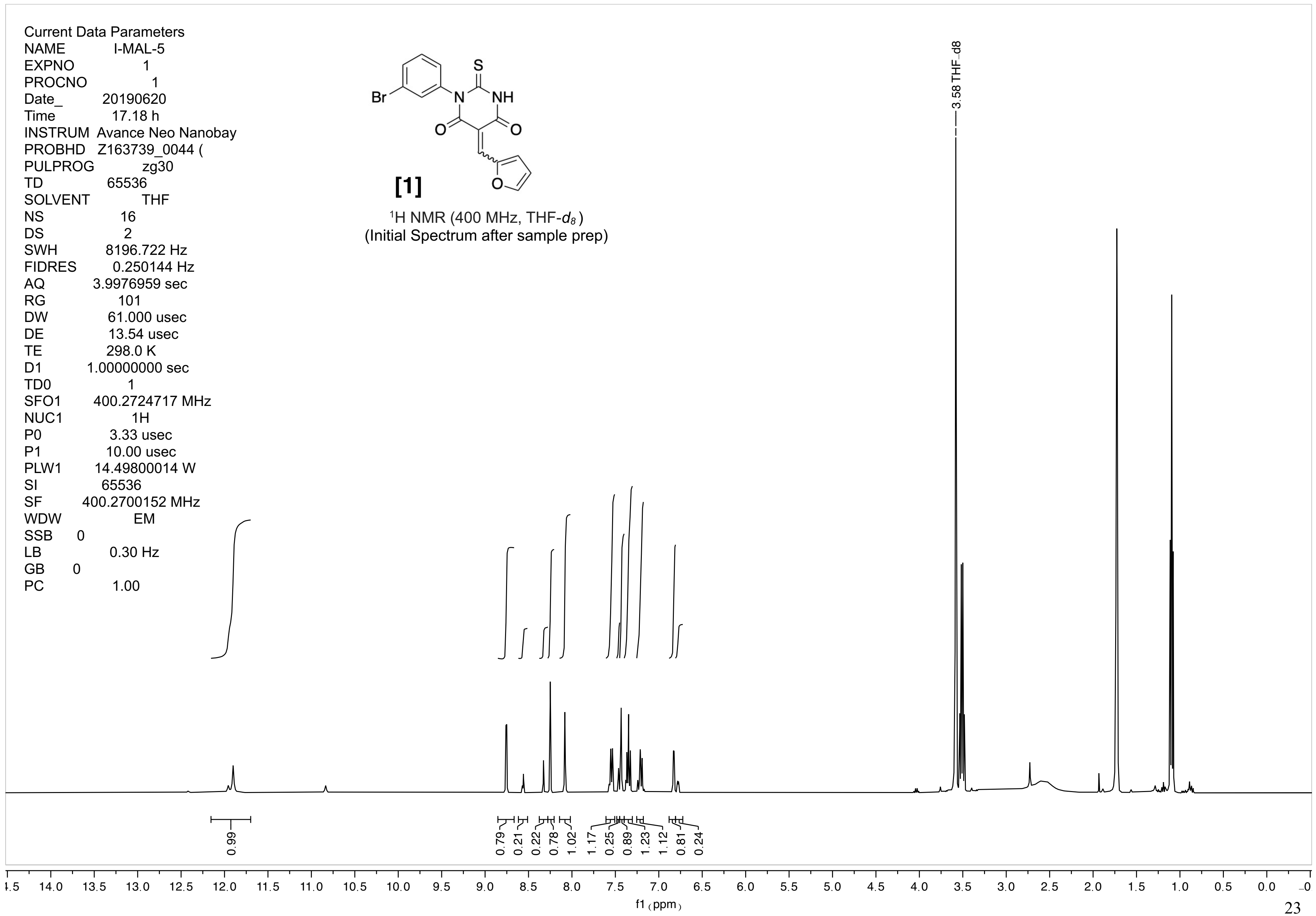
Current Data Parameters
NAME I-MAL-5
EXPNO 1
PROCNO 1
Date_ 20190620
Time 17.18 h
INSTRUM Avance Neo Nanobay
PROBHD Z163739_0044 (
PULPROG zg30
TD 65536
SOLVENT THF
NS 16
DS 2
SWH 8196.722 Hz
FIDRES 0.250144 Hz
AQ 3.9976959 sec
RG 101
DW 61.000 usec
DE 13.54 usec
TE 298.0 K
D1 1.00000000 sec
TD0 1
SFO1 400.2724717 MHz
NUC1 1H
P0 3.33 usec
P1 10.00 usec
PLW1 14.49800014 W
SI 65536
SF 400.2700152 MHz
WDW EM
SSB 0
LB 0.30 Hz
GB 0
PC 1.00
Br
S
N
NH
O
O
O
[1]
1H NMR (400 MHz, THF-d8)
(Initial Spectrum after sample prep)
3.58 THF_d8
0.99
0.79
0.21
0.22
0.78
1.02
1.17
0.25
0.89
1.23
1.12
0.81
0.24
14.0 13.5 13.0 12.5 12.0 11.5 11.0 10.5 10.0 9.5 9.0 8.5 8.0 7.5 7.0 6.5 6.0 5.5 5.0 4.5 4.0 3.5 3.0 2.5 2.0 1.5 1.0 0.5 0.0
f1 (ppm)

1-MAL-5

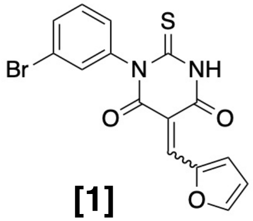

[1]

$^{1}$H NMR (400 MHz, THF-$d_8$)
(Initial Spectrum after sample prep.)

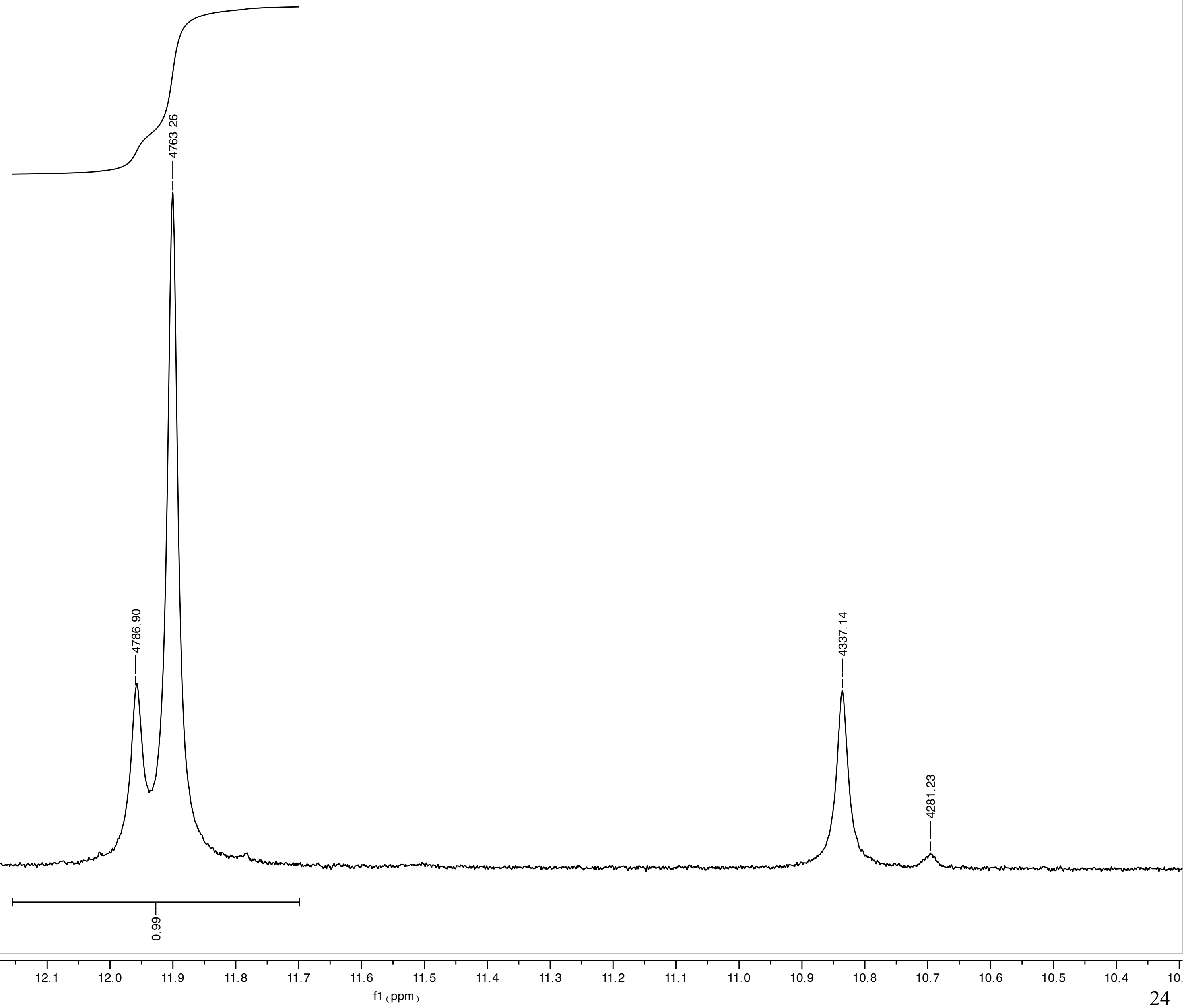

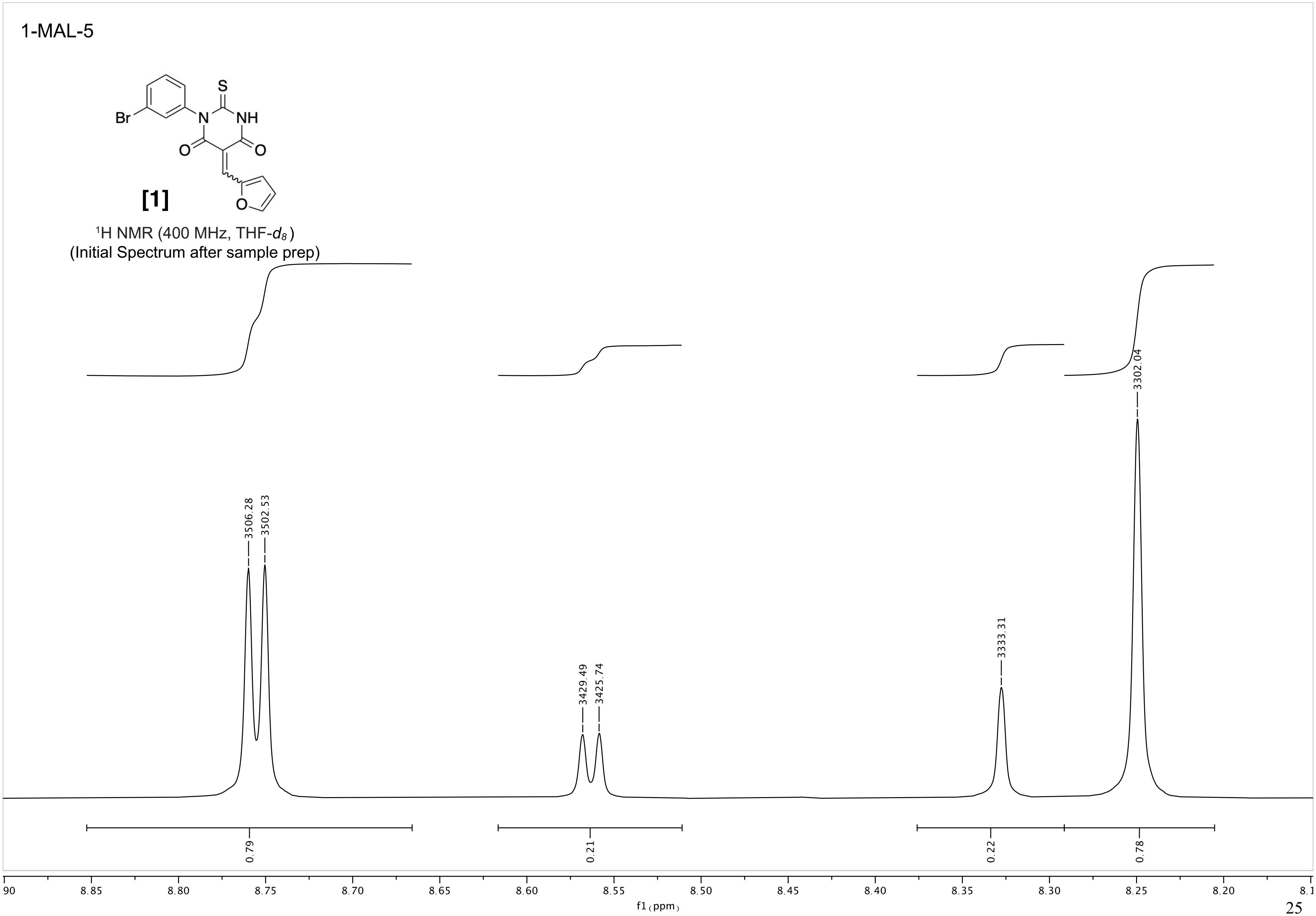
1-MAL-5
S
NH
Br
N
O
O
O
[1]
$^1$H NMR (400 MHz, THF-$d_8$)
(Initial Spectrum after sample prep)
3506.28
3502.53
3429.49
3425.74
3333.31
3302.04
0.79
0.21
0.22
0.78
f1 (ppm)

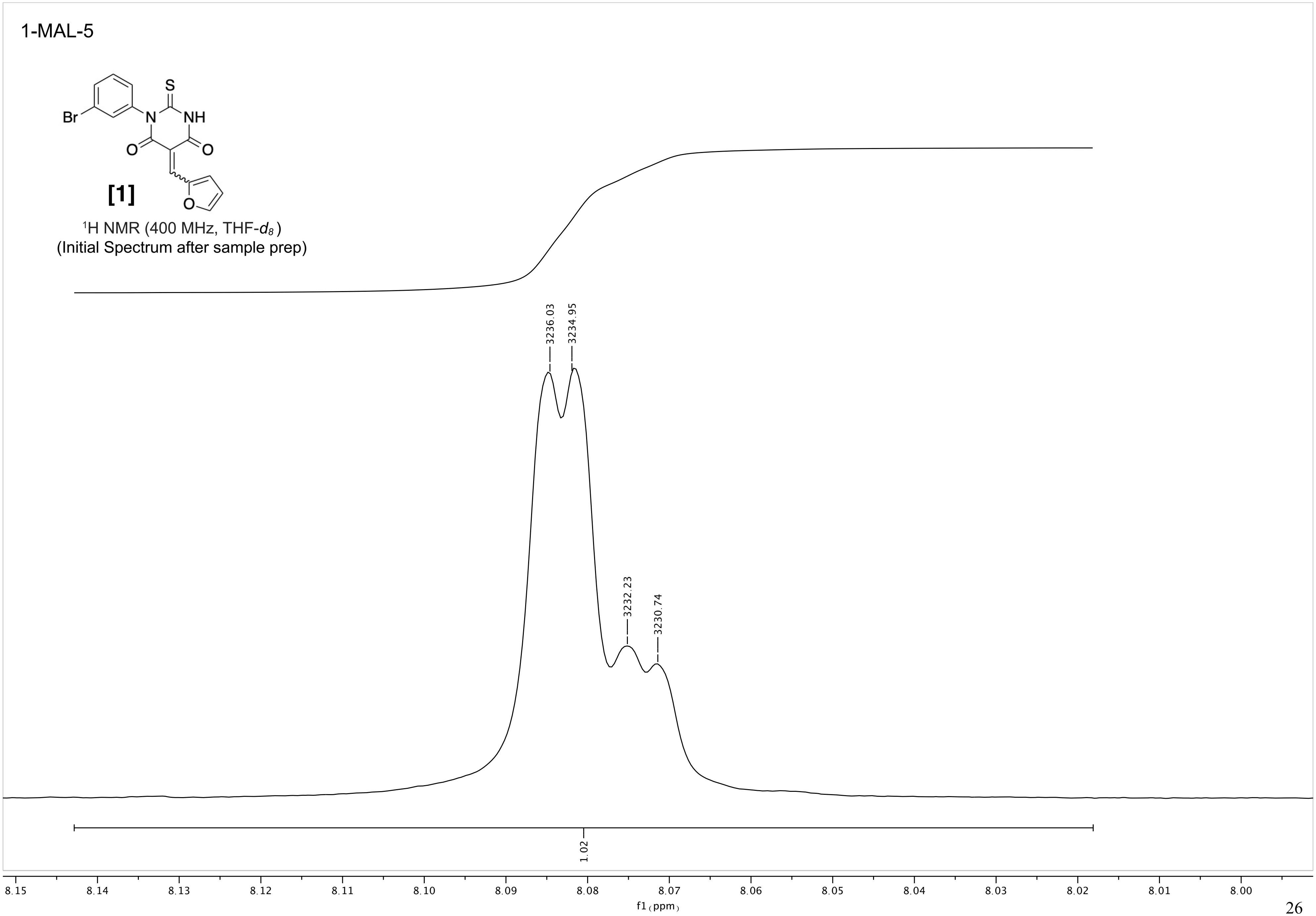
1-MAL-5
[1]
$^1$H NMR (400 MHz, THF-$d_8$)
(Initial Spectrum after sample prep)
3236.03
3234.95
3232.23
3230.74
1.02
f1 (ppm)

1-MAL-5

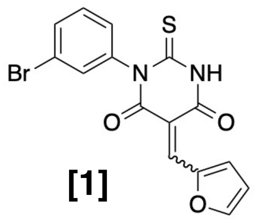

[1]

$^{1}$H NMR (400 MHz, THF-$d_8$)
(Initial Spectrum after sample prep)

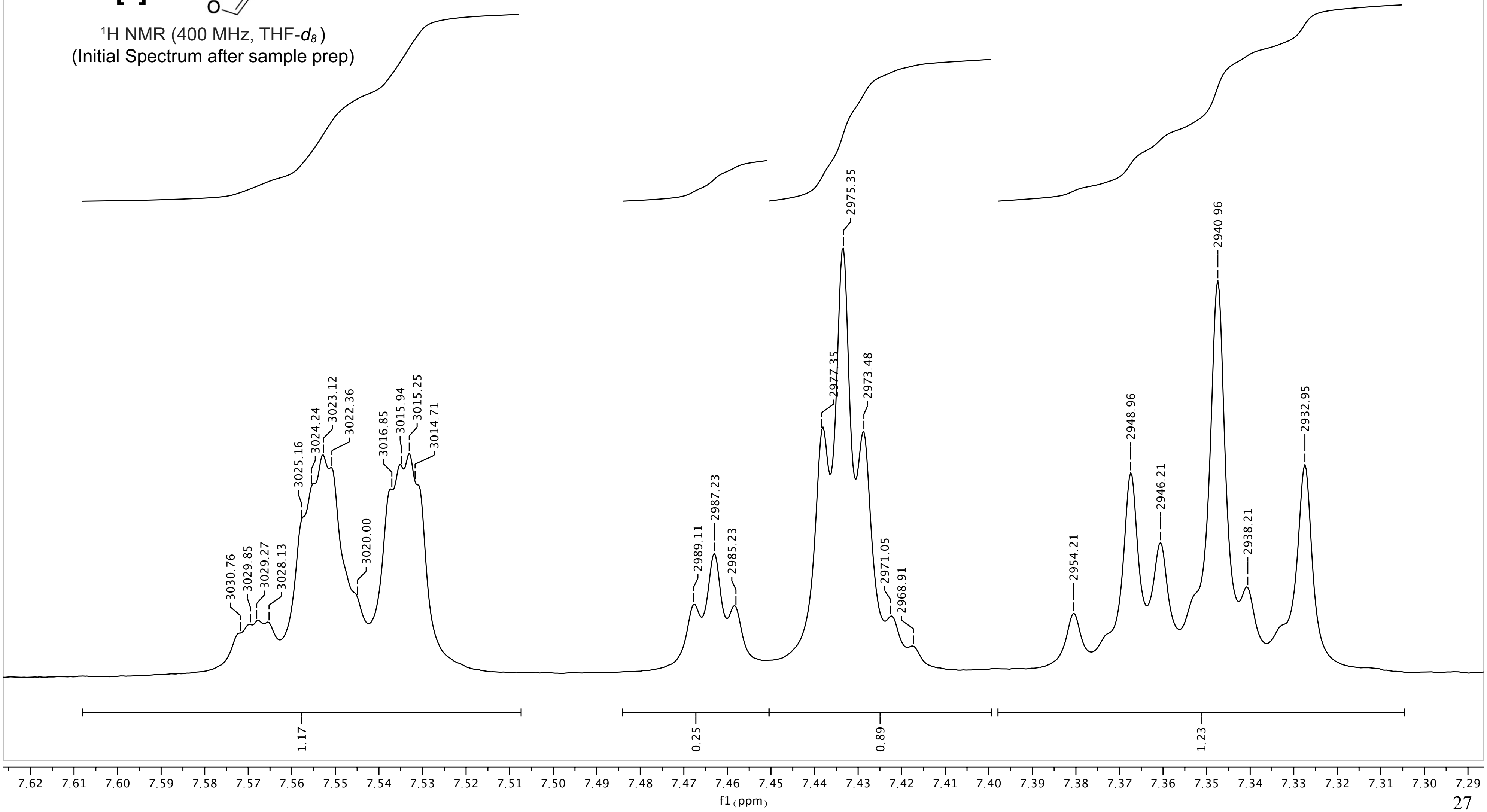

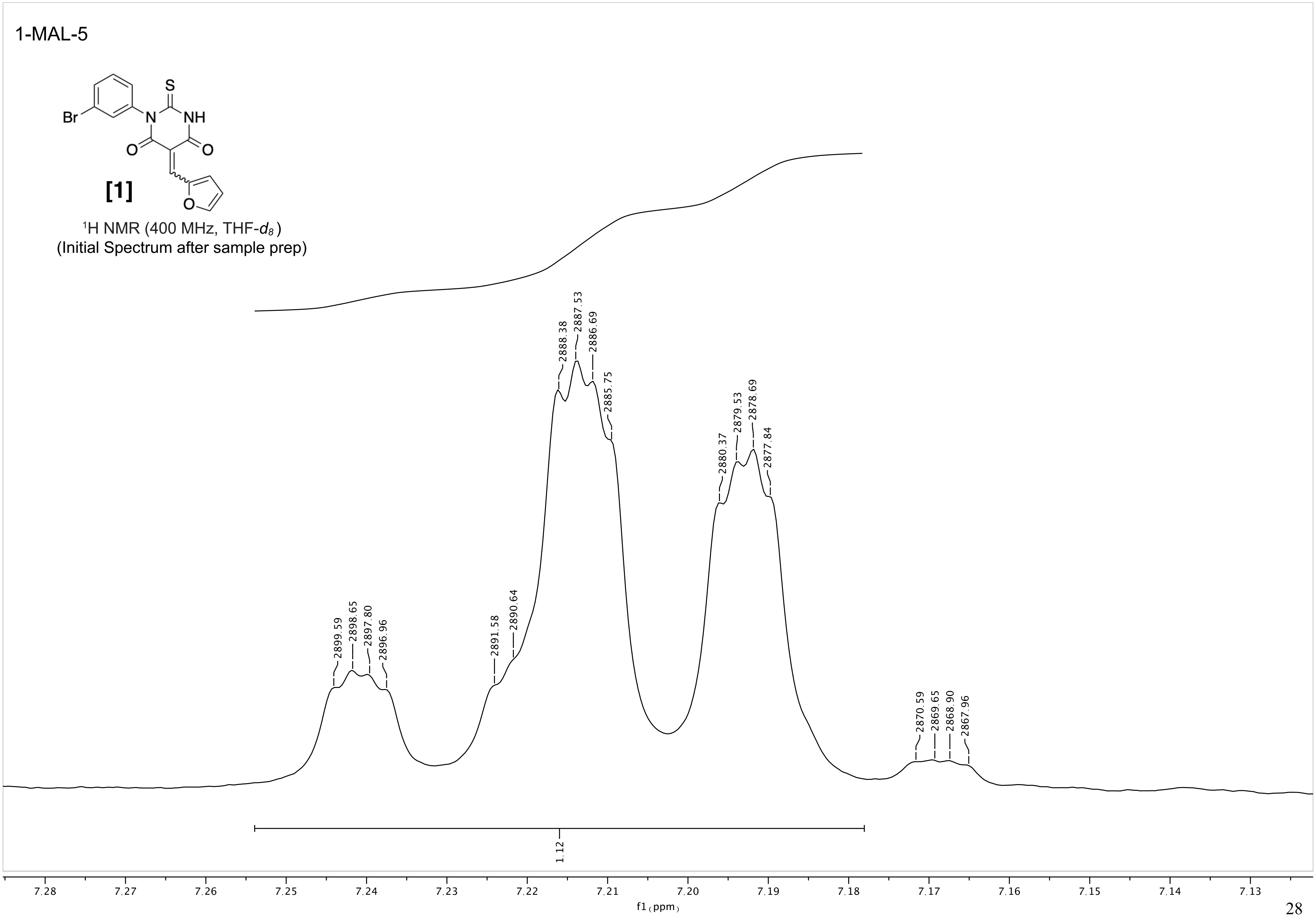
1-MAL-5
Br
S
NH
N
O
O
O
[1]
$^1$H NMR (400 MHz, THF-$d_8$)
(Initial Spectrum after sample prep)
2899.59
2898.65
2897.80
2896.96
2891.58
2890.64
2888.38
2887.53
2886.69
2885.75
2880.37
2879.53
2878.69
2877.84
2870.59
2869.65
2868.90
2867.96
1.12
7.28
7.27
7.26
7.25
7.24
7.23
7.22
7.21
7.20
7.19
7.18
7.17
7.16
7.15
7.14
7.13
f1 (ppm)

1-MAL-5

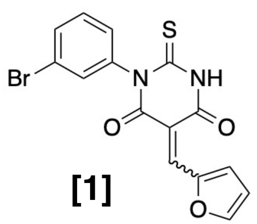

**[1]**

$^{1}$H NMR (400 MHz, THF-$d_8$)
(Initial Spectrum after sample prep)

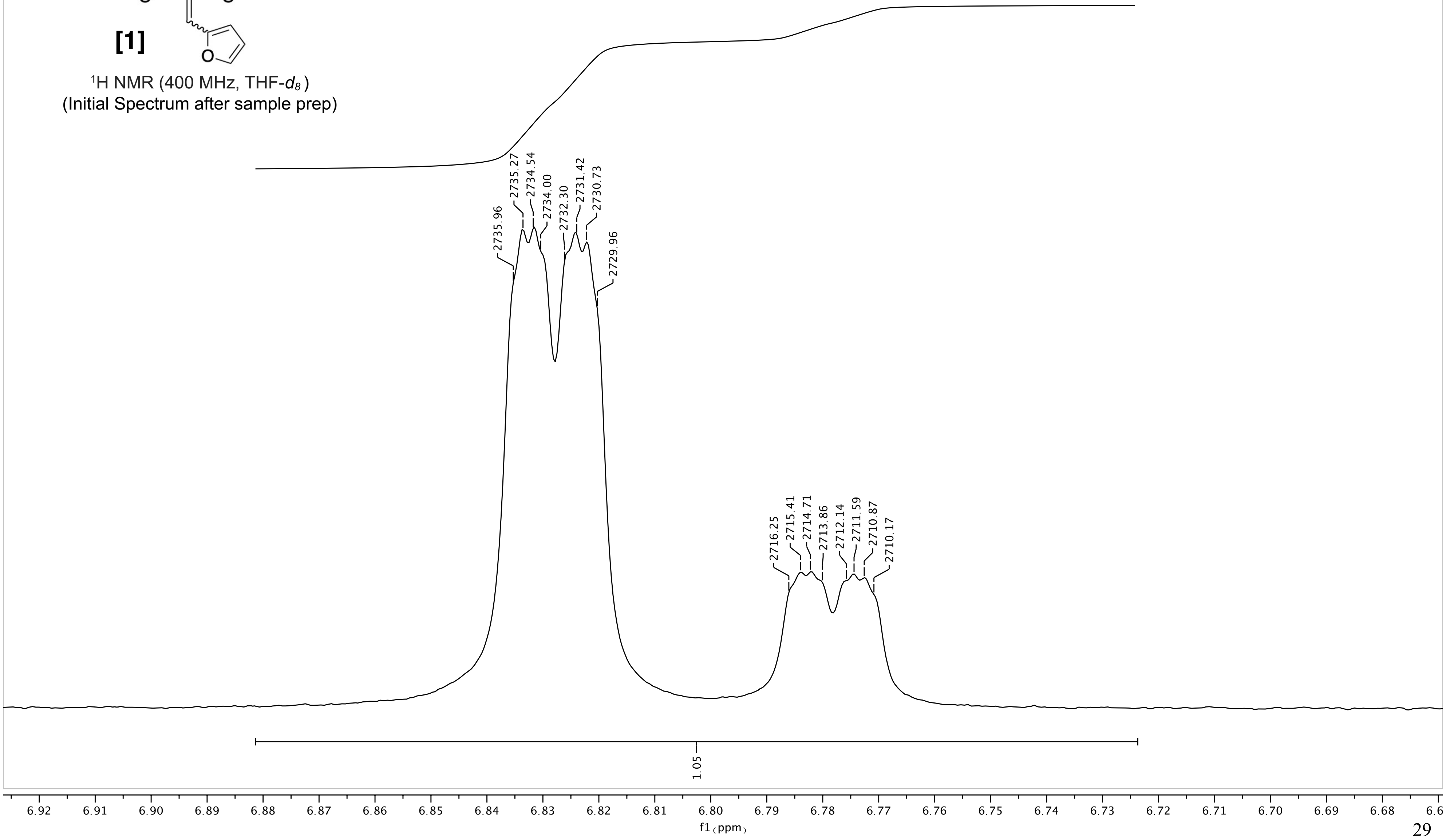

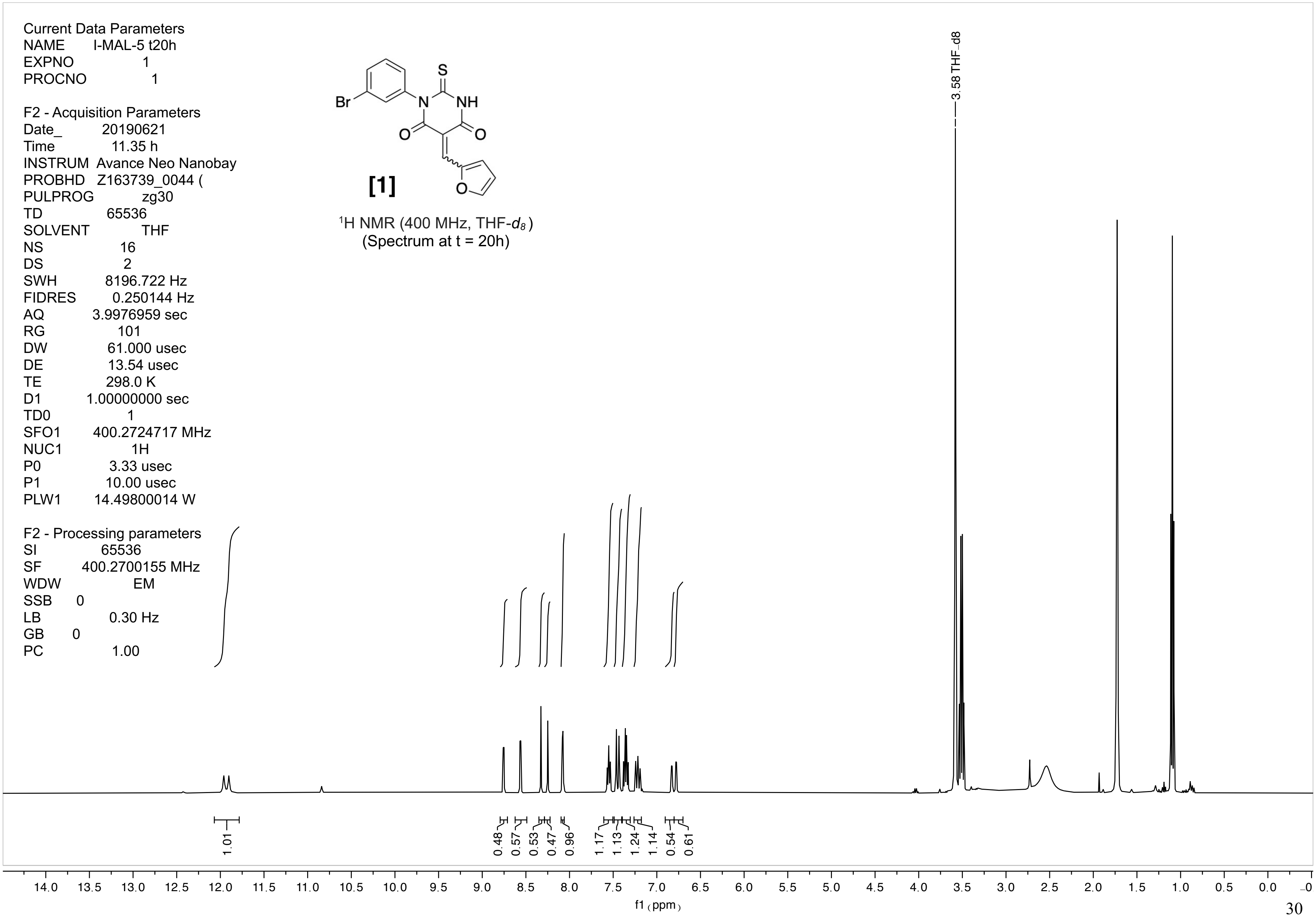
Current Data Parameters
NAME I-MAL-5 t20h
EXPNO 1
PROCNO 1
F2 - Acquisition Parameters
Date_ 20190621
Time 11.35 h
INSTRUM Avance Neo Nanobay
PROBHD Z163739_0044 (
PULPROG zg30
TD 65536
SOLVENT THF
NS 16
DS 2
SWH 8196.722 Hz
FIDRES 0.250144 Hz
AQ 3.9976959 sec
RG 101
DW 61.000 usec
DE 13.54 usec
TE 298.0 K
D1 1.00000000 sec
TD0 1
SFO1 400.2724717 MHz
NUC1 1H
P0 3.33 usec
P1 10.00 usec
PLW1 14.49800014 W
F2 - Processing parameters
SI 65536
SF 400.2700155 MHz
WDW EM
SSB 0
LB 0.30 Hz
GB 0
PC 1.00
[1]
1H NMR (400 MHz, THF-d8)
(Spectrum at t = 20h)
3.58 THF_d8
1.01
0.48
0.57
0.53
0.47
0.96
1.17
1.13
1.24
1.14
0.54
0.61
f1 (ppm)

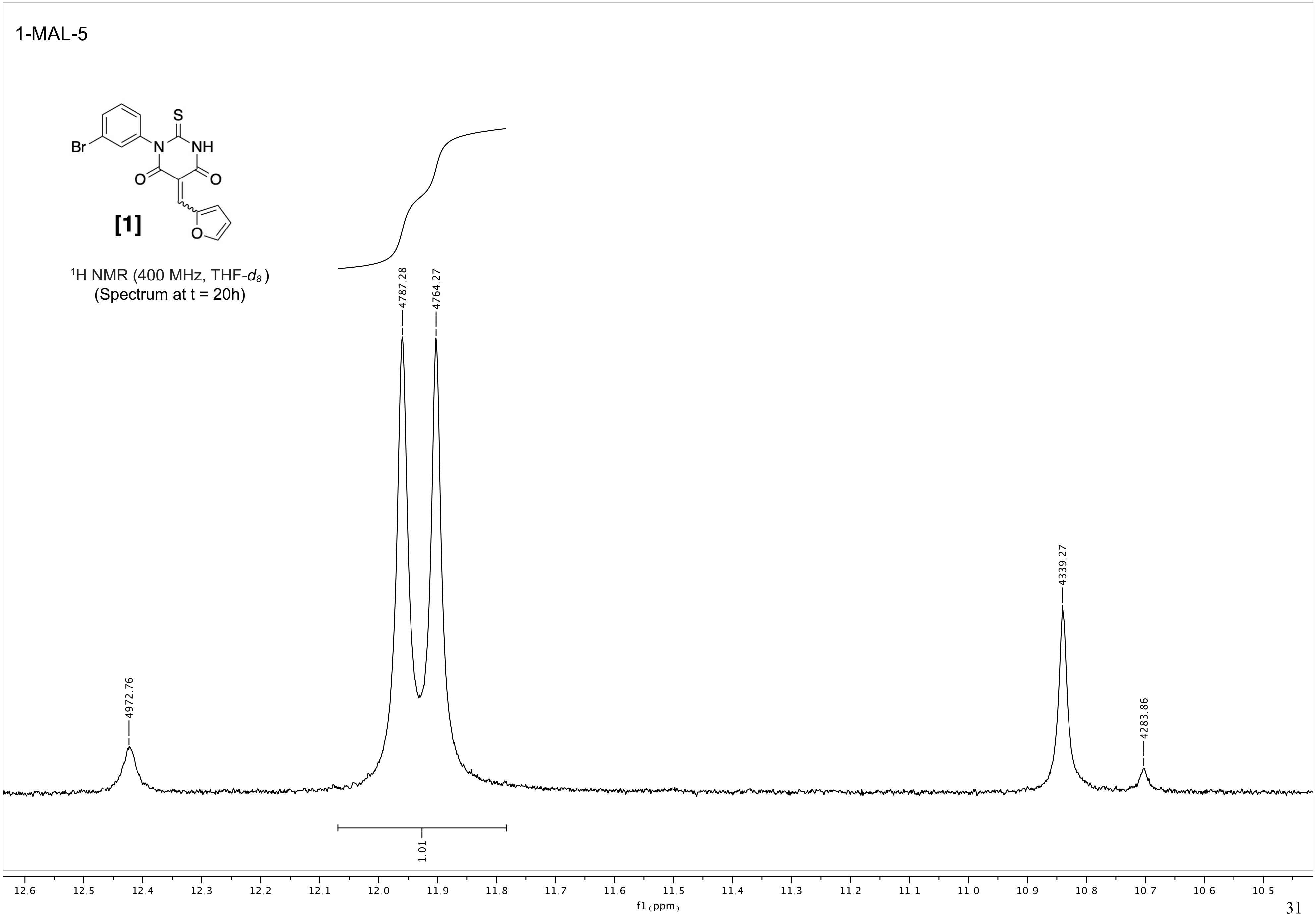
1-MAL-5
S
NH
N
Br
O
O
O
[1]
$^1$H NMR (400 MHz, THF-$d_8$)
(Spectrum at t = 20h)
4972.76
4787.28
4764.27
4339.27
4283.86
1.01
12.6
12.5
12.4
12.3
12.2
12.1
12.0
11.9
11.8
11.7
11.6
11.5
11.4
11.3
11.2
11.1
11.0
10.9
10.8
10.7
10.6
10.5
f1 (ppm)

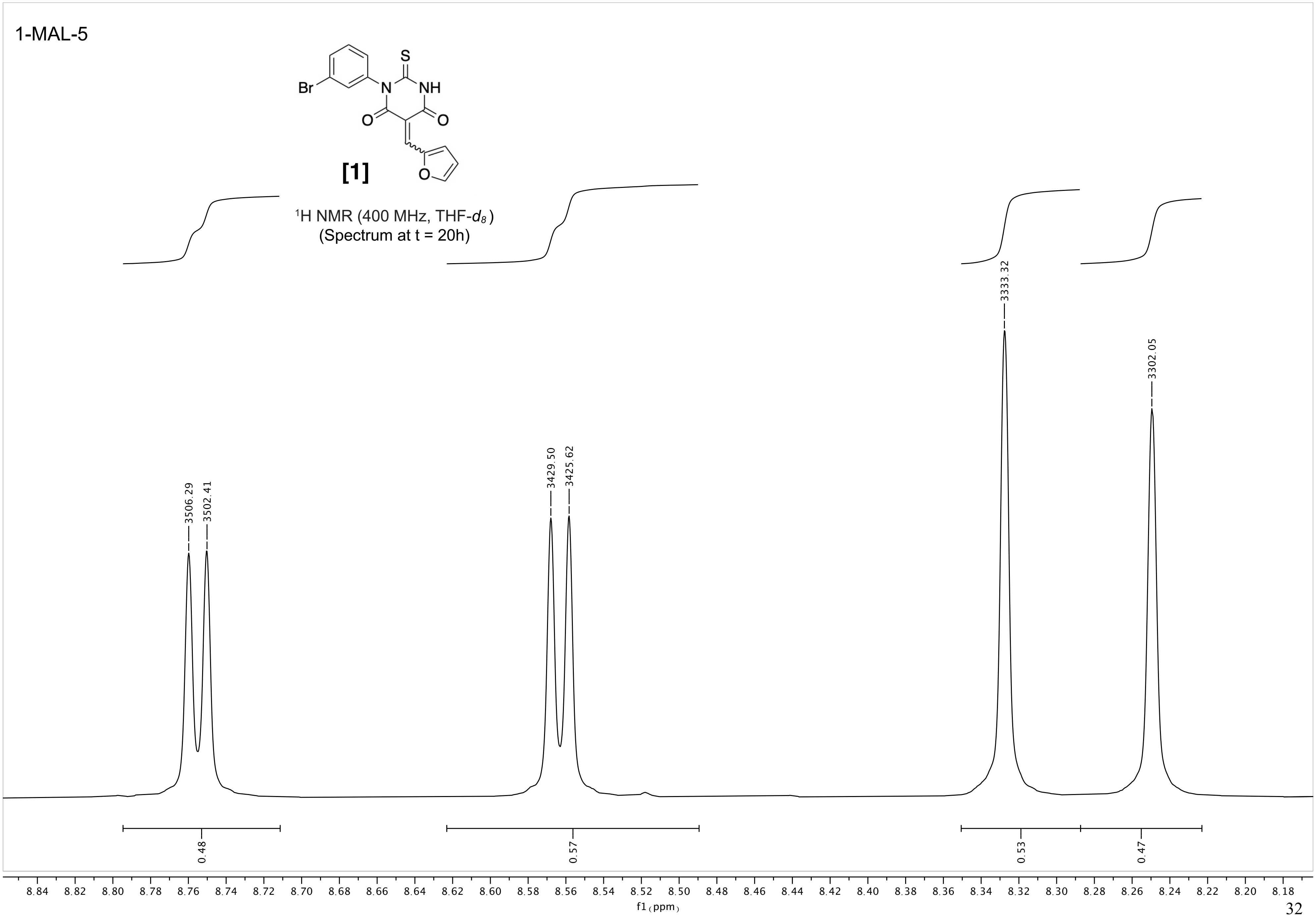
1-MAL-5
Br
S
N
NH
O
O
O
[1]
1H NMR (400 MHz, THF-d8)
(Spectrum at t = 20h)
3506.29
3502.41
3429.50
3425.62
3333.32
3302.05
0.48
0.57
0.53
0.47
8.84 8.82 8.80 8.78 8.76 8.74 8.72 8.70 8.68 8.66 8.64 8.62 8.60 8.58 8.56 8.54 8.52 8.50 8.48 8.46 8.44 8.42 8.40 8.38 8.36 8.34 8.32 8.30 8.28 8.26 8.24 8.22 8.20 8.18
f1 (ppm)

1-MAL-5

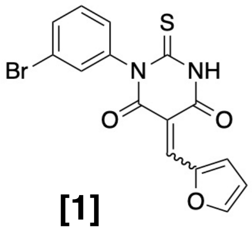

**[1]**

$^1$H NMR (400 MHz, THF-$d_8$)
(Spectrum at t = 20h)

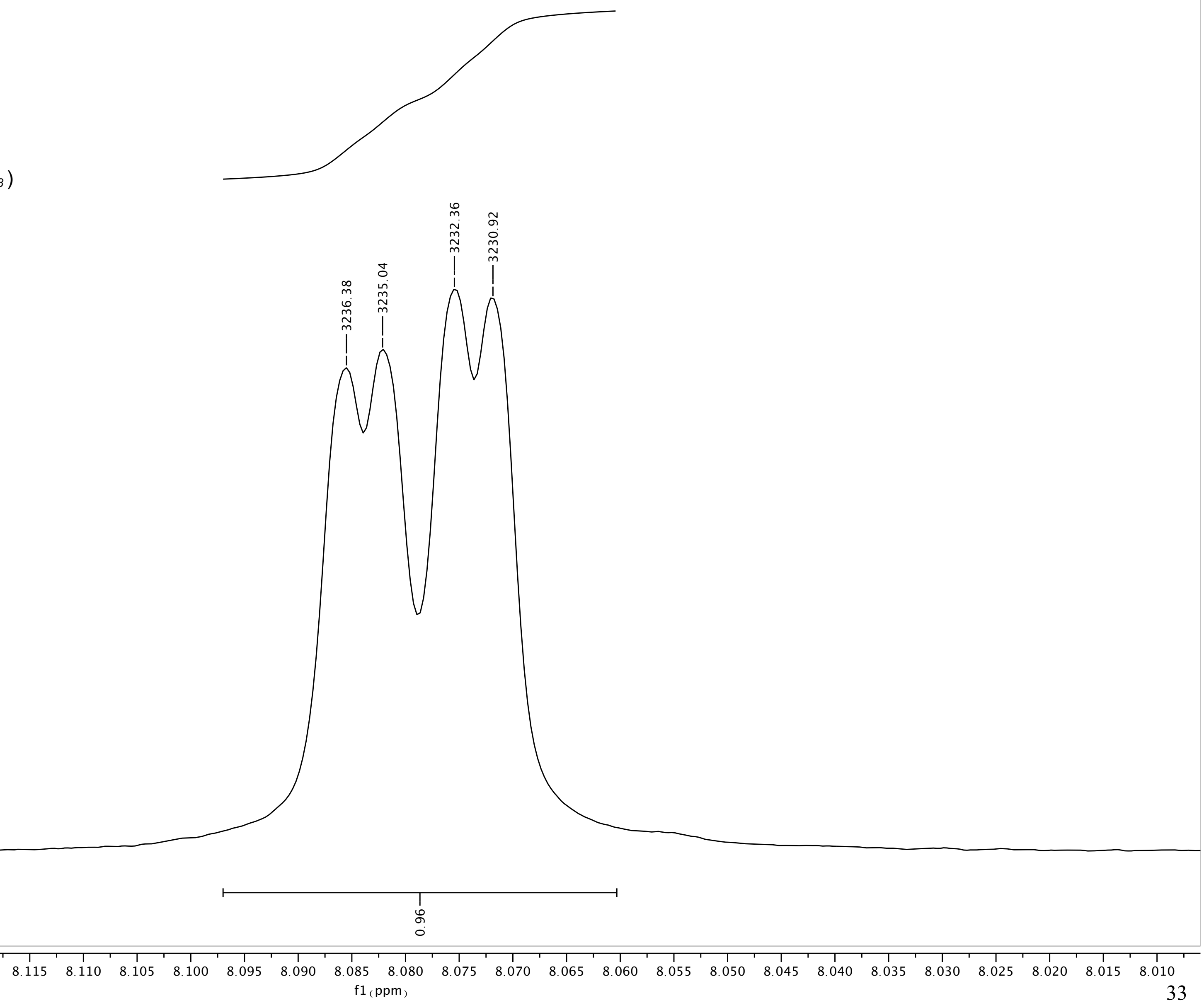

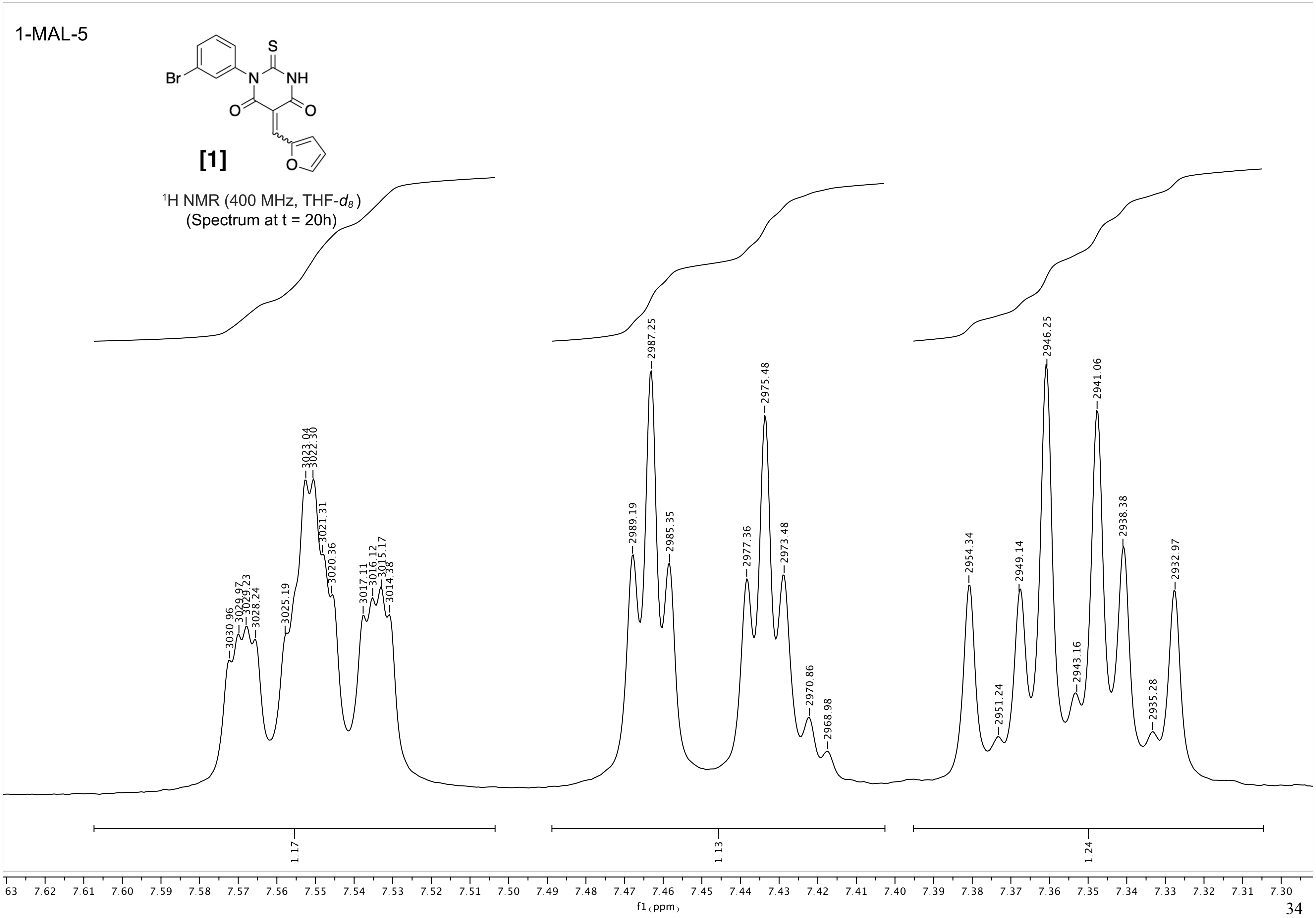
1-MAL-5
[1]
1H NMR (400 MHz, THF-d8)
(Spectrum at t = 20h)
f1 (ppm)

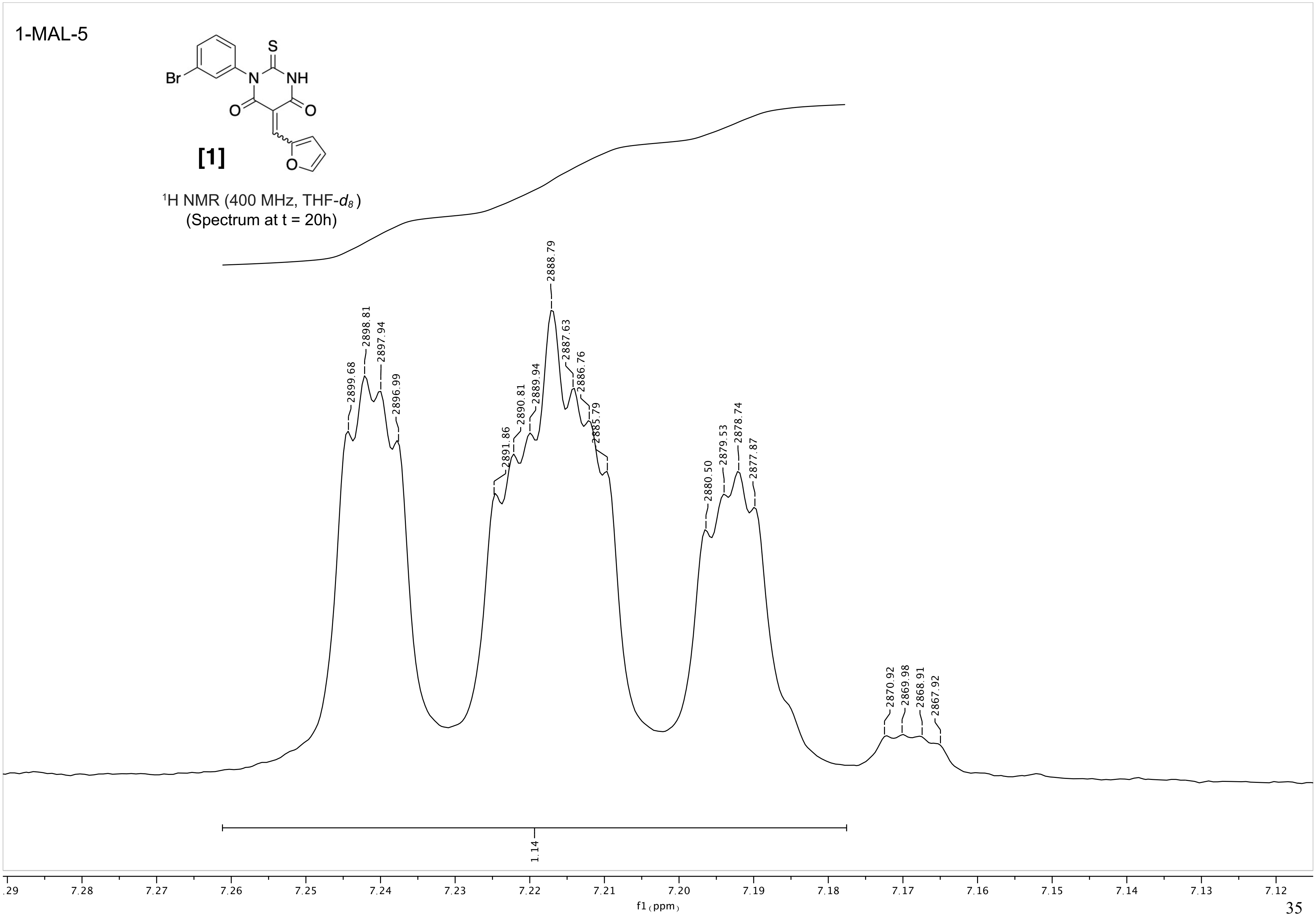
1-MAL-5
Br
N
NH
S
O
O
O
[1]
1H NMR (400 MHz, THF-d8)
(Spectrum at t = 20h)
2899.68
2898.81
2897.94
2896.99
2891.86
2890.81
2889.94
2888.79
2887.63
2886.76
2885.79
2880.50
2879.53
2878.74
2877.87
2870.92
2869.98
2868.91
2867.92
1.14
.29 7.28 7.27 7.26 7.25 7.24 7.23 7.22 7.21 7.20 7.19 7.18 7.17 7.16 7.15 7.14 7.13 7.12
f1 (ppm)

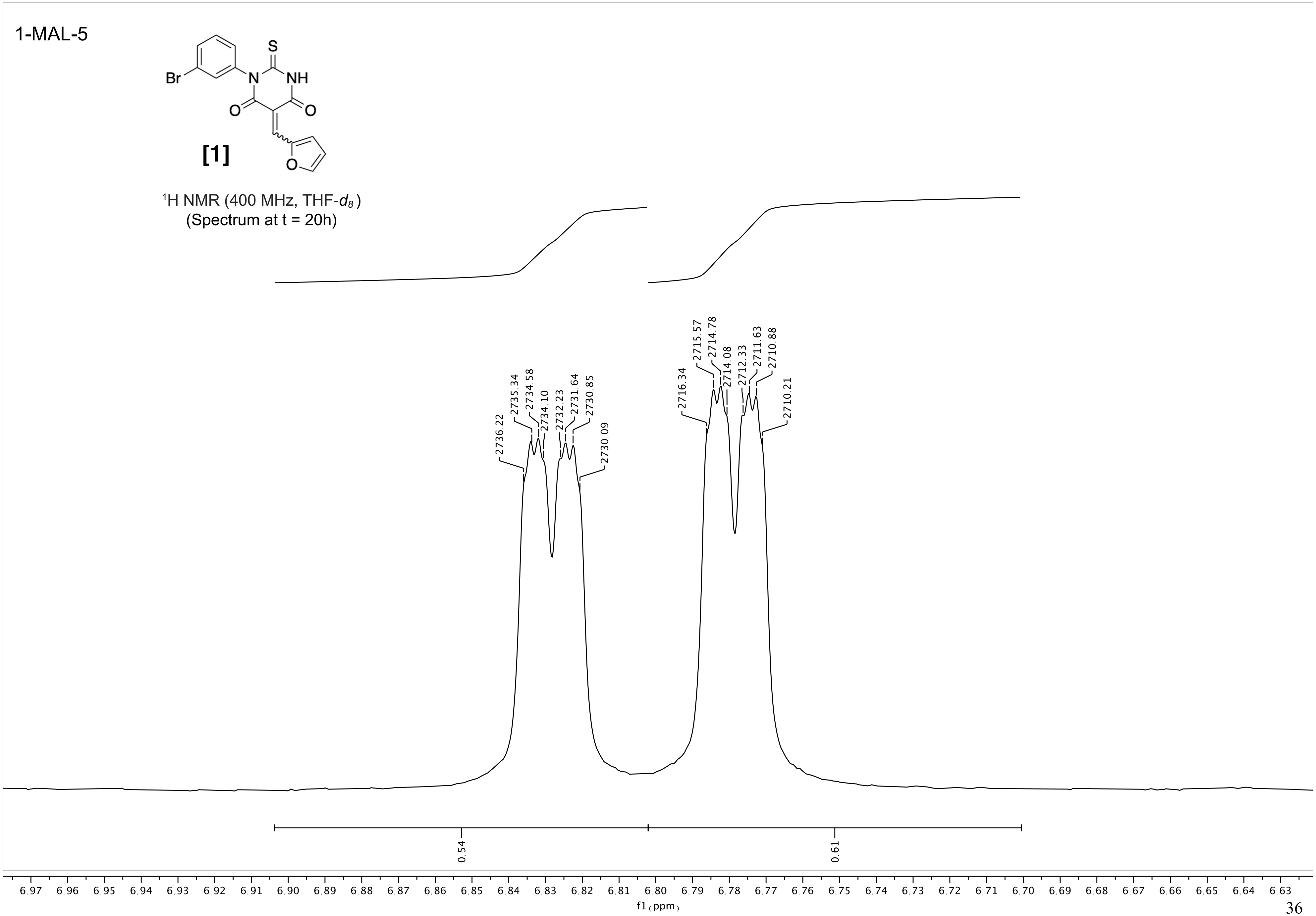
1-MAL-5
Br
N
S
NH
O
O
O
[1]
¹H NMR (400 MHz, THF-d₈)
(Spectrum at t = 20h)
2736.22
2735.34
2734.58
2734.10
2732.23
2731.64
2730.85
2730.09
2716.34
2715.57
2714.78
2714.08
2712.33
2711.63
2710.88
2710.21
0.54
0.61
6.97 6.96 6.95 6.94 6.93 6.92 6.91 6.90 6.89 6.88 6.87 6.86 6.85 6.84 6.83 6.82 6.81 6.80 6.79 6.78 6.77 6.76 6.75 6.74 6.73 6.72 6.71 6.70 6.69 6.68 6.67 6.66 6.65 6.64 6.63
f1 (ppm)

Current Data Parameters
NAME SQL SMFH2-M THF
EXPNO 1
PROCNO 1
Date_ 20190620
Time 10.44 h
INSTRUM Avance Neo Nanobay
PROBHD Z163739_0044 (
PULPROG zg30
TD 65536
SOLVENT THF
NS 16
DS 2
SWH 8196.722 Hz
FIDRES 0.250144 Hz
AQ 3.9976959 sec
RG 101
DW 61.000 usec
DE 13.54 usec
TE 298.0 K
D1 1.00000000 sec
TD0 1
SFO1 400.2724717 MHz
NUC1 1H
P0 3.33 usec
P1 10.00 usec
PLW1 14.49800014 W
SI 65536
SF 400.2700000 MHz
WDW EM
SSB 0
LB 0.30 Hz
GB 0
PC 1.00

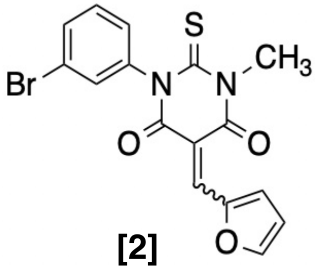

**[2]**

$^{1}$H NMR (400 MHz, THF-$d_8$)

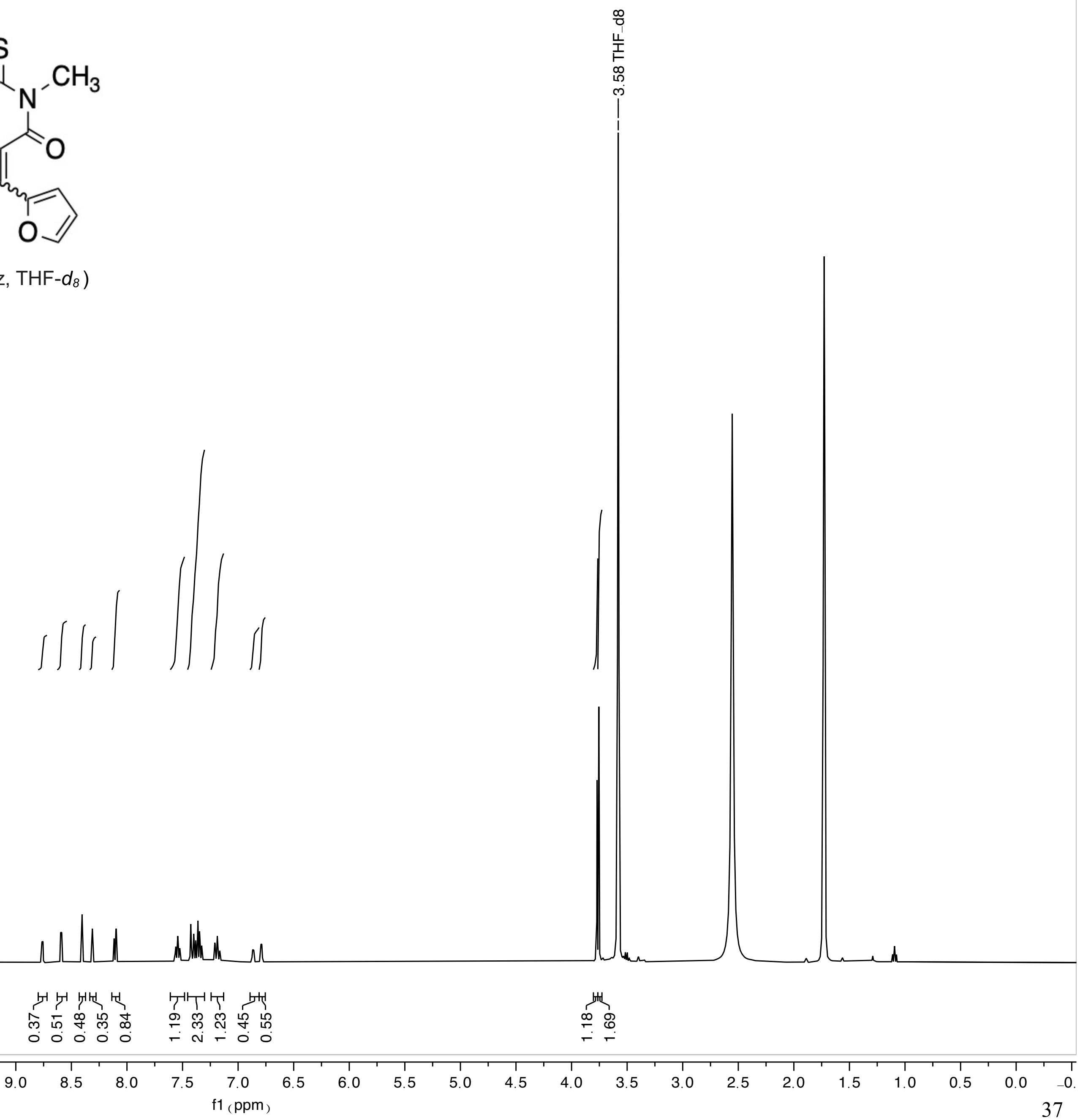

SQL-SMFH2-M

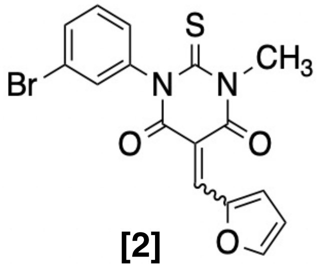

[2]

1H NMR (400 MHz, THF-$d_8$)

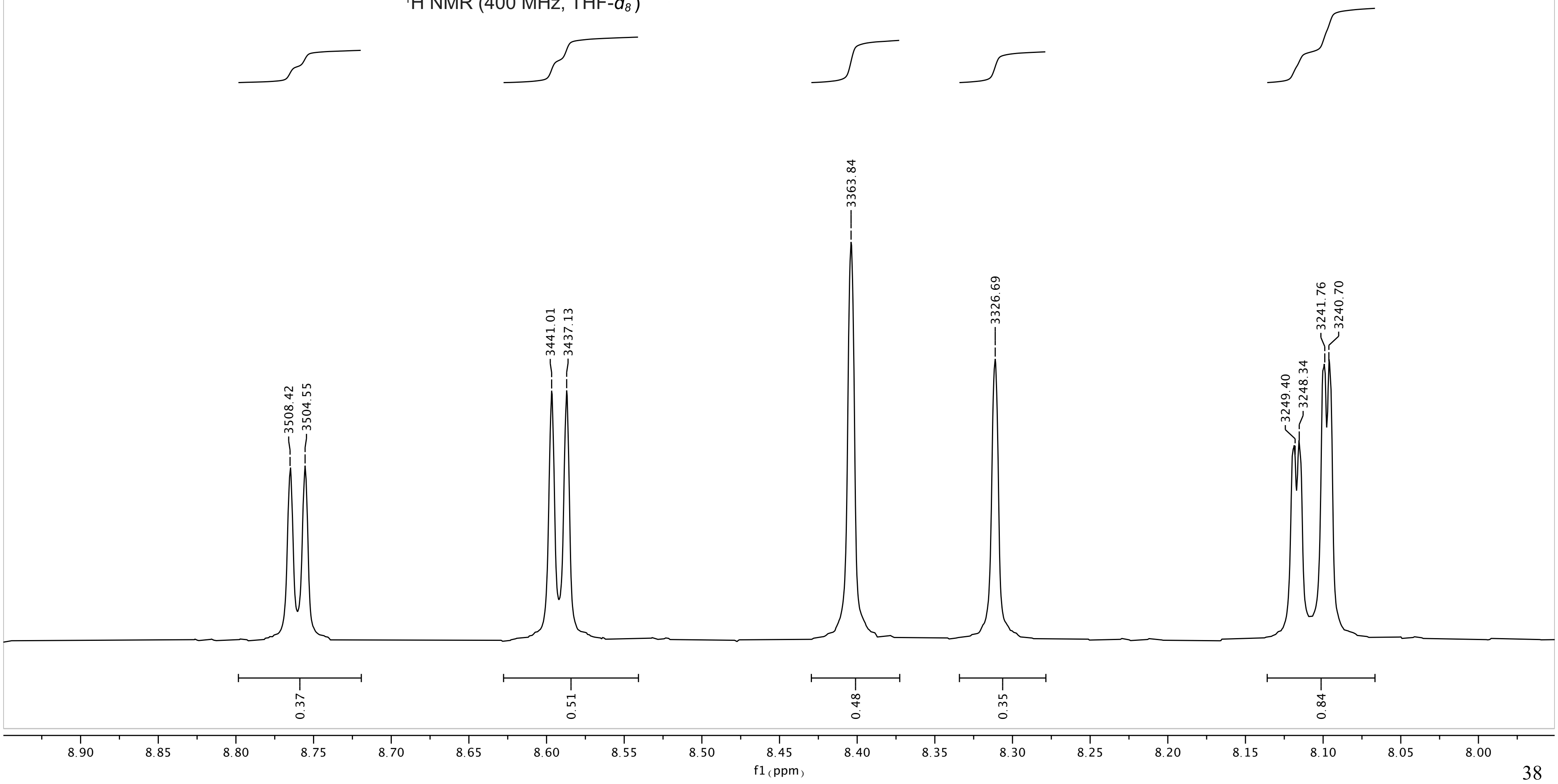

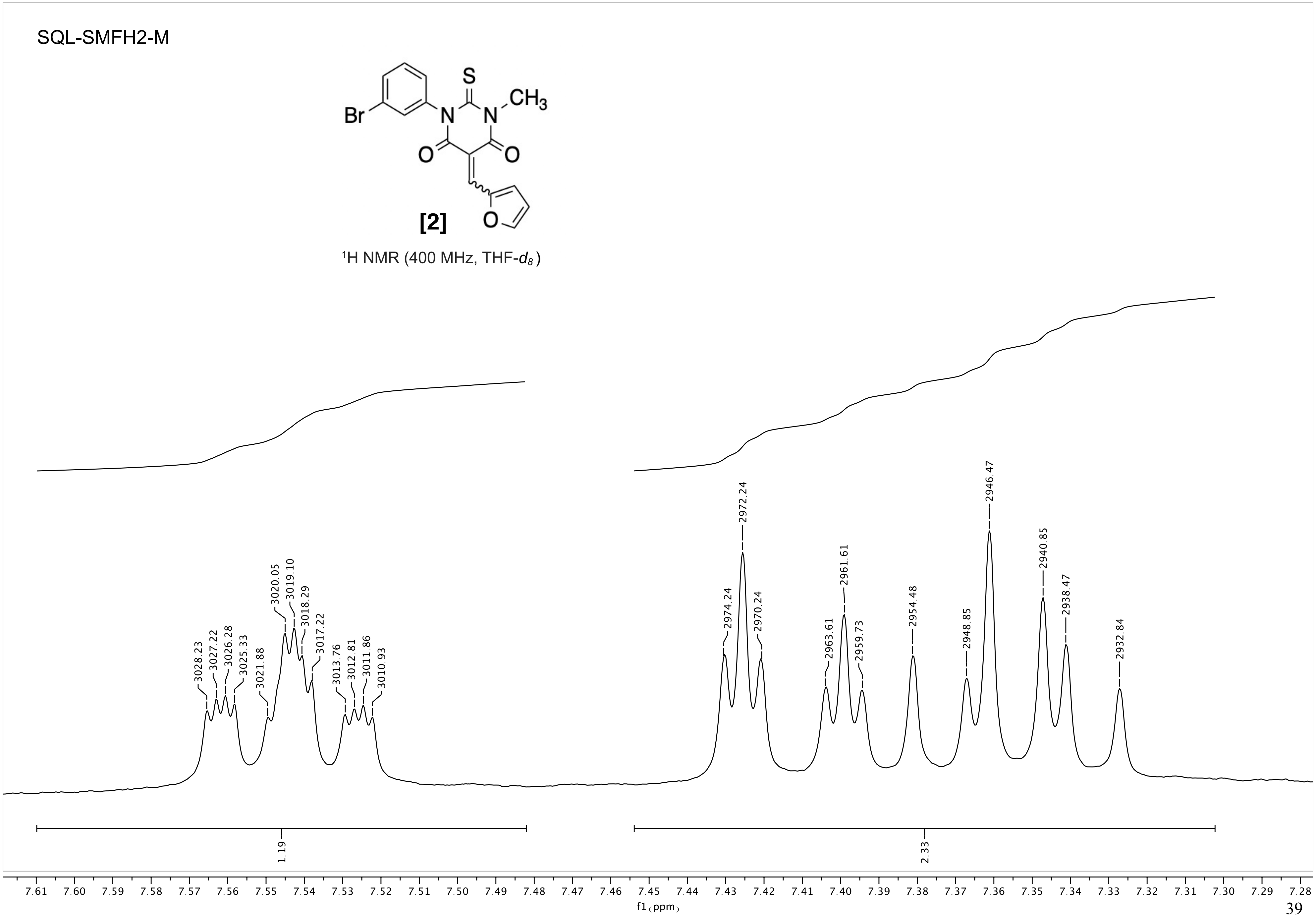
SQL-SMFH2-M
[2]
¹H NMR (400 MHz, THF-d8)
f1 (ppm)
1.19
2.33

SQL-SMFH2-M

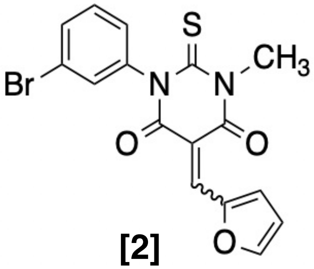

[2]

$^{1}$H NMR (400 MHz, THF-$d_8$)

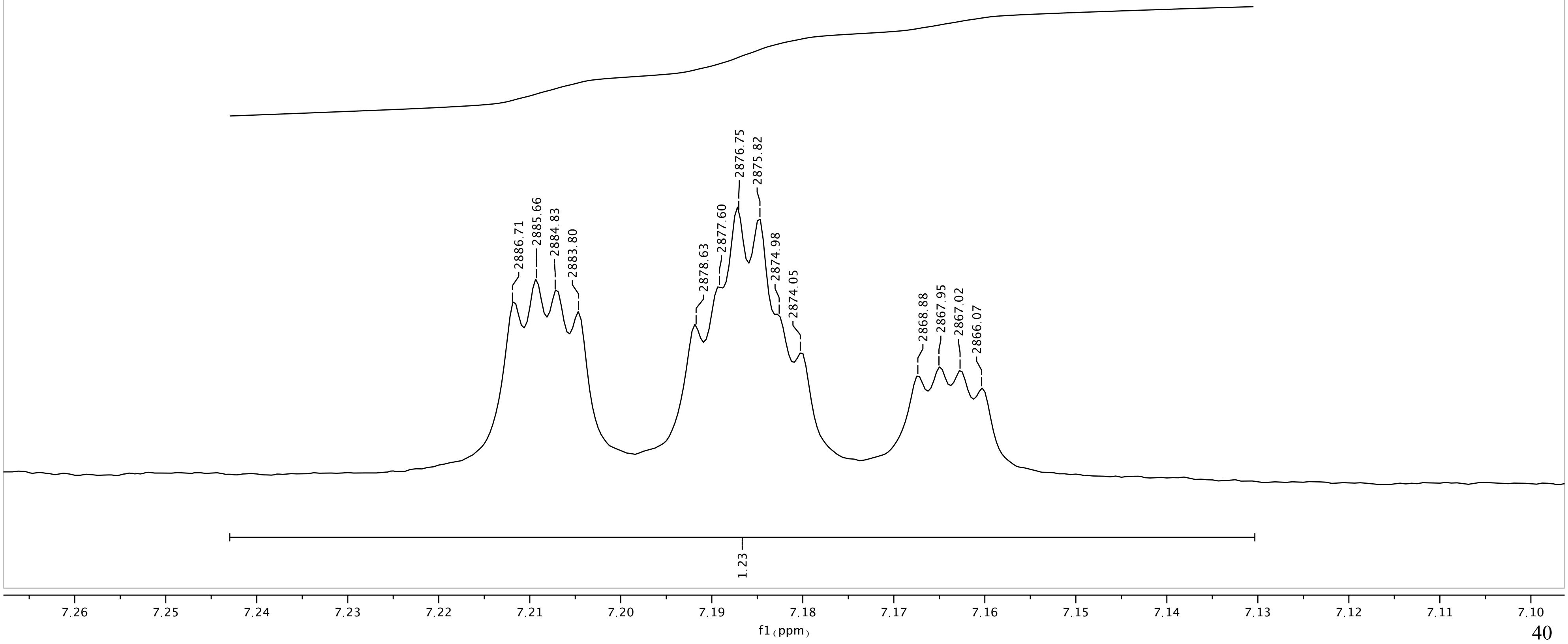

SQL-SMFH2-M

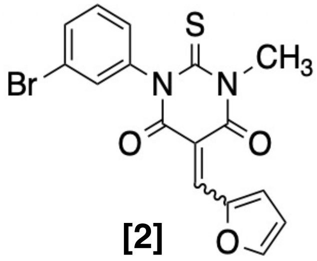

[2]

$^1H$ NMR (400 MHz, THF-$d_8$)

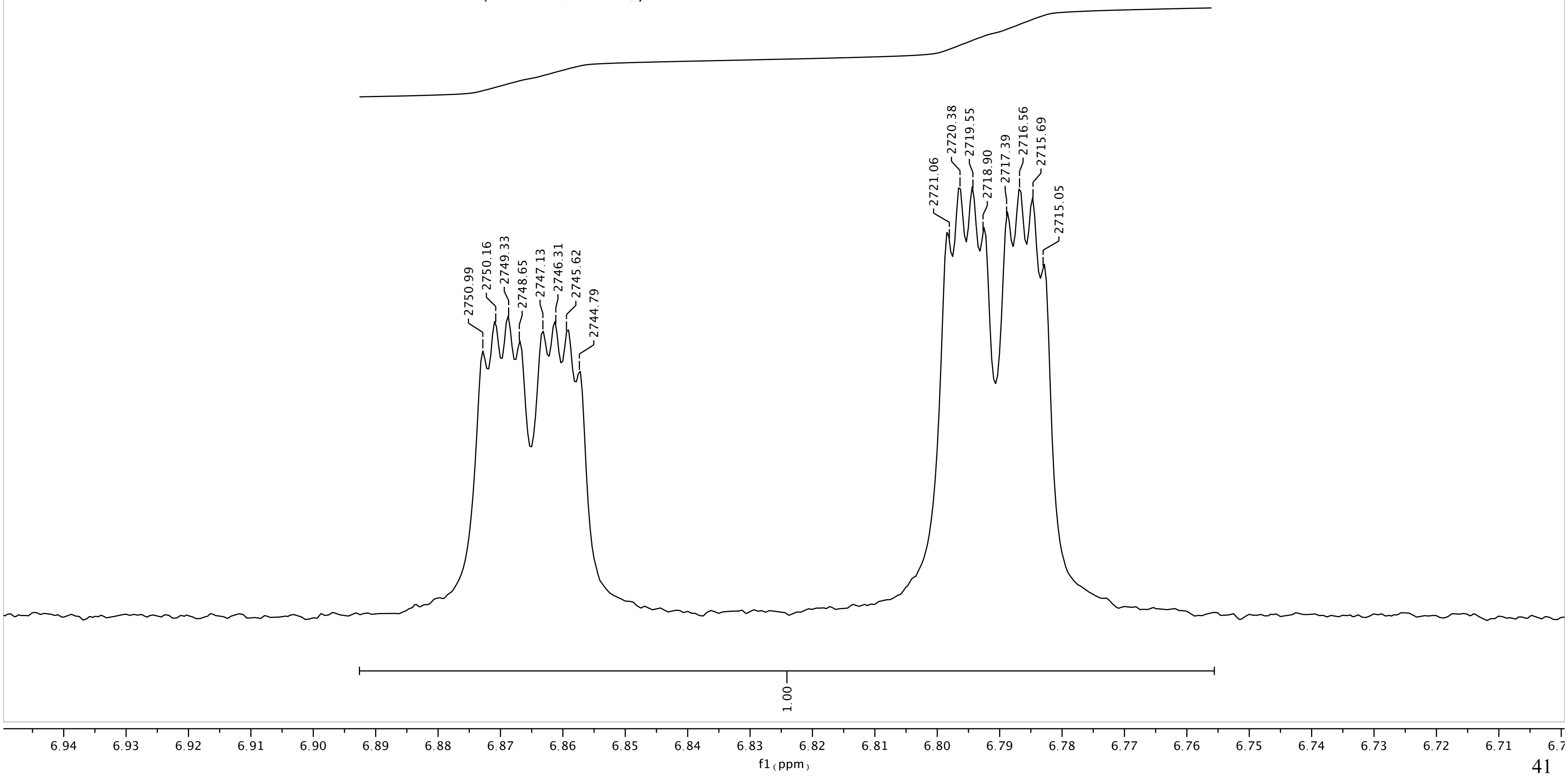

SQL-SMFH2-M

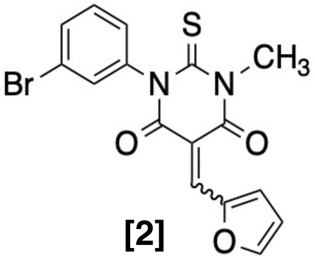

[2]

$^{1}$H NMR (400 MHz, THF-$d_8$)

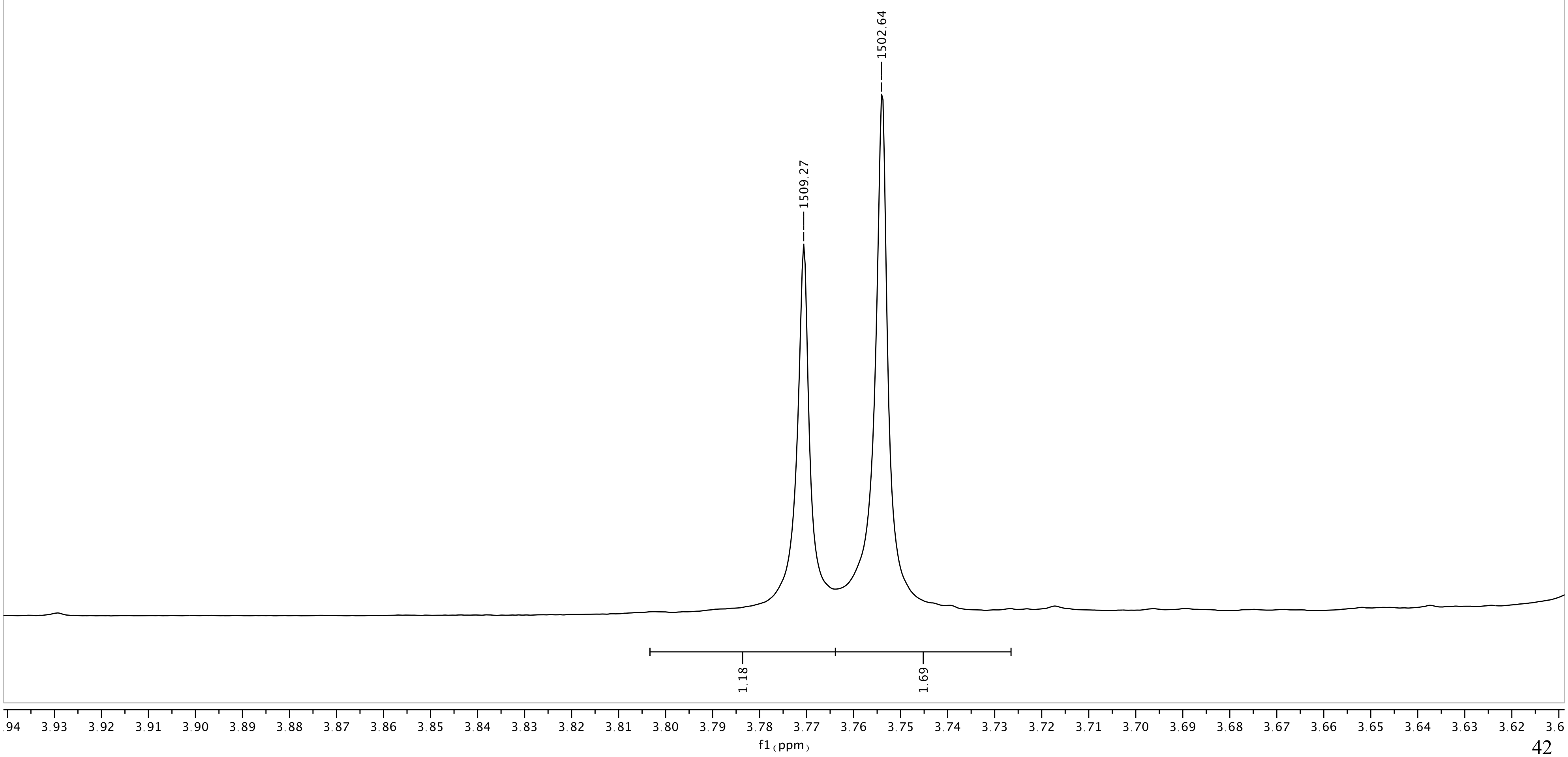

Current Data Parameters
NAME I-KGS-77
EXPNO 5
PROCNO 1
Date_ 20190626
Time 16.41 h
INSTRUM Avance Neo Nanobay
PROBHD Z163739_0044 (
PULPROG zg30
TD 65536
SOLVENT CDCl3
NS 32
DS 2
SWH 8196.722 Hz
FIDRES 0.250144 Hz
AQ 3.9976959 sec
RG 101
DW 61.000 usec
DE 13.54 usec
TE 298.0 K
D1 1.00000000 sec
TD0 1
SFO1 400.2724717 MHz
NUC1 1H
P0 3.33 usec
P1 10.00 usec
PLW1 14.49800014 W
SI 65536
SF 400.2700094 MHz
WDW EM
SSB 0
LB 0.30 Hz
GB 0
PC 1.00

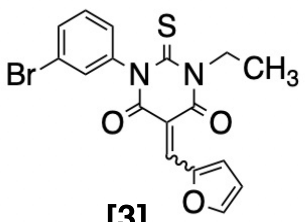

[3]

$^1$H NMR(400 MHz, $CDCl_3$)

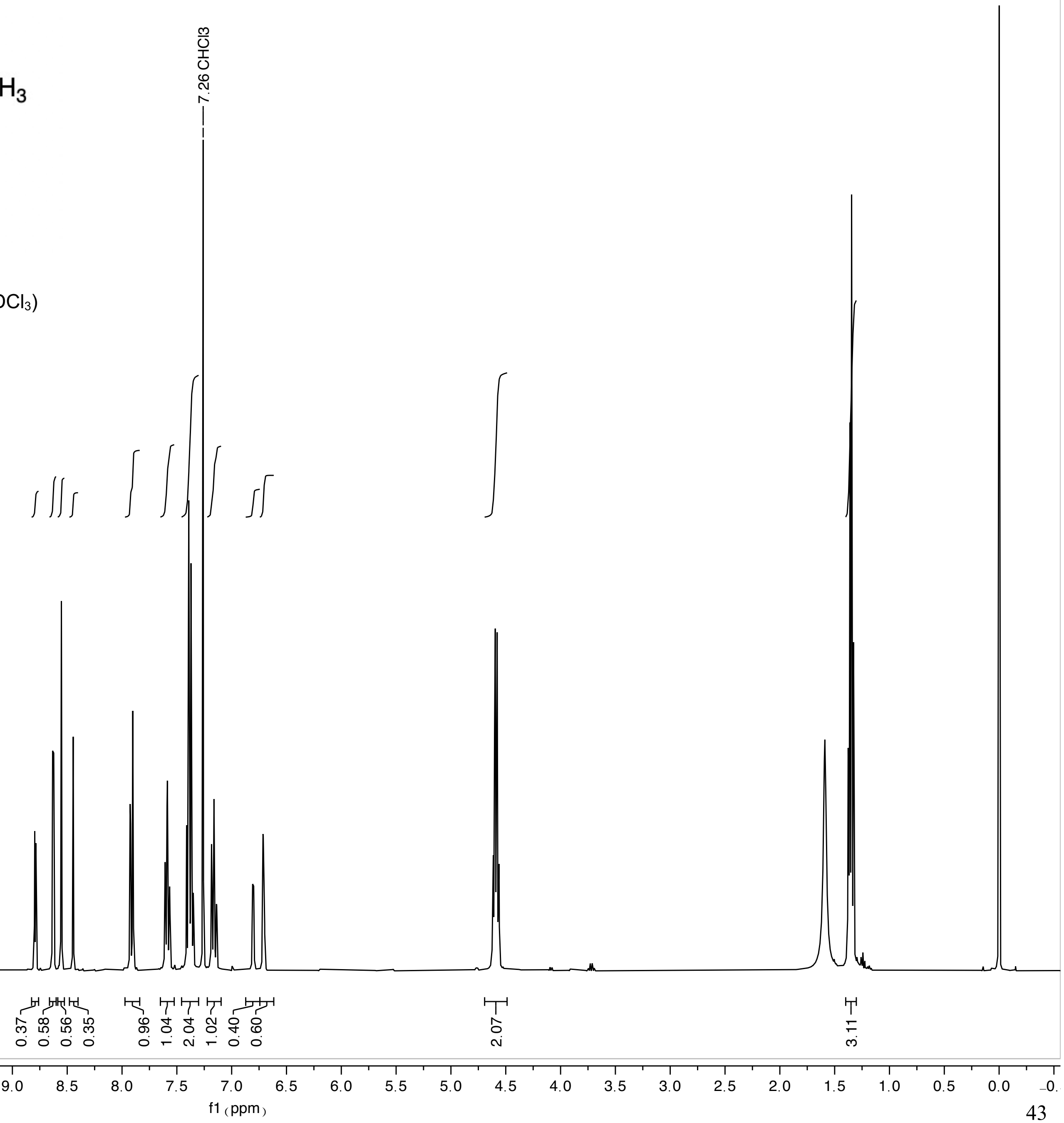

I-KGS-77

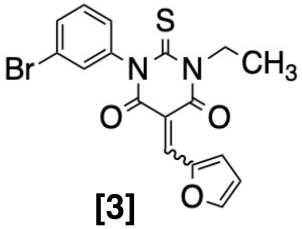

**[3]**

$^1$H NMR(400 MHz, $CDCl_3$)

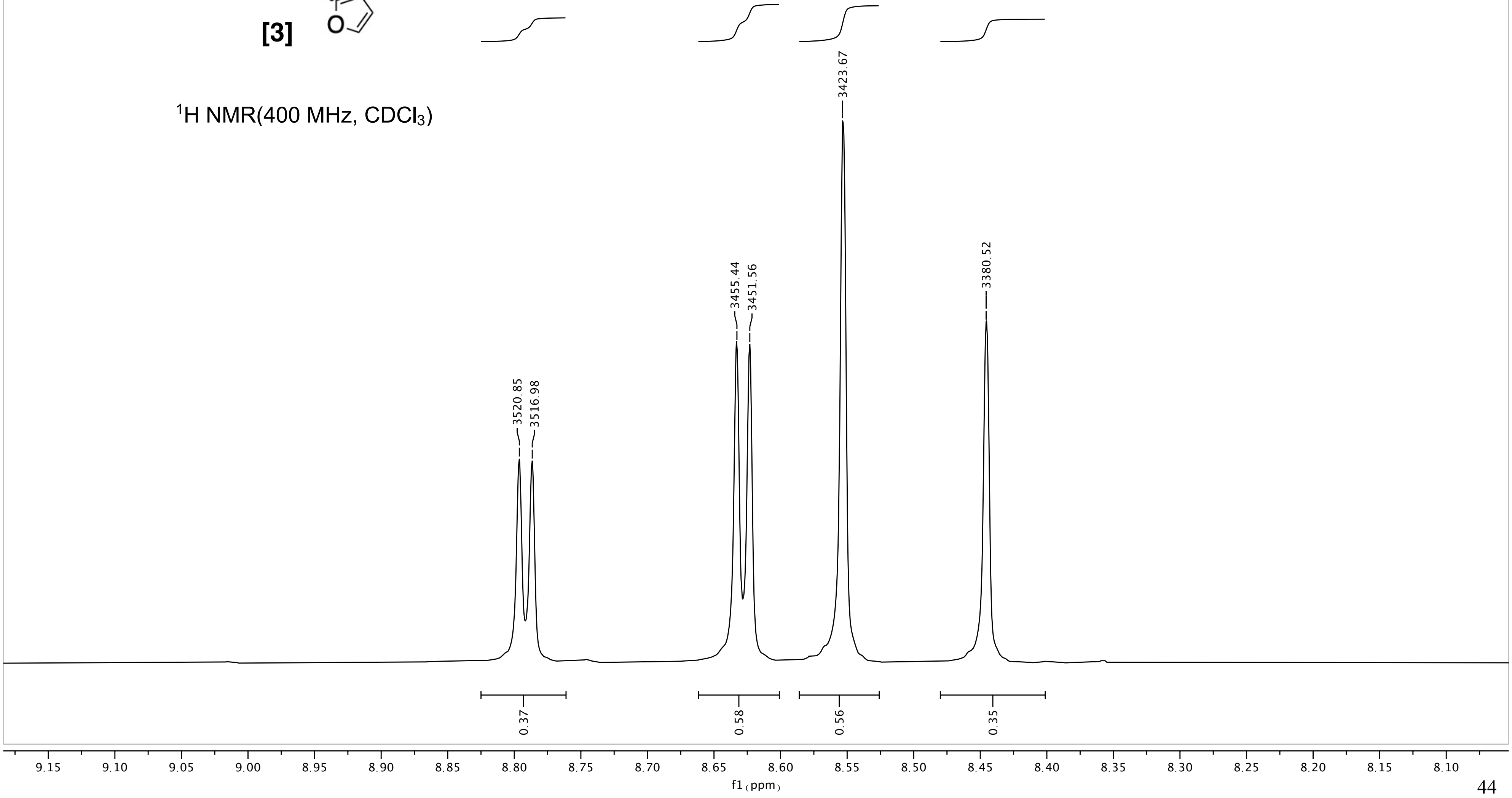

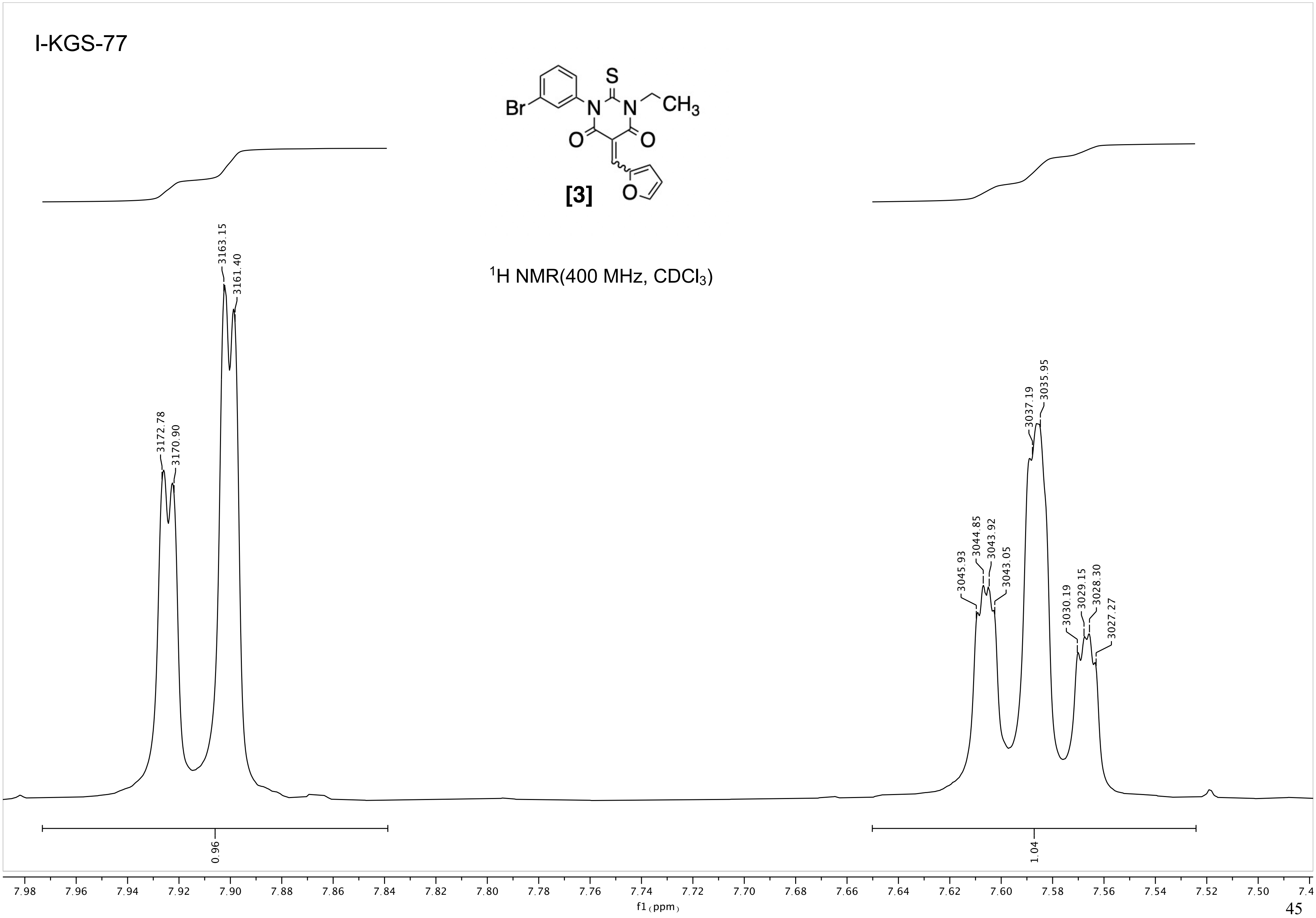
I-KGS-77
Br
N
S
N
CH3
O
O
O
[3]
1H NMR(400 MHz, CDCl3)
3172.78
3170.90
3163.15
3161.40
3045.93
3044.85
3043.92
3043.05
3037.19
3035.95
3030.19
3029.15
3028.30
3027.27
0.96
1.04
7.98 7.96 7.94 7.92 7.90 7.88 7.86 7.84 7.82 7.80 7.78 7.76 7.74 7.72 7.70 7.68 7.66 7.64 7.62 7.60 7.58 7.56 7.54 7.52 7.50 7.4
f1 (ppm)

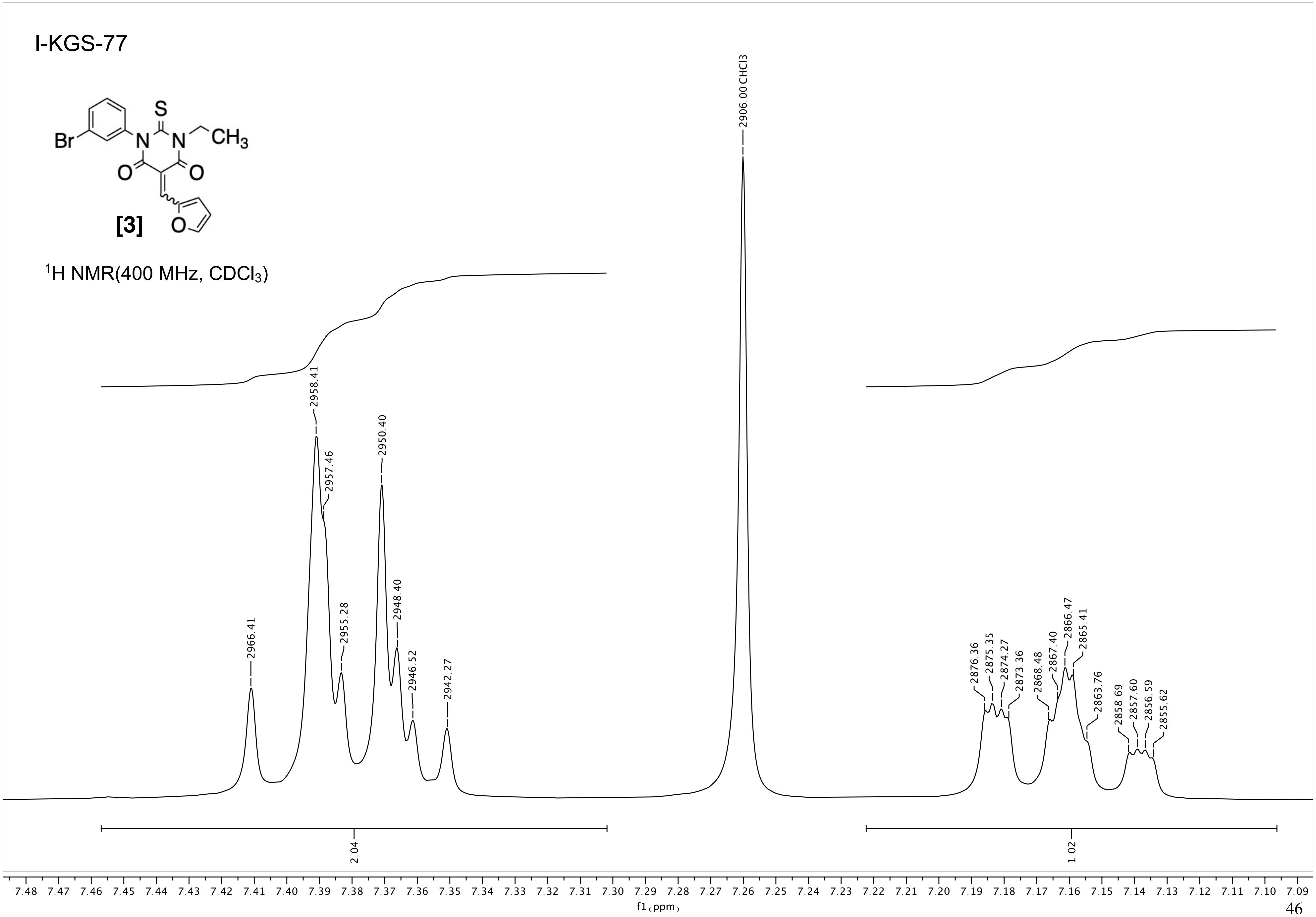
I-KGS-77
S
Br
N
N
CH3
O
O
O
[3]
1H NMR(400 MHz, CDCl3)
2906.00 CHCl3
2966.41
2958.41
2957.46
2955.28
2950.40
2948.40
2946.52
2942.27
2876.36
2875.35
2874.27
2873.36
2868.48
2867.40
2866.47
2865.41
2863.76
2858.69
2857.60
2856.59
2855.62
2.04
1.02
7.48 7.47 7.46 7.45 7.44 7.43 7.42 7.41 7.40 7.39 7.38 7.37 7.36 7.35 7.34 7.33 7.32 7.31 7.30 7.29 7.28 7.27 7.26 7.25 7.24 7.23 7.22 7.21 7.20 7.19 7.18 7.17 7.16 7.15 7.14 7.13 7.12 7.11 7.10 7.09
f1 (ppm)

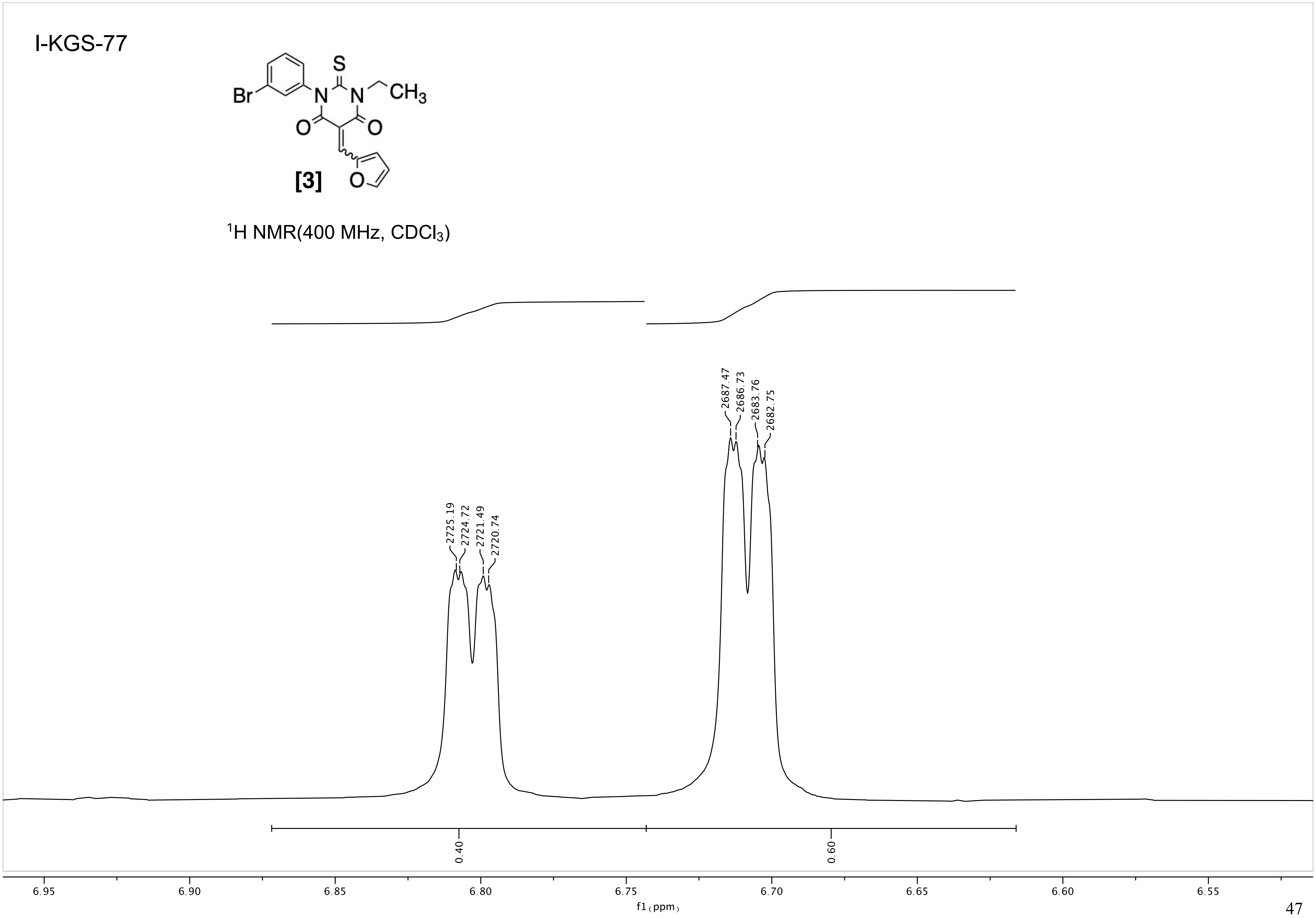

I-KGS-77
[3]
$^1$H NMR(400 MHz, CDCl$_3$)
2725.19
2724.72
2721.49
2720.74
2687.47
2686.73
2683.76
2682.75
0.40
0.60
6.95
6.90
6.85
6.80
6.75
6.70
6.65
6.60
6.55
f1 (ppm)

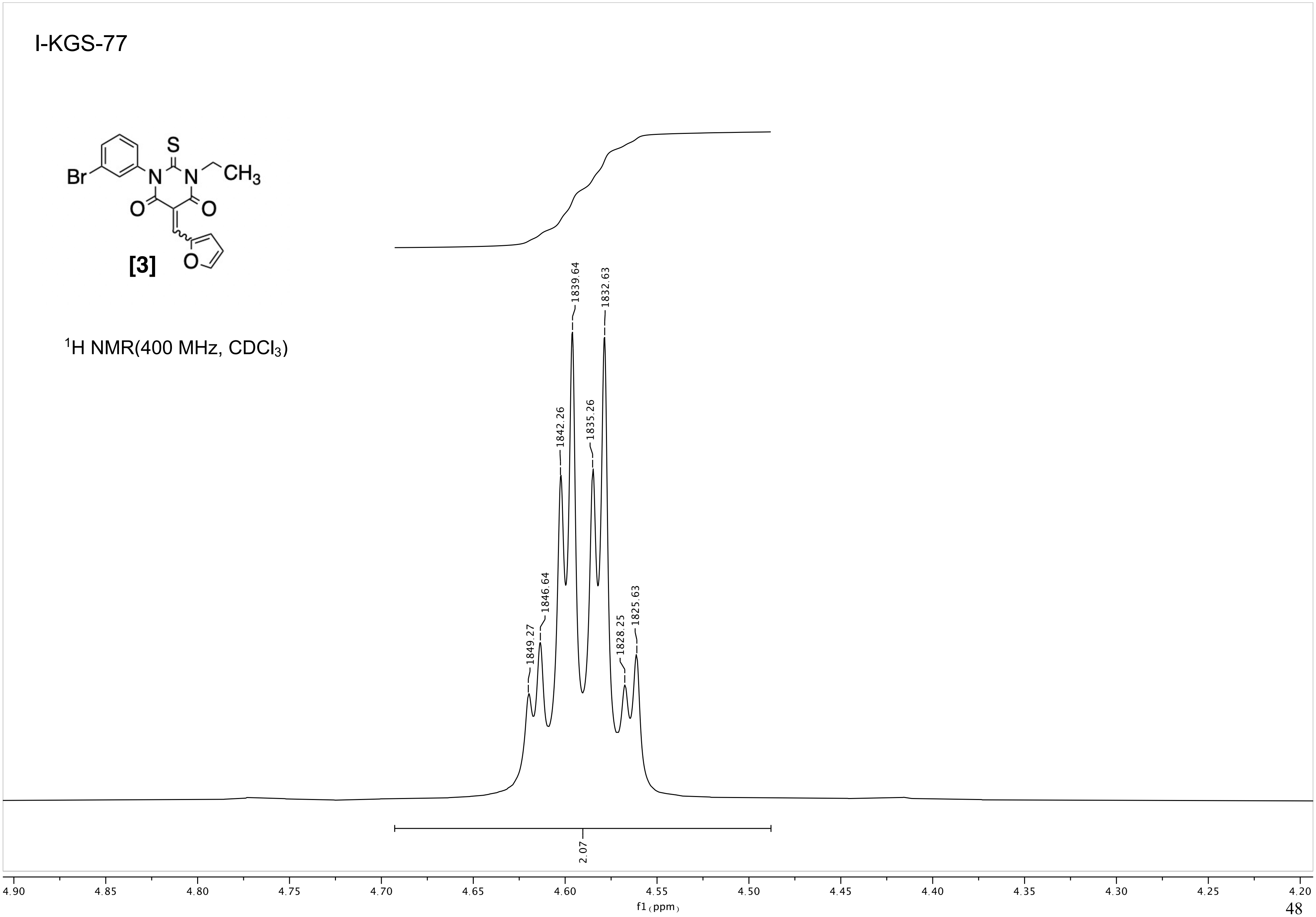
I-KGS-77
[3]
Br
S
N
CH3
O
¹H NMR(400 MHz, CDCl3)
1849.27
1846.64
1842.26
1839.64
1835.26
1832.63
1828.25
1825.63
2.07
4.90
4.85
4.80
4.75
4.70
4.65
4.60
4.55
4.50
4.45
4.40
4.35
4.30
4.25
4.20
f1 (ppm)

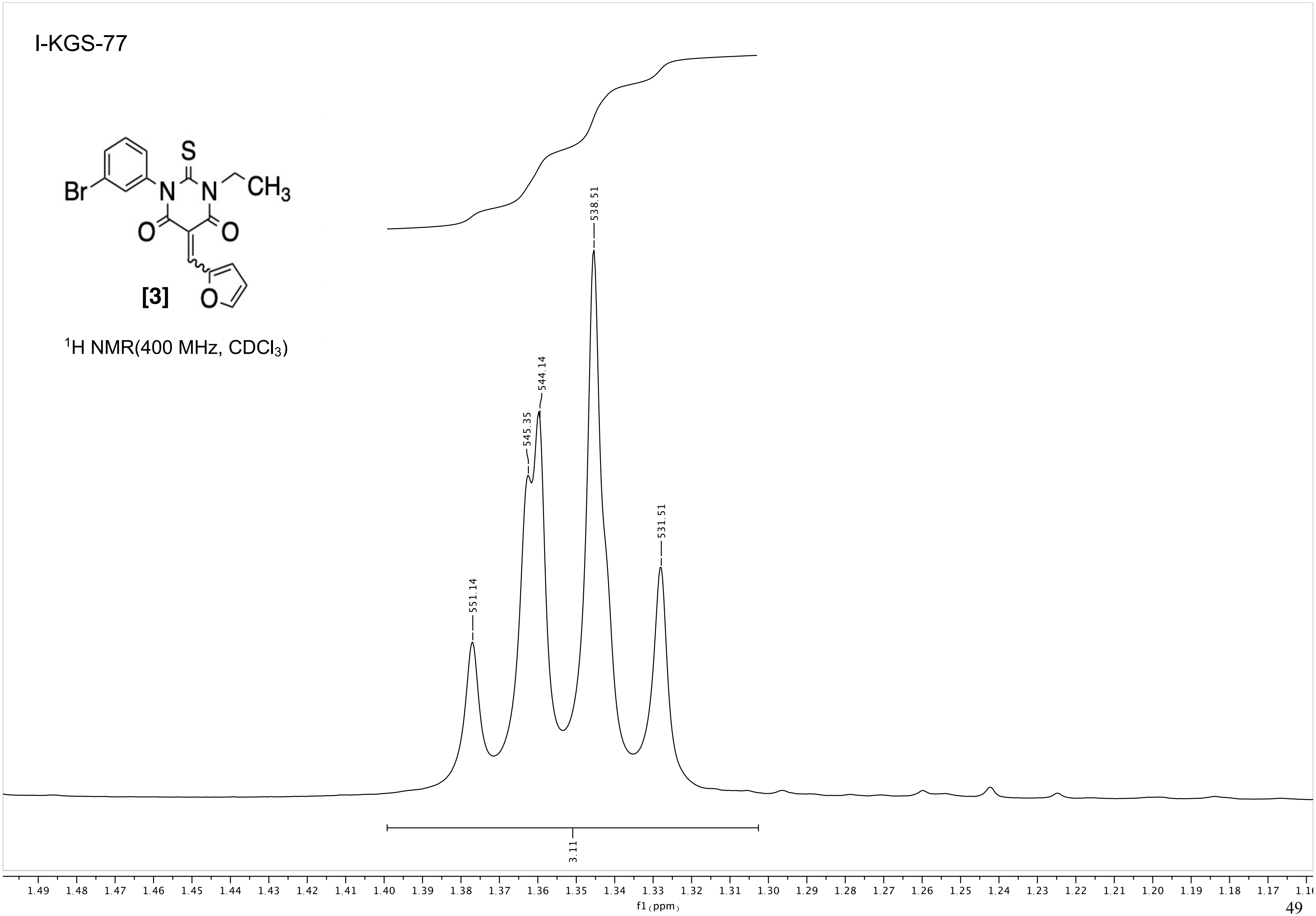

I-KGS-77
Br
N
S
N
CH3
O
O
[3]
O
1H NMR(400 MHz, CDCl3)
551.14
545.35
544.14
538.51
531.51
3.11
1.49 1.48 1.47 1.46 1.45 1.44 1.43 1.42 1.41 1.40 1.39 1.38 1.37 1.36 1.35 1.34 1.33 1.32 1.31 1.30 1.29 1.28 1.27 1.26 1.25 1.24 1.23 1.22 1.21 1.20 1.19 1.18 1.17
f1 (ppm)

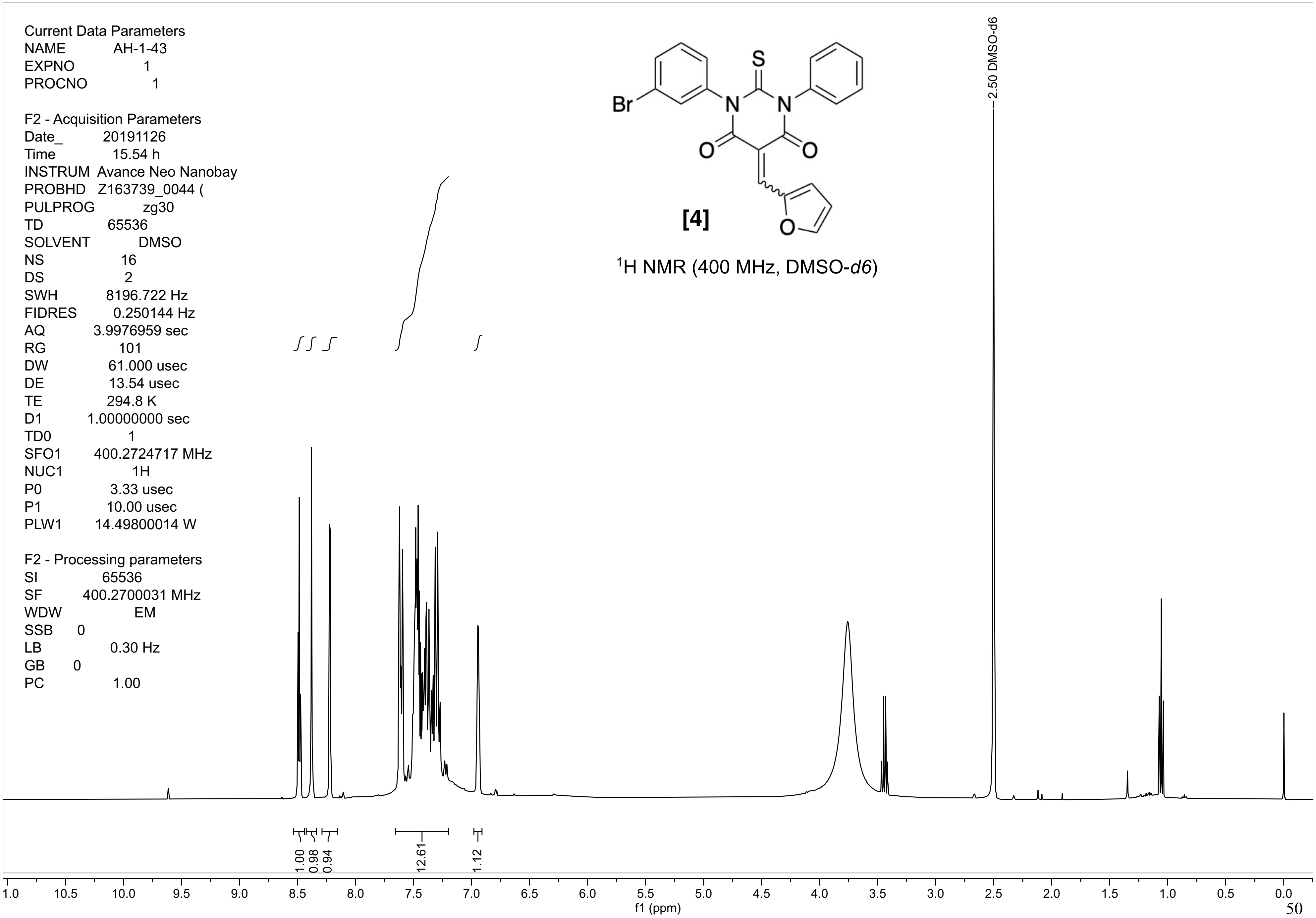

Current Data Parameters
NAME AH-1-43
EXPNO 1
PROCNO 1
F2 - Acquisition Parameters
Date_ 20191126
Time 15.54 h
INSTRUM Avance Neo Nanobay
PROBHD Z163739_0044 (
PULPROG zg30
TD 65536
SOLVENT DMSO
NS 16
DS 2
SWH 8196.722 Hz
FIDRES 0.250144 Hz
AQ 3.9976959 sec
RG 101
DW 61.000 usec
DE 13.54 usec
TE 294.8 K
D1 1.00000000 sec
TD0 1
SFO1 400.2724717 MHz
NUC1 1H
P0 3.33 usec
P1 10.00 usec
PLW1 14.49800014 W
F2 - Processing parameters
SI 65536
SF 400.2700031 MHz
WDW EM
SSB 0
LB 0.30 Hz
GB 0
PC 1.00
Br
S
N
N
O
O
O
[4]
¹H NMR (400 MHz, DMSO-d6)
2.50 DMSO-d6
1.00
0.98
0.94
12.61
1.12
f1 (ppm)

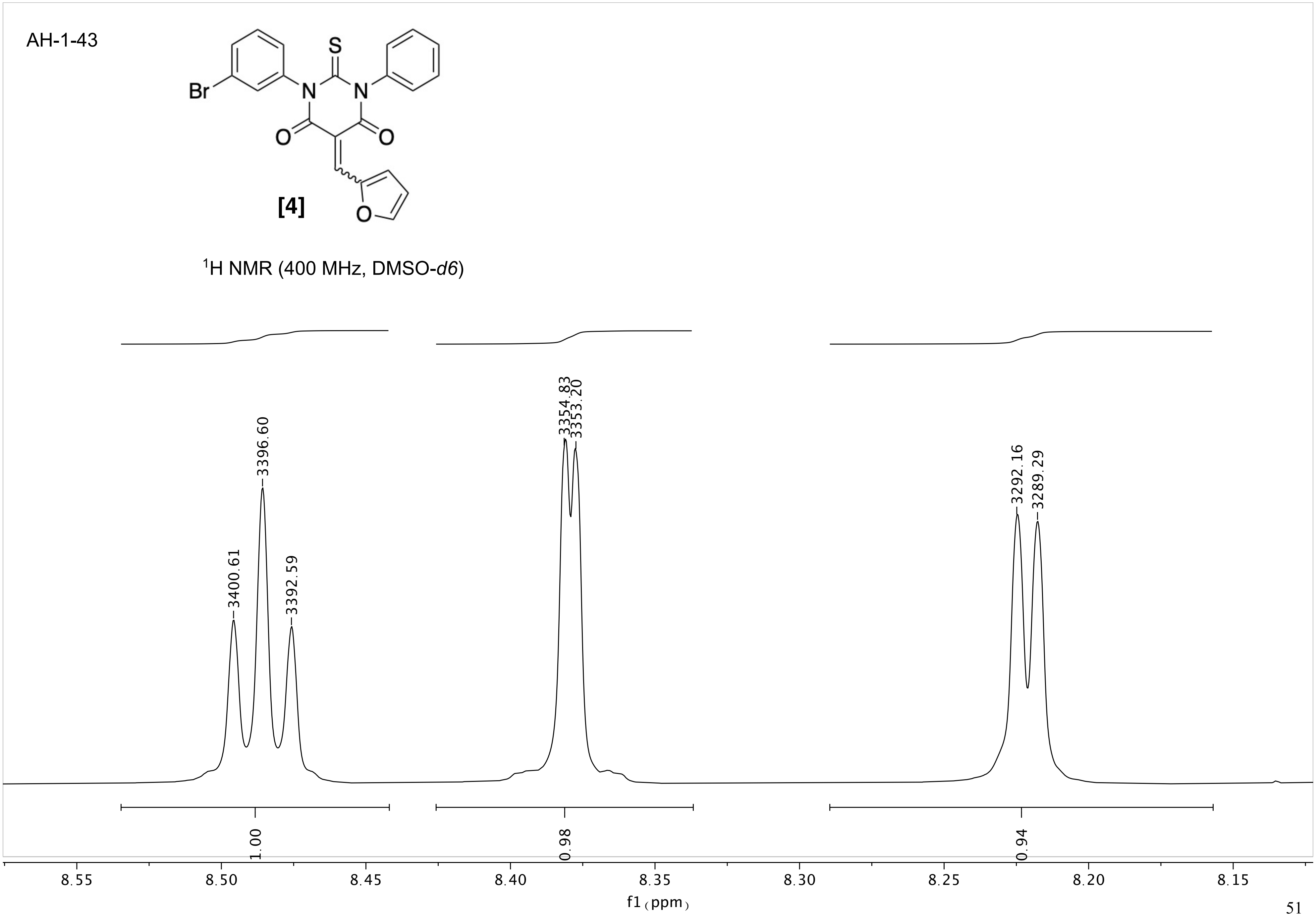

AH-1-43
Br
S
N
N
O
O
O
[4]
$^1$H NMR (400 MHz, DMSO-*d6*)
3400.61
3396.60
3392.59
3354.83
3353.20
3292.16
3289.29
1.00
0.98
0.94
8.55
8.50
8.45
8.40
8.35
8.30
8.25
8.20
8.15
f1 (ppm)

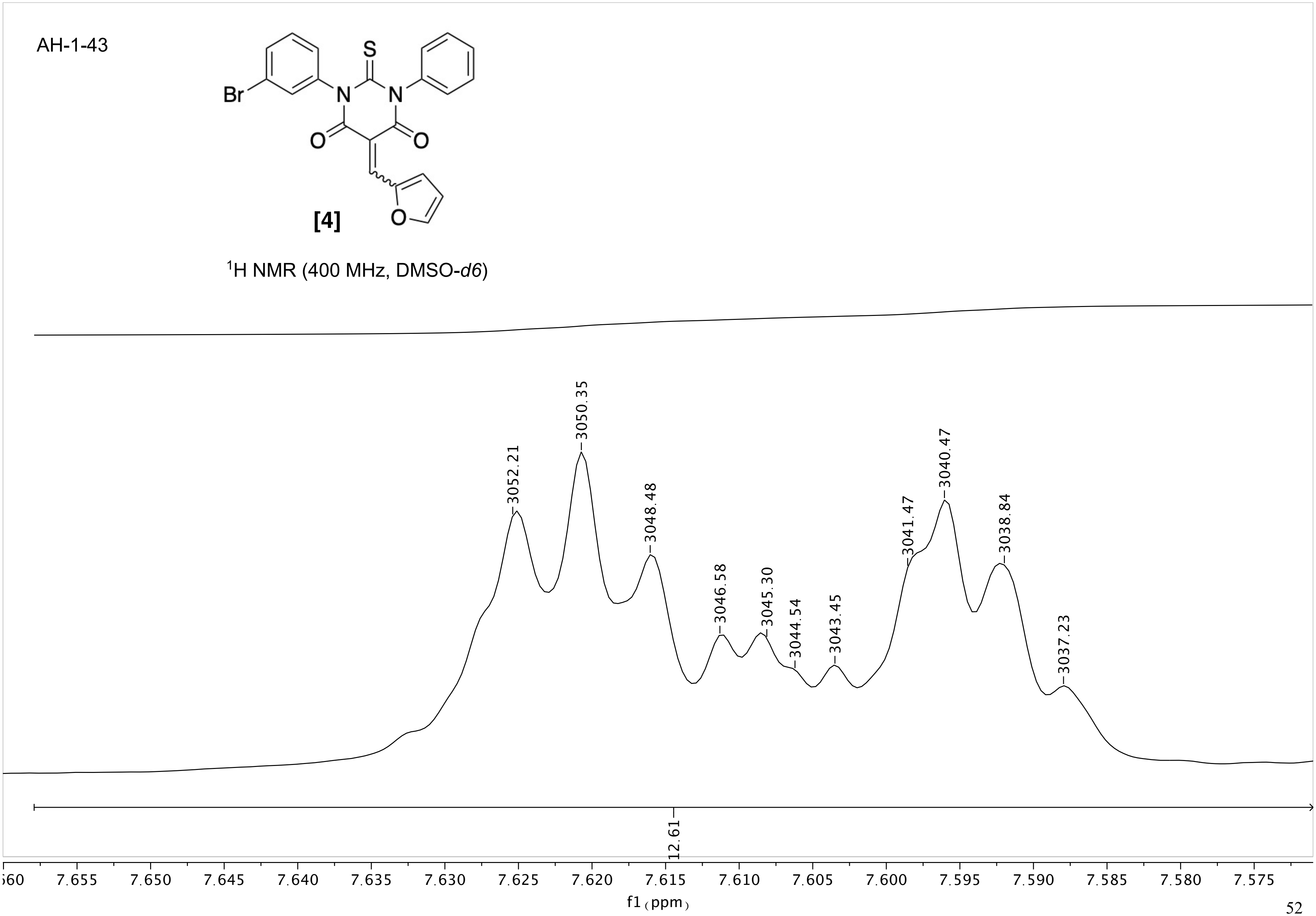
AH-1-43
[4]
1H NMR (400 MHz, DMSO-d6)
3052.21
3050.35
3048.48
3046.58
3045.30
3044.54
3043.45
3041.47
3040.47
3038.84
3037.23
12.61
7.655 7.650 7.645 7.640 7.635 7.630 7.625 7.620 7.615 7.610 7.605 7.600 7.595 7.590 7.585 7.580 7.575
f1 (ppm)

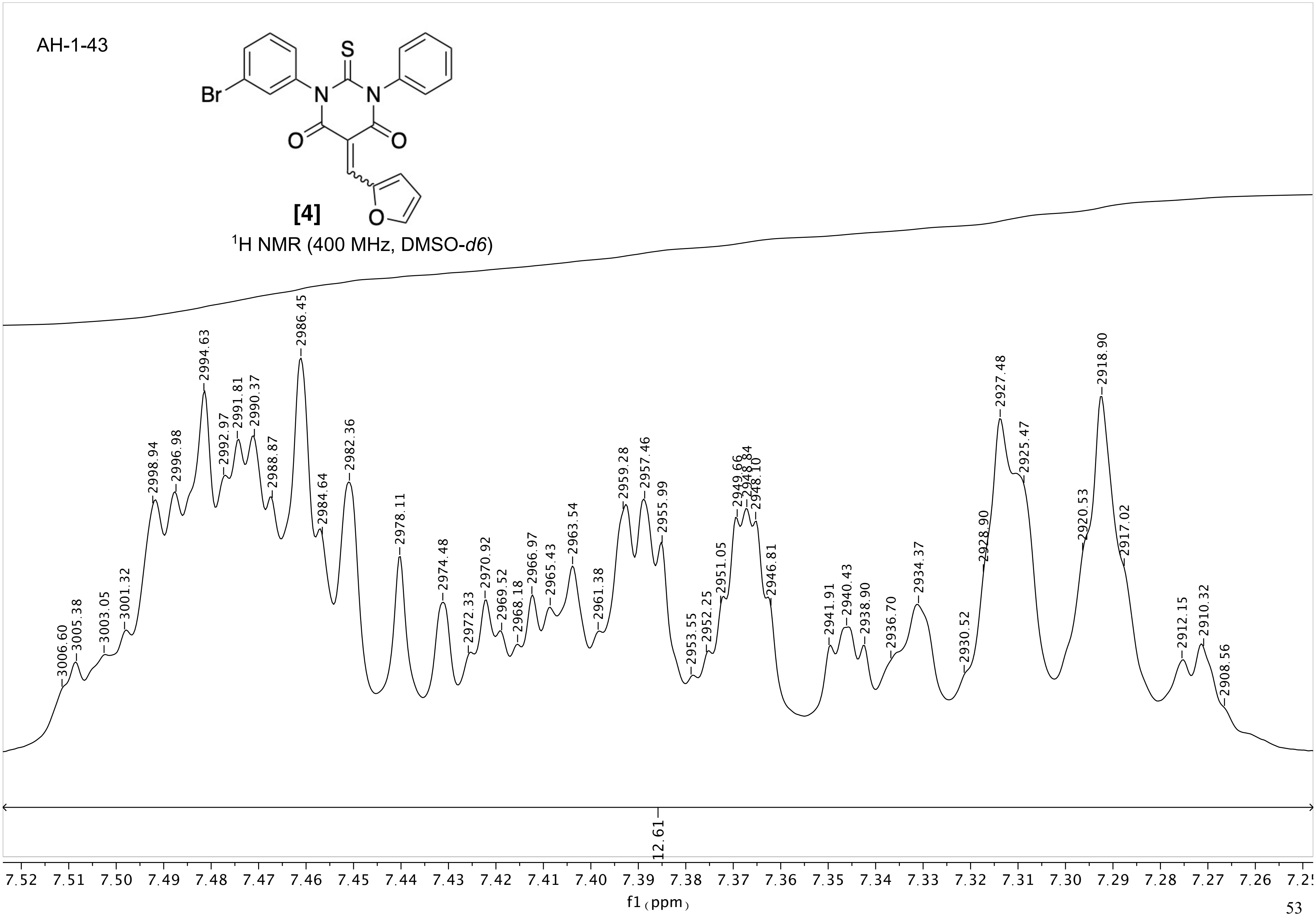
AH-1-43
[4]
1H NMR (400 MHz, DMSO-d6)
12.61
f1 (ppm)

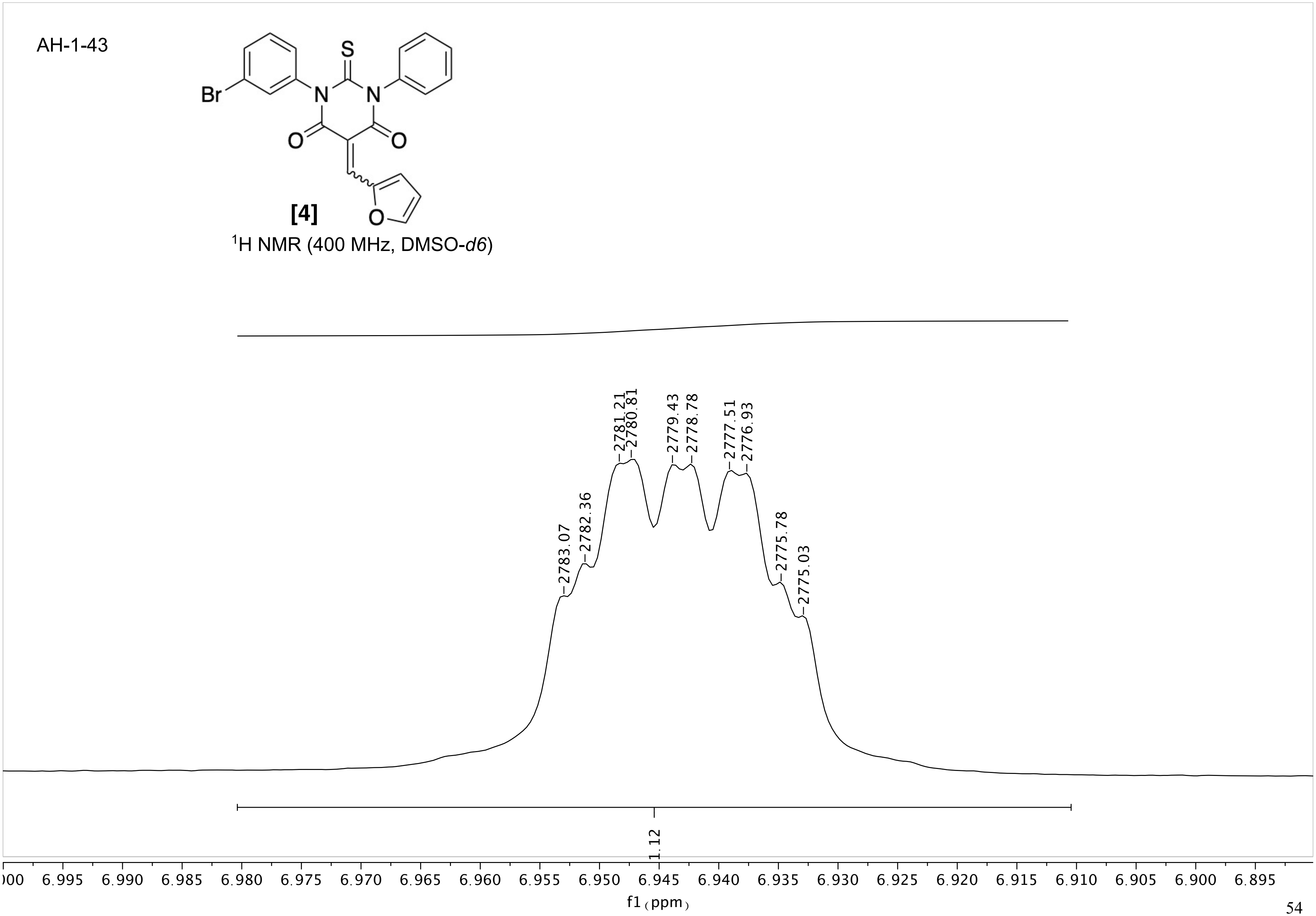
AH-1-43
S
N
N
Br
O
O
O
[4]
1H NMR (400 MHz, DMSO-d6)
2783.07
2782.36
2781.21
2780.81
2779.43
2778.78
2777.51
2776.93
2775.78
2775.03
1.12
f1 (ppm)

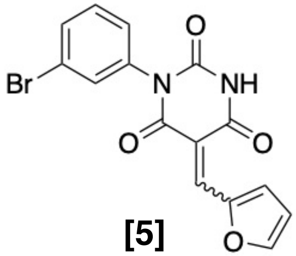

**[5]**

$^{1}$H NMR (400 MHz, DMSO-*d6*)

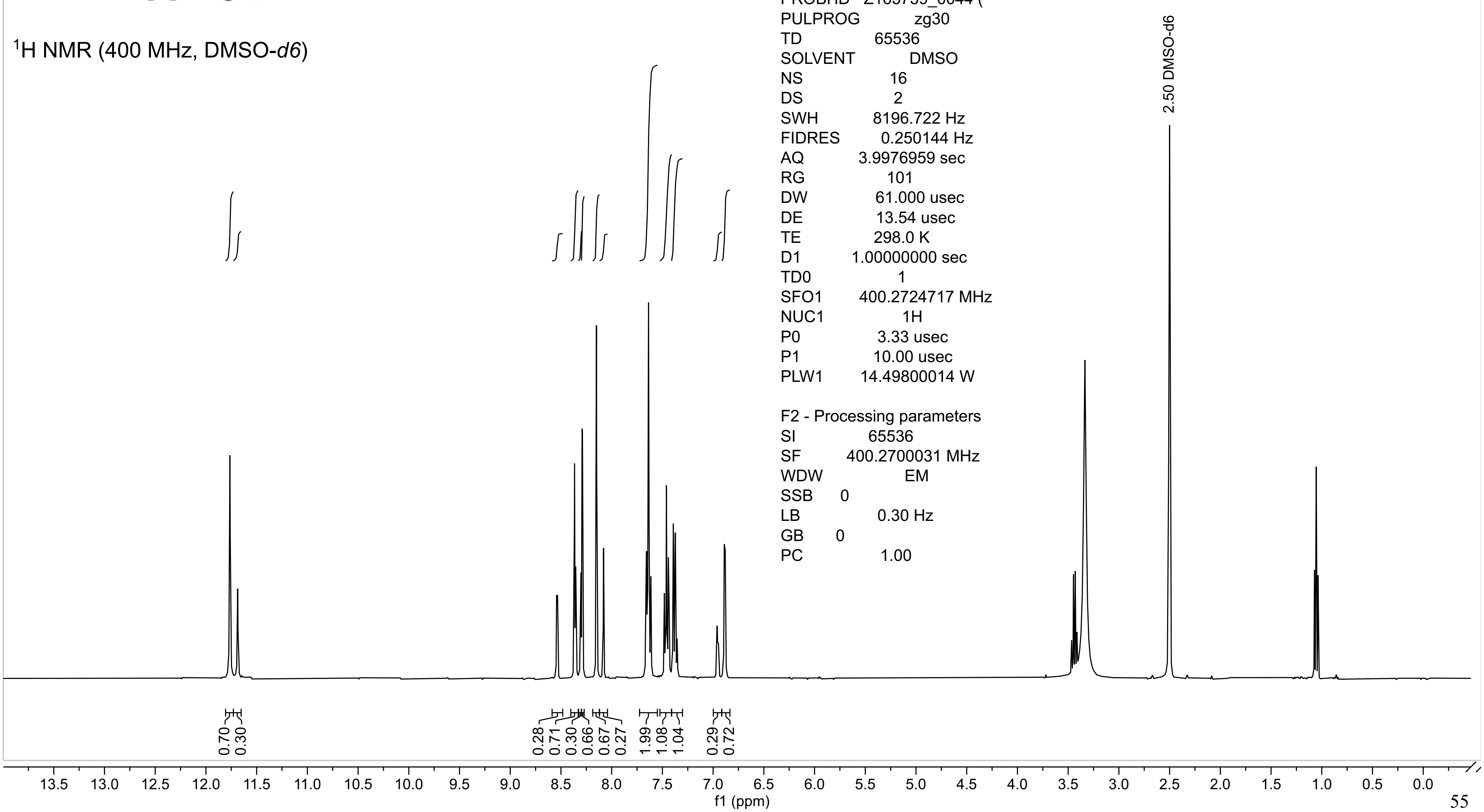

Current Data Parameters
NAME I-MAL-28
EXPNO 1
PROCNO 1

F2 - Acquisition Parameters
Date_ 20190702
Time 13.35 h
INSTRUM Avance Neo Nanobay
PROBHD Z163739_0044 (
PULPROG zg30
TD 65536
SOLVENT DMSO
NS 16
DS 2
SWH 8196.722 Hz
FIDRES 0.250144 Hz
AQ 3.9976959 sec
RG 101
DW 61.000 usec
DE 13.54 usec
TE 298.0 K
D1 1.00000000 sec
TD0 1
SFO1 400.2724717 MHz
NUC1 1H
P0 3.33 usec
P1 10.00 usec
PLW1 14.49800014 W

F2 - Processing parameters
SI 65536
SF 400.2700031 MHz
WDW EM
SSB 0
LB 0.30 Hz
GB 0
PC 1.00

I-MAL-28

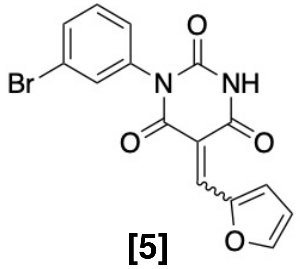

[5]

$^1$H NMR (400 MHz, DMSO-*d6*)

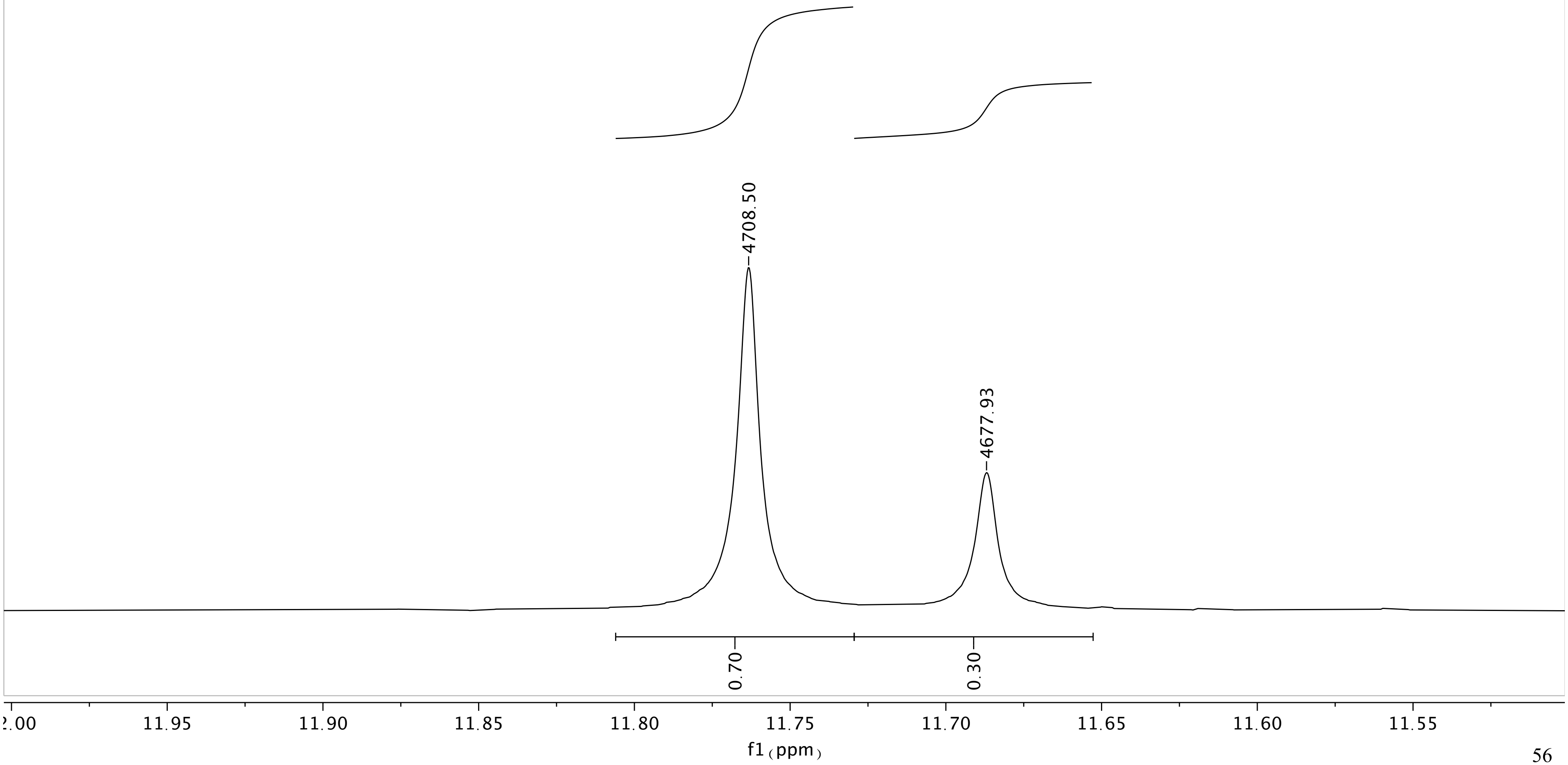

I-MAL-28

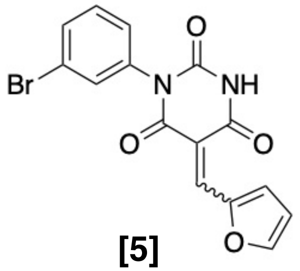

**[5]**

$^1$H NMR (400 MHz, DMSO-*d6*)

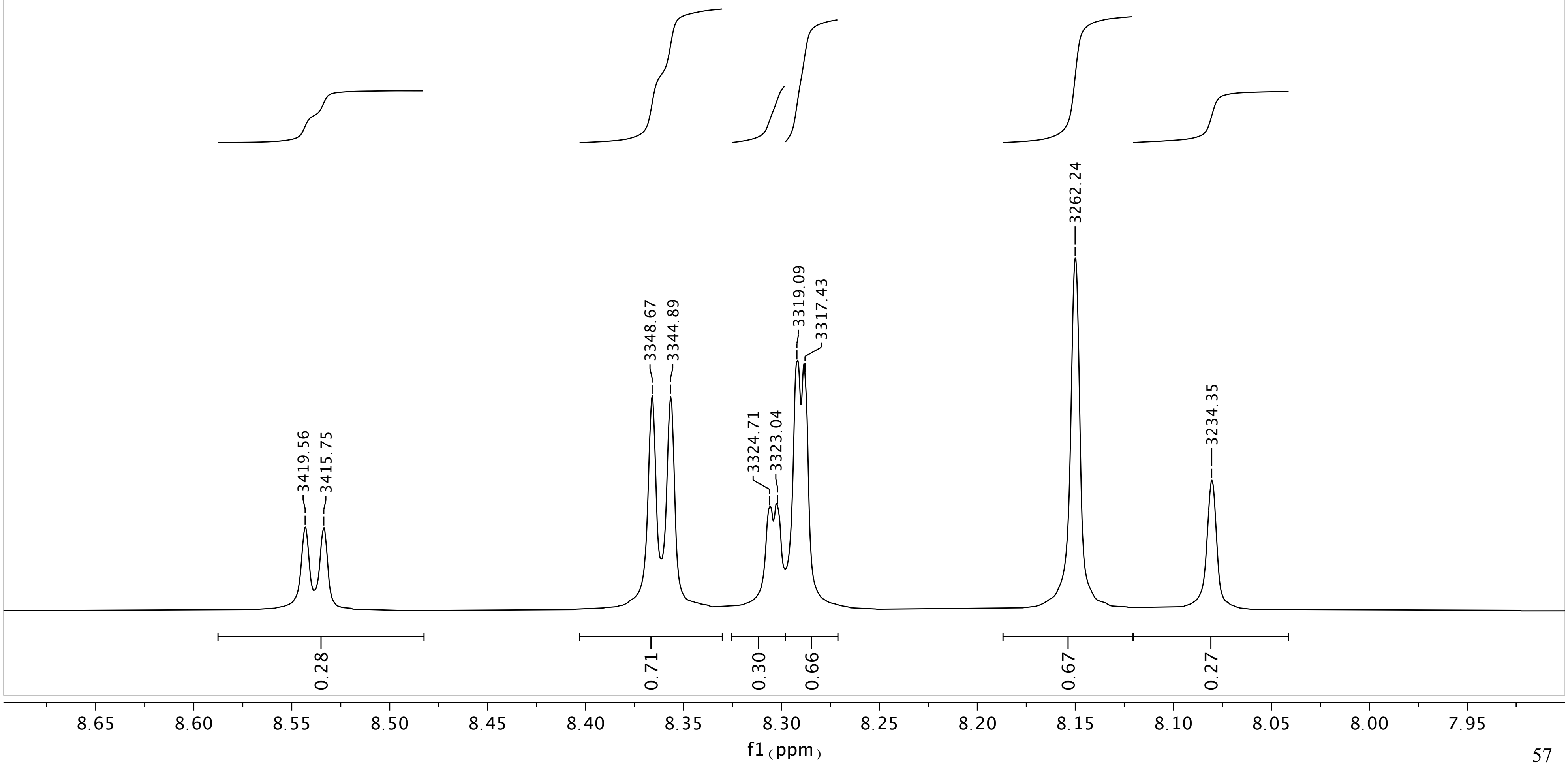

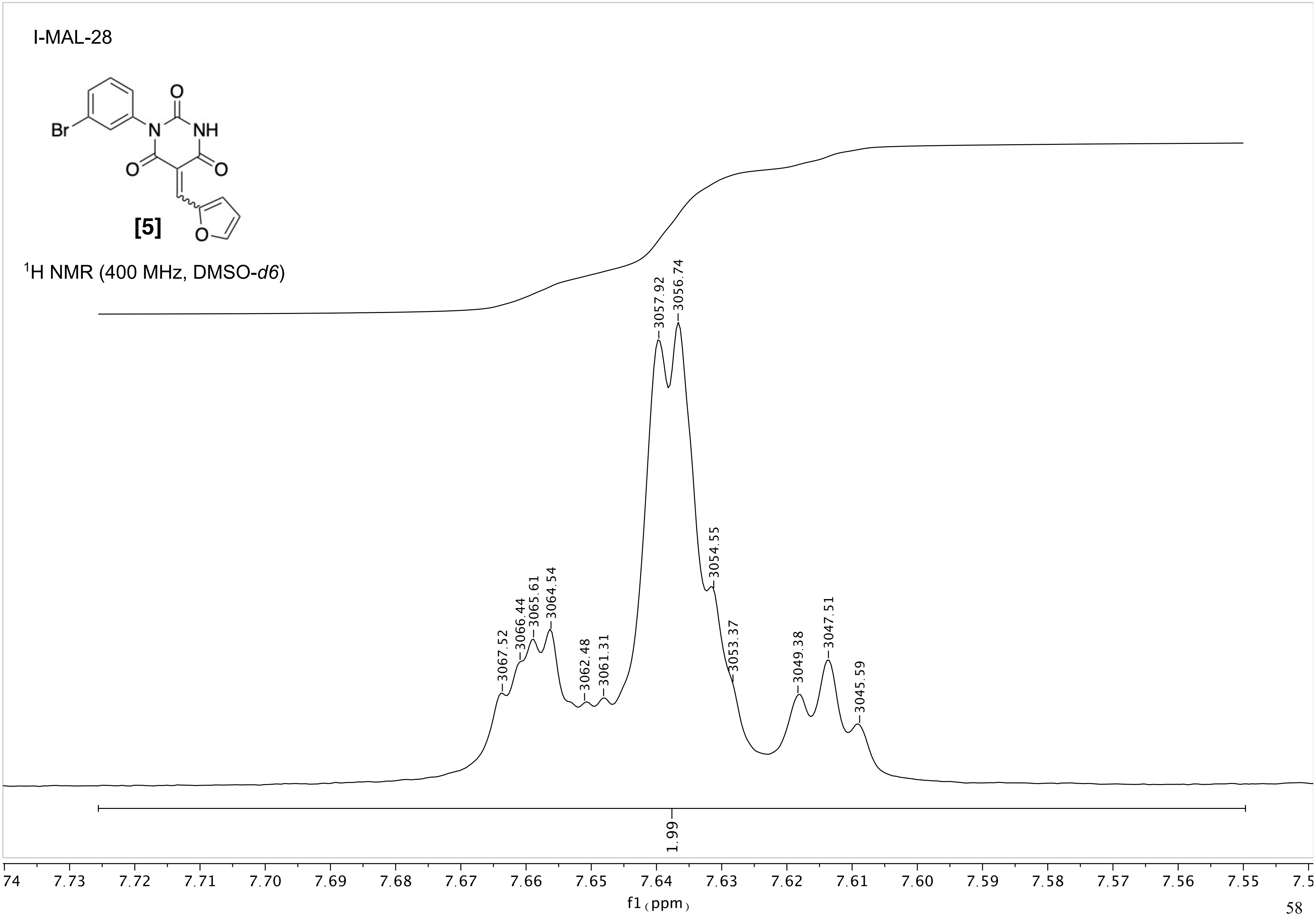
I-MAL-28
Br
O
N
NH
O
O
[5]
O
1H NMR (400 MHz, DMSO-d6)
3057.92
3056.74
3054.55
3053.37
3067.52
3066.44
3065.61
3064.54
3062.48
3061.31
3049.38
3047.51
3045.59
1.99
74
7.73
7.72
7.71
7.70
7.69
7.68
7.67
7.66
7.65
7.64
7.63
7.62
7.61
7.60
7.59
7.58
7.57
7.56
7.55
7.5
f1 (ppm)

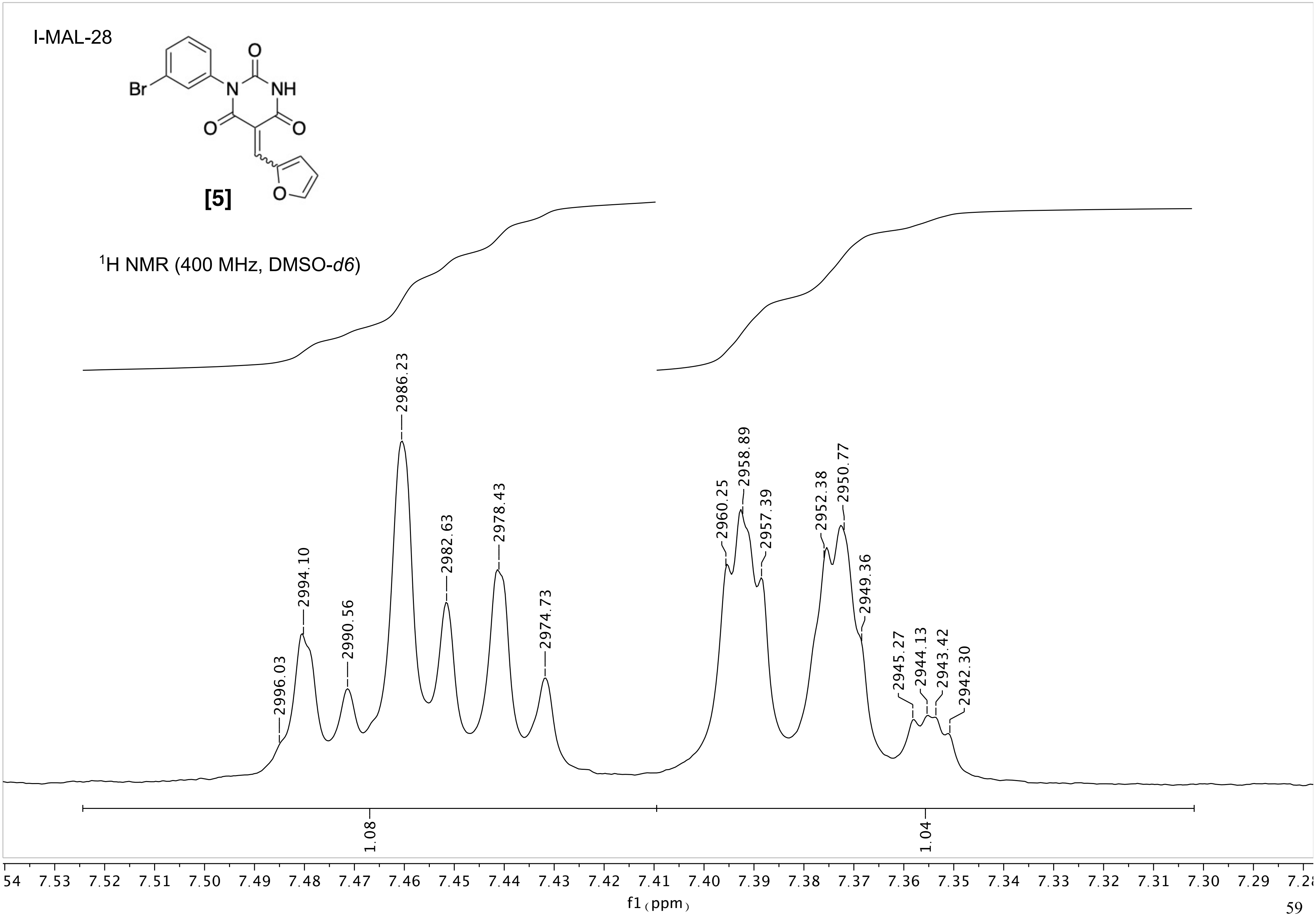
I-MAL-28
[5]
$^1$H NMR (400 MHz, DMSO-$d6$)
2996.03
2994.10
2990.56
2986.23
2982.63
2978.43
2974.73
2960.25
2958.89
2957.39
2952.38
2950.77
2949.36
2945.27
2944.13
2943.42
2942.30
1.08
1.04
f1 (ppm)

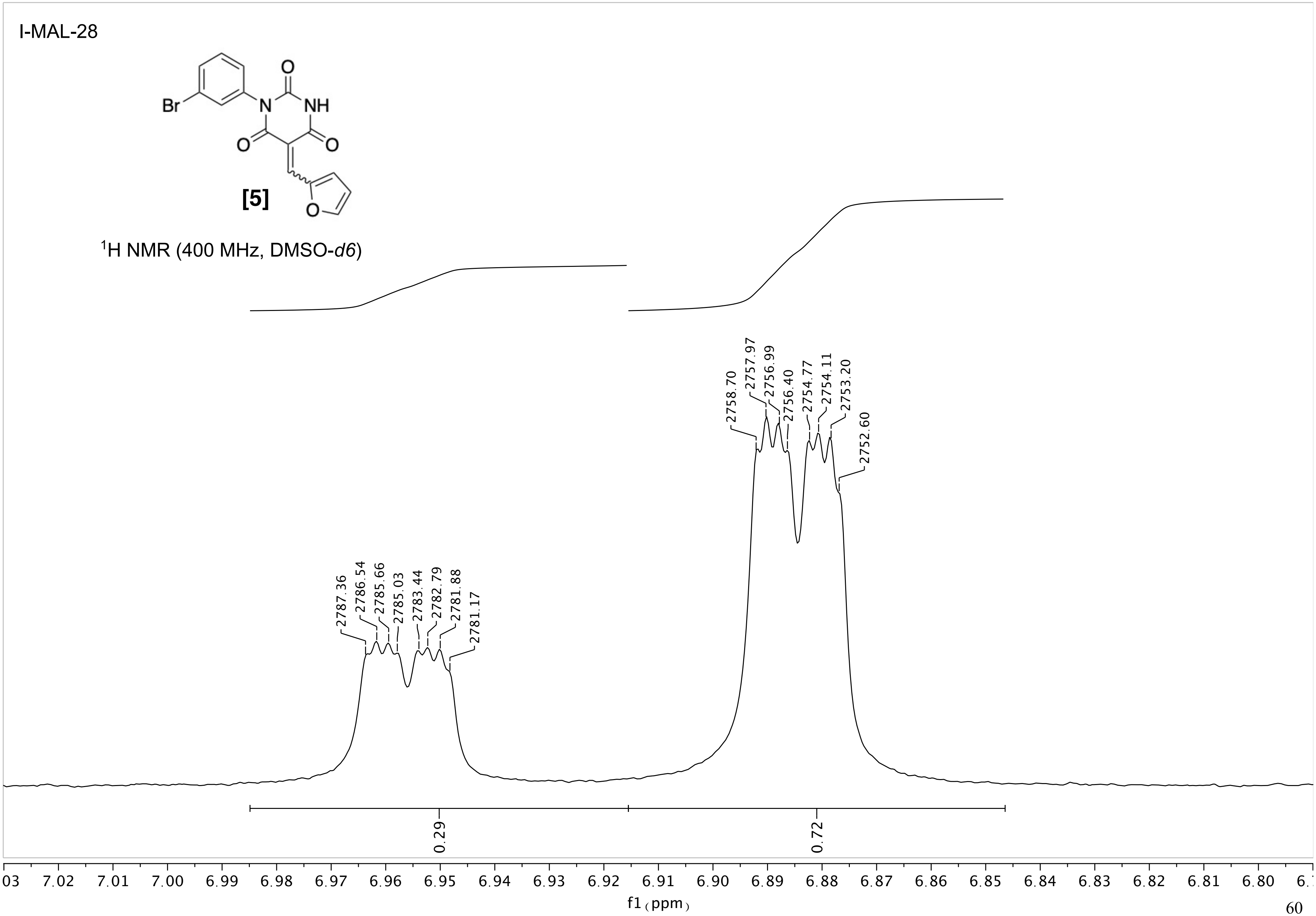
I-MAL-28
Br
N
NH
O
O
O
O
[5]
1H NMR (400 MHz, DMSO-d6)
2787.36
2786.54
2785.66
2785.03
2783.44
2782.79
2781.88
2781.17
2758.70
2757.97
2756.99
2756.40
2754.77
2754.11
2753.20
2752.60
0.29
0.72
f1 (ppm)

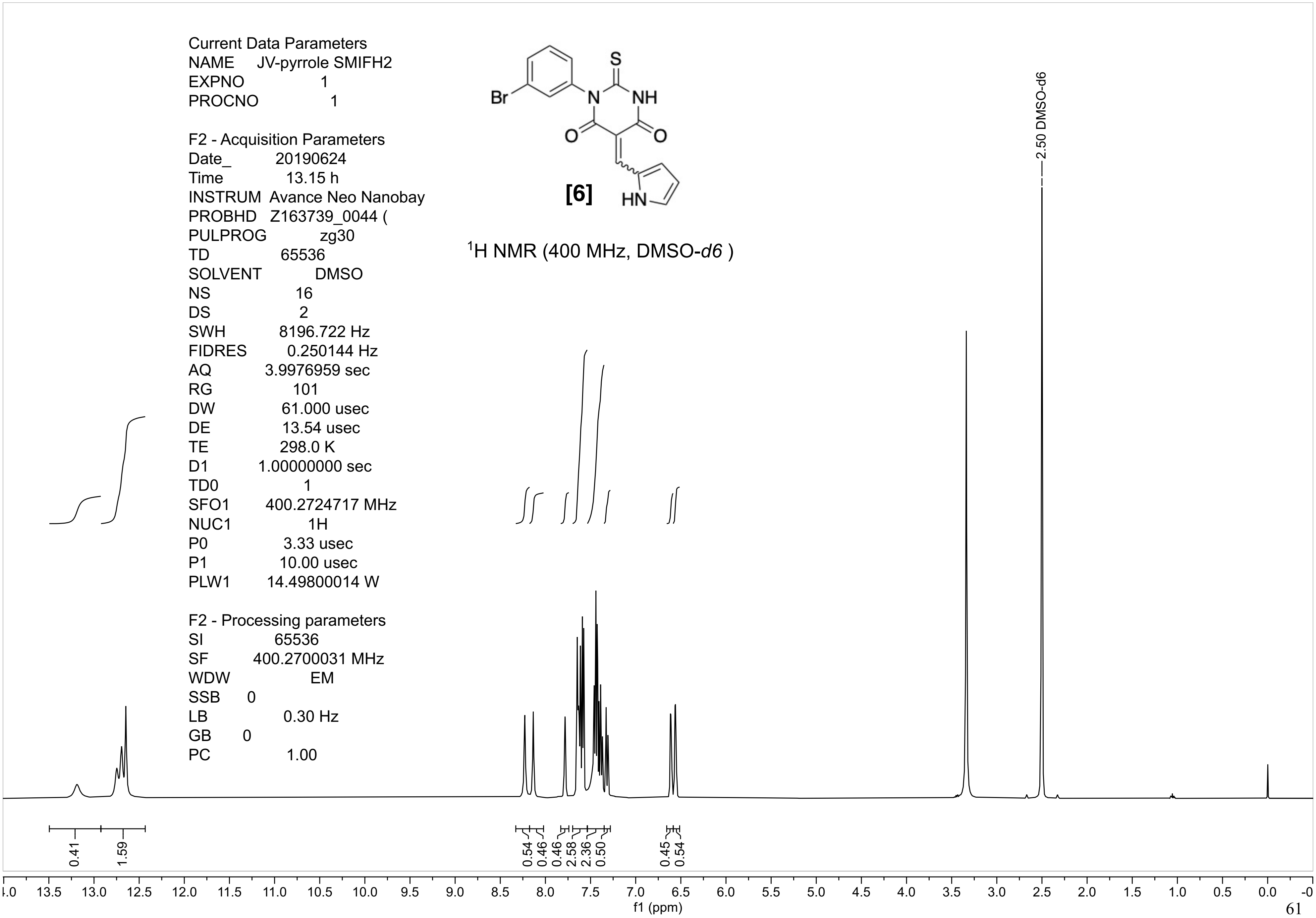
Current Data Parameters
NAME JV-pyrrole SMIFH2
EXPNO 1
PROCNO 1
F2 - Acquisition Parameters
Date_ 20190624
Time 13.15 h
INSTRUM Avance Neo Nanobay
PROBHD Z163739_0044 (
PULPROG zg30
TD 65536
SOLVENT DMSO
NS 16
DS 2
SWH 8196.722 Hz
FIDRES 0.250144 Hz
AQ 3.9976959 sec
RG 101
DW 61.000 usec
DE 13.54 usec
TE 298.0 K
D1 1.00000000 sec
TD0 1
SFO1 400.2724717 MHz
NUC1 1H
P0 3.33 usec
P1 10.00 usec
PLW1 14.49800014 W
F2 - Processing parameters
SI 65536
SF 400.2700031 MHz
WDW EM
SSB 0
LB 0.30 Hz
GB 0
PC 1.00
Br
S
N
NH
O
O
HN
[6]
$^1$H NMR (400 MHz, DMSO-*d6* )
2.50 DMSO-d6
0.41
1.59
0.54
0.46
0.46
2.58
2.36
0.50
0.45
0.54
13.5 13.0 12.5 12.0 11.5 11.0 10.5 10.0 9.5 9.0 8.5 8.0 7.5 7.0 6.5 6.0 5.5 5.0 4.5 4.0 3.5 3.0 2.5 2.0 1.5 1.0 0.5 0.0
f1 (ppm)

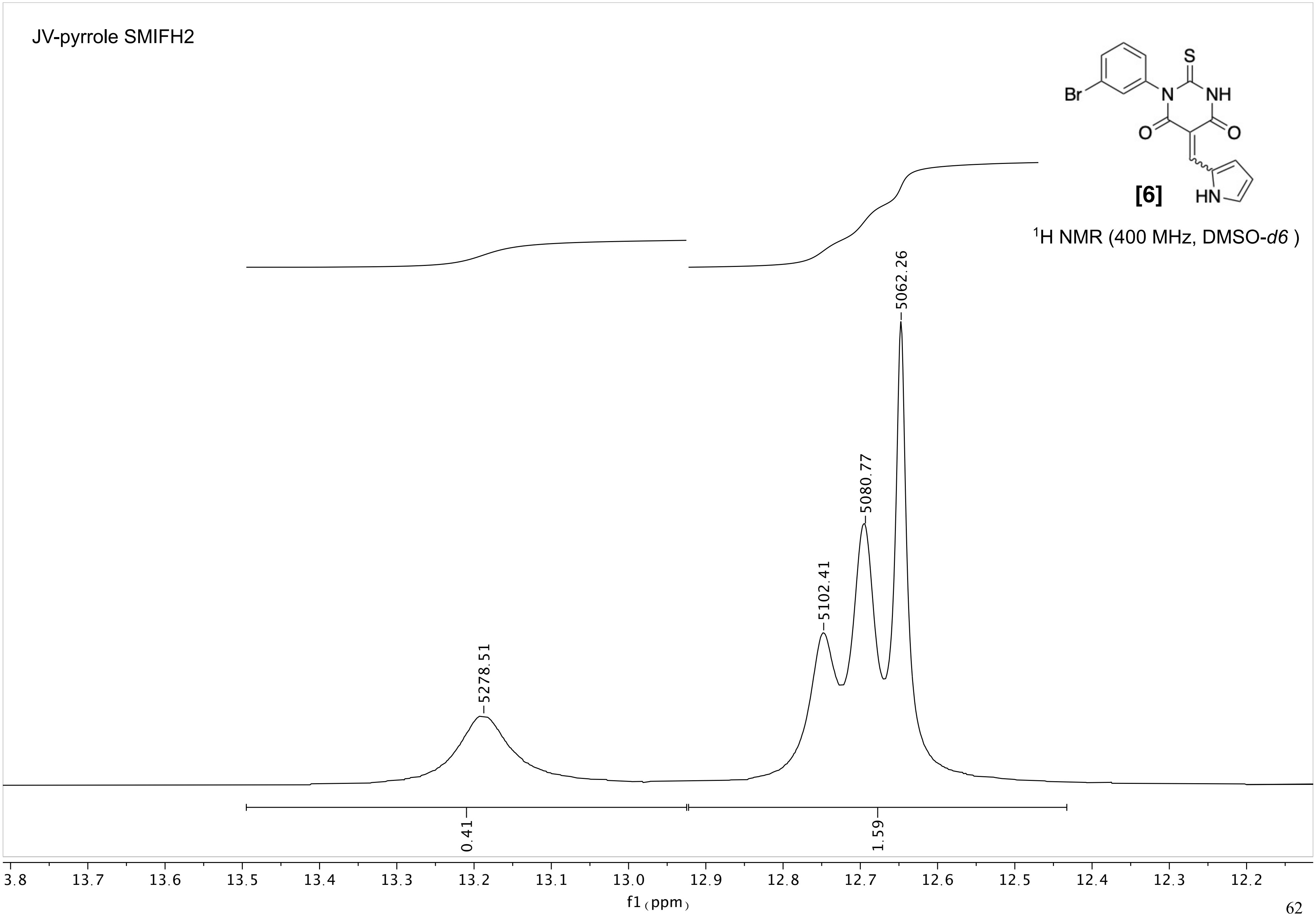
JV-pyrrole SMIFH2
Br
N
S
NH
O
O
HN
[6]
¹H NMR (400 MHz, DMSO-d6 )
5278.51
5102.41
5080.77
5062.26
0.41
1.59
3.8
13.7
13.6
13.5
13.4
13.3
13.2
13.1
13.0
12.9
12.8
12.7
12.6
12.5
12.4
12.3
12.2
f1 (ppm)

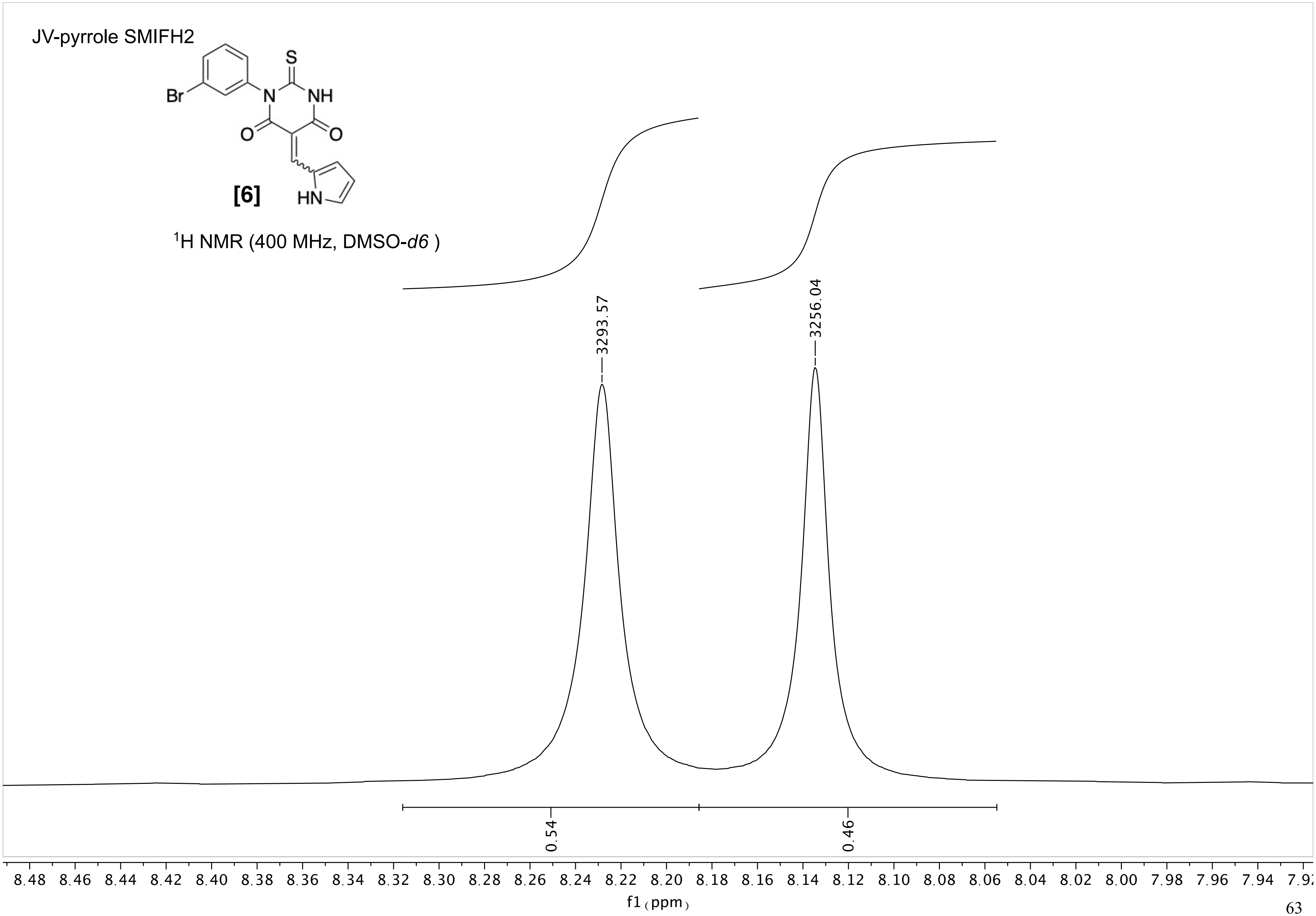
JV-pyrrole SMIFH2
Br
N
S
NH
O
O
HN
[6]
1H NMR (400 MHz, DMSO-d6 )
3293.57
3256.04
0.54
0.46
8.48
8.46
8.44
8.42
8.40
8.38
8.36
8.34
8.32
8.30
8.28
8.26
8.24
8.22
8.20
8.18
8.16
8.14
8.12
8.10
8.08
8.06
8.04
8.02
8.00
7.98
7.96
7.94
f1 (ppm)

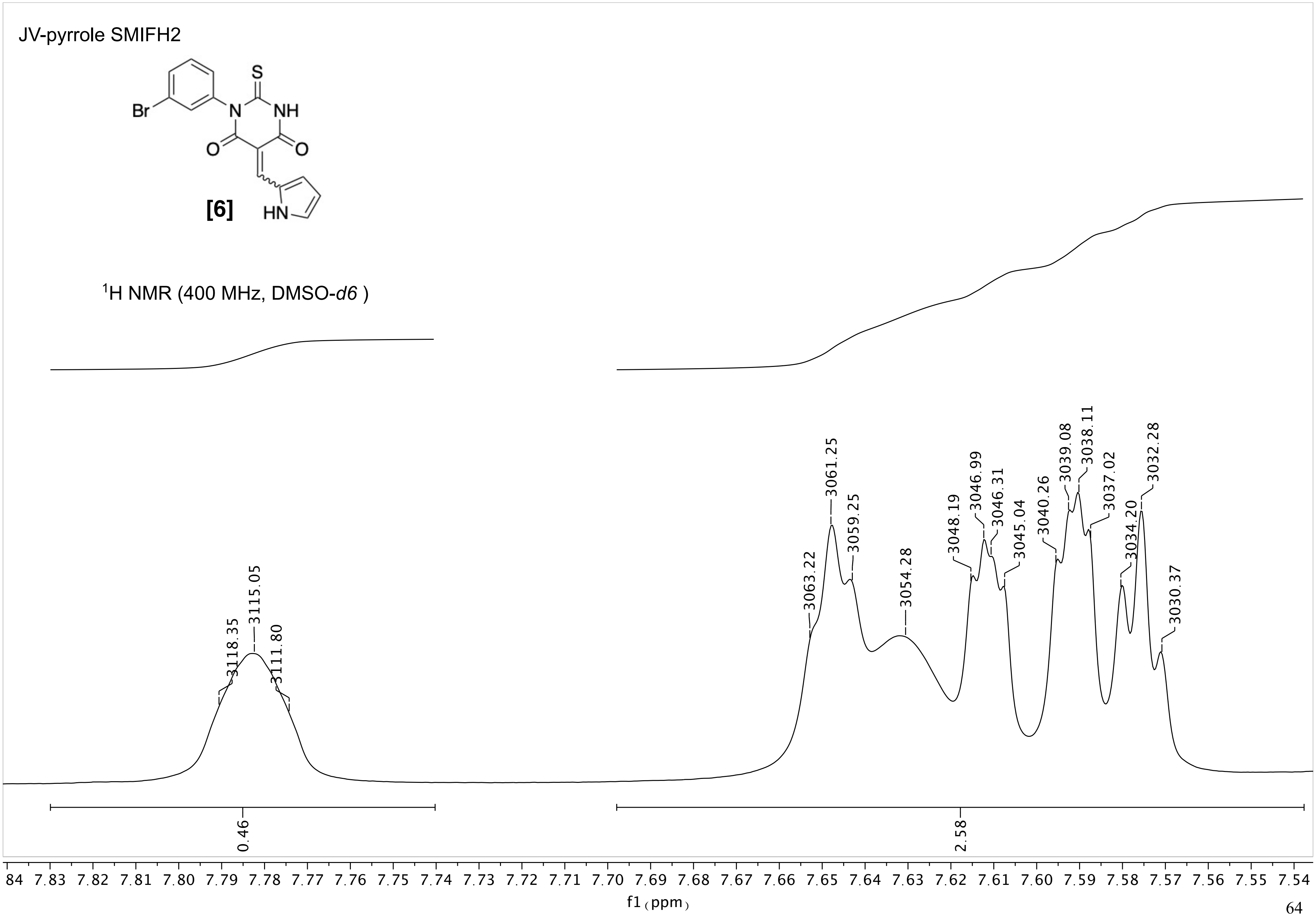
JV-pyrrole SMIFH2
S
NH
Br
N
O
O
HN
[6]
$^1$H NMR (400 MHz, DMSO-*d6* )
3118.35
3115.05
3111.80
3063.22
3061.25
3059.25
3054.28
3048.19
3046.99
3046.31
3045.04
3040.26
3039.08
3038.11
3037.02
3034.20
3032.28
3030.37
0.46
2.58
84 7.83 7.82 7.81 7.80 7.79 7.78 7.77 7.76 7.75 7.74 7.73 7.72 7.71 7.70 7.69 7.68 7.67 7.66 7.65 7.64 7.63 7.62 7.61 7.60 7.59 7.58 7.57 7.56 7.55 7.54
f1 (ppm)

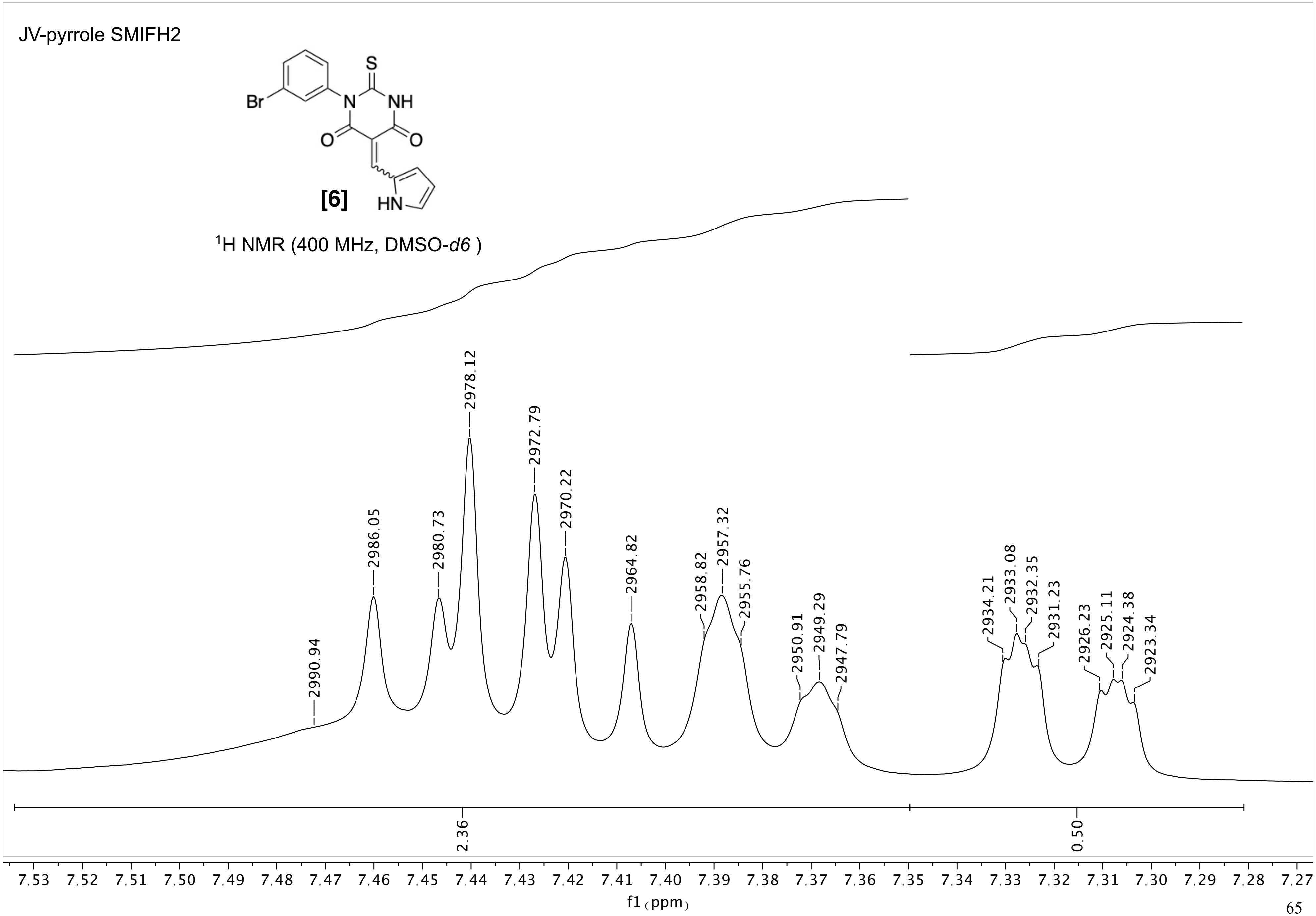
JV-pyrrole SMIFH2
[6]
$^1$H NMR (400 MHz, DMSO-*d6* )
2990.94
2986.05
2980.73
2978.12
2972.79
2970.22
2964.82
2958.82
2957.32
2955.76
2950.91
2949.29
2947.79
2934.21
2933.08
2932.35
2931.23
2926.23
2925.11
2924.38
2923.34
2.36
0.50
7.53 7.52 7.51 7.50 7.49 7.48 7.47 7.46 7.45 7.44 7.43 7.42 7.41 7.40 7.39 7.38 7.37 7.36 7.35 7.34 7.33 7.32 7.31 7.30 7.29 7.28 7.27
f1 (ppm)

JV-pyrrole SMIFH2

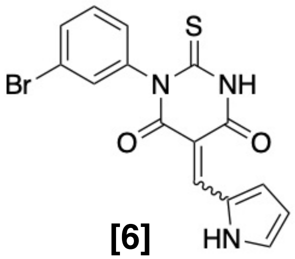

**[6]**

$^{1}$H NMR (400 MHz, DMSO-*d6* )

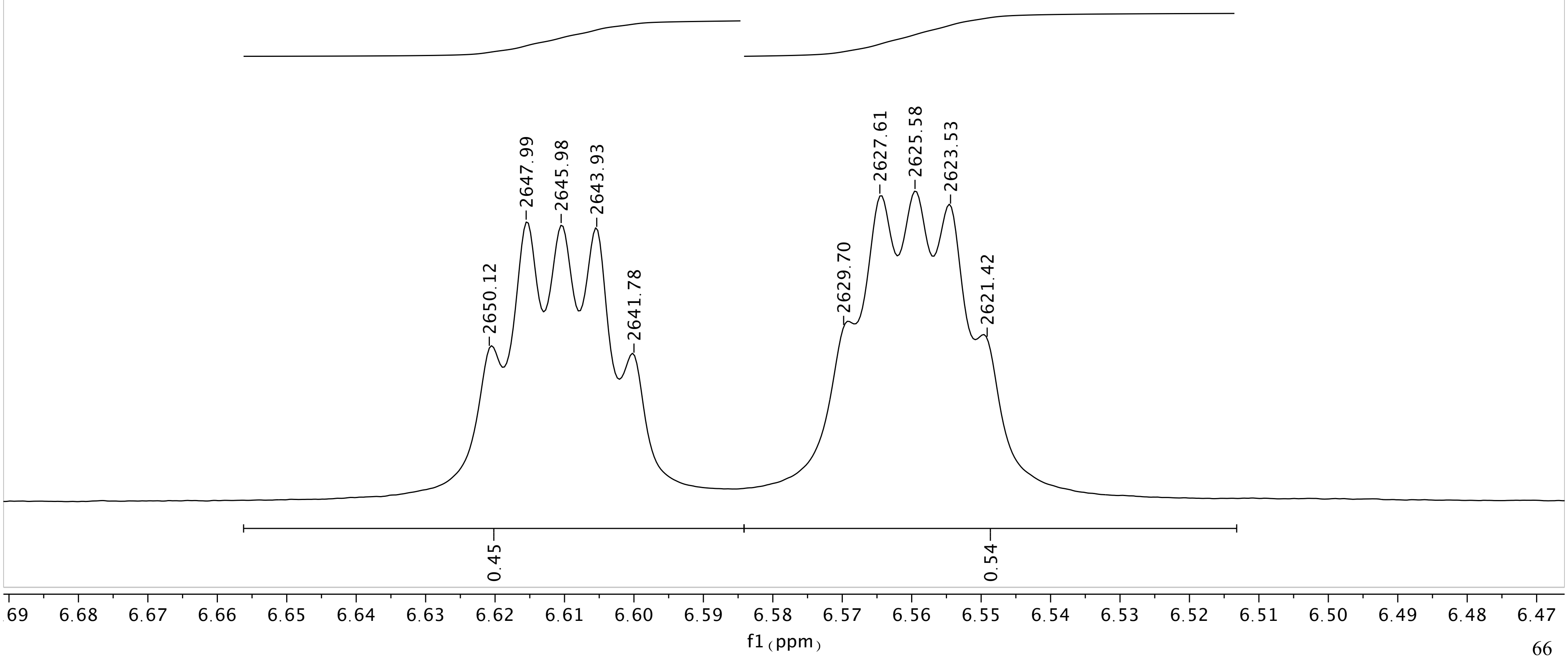

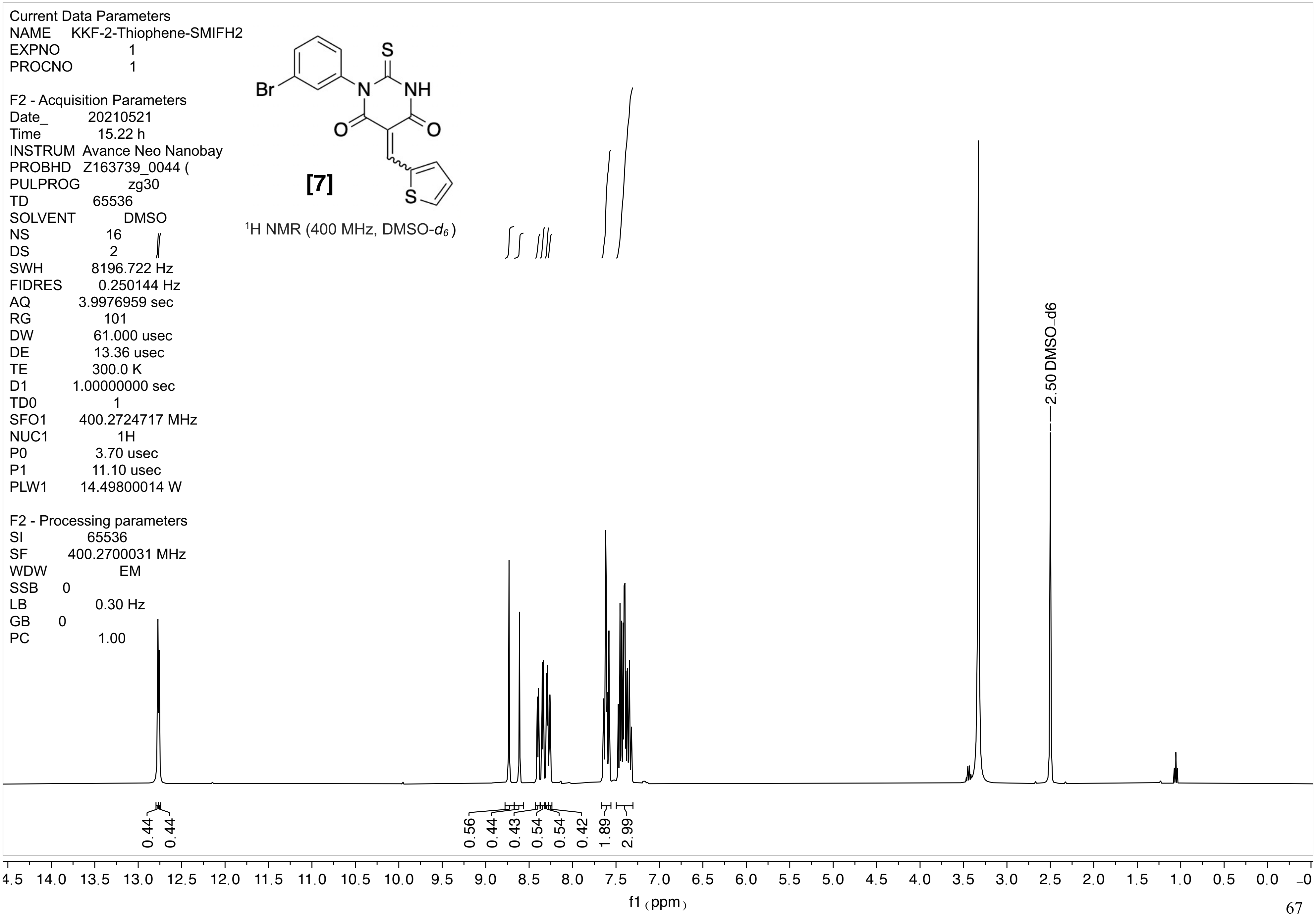
Current Data Parameters
NAME KKF-2-Thiophene-SMIFH2
EXPNO 1
PROCNO 1
F2 - Acquisition Parameters
Date_ 20210521
Time 15.22 h
INSTRUM Avance Neo Nanobay
PROBHD Z163739_0044 (
PULPROG zg30
TD 65536
SOLVENT DMSO
NS 16
DS 2
SWH 8196.722 Hz
FIDRES 0.250144 Hz
AQ 3.9976959 sec
RG 101
DW 61.000 usec
DE 13.36 usec
TE 300.0 K
D1 1.00000000 sec
TD0 1
SFO1 400.2724717 MHz
NUC1 1H
P0 3.70 usec
P1 11.10 usec
PLW1 14.49800014 W
F2 - Processing parameters
SI 65536
SF 400.2700031 MHz
WDW EM
SSB 0
LB 0.30 Hz
GB 0
PC 1.00
Br
N
NH
S
O
O
S
[7]
1H NMR (400 MHz, DMSO-d6)
2.50 DMSO_d6
0.44
0.44
0.56
0.44
0.43
0.54
0.54
0.42
1.89
2.99
4.5 14.0 13.5 13.0 12.5 12.0 11.5 11.0 10.5 10.0 9.5 9.0 8.5 8.0 7.5 7.0 6.5 6.0 5.5 5.0 4.5 4.0 3.5 3.0 2.5 2.0 1.5 1.0 0.5 0.0 -0
f1 (ppm)

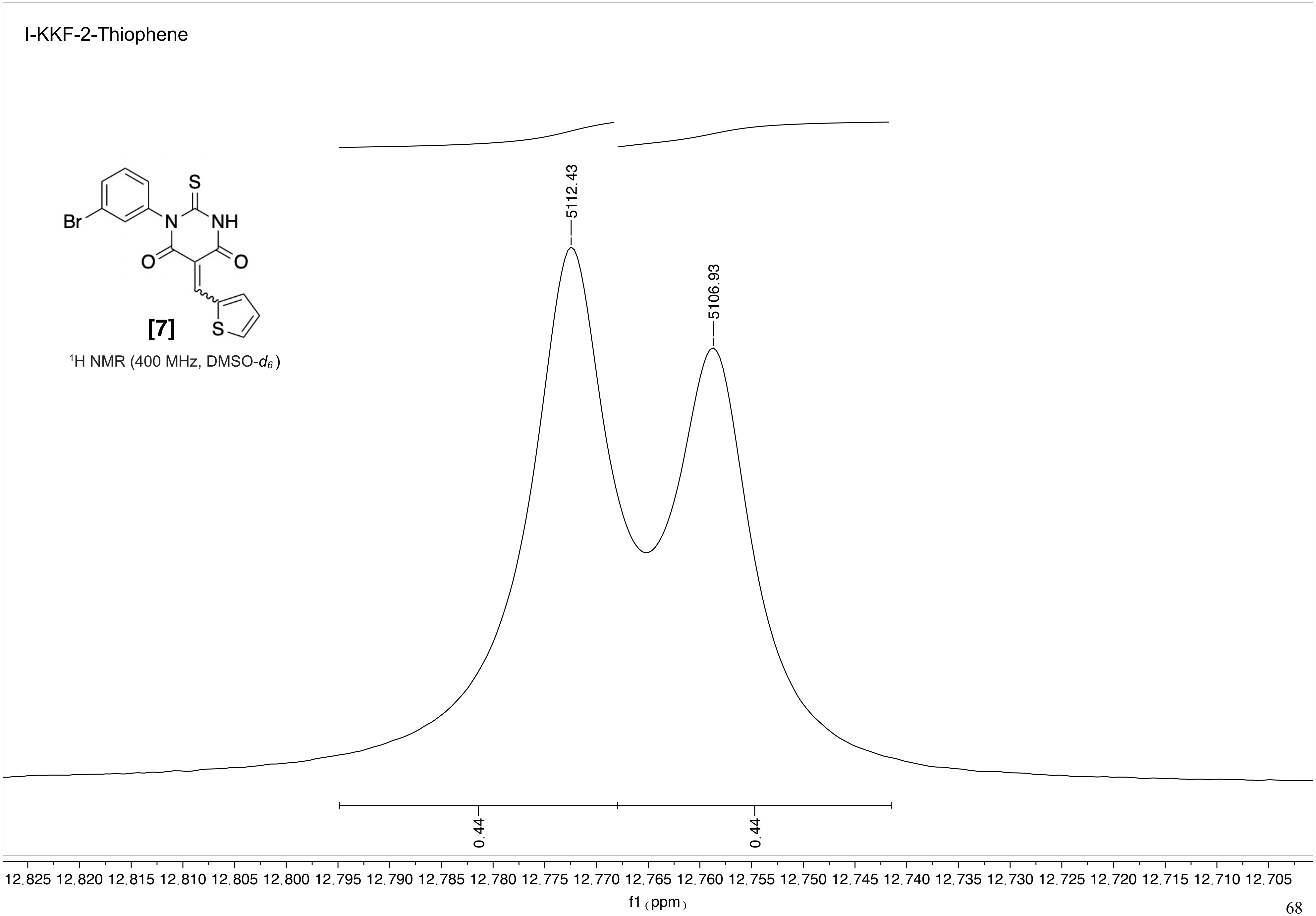

I-KKF-2-Thiophene
Br
N
NH
S
O
O
S
[7]
1H NMR (400 MHz, DMSO-d6)
5112.43
5106.93
0.44
0.44
12.825 12.820 12.815 12.810 12.805 12.800 12.795 12.790 12.785 12.780 12.775 12.770 12.765 12.760 12.755 12.750 12.745 12.740 12.735 12.730 12.725 12.720 12.715 12.710 12.705
f1 (ppm)

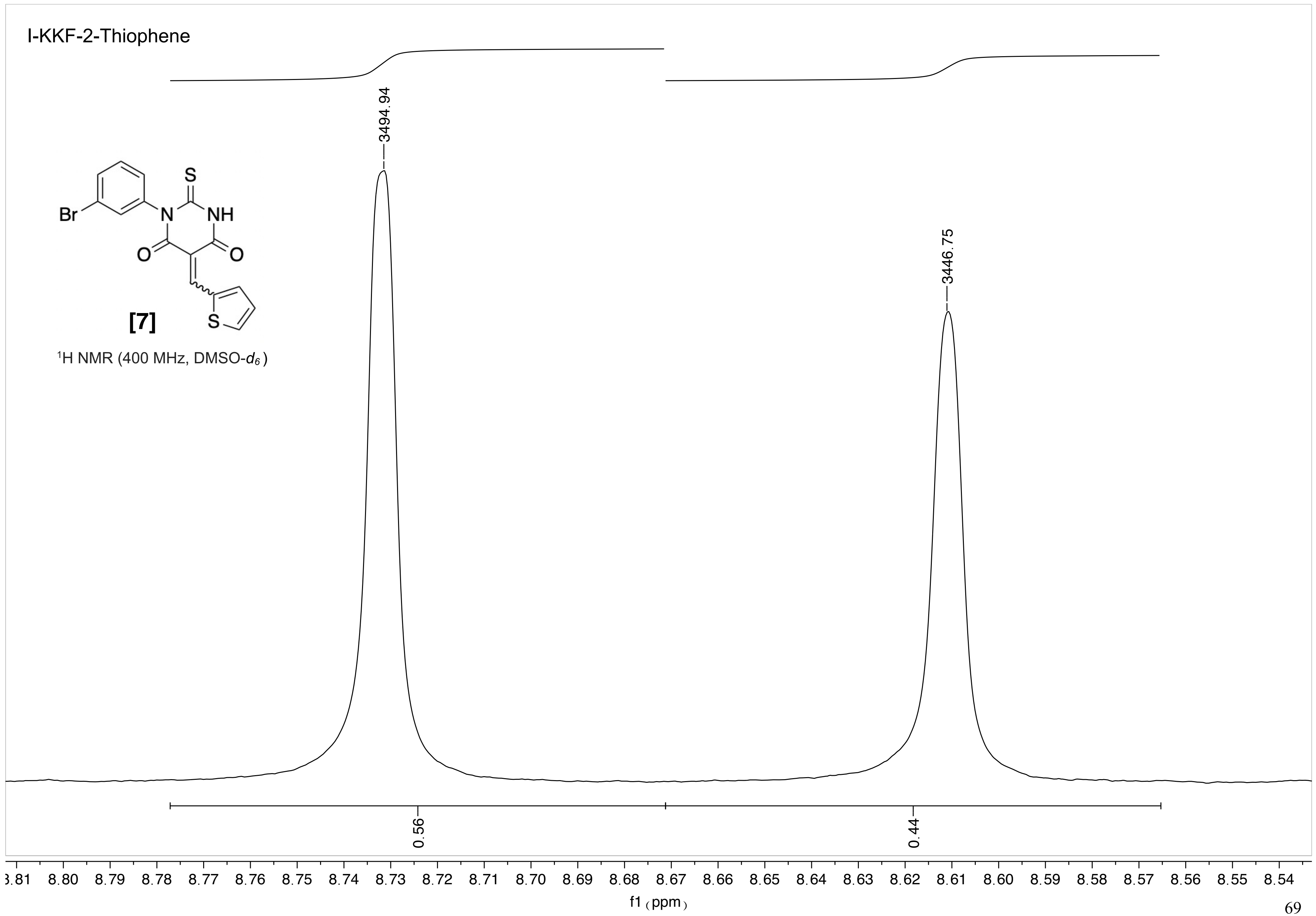
I-KKF-2-Thiophene
3494.94
3446.75
Br
N
NH
S
O
O
S
[7]
1H NMR (400 MHz, DMSO-d6)
0.56
0.44
8.81 8.80 8.79 8.78 8.77 8.76 8.75 8.74 8.73 8.72 8.71 8.70 8.69 8.68 8.67 8.66 8.65 8.64 8.63 8.62 8.61 8.60 8.59 8.58 8.57 8.56 8.55 8.54
f1 (ppm)

I-KKF-2-Thiophene

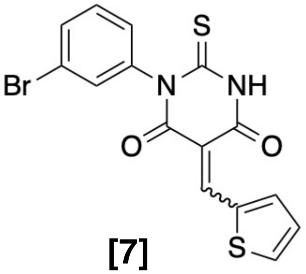

**[7]**

$^{1}$H NMR (400 MHz, DMSO-$d_6$)

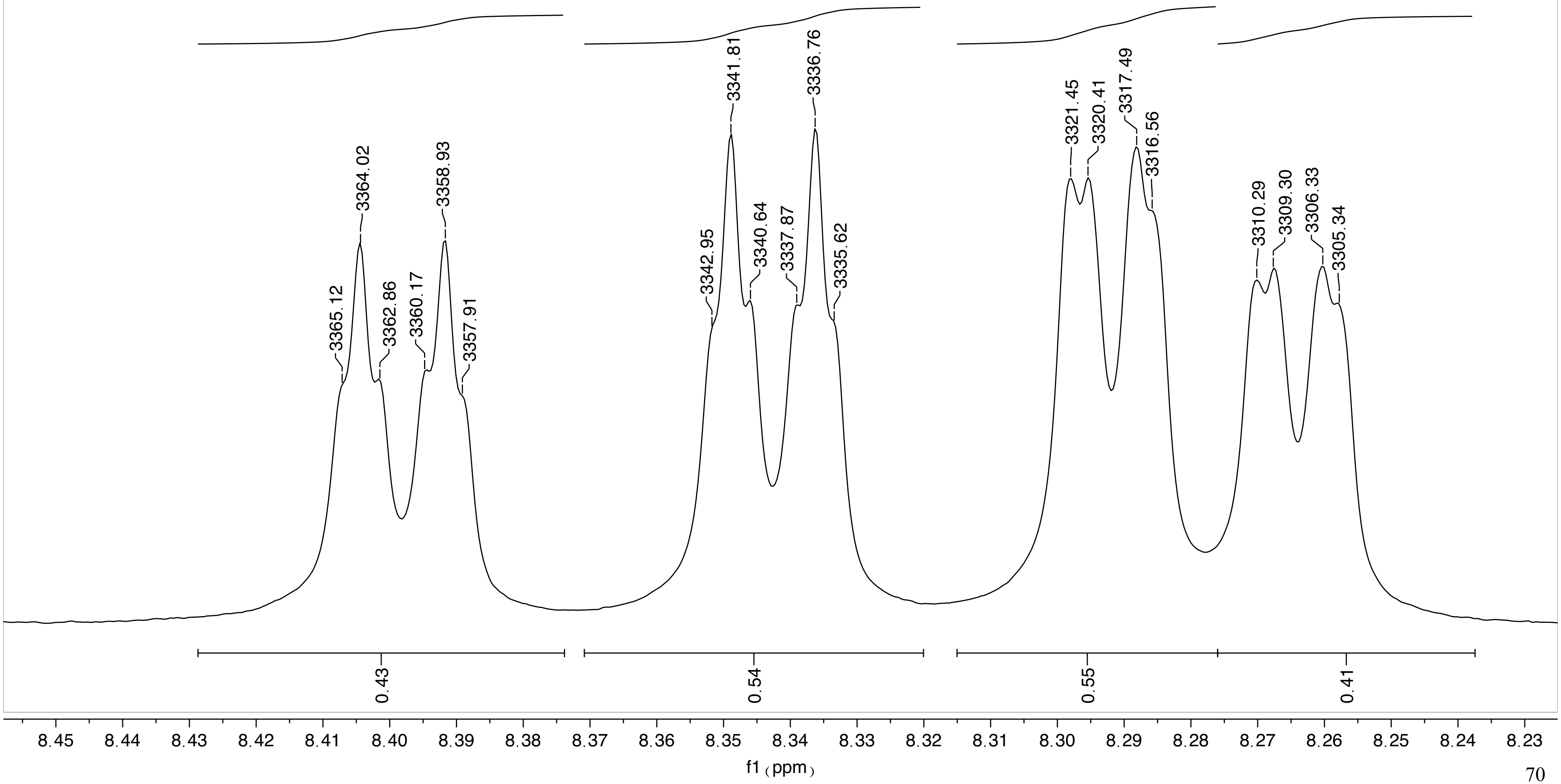

I-KKF-2-Thiophene

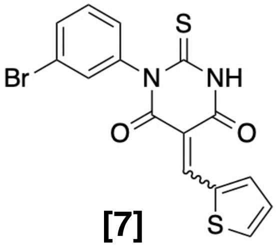

[7]

$^{1}$H NMR (400 MHz, DMSO-$d_6$)

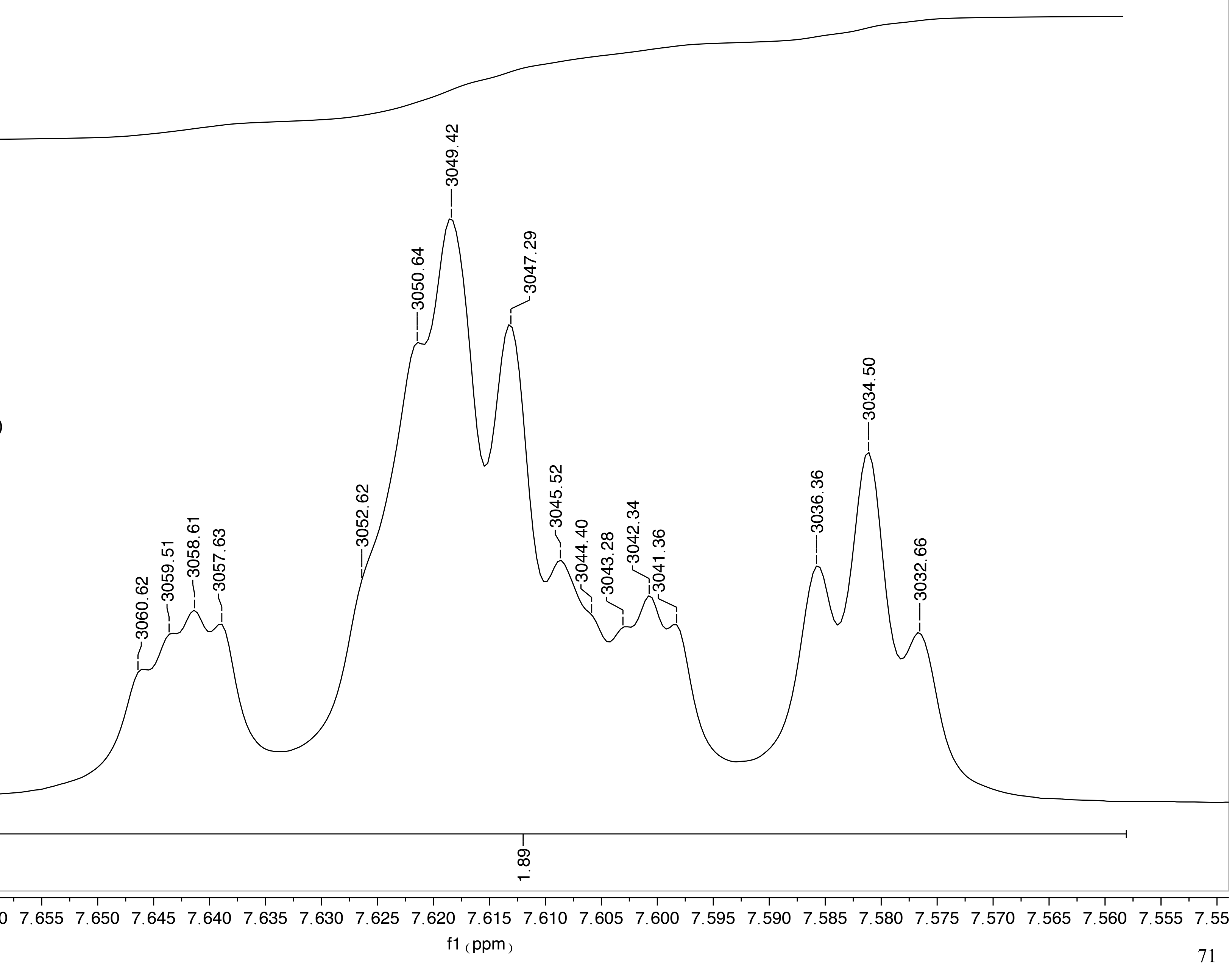

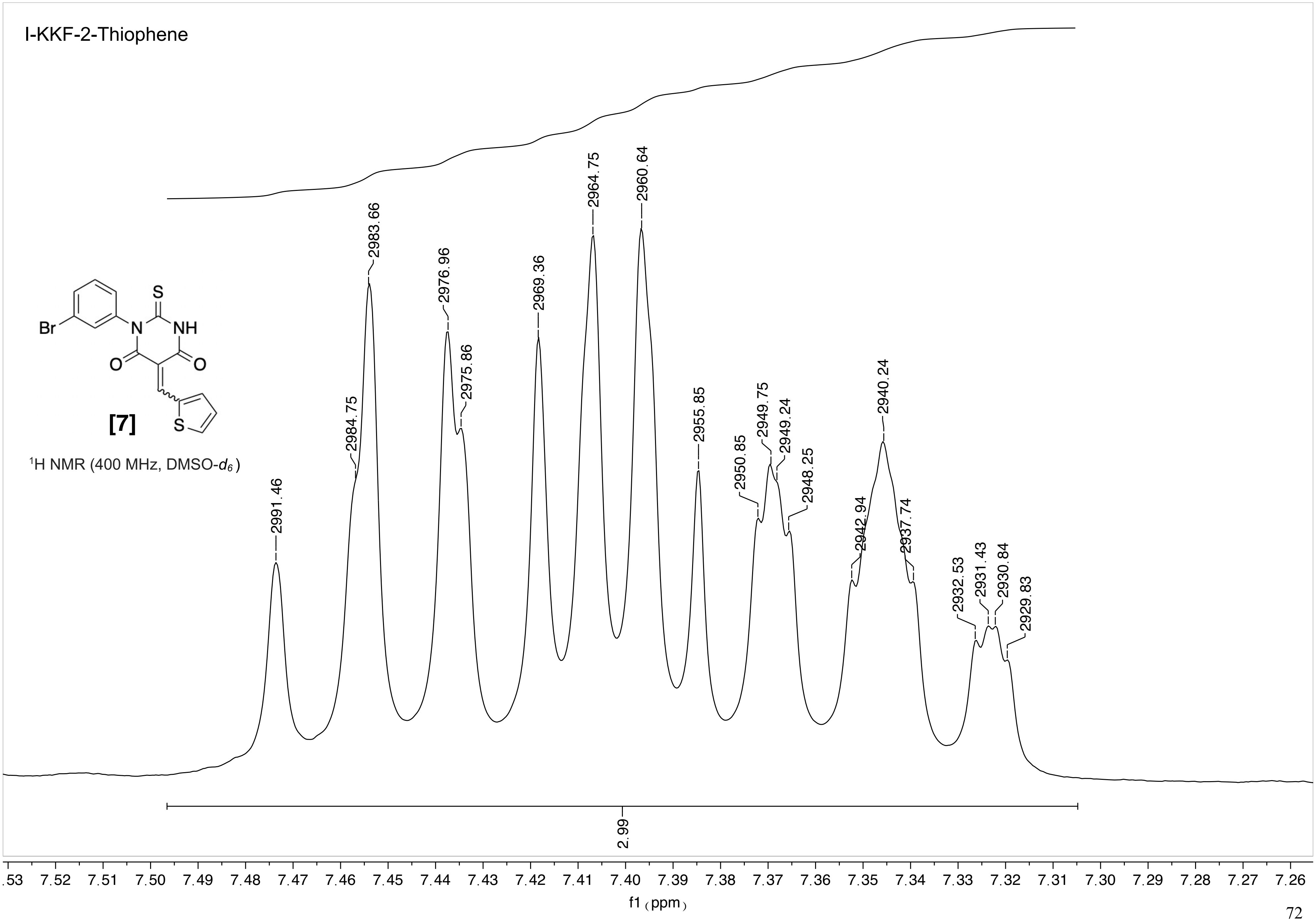

I-KKF-2-Thiophene
Br
N
NH
S
O
O
S
[7]
1H NMR (400 MHz, DMSO-d6)
2991.46
2984.75
2983.66
2976.96
2975.86
2969.36
2964.75
2960.64
2955.85
2950.85
2949.75
2949.24
2948.25
2942.94
2940.24
2937.74
2932.53
2931.43
2930.84
2929.83
2.99
7.53 7.52 7.51 7.50 7.49 7.48 7.47 7.46 7.45 7.44 7.43 7.42 7.41 7.40 7.39 7.38 7.37 7.36 7.35 7.34 7.33 7.32 7.31 7.30 7.29 7.28 7.27 7.26
f1 (ppm)

Current Data Parameters
NAME WX-I-50
EXPNO 4
PROCNO 1

F2 - Acquisition Parameters
Date_ 20191017
Time 15.50 h
INSTRUM Avance Neo
Nanobay
PROBHD Z163739_0044 (
PULPROG zg30
TD 65536
SOLVENT DMSO
NS 16
DS 2
SWH 8196.722 Hz
FIDRES 0.250144 Hz
AQ 3.9976959 sec
RG 101
DW 61.000 usec
DE 13.54 usec
TE 295.9 K
D1 1.00000000 sec
TD0 1
SFO1 400.2724717 MHz
NUC1 1H
P0 3.33 usec
P1 10.00 usec
PLW1 14.49800014 W

F2 - Processing parameters
SI 65536
SF 400.2700024 MHz
WDW EM
SSB 0
LB 0.30 Hz
GB 0
PC 1.00

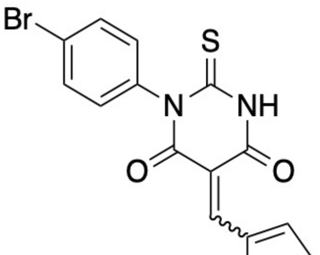

[8]

$^1$H NMR (400 MHz, DMSO-$d_6$)

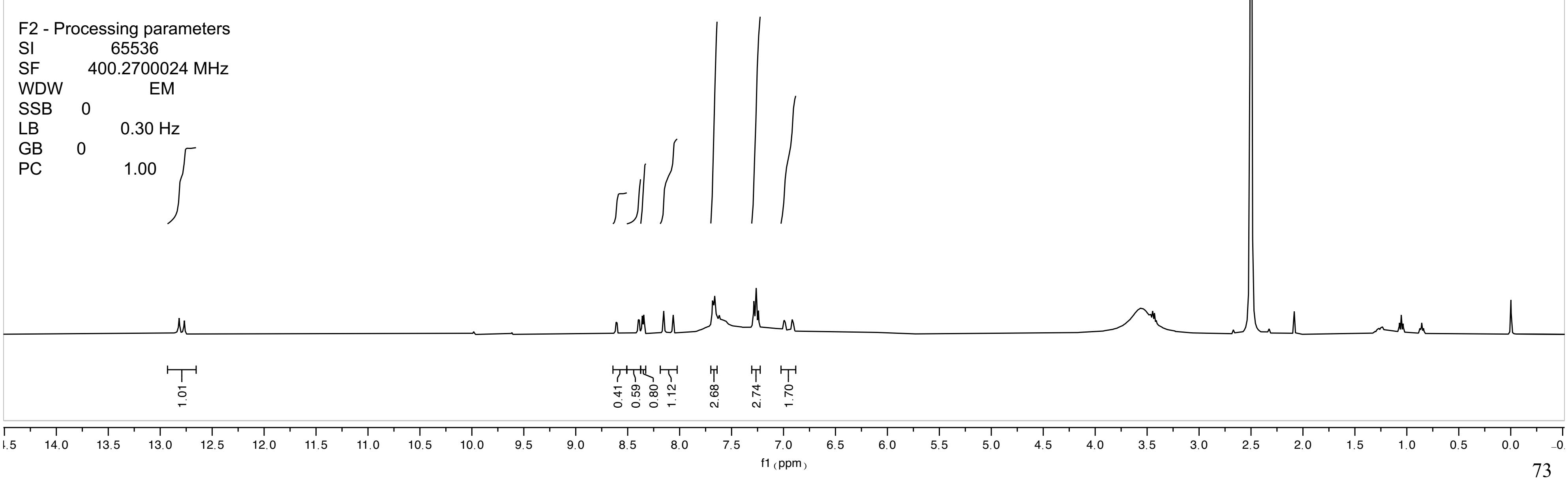

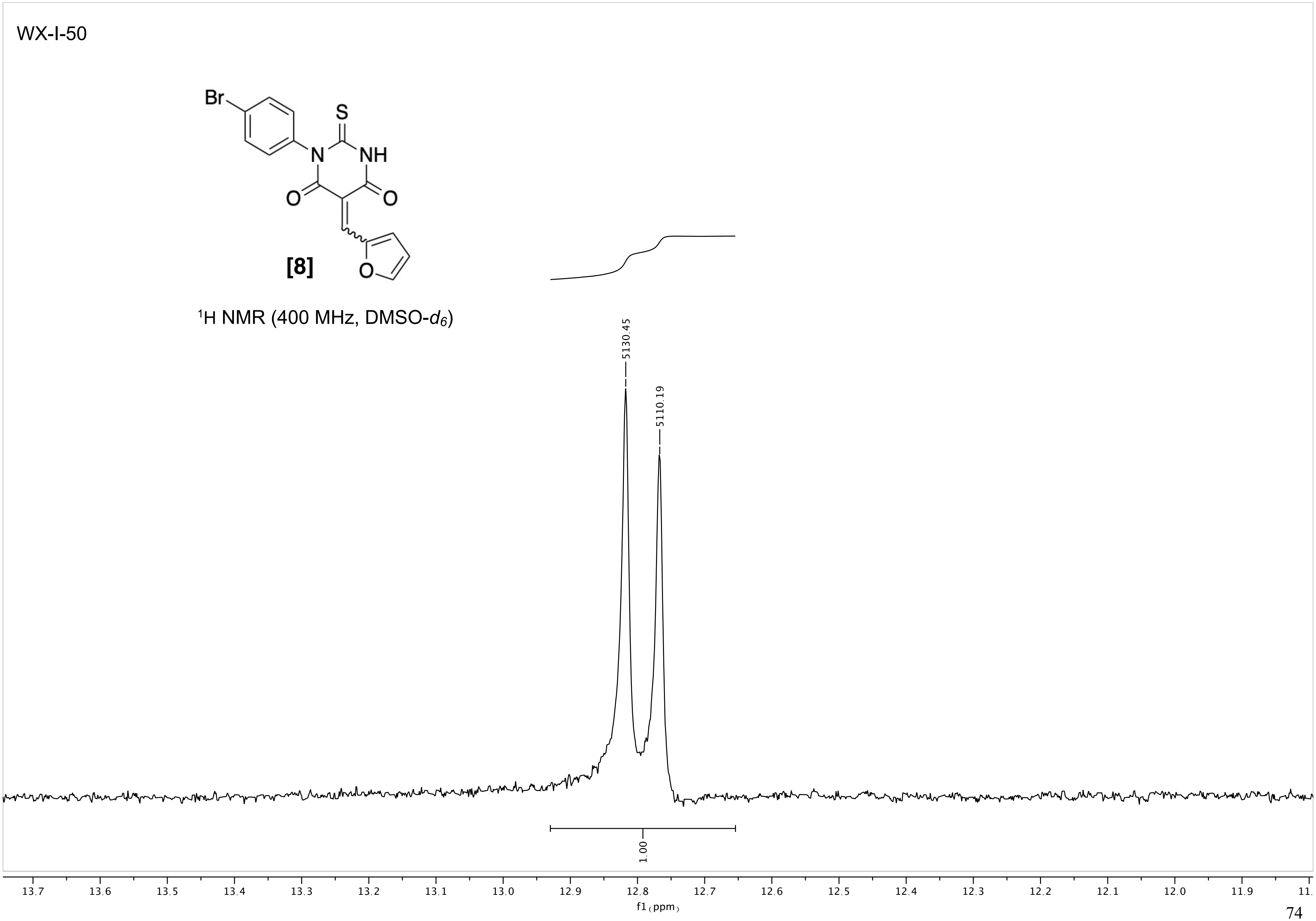
WX-I-50
Br
S
N
NH
O
O
O
[8]
¹H NMR (400 MHz, DMSO-d6)
5130.45
5110.19
1.00
13.7
13.6
13.5
13.4
13.3
13.2
13.1
13.0
12.9
12.8
12.7
12.6
12.5
12.4
12.3
12.2
12.1
12.0
11.9
f1 (ppm)

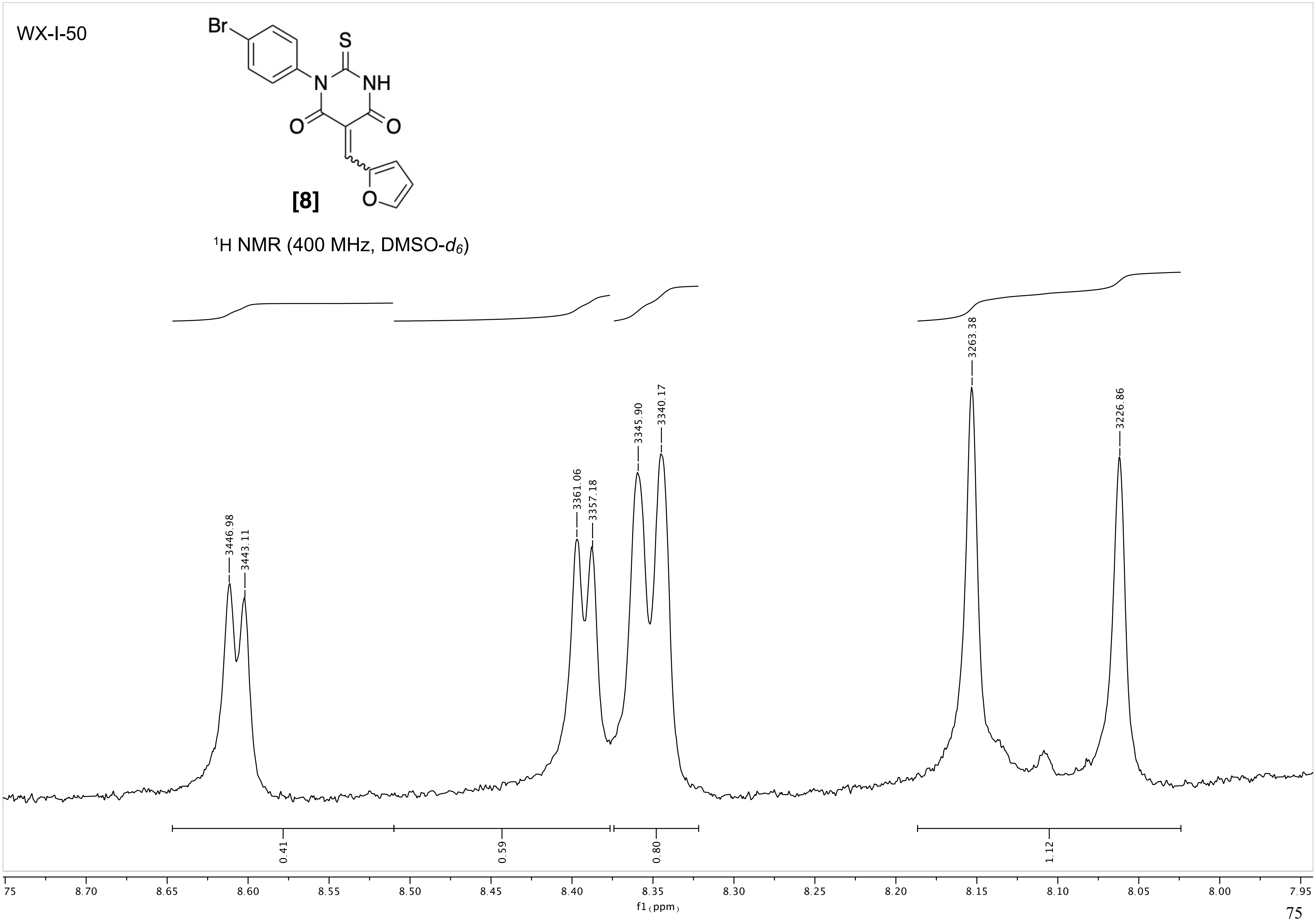
WX-I-50
[8]
$^1$H NMR (400 MHz, DMSO-$d_6$)
3446.98
3443.11
3361.06
3357.18
3345.90
3340.17
3263.38
3226.86
0.41
0.59
0.80
1.12
f1 (ppm)

WX-I-50

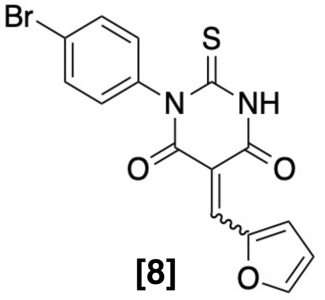

**[8]**

$^1$H NMR (400 MHz, DMSO-$d_6$)

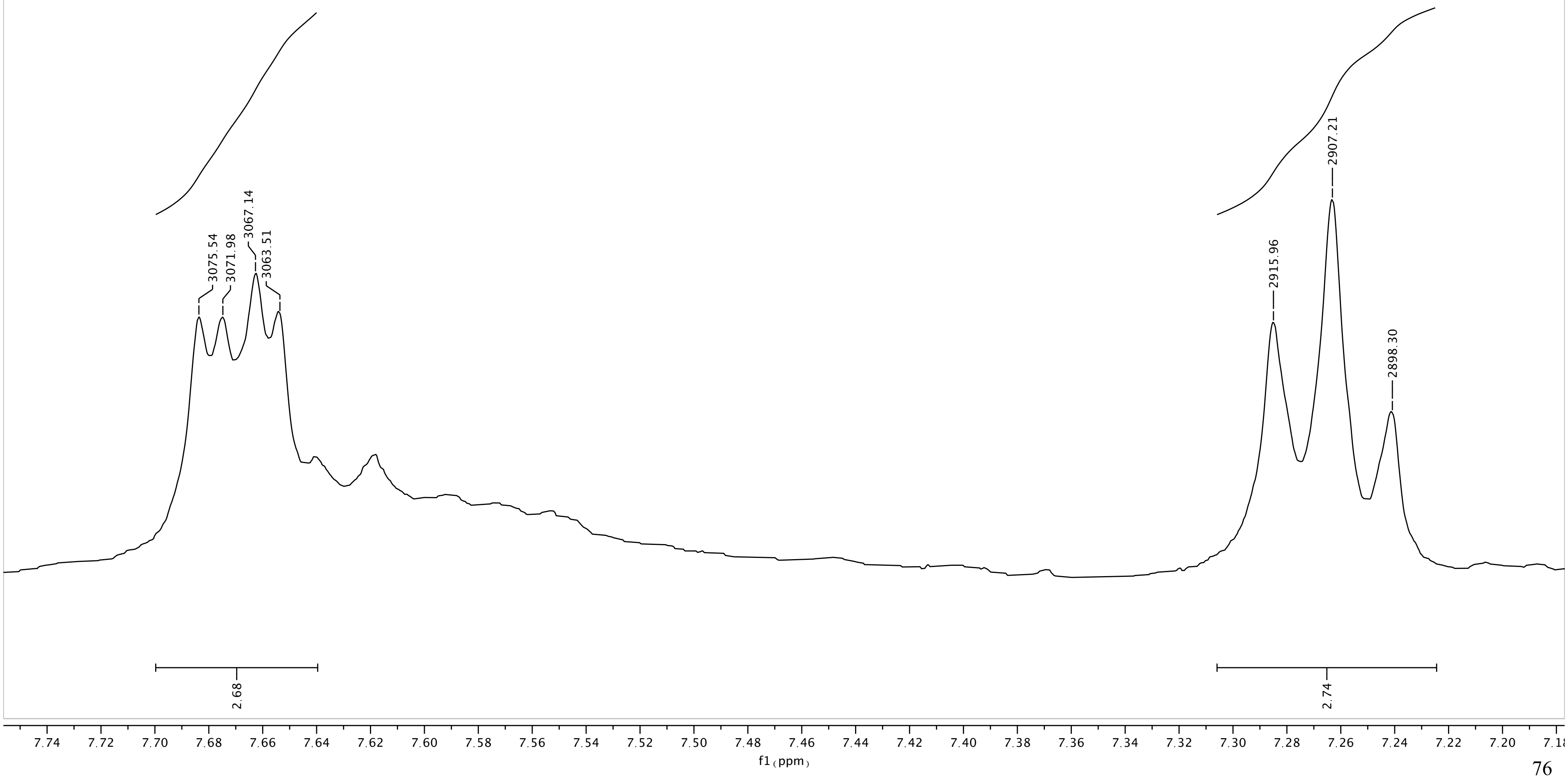

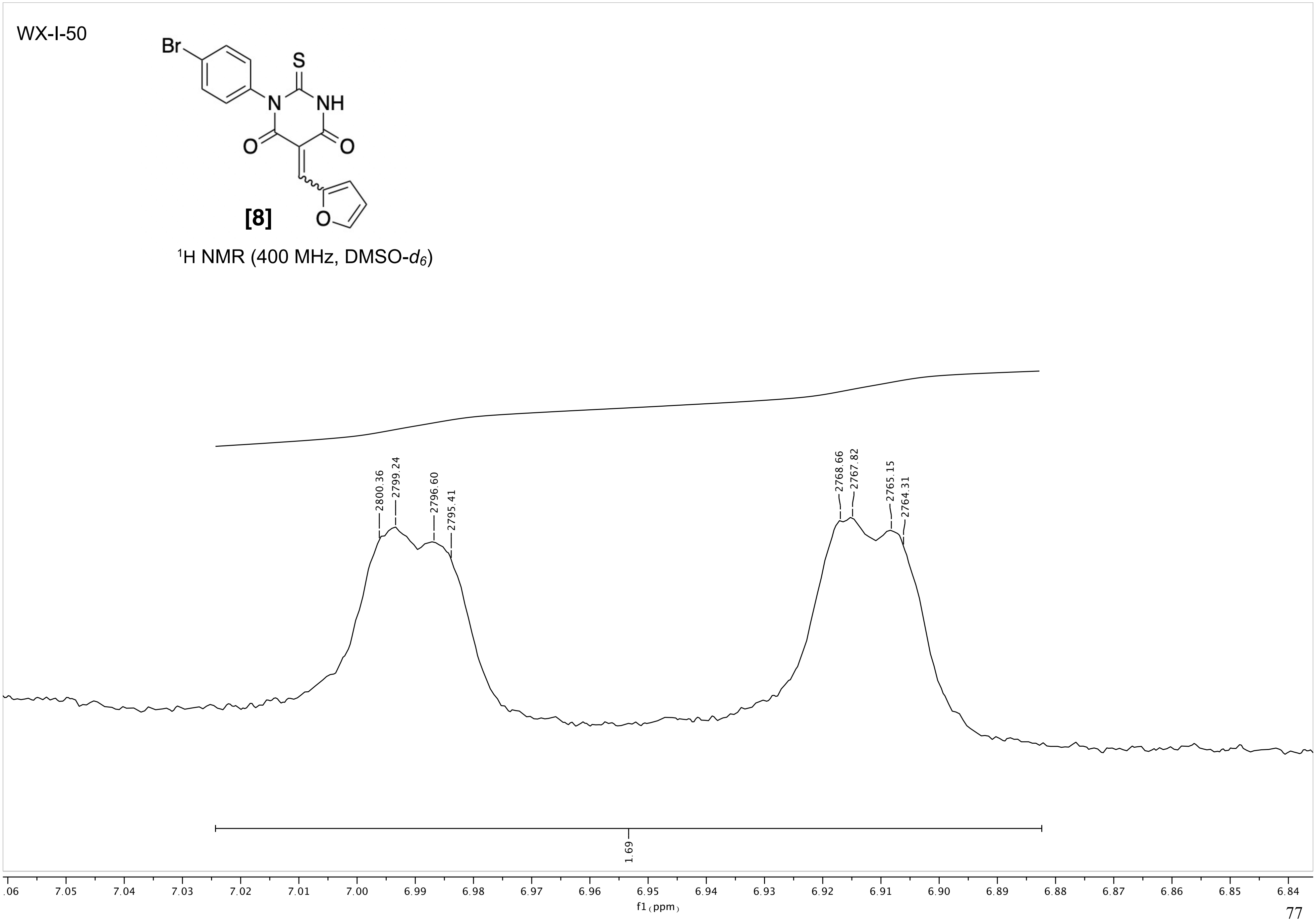
WX-I-50
Br
S
N
NH
O
O
O
[8]
1H NMR (400 MHz, DMSO-d6)
2800.36
2799.24
2796.60
2795.41
2768.66
2767.82
2765.15
2764.31
1.69
7.05
7.04
7.03
7.02
7.01
7.00
6.99
6.98
6.97
6.96
6.95
6.94
6.93
6.92
6.91
6.90
6.89
6.88
6.87
6.86
6.85
6.84
f1 (ppm)

NAME MVP-1-026
EXPNO 1
PROCNO 1
Date_ 20161103
Time 17.17
INSTRUM spect
PROBHD 5 mm BBO BB-1H
PULPROG zg30
TD 65536
SOLVENT DMSO
NS 16
DS 2
SWH 6172.839 Hz
FIDRES 0.094190 Hz
AQ 5.3084660 sec
RG 256
DW 81.000 usec
DE 6.50 usec
TE 294.1 K
D1 1.00000000 sec
TD0 1

======== CHANNEL f1 ========
NUC1 1H
P1 7.10 usec
PL1 3.00 dB
SFO1 300.1318534 MHz
SI 32768
SF 300.1300000 MHz
WDW EM
SSB 0
LB 0.30 Hz
GB 0
PC 1.00

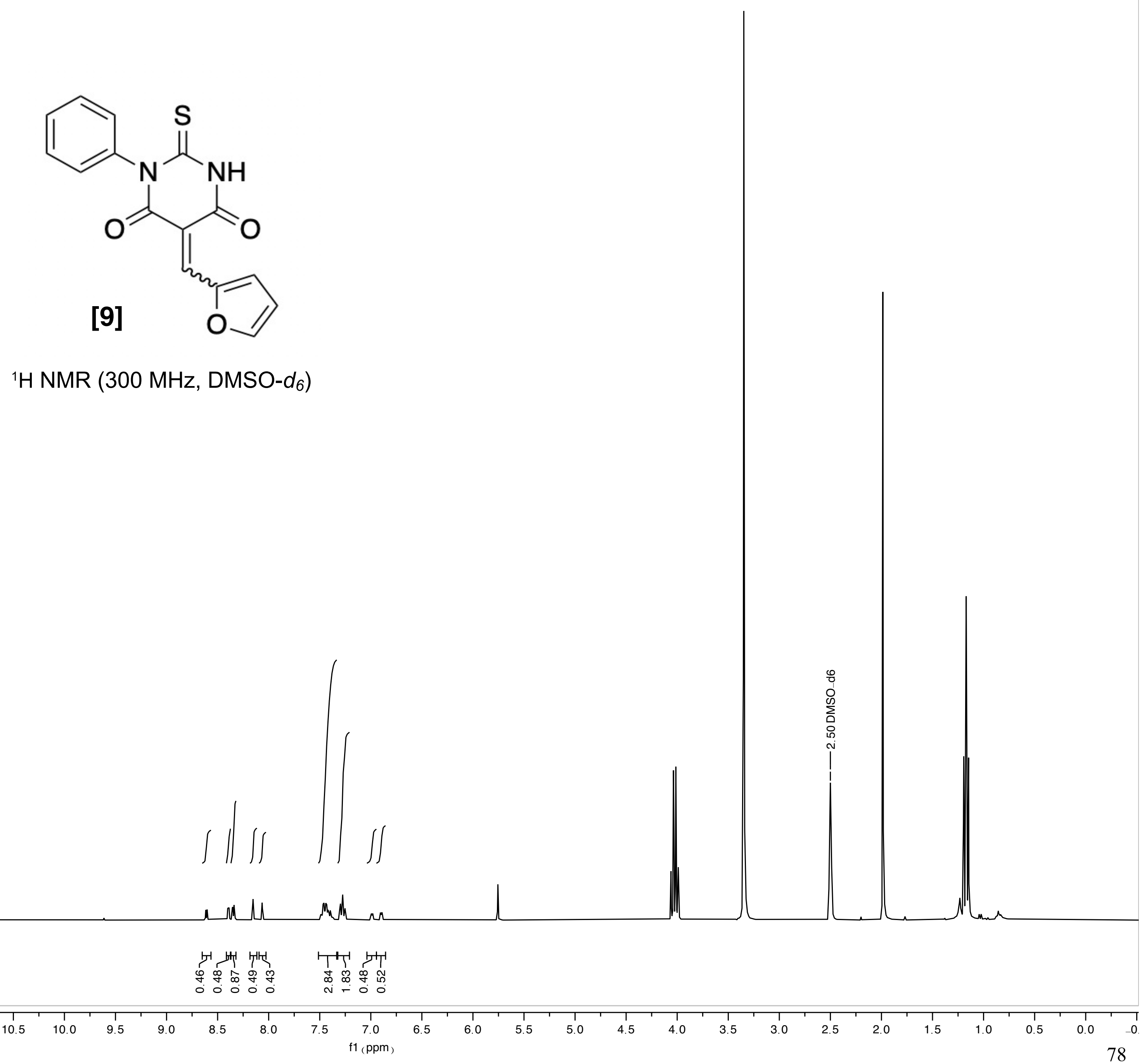

**[9]**

$^1$H NMR (300 MHz, DMSO-$d_6$)

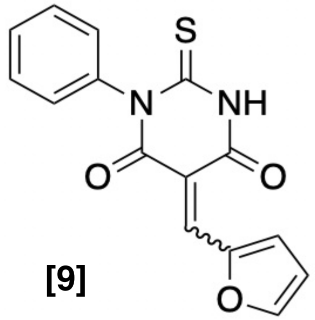

[9]

$^1$H NMR (300 MHz, DMSO-$d_6$)

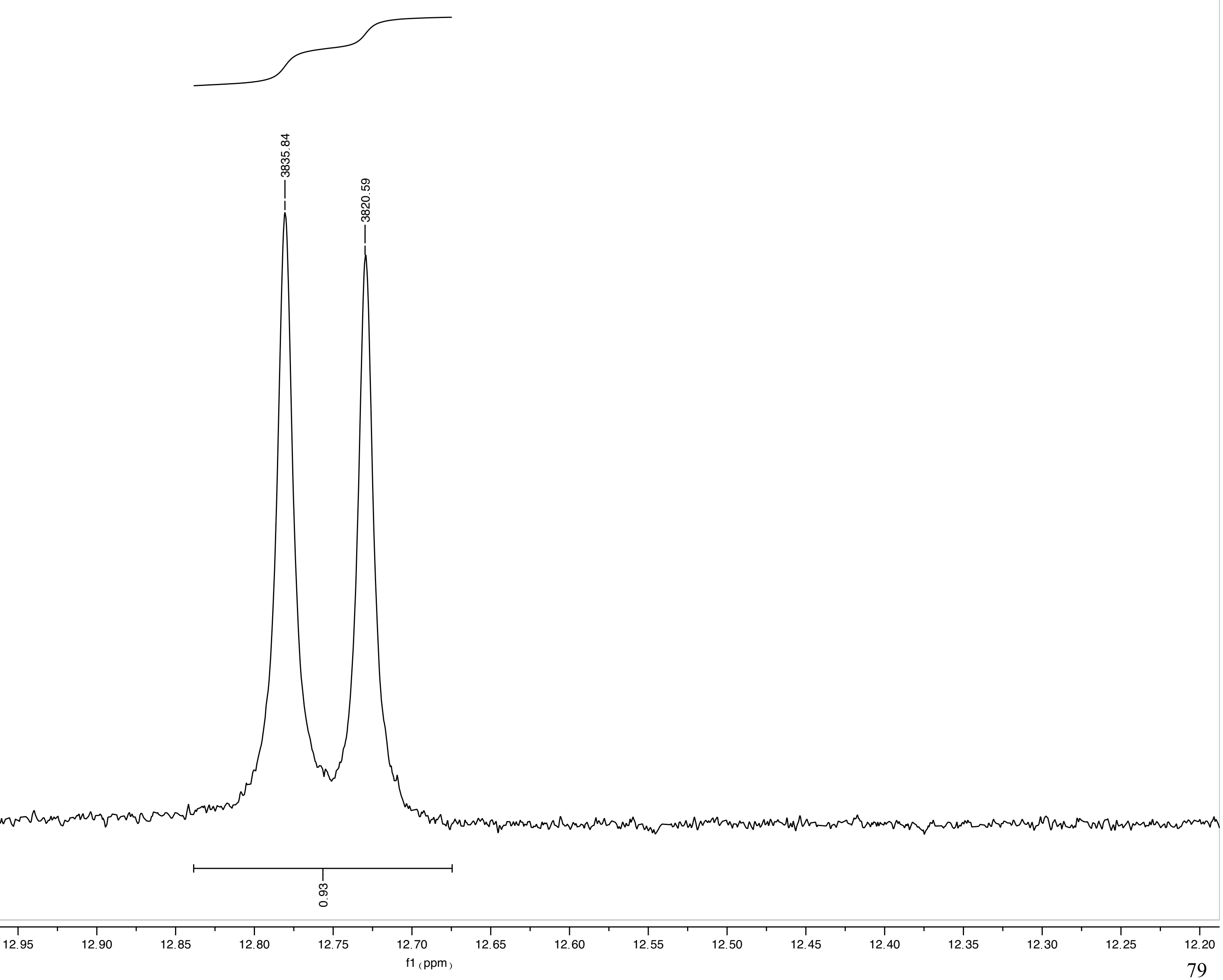

MVP-1-026

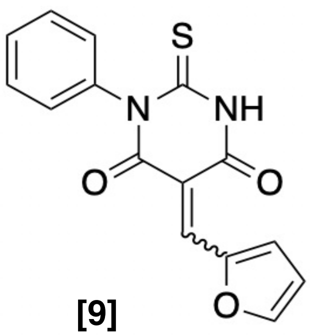

[9]

$^{1}$H NMR (300 MHz, DMSO-$d_6$)

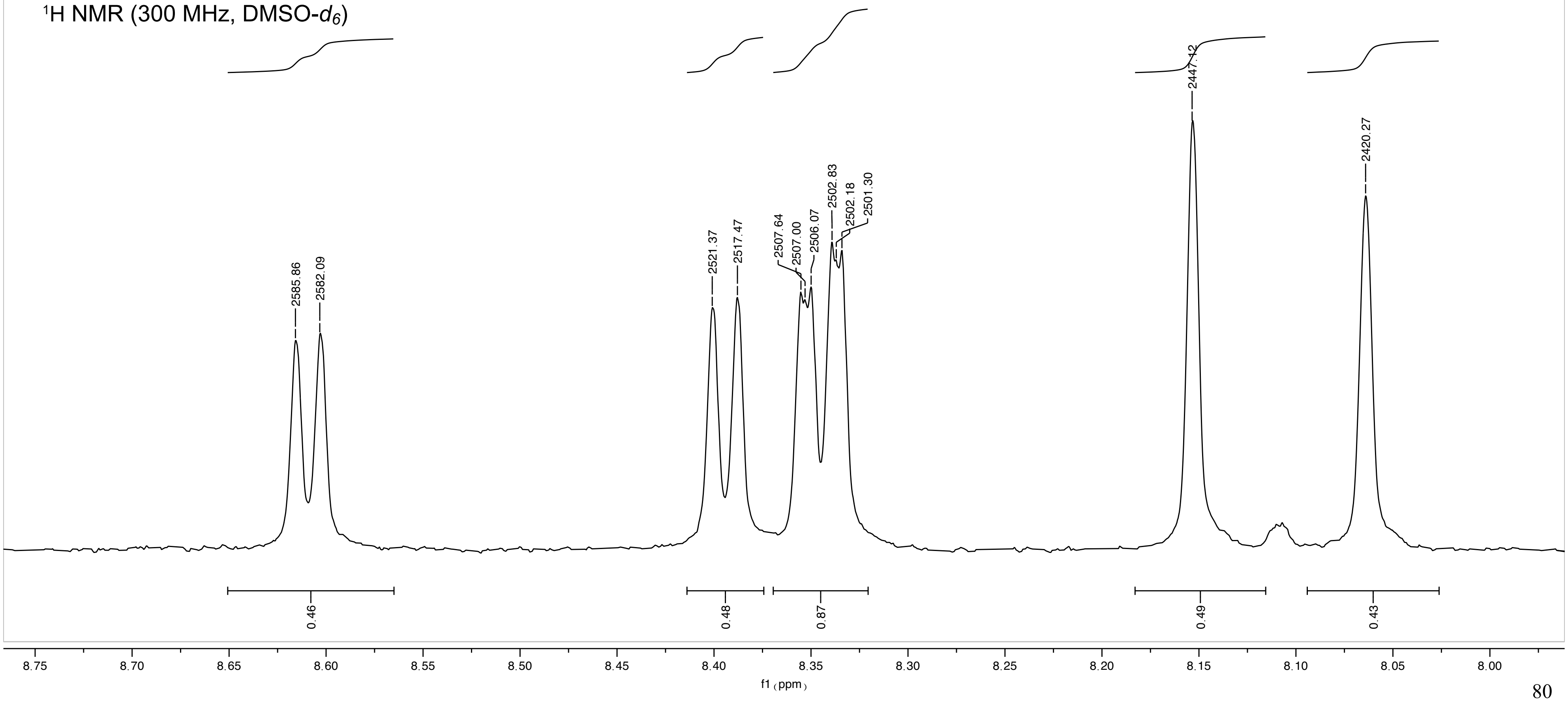

MVP-1-026

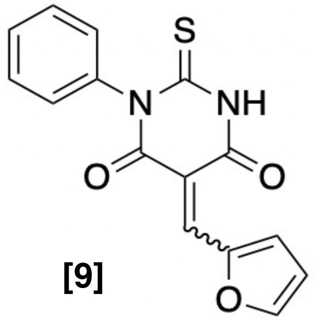

[9]

$^1$H NMR (300 MHz, DMSO-$d_6$)

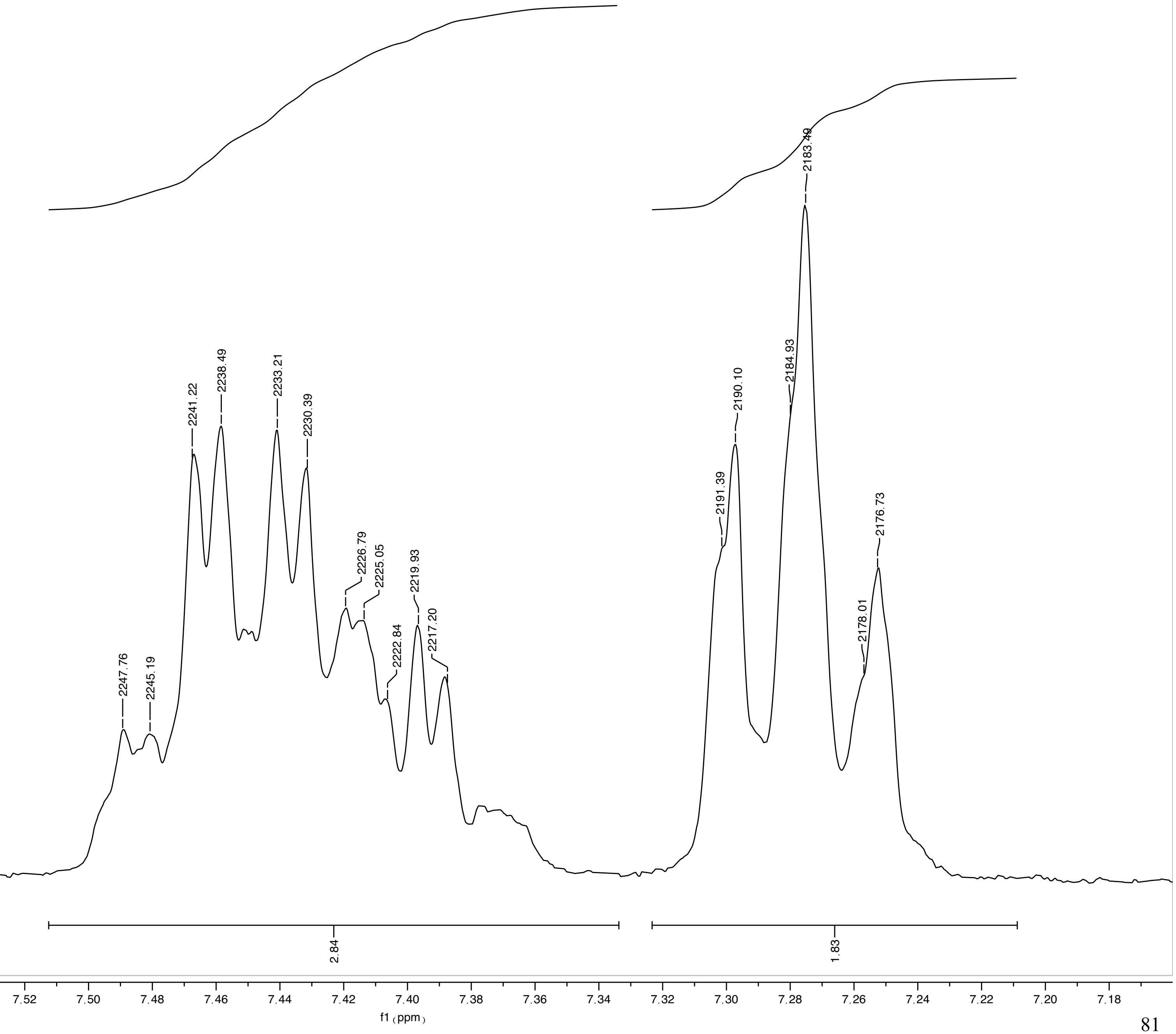

MVP-1-026

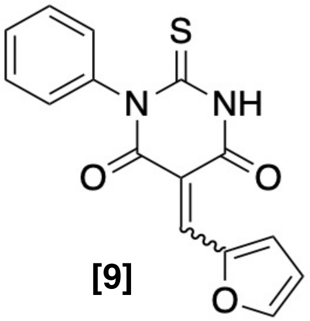

**[9]**

$^1$H NMR (300 MHz, DMSO-$d_6$)

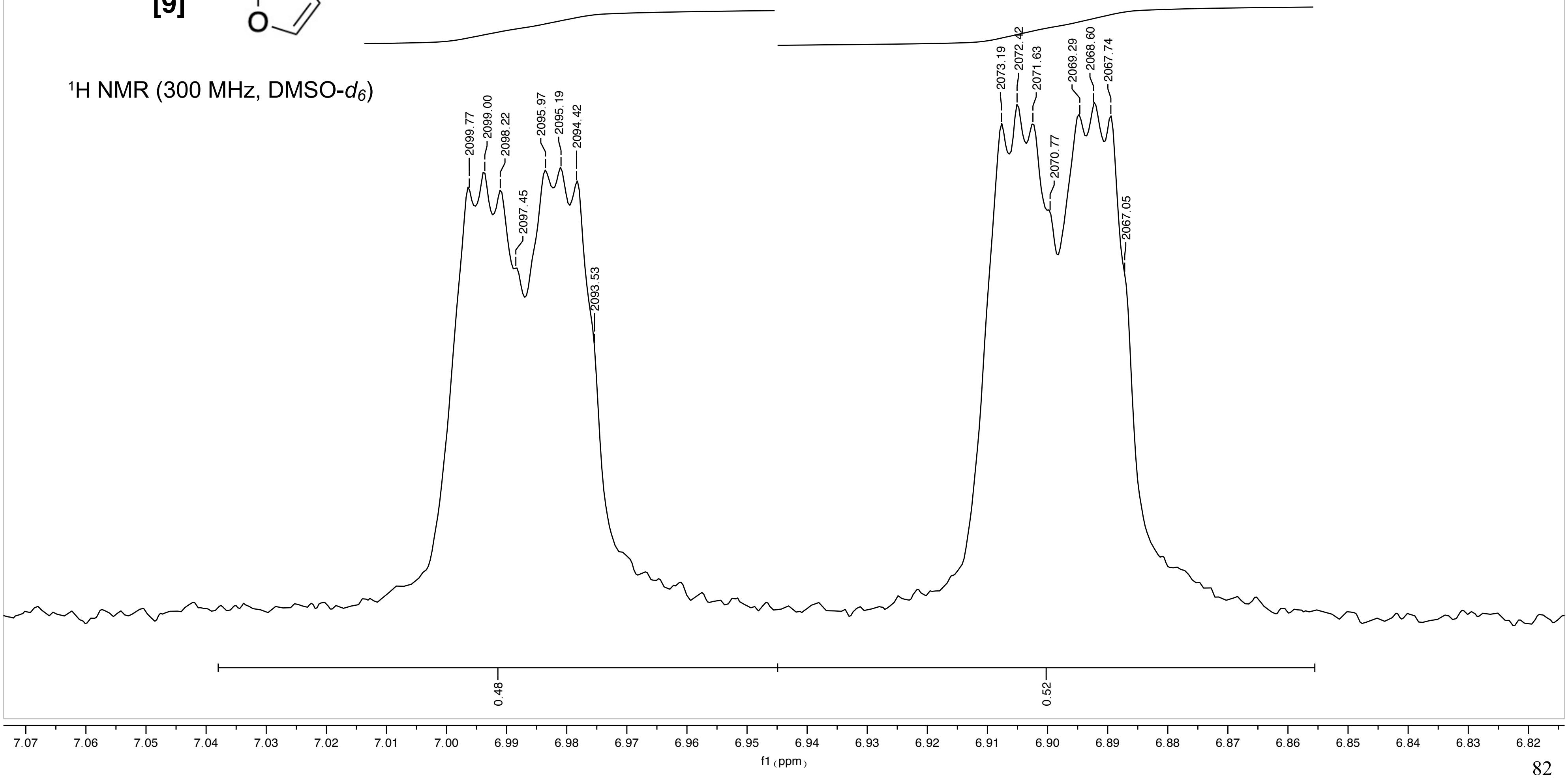

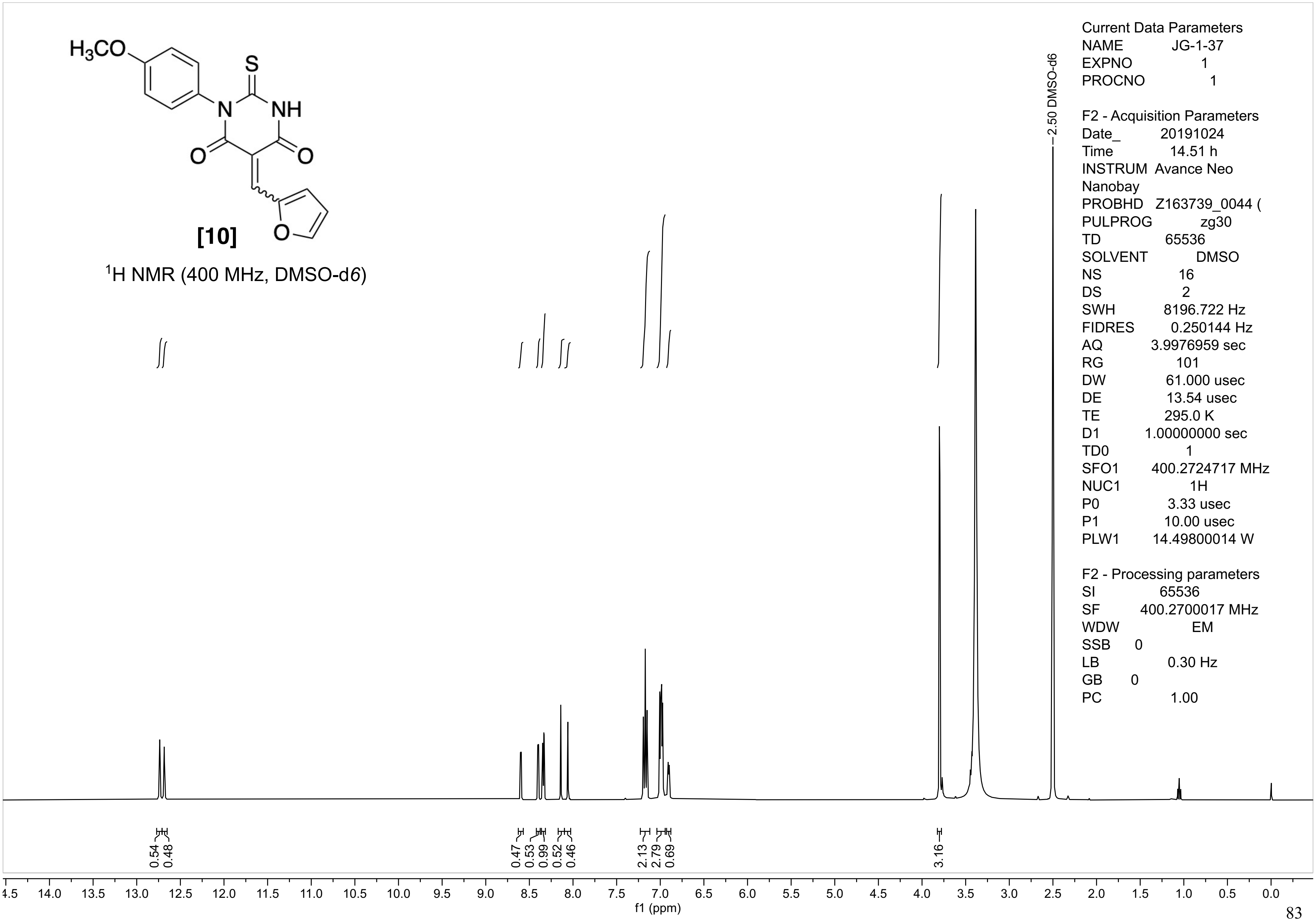
H3CO
S
N
NH
O
O
O
[10]
1H NMR (400 MHz, DMSO-d6)
2.50 DMSO-d6
Current Data Parameters
NAME JG-1-37
EXPNO 1
PROCNO 1
F2 - Acquisition Parameters
Date_ 20191024
Time 14.51 h
INSTRUM Avance Neo
Nanobay
PROBHD Z163739_0044 (
PULPROG zg30
TD 65536
SOLVENT DMSO
NS 16
DS 2
SWH 8196.722 Hz
FIDRES 0.250144 Hz
AQ 3.9976959 sec
RG 101
DW 61.000 usec
DE 13.54 usec
TE 295.0 K
D1 1.00000000 sec
TD0 1
SFO1 400.2724717 MHz
NUC1 1H
P0 3.33 usec
P1 10.00 usec
PLW1 14.49800014 W
F2 - Processing parameters
SI 65536
SF 400.2700017 MHz
WDW EM
SSB 0
LB 0.30 Hz
GB 0
PC 1.00
0.54
0.48
0.47
0.53
0.99
0.52
0.46
2.13
2.79
0.69
3.16
f1 (ppm)

JG-1-37

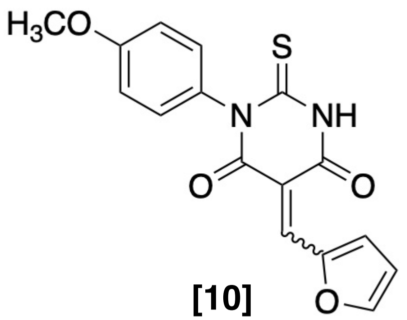

[10]

$^1$H NMR (400 MHz, DMSO-d$_6$)

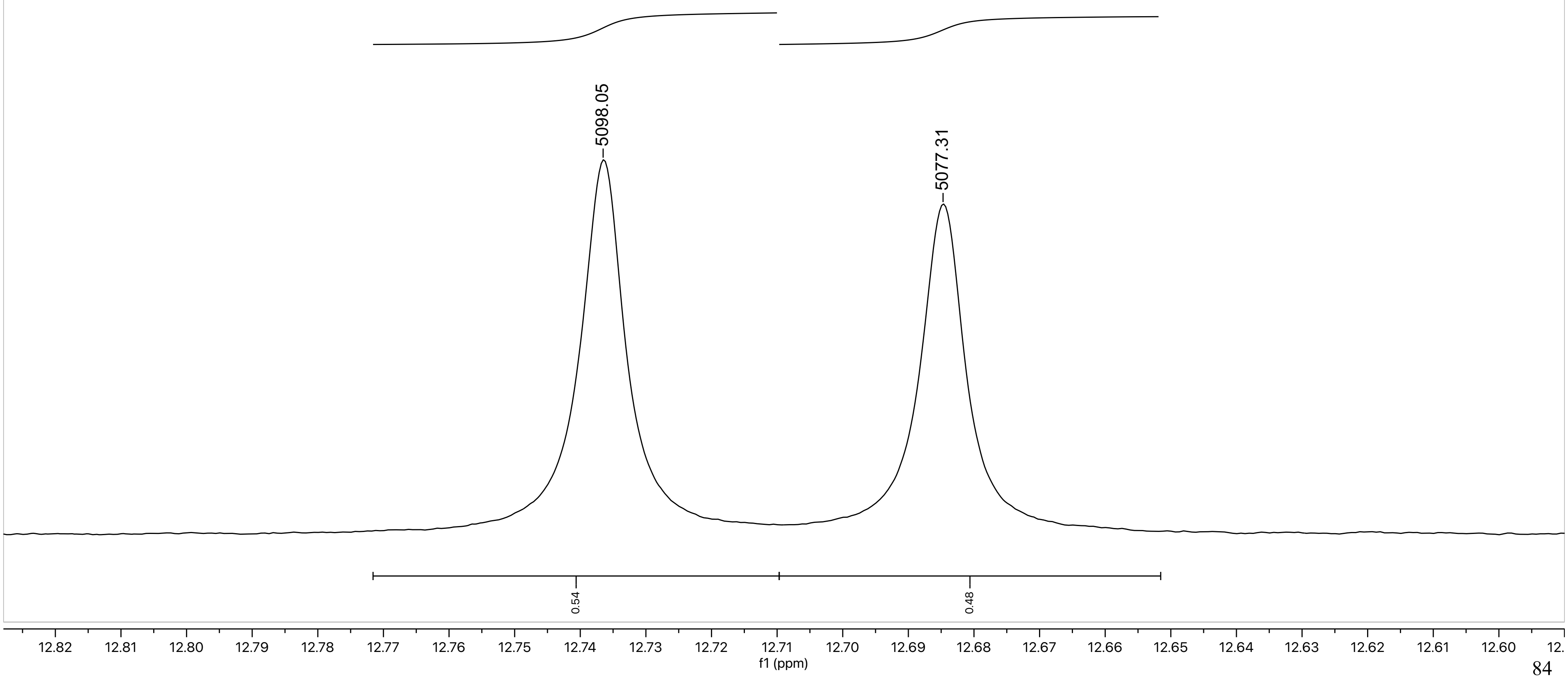

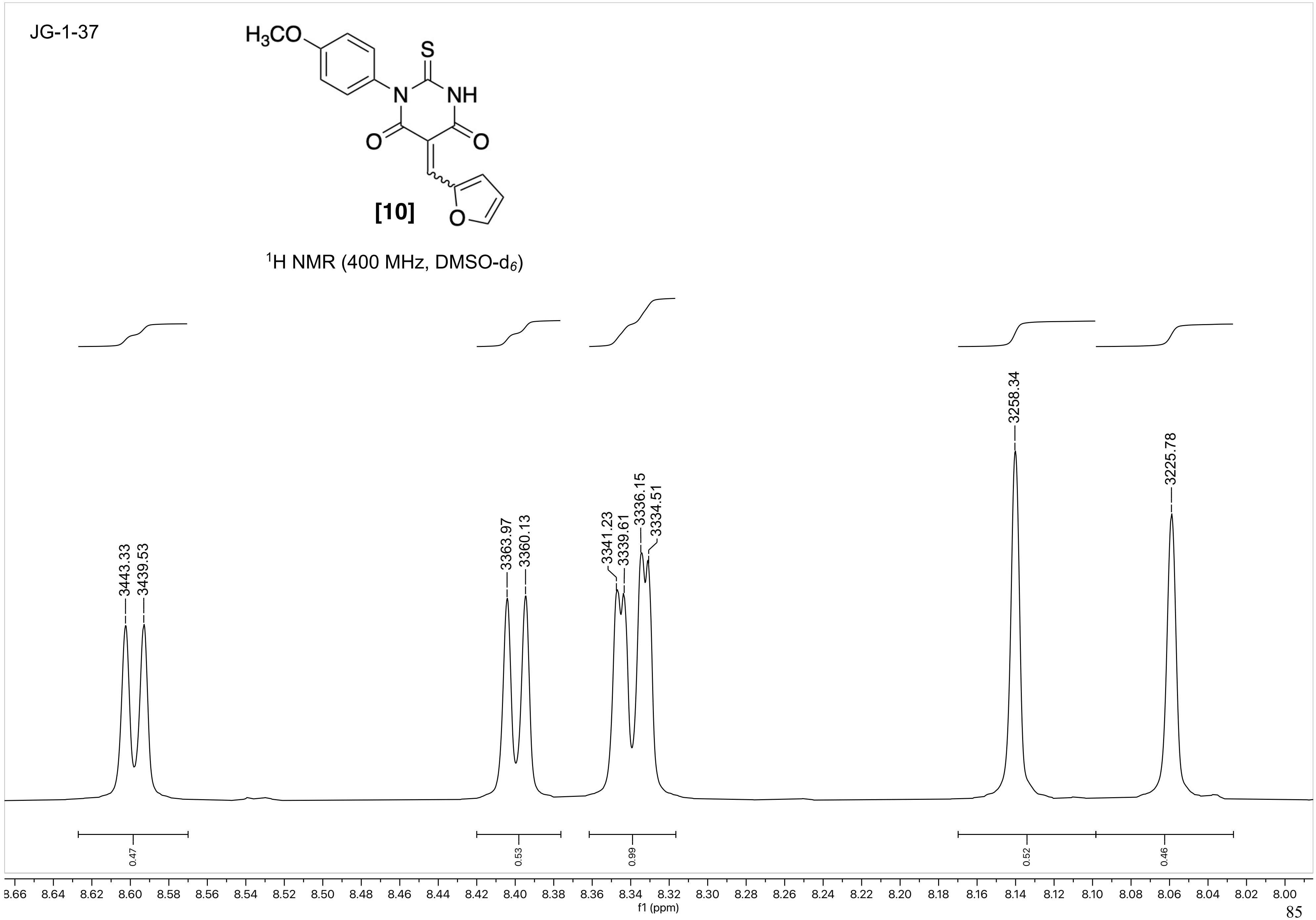
JG-1-37
H3CO
S
N
NH
O
O
O
[10]
1H NMR (400 MHz, DMSO-d6)
3443.33
3439.53
3363.97
3360.13
3341.23
3339.61
3336.15
3334.51
3258.34
3225.78
0.47
0.53
0.99
0.52
0.46
f1 (ppm)

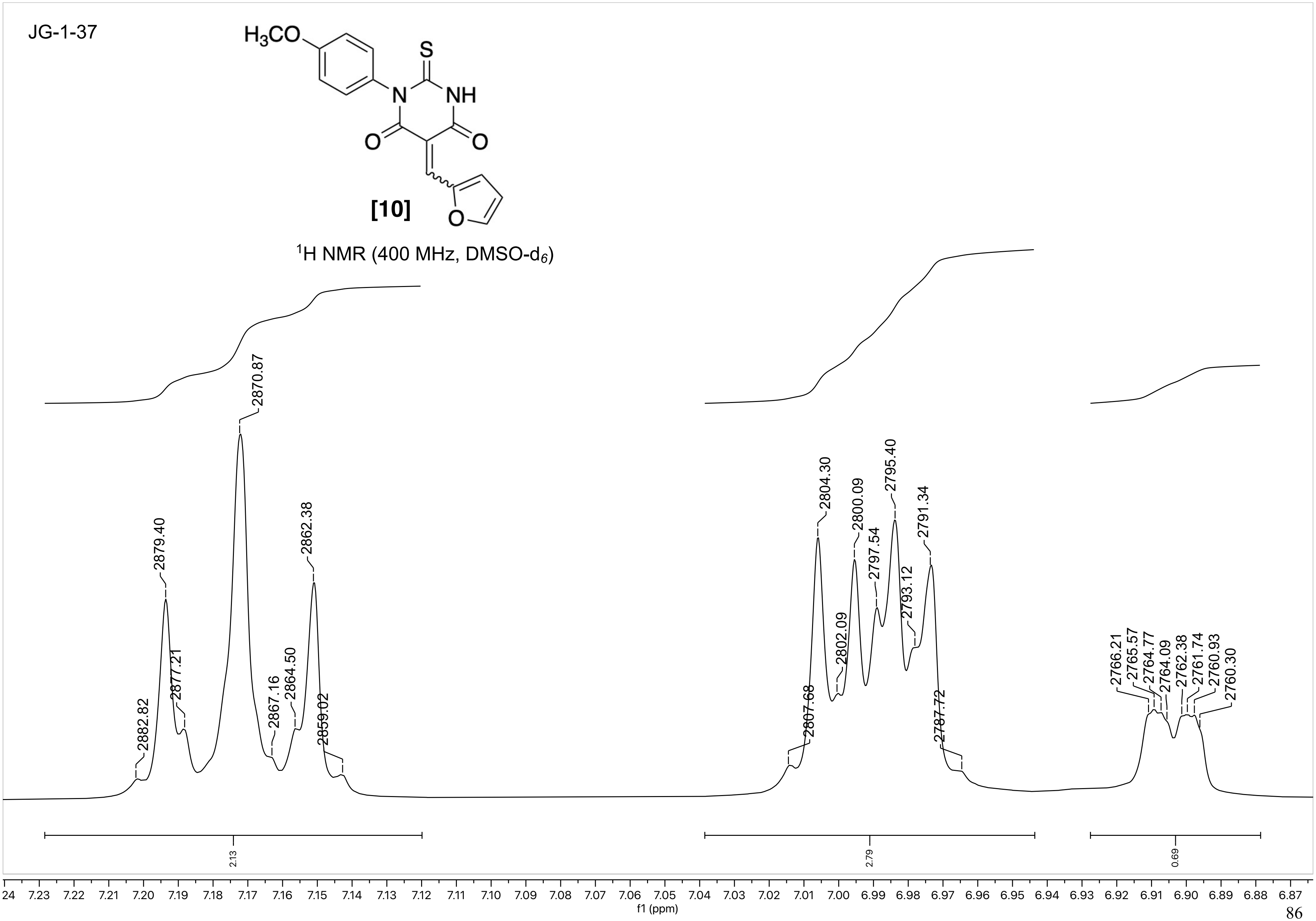
JG-1-37
H3CO
S
N
NH
O
O
O
[10]
1H NMR (400 MHz, DMSO-d6)
2882.82
2879.40
2877.21
2870.87
2867.16
2864.50
2862.38
2859.02
2807.68
2804.30
2802.09
2800.09
2797.54
2795.40
2793.12
2791.34
2787.72
2766.21
2765.57
2764.77
2764.09
2762.38
2761.74
2760.93
2760.30
2.13
2.79
0.69
f1 (ppm)

JG-1-37

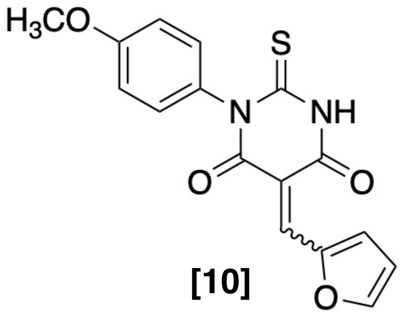

**[10]**

$^{1}$H NMR (400 MHz, DMSO-$d_6$)

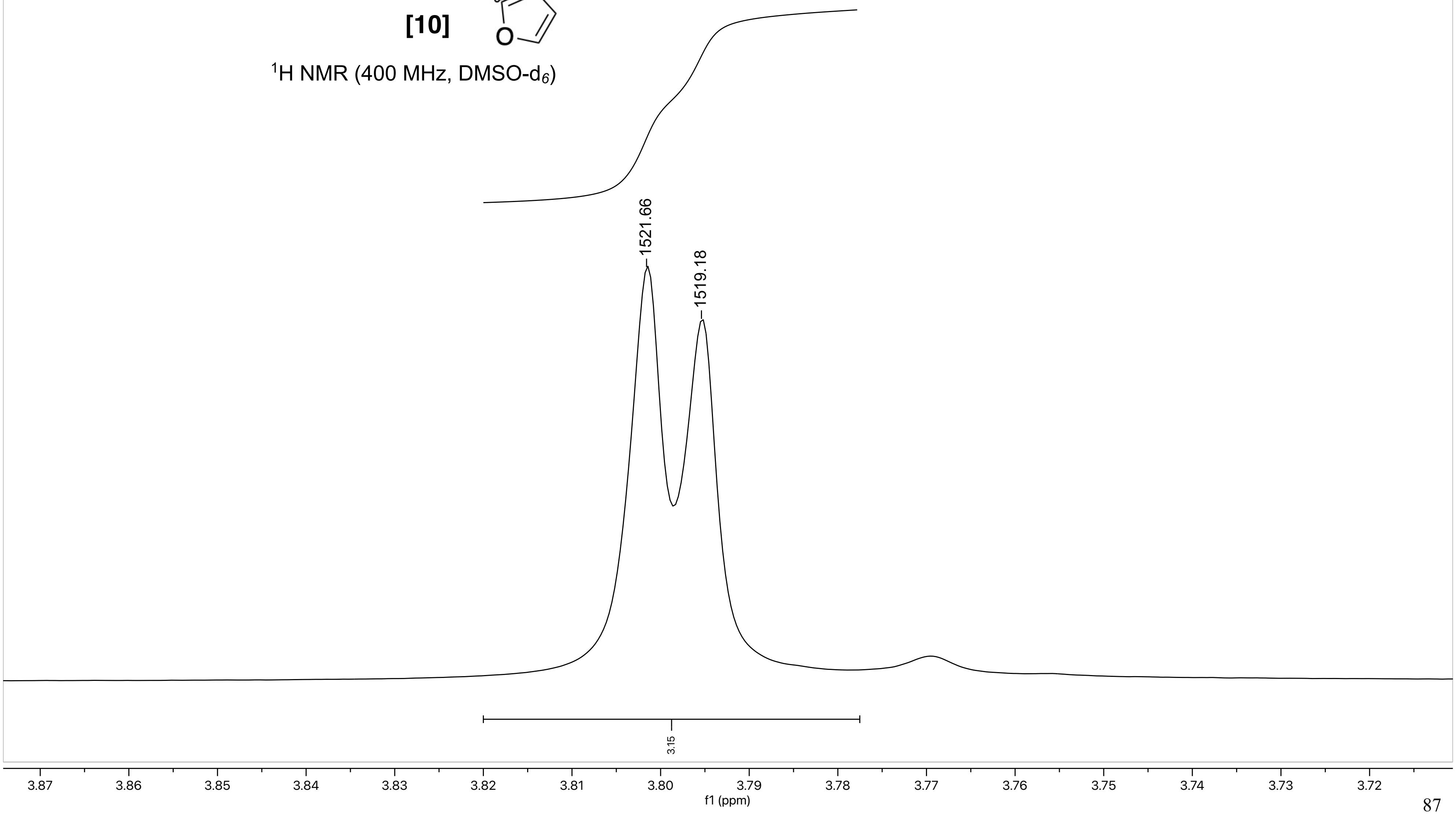

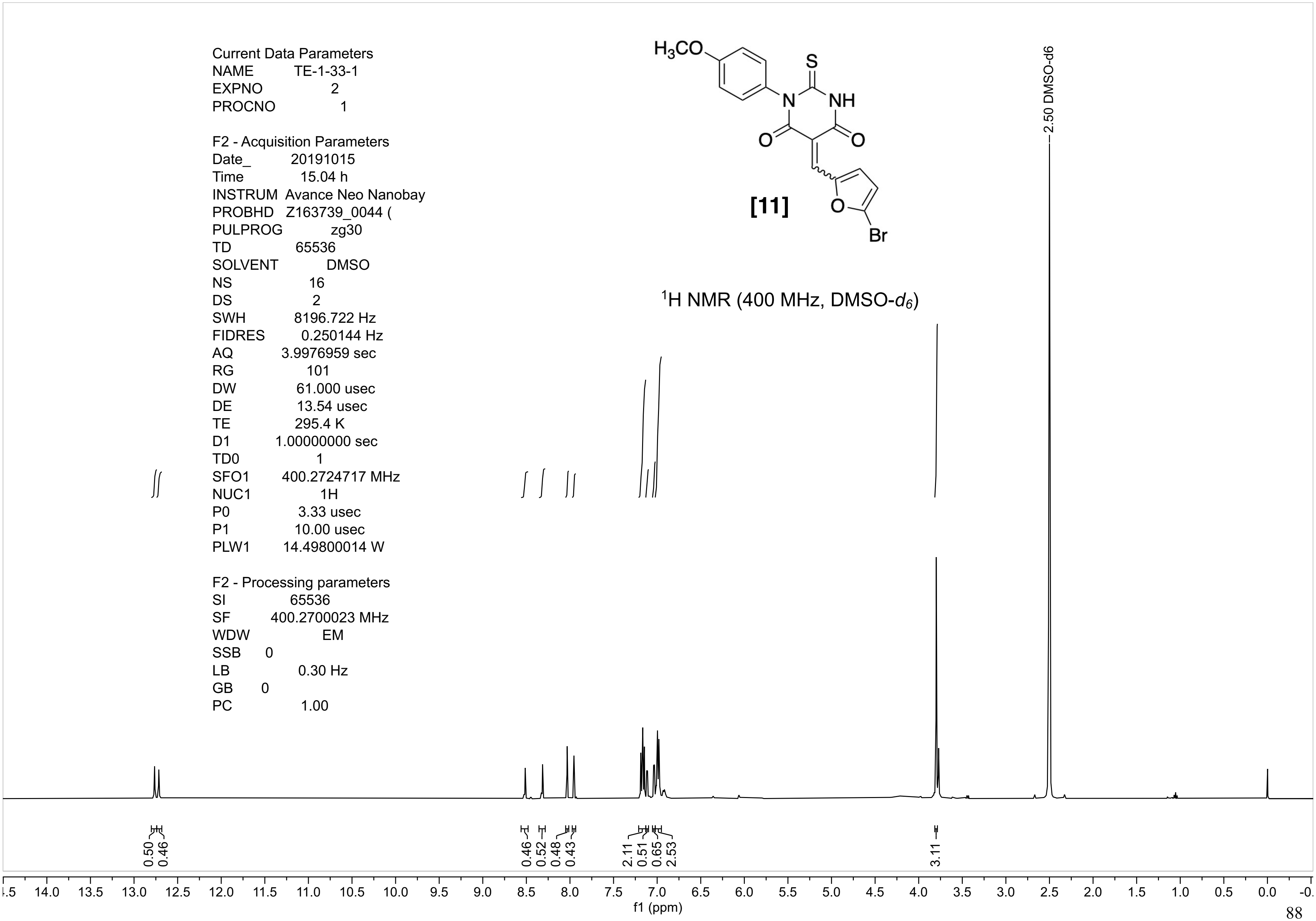
Current Data Parameters
NAME TE-1-33-1
EXPNO 2
PROCNO 1
F2 - Acquisition Parameters
Date_ 20191015
Time 15.04 h
INSTRUM Avance Neo Nanobay
PROBHD Z163739_0044 (
PULPROG zg30
TD 65536
SOLVENT DMSO
NS 16
DS 2
SWH 8196.722 Hz
FIDRES 0.250144 Hz
AQ 3.9976959 sec
RG 101
DW 61.000 usec
DE 13.54 usec
TE 295.4 K
D1 1.00000000 sec
TD0 1
SFO1 400.2724717 MHz
NUC1 1H
P0 3.33 usec
P1 10.00 usec
PLW1 14.49800014 W
F2 - Processing parameters
SI 65536
SF 400.2700023 MHz
WDW EM
SSB 0
LB 0.30 Hz
GB 0
PC 1.00
H3CO
S
N
NH
O
O
O
Br
[11]
1H NMR (400 MHz, DMSO-d6)
2.50 DMSO-d6
0.50
0.46
0.46
0.52
0.48
0.43
2.11
0.51
0.65
2.53
3.11
14.0 13.5 13.0 12.5 12.0 11.5 11.0 10.5 10.0 9.5 9.0 8.5 8.0 7.5 7.0 6.5 6.0 5.5 5.0 4.5 4.0 3.5 3.0 2.5 2.0 1.5 1.0 0.5 0.0
f1 (ppm)

TE-1-33-1

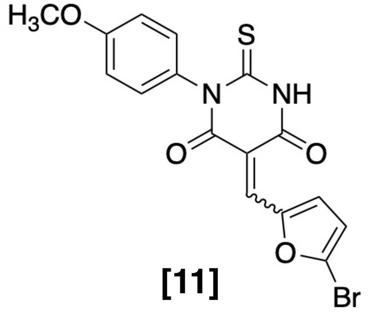

**[11]**

$^{1}H$ NMR (400 MHz, DMSO-$d_6$)

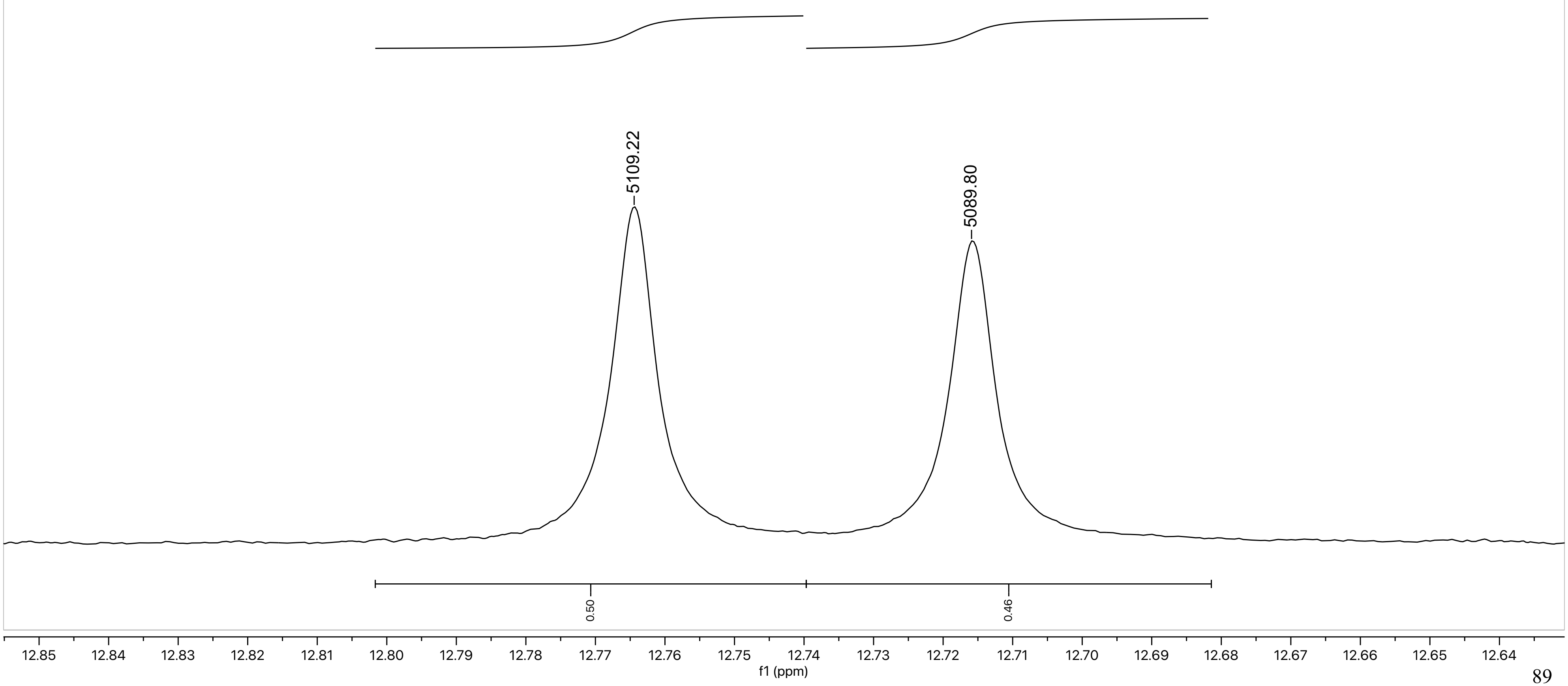

TE-1-33-1

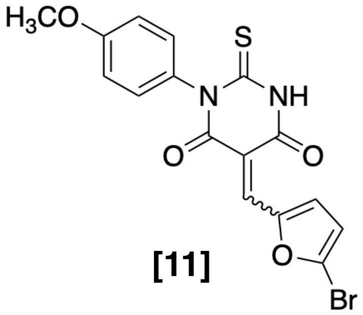

**[11]**

$^{1}H$ NMR (400 MHz, DMSO-$d_6$)

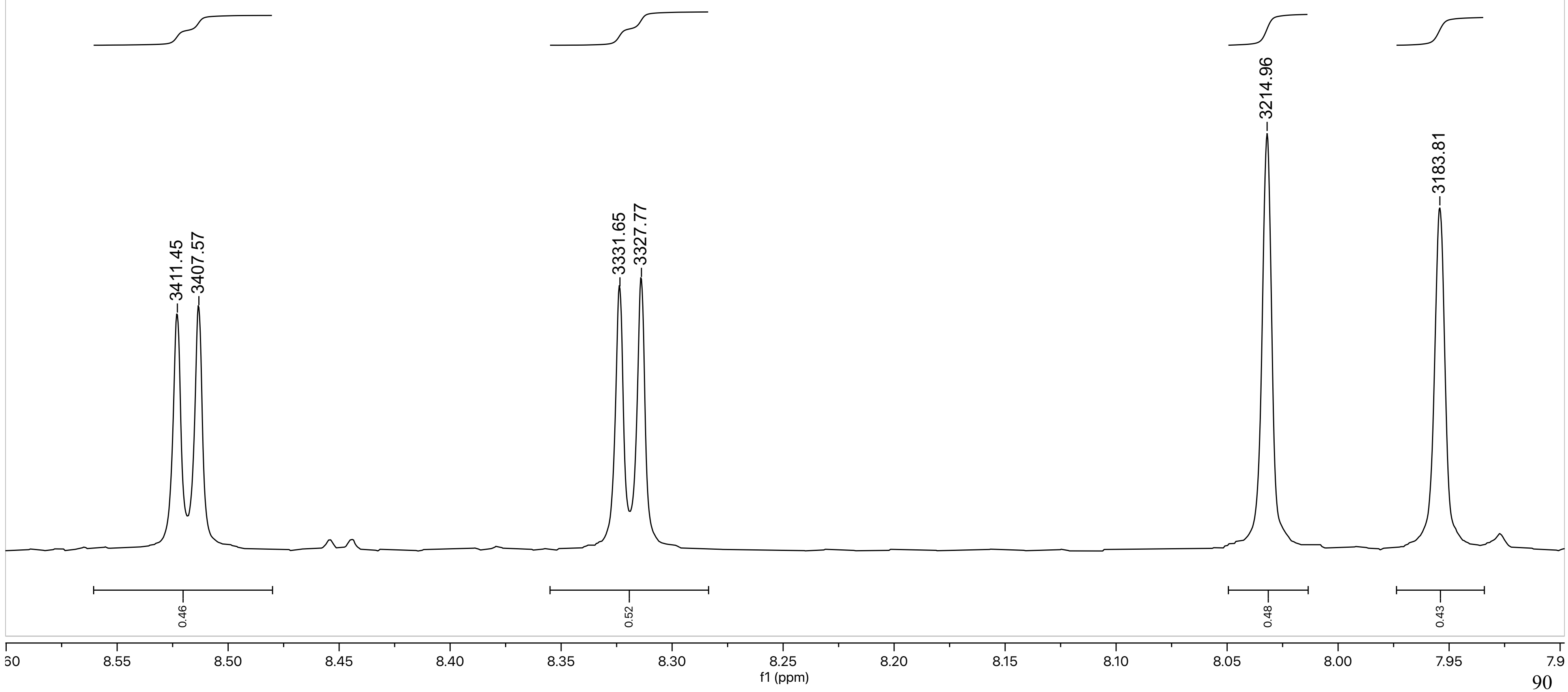

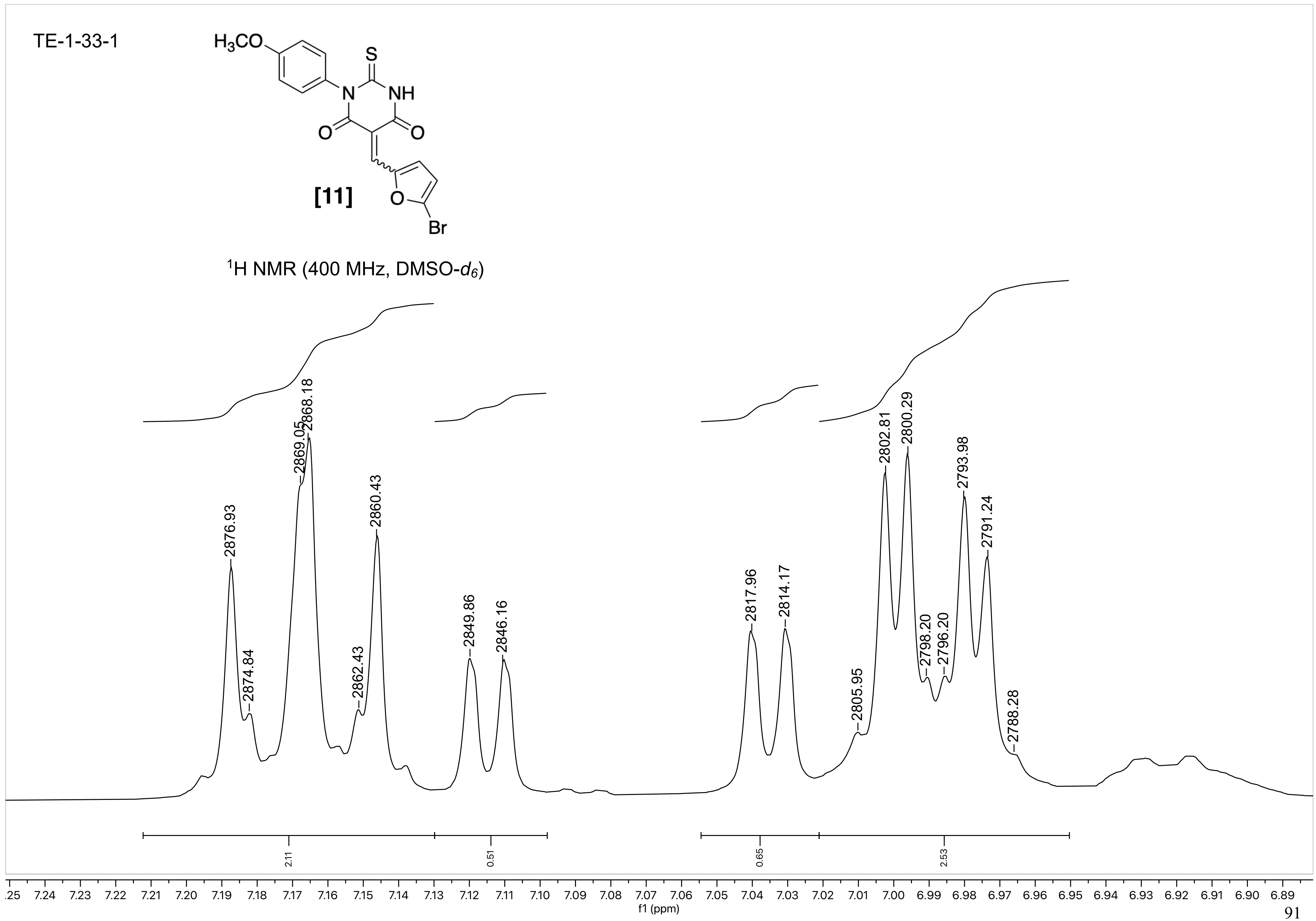

TE-1-33-1
H3CO
S
N
NH
O
O
O
Br
[11]
1H NMR (400 MHz, DMSO-d6)
2876.93
2874.84
2869.05
2868.18
2862.43
2860.43
2849.86
2846.16
2817.96
2814.17
2805.95
2802.81
2800.29
2798.20
2796.20
2793.98
2791.24
2788.28
2.11
0.51
0.65
2.53
.25 7.24 7.23 7.22 7.21 7.20 7.19 7.18 7.17 7.16 7.15 7.14 7.13 7.12 7.11 7.10 7.09 7.08 7.07 7.06 7.05 7.04 7.03 7.02 7.01 7.00 6.99 6.98 6.97 6.96 6.95 6.94 6.93 6.92 6.91 6.90 6.89
f1 (ppm)

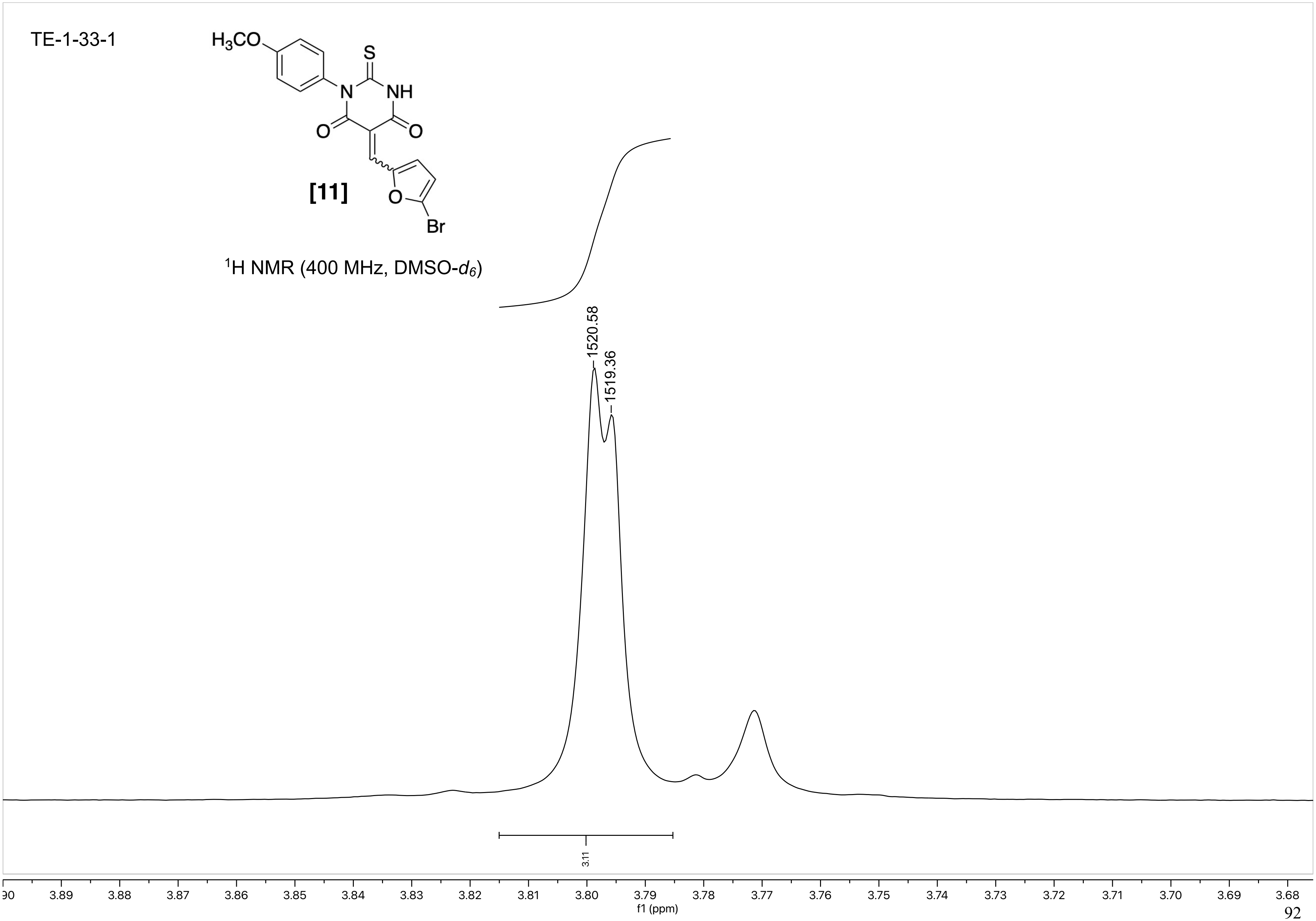
TE-1-33-1
H3CO
S
N
NH
O
O
[11]
O
Br
1H NMR (400 MHz, DMSO-d6)
1520.58
1519.36
3.11
3.90 3.89 3.88 3.87 3.86 3.85 3.84 3.83 3.82 3.81 3.80 3.79 3.78 3.77 3.76 3.75 3.74 3.73 3.72 3.71 3.70 3.69 3.68
f1 (ppm)

NAME SW1-30
EXPNO 1
PROCNO 1

F2 - Acquisition Parameters
Date_ 20191015
Time 16.27 h
INSTRUM Avance Neo Nanobay
PROBHD Z163739_0044 (
PULPROG zg30
TD 65536
SOLVENT DMSO
NS 16
DS 2
SWH 8196.722 Hz
FIDRES 0.250144 Hz
AQ 3.9976959 sec
RG 101
DW 61.000 usec
DE 13.54 usec
TE 295.2 K
D1 1.00000000 sec
TD0 1
SFO1 400.2724717 MHz
NUC1 1H
P0 3.33 usec
P1 10.00 usec
PLW1 14.49800014 W

F2 - Processing parameters
SI 65536
SF 400.2700017 MHz
WDW EM
SSB 0
LB 0.30 Hz
GB 0
PC 1.00

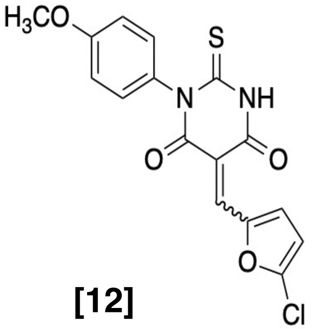

**[12]**

$^{1}$H NMR (400 MHz, DMSO-$d_6$)

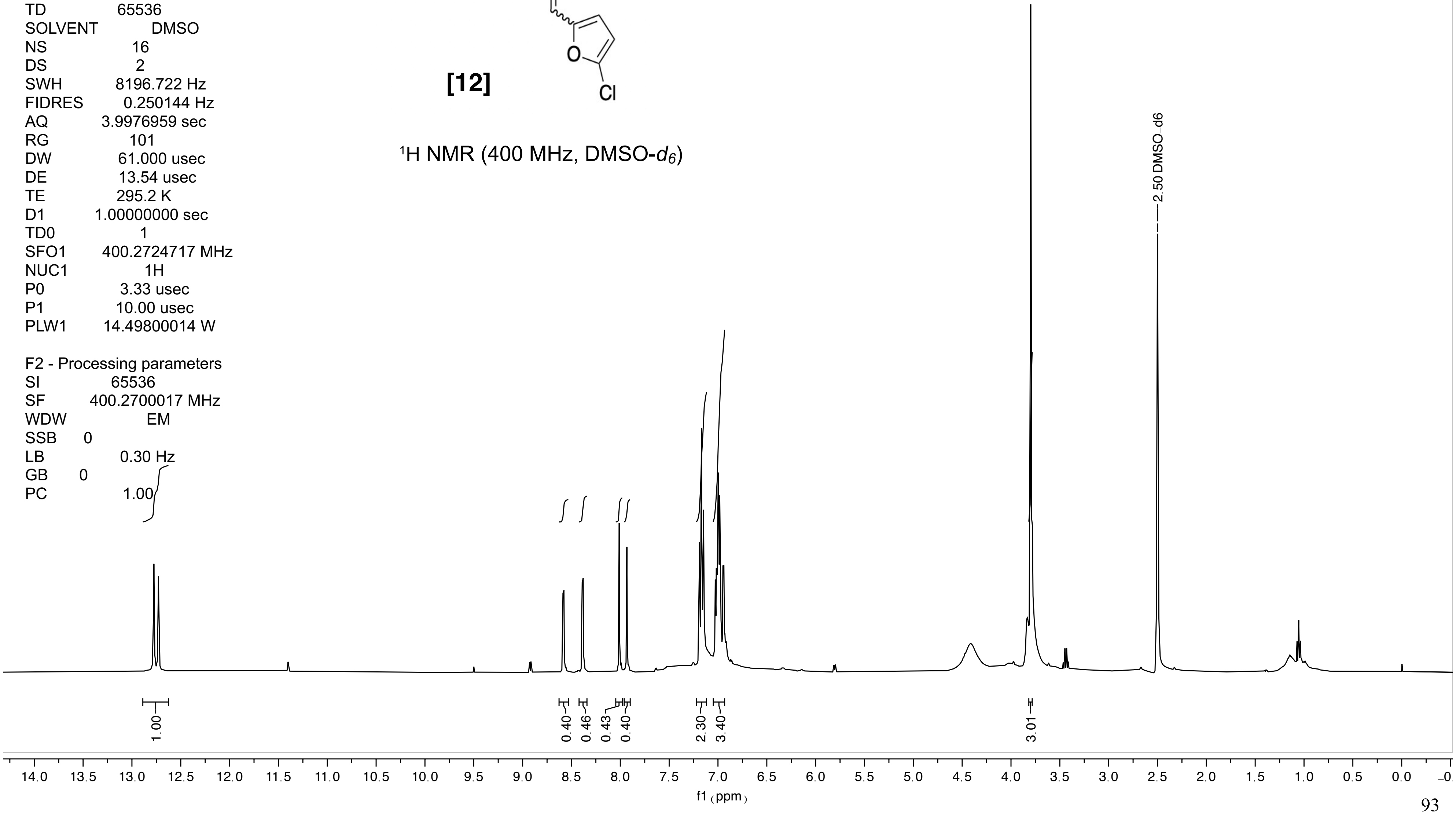

SW-I-30

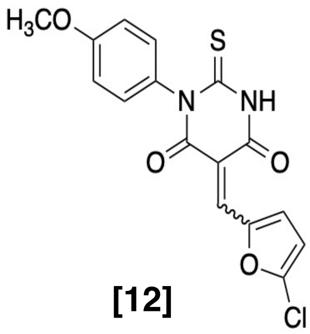

**[12]**

$^1$H NMR (400 MHz, DMSO-$d_6$)

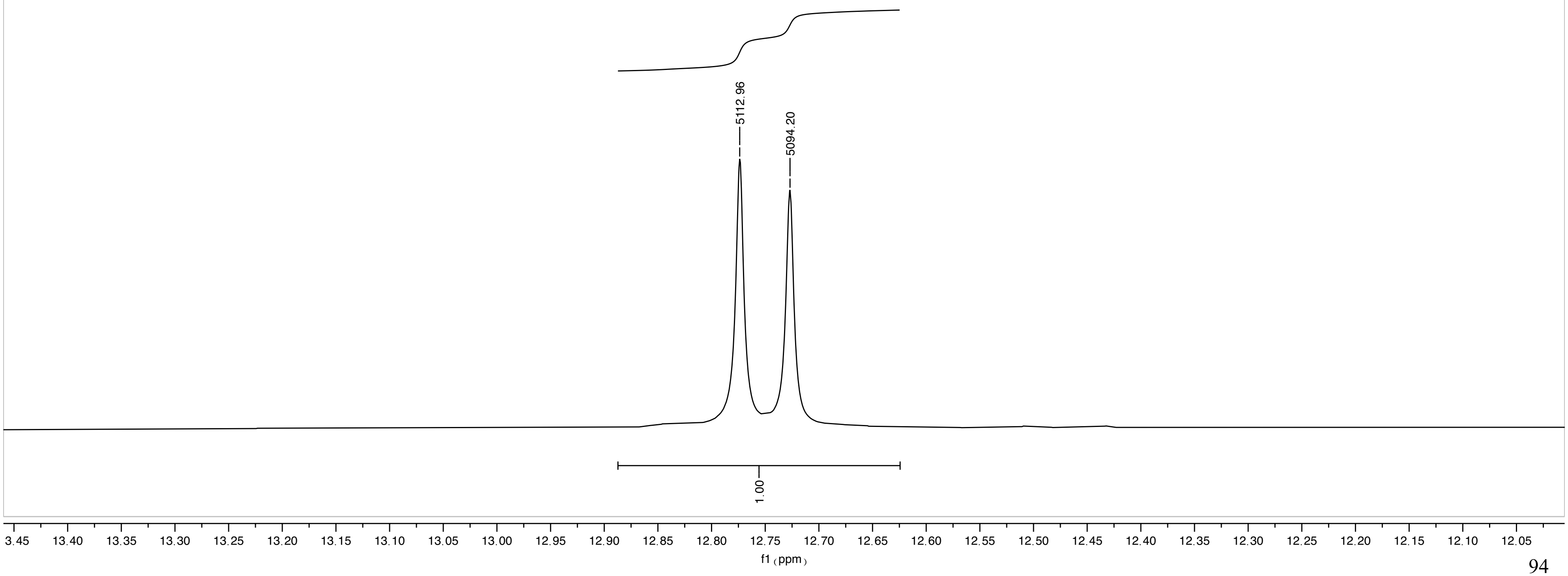

SW-I-30

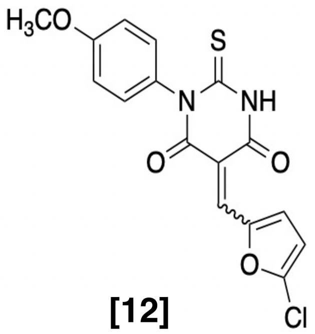

**[12]**

[1]H NMR (400 MHz, DMSO-$d_6$)

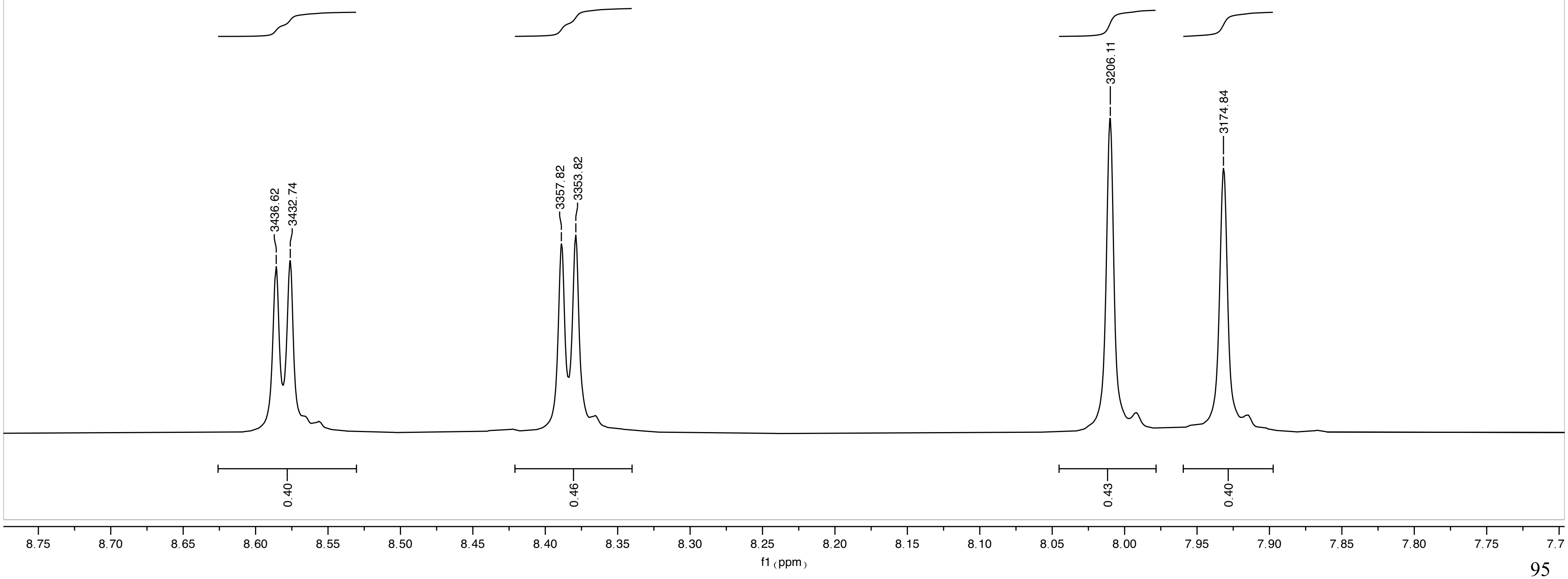

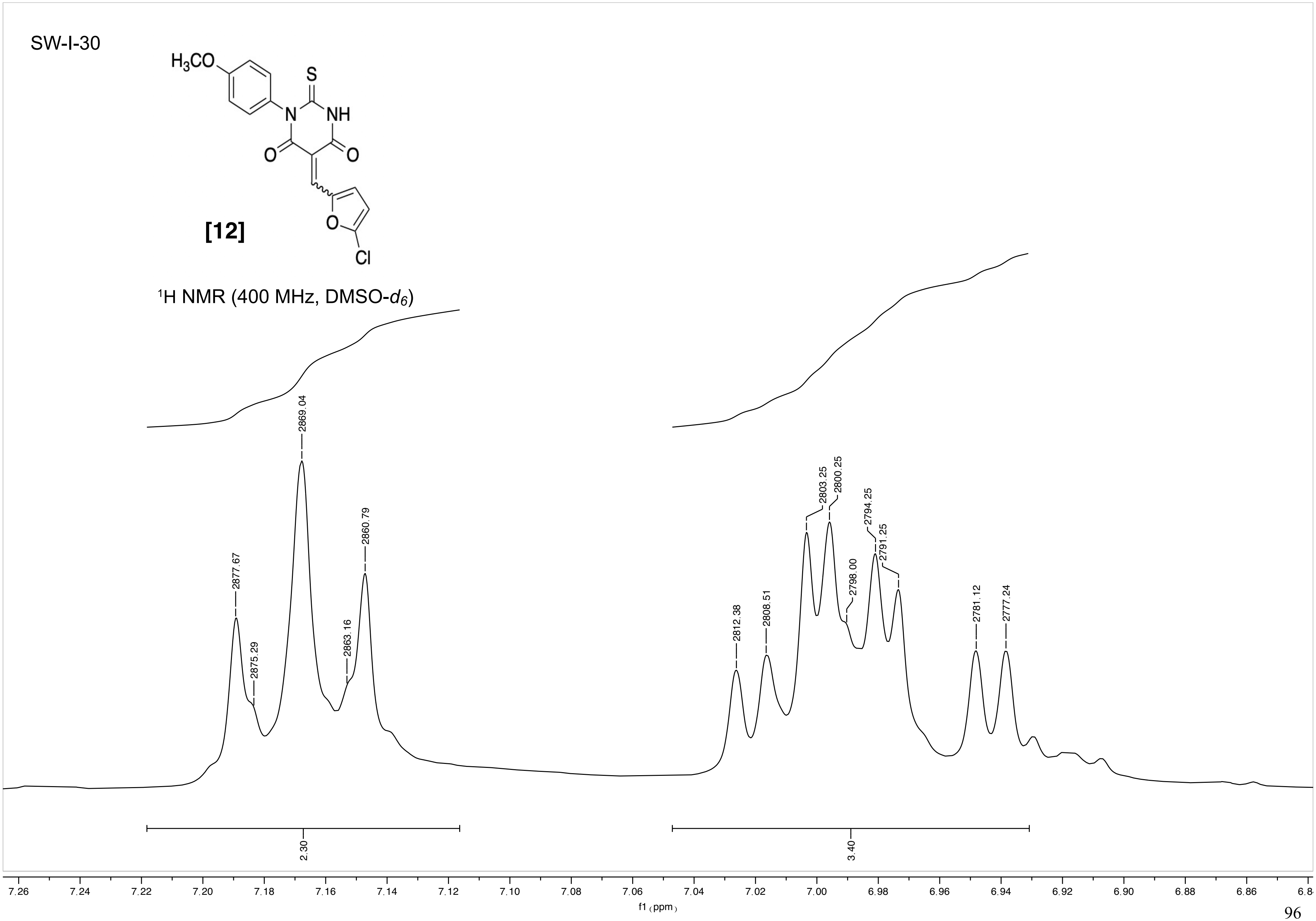
SW-I-30
$H_3CO$
S
N
NH
O
O
O
Cl
[12]
$^1$H NMR (400 MHz, DMSO-$d_6$)
2877.67
2875.29
2869.04
2863.16
2860.79
2812.38
2808.51
2803.25
2800.25
2798.00
2794.25
2791.25
2781.12
2777.24
2.30
3.40
7.26 7.24 7.22 7.20 7.18 7.16 7.14 7.12 7.10 7.08 7.06 7.04 7.02 7.00 6.98 6.96 6.94 6.92 6.90 6.88 6.86
f1 (ppm)

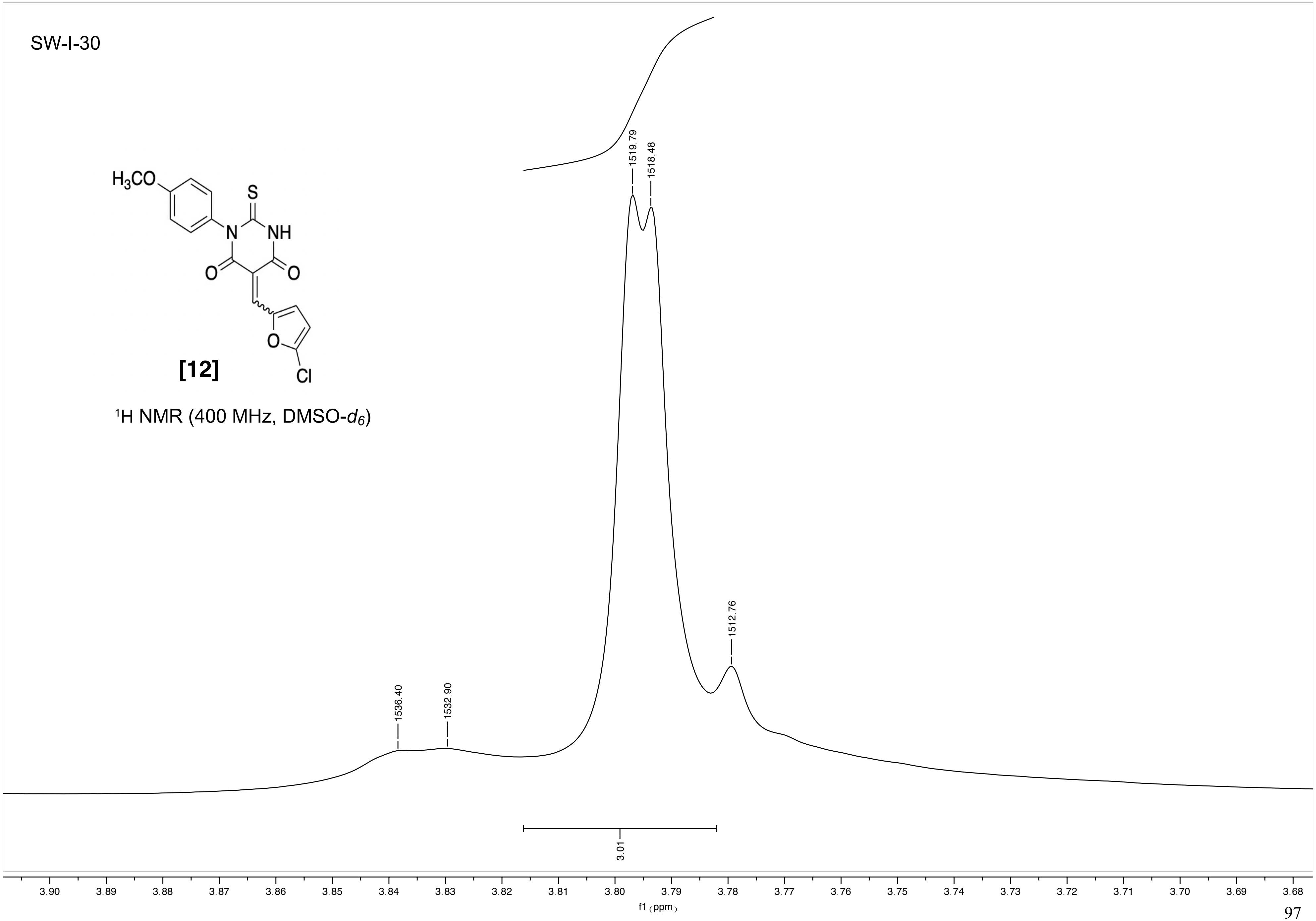
SW-I-30
H3CO
S
N
NH
O
O
O
Cl
[12]
1H NMR (400 MHz, DMSO-d6)
1519.79
1518.48
1512.76
1536.40
1532.90
3.01
3.90
3.89
3.88
3.87
3.86
3.85
3.84
3.83
3.82
3.81
3.80
3.79
3.78
3.77
3.76
3.75
3.74
3.73
3.72
3.71
3.70
3.69
3.68
f1 (ppm)

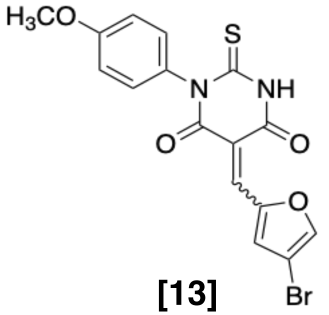

**[13]**

$^1H$ NMR (400 MHz, $CDCl_3$)

Current Data Parameters
NAME I-JG-23
EXPNO 5
PROCNO 1

F2 - Acquisition Parameters
Date_ 20211027
Time 16.08 h
INSTRUM Avance Neo Nanobay
PROBHD Z163739_0044 (
PULPROG zg30
TD 65536
SOLVENT CDCl3
NS 16
DS 2
SWH 8196.722 Hz
FIDRES 0.250144 Hz
AQ 3.9976959 sec
RG 101
DW 61.000 usec
DE 13.24 usec
TE 300.0 K
D1 1.00000000 sec
TD0 1
SFO1 400.2724717 MHz
NUC1 1H
P0 3.92 usec
P1 11.75 usec
PLW1 14.49800014 W

F2 - Processing parameters
SI 65536
SF 400.2700061 MHz
WDW EM
SSB 0
LB 0.30 Hz
GB 0
PC 1.00

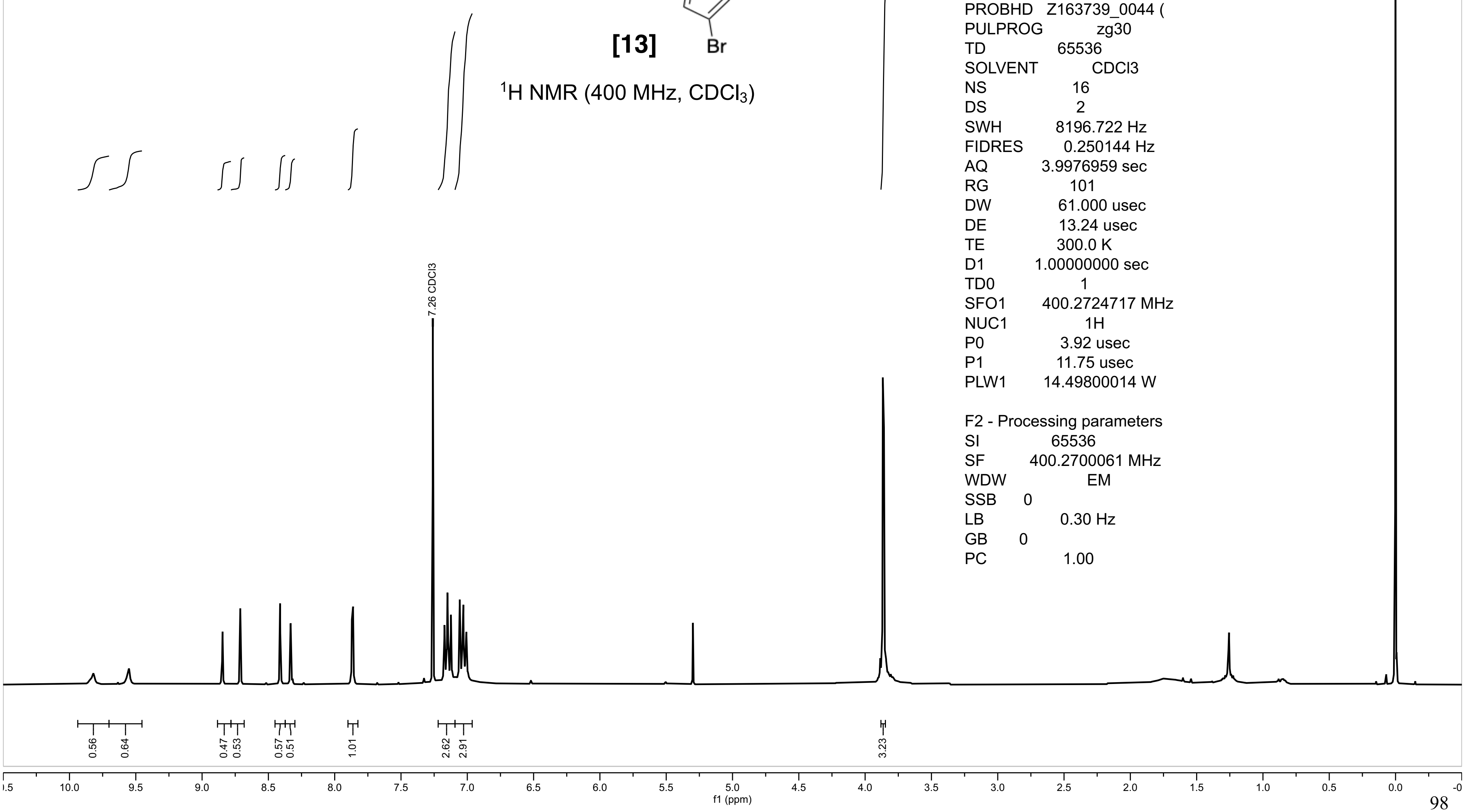

I-JG-23

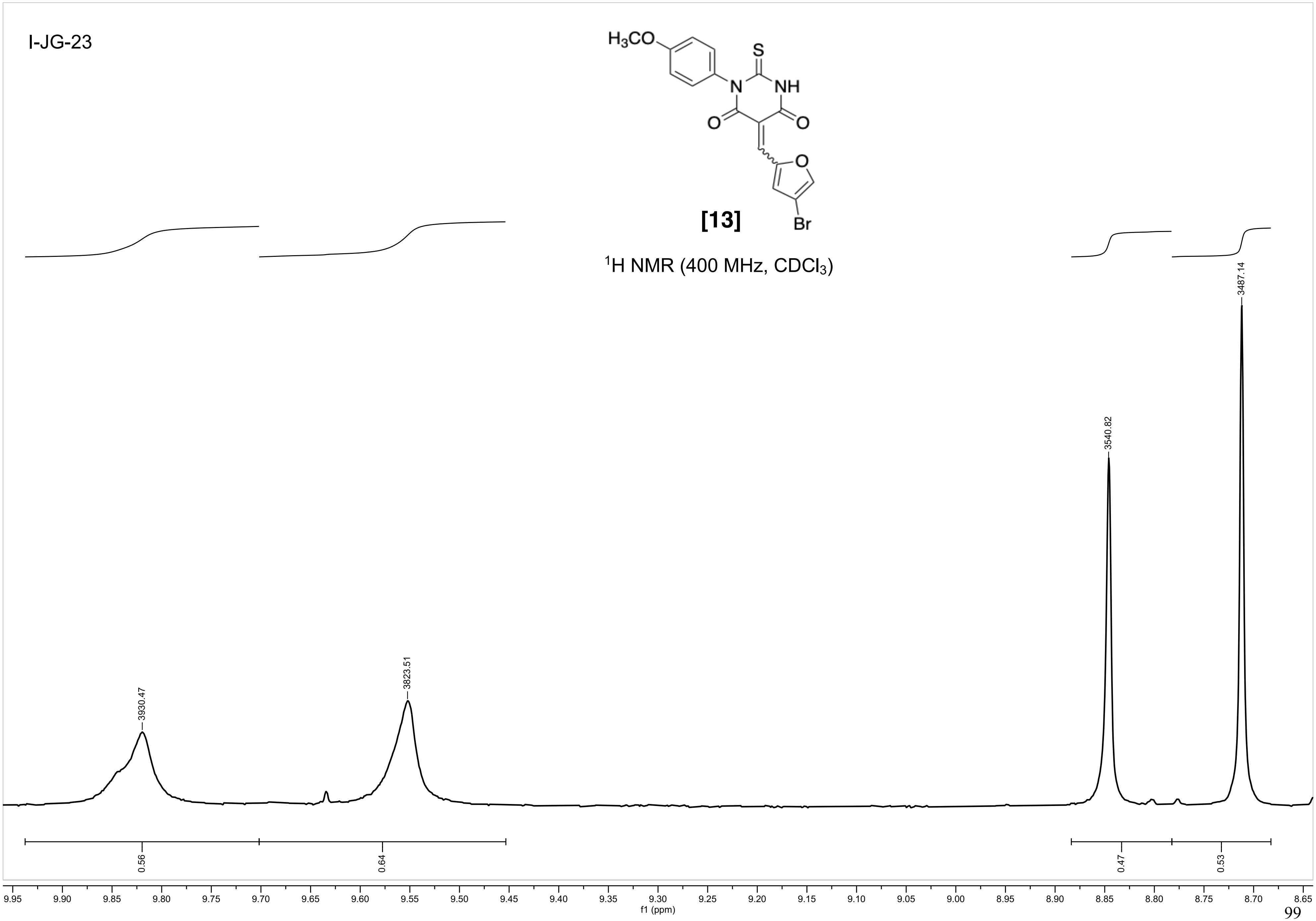
H3CO
S
N
NH
O
O
O
Br
[13]
1H NMR (400 MHz, CDCl3)
3930.47
3823.51
3540.82
3487.14
0.56
0.64
0.47
0.53
9.95
9.90
9.85
9.80
9.75
9.70
9.65
9.60
9.55
9.50
9.45
9.40
9.35
9.30
9.25
9.20
9.15
9.10
9.05
9.00
8.95
8.90
8.85
8.80
8.75
8.70
8.65
f1 (ppm)

I-JG-23

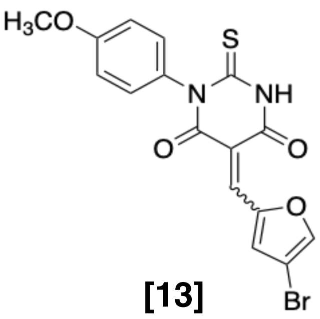

[13]

$^{1}$H NMR (400 MHz, $CDCl_3$)

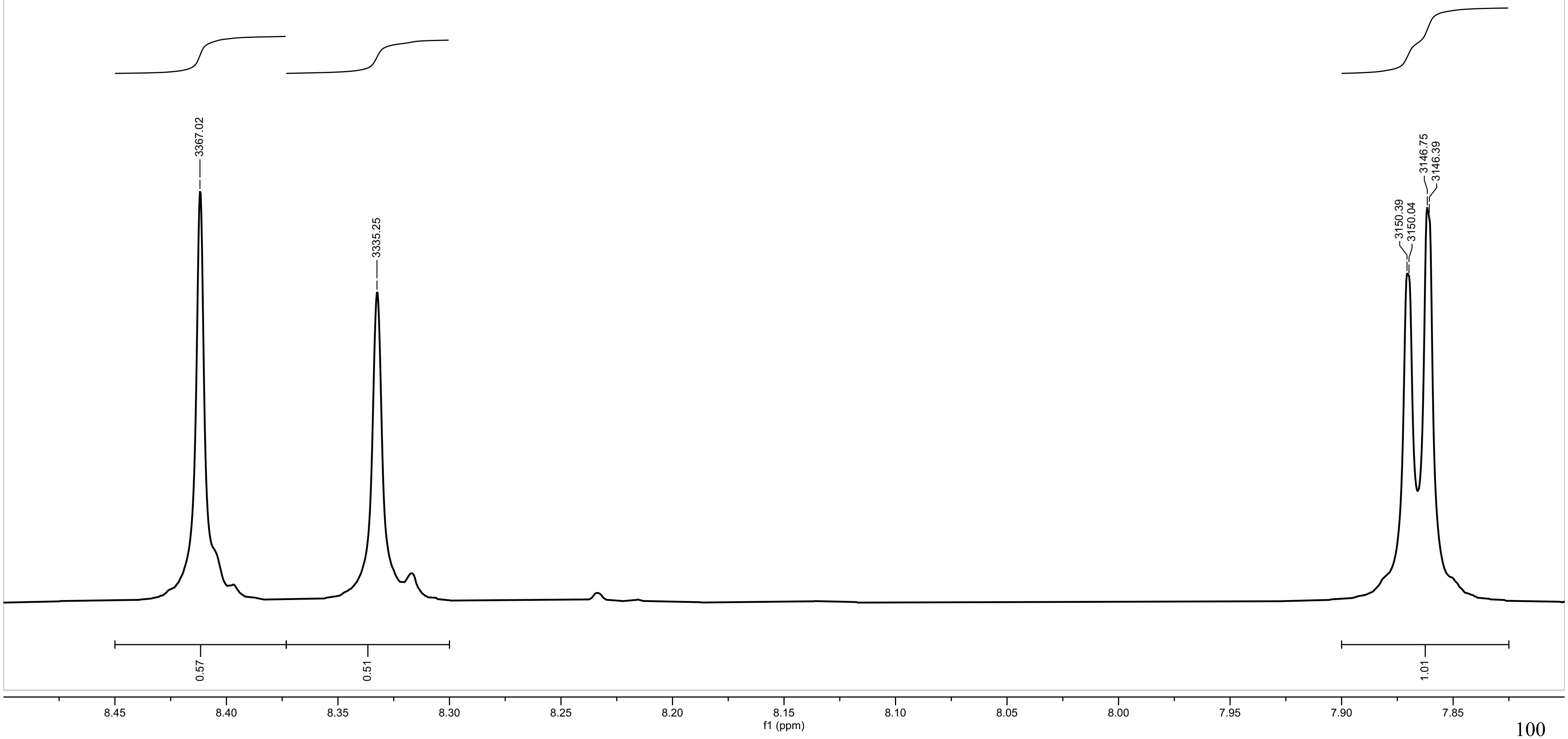

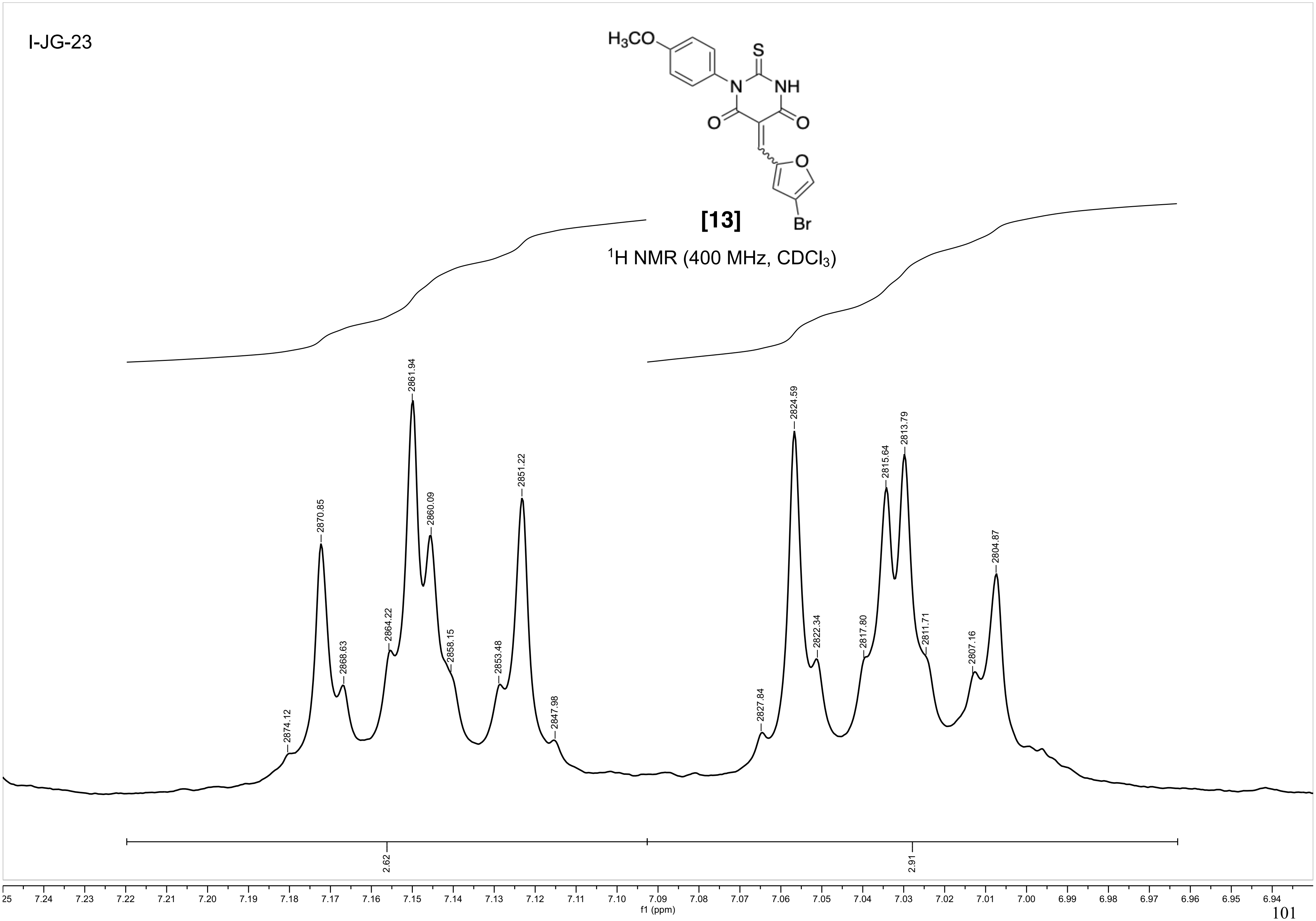
I-JG-23
H3CO
S
N
NH
O
O
O
Br
[13]
1H NMR (400 MHz, CDCl3)
2874.12
2870.85
2868.63
2864.22
2861.94
2860.09
2858.15
2853.48
2851.22
2847.98
2827.84
2824.59
2822.34
2817.80
2815.64
2813.79
2811.71
2807.16
2804.87
2.62
2.91
f1 (ppm)

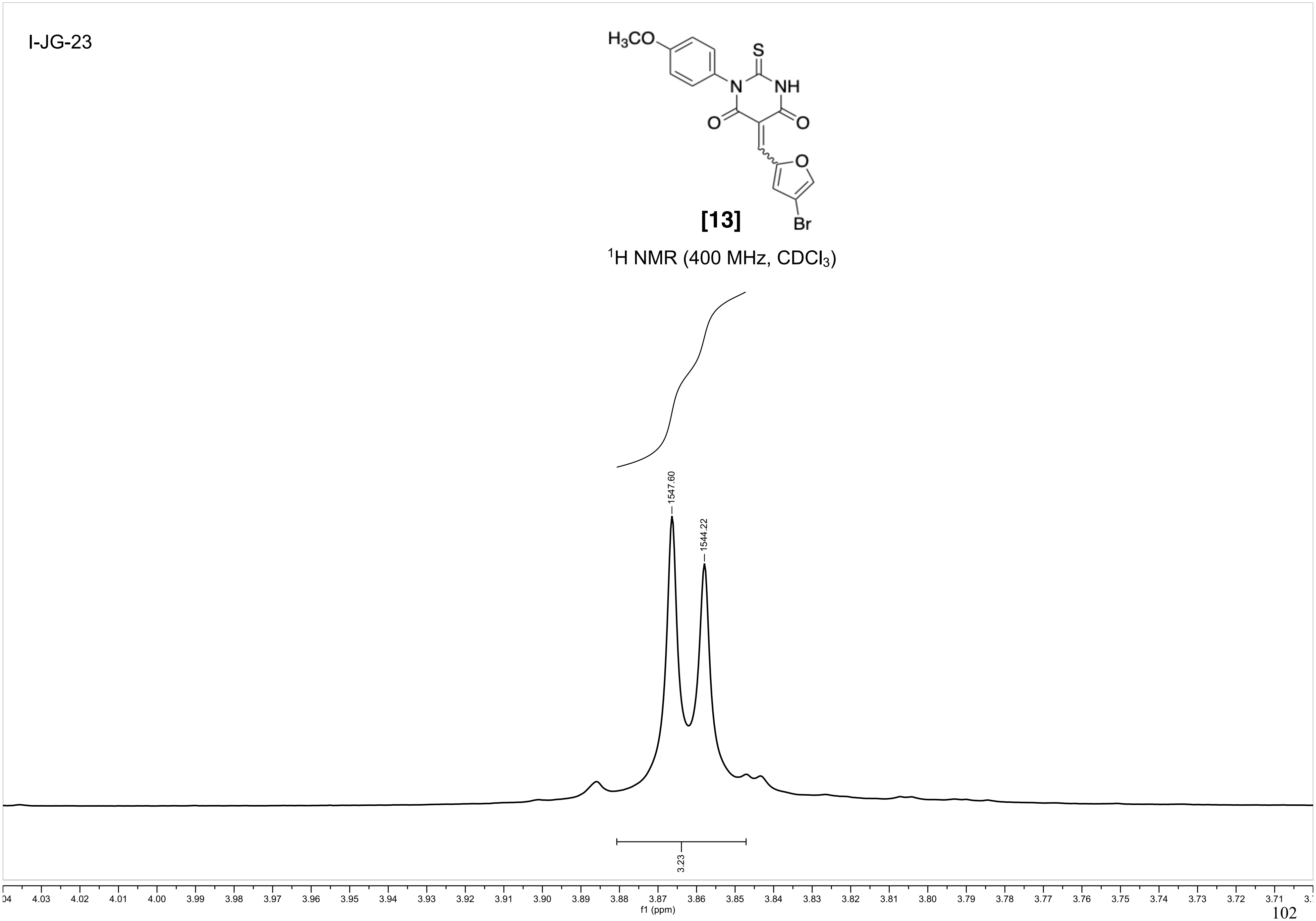
I-JG-23
H3CO
S
N
NH
O
O
O
Br
[13]
1H NMR (400 MHz, CDCl3)
1547.60
1544.22
3.23
f1 (ppm)

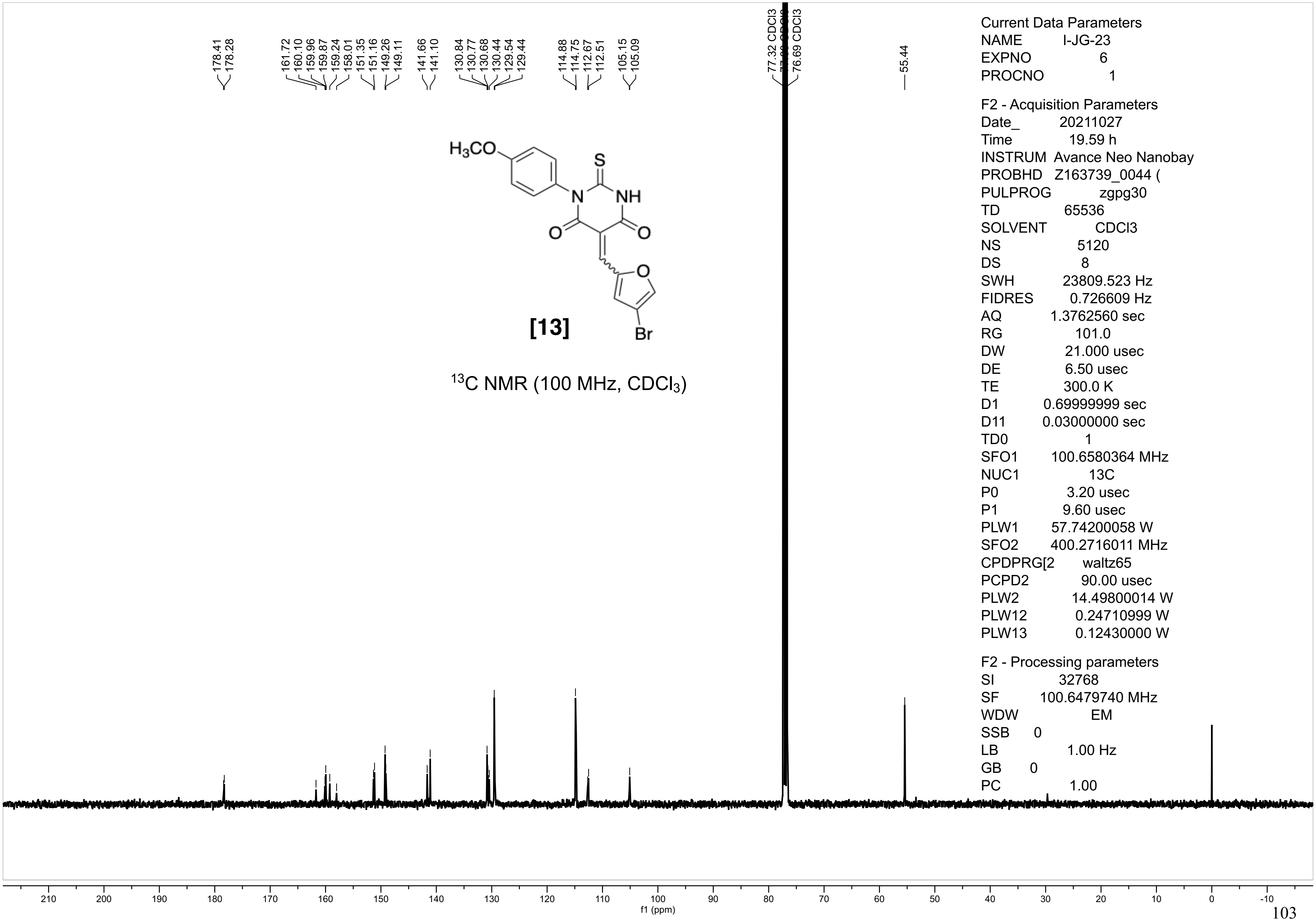
178.41
178.28
161.72
160.10
159.96
159.87
159.24
158.01
151.35
151.16
149.26
149.11
141.66
141.10
130.84
130.77
130.68
130.44
129.54
129.44
114.88
114.75
112.67
112.51
105.15
105.09
77.32 CDCl3
76.69 CDCl3
55.44
H3CO
S
N
NH
O
O
O
Br
[13]
13C NMR (100 MHz, CDCl3)
Current Data Parameters
NAME I-JG-23
EXPNO 6
PROCNO 1
F2 - Acquisition Parameters
Date_ 20211027
Time 19.59 h
INSTRUM Avance Neo Nanobay
PROBHD Z163739_0044 (
PULPROG zgpg30
TD 65536
SOLVENT CDCl3
NS 5120
DS 8
SWH 23809.523 Hz
FIDRES 0.726609 Hz
AQ 1.3762560 sec
RG 101.0
DW 21.000 usec
DE 6.50 usec
TE 300.0 K
D1 0.69999999 sec
D11 0.03000000 sec
TD0 1
SFO1 100.6580364 MHz
NUC1 13C
P0 3.20 usec
P1 9.60 usec
PLW1 57.74200058 W
SFO2 400.2716011 MHz
CPDPRG[2 waltz65
PCPD2 90.00 usec
PLW2 14.49800014 W
PLW12 0.24710999 W
PLW13 0.12430000 W
F2 - Processing parameters
SI 32768
SF 100.6479740 MHz
WDW EM
SSB 0
LB 1.00 Hz
GB 0
PC 1.00
210 200 190 180 170 160 150 140 130 120 110 100 90 80 70 60 50 40 30 20 10 0 -10
f1 (ppm)

I-JG-23

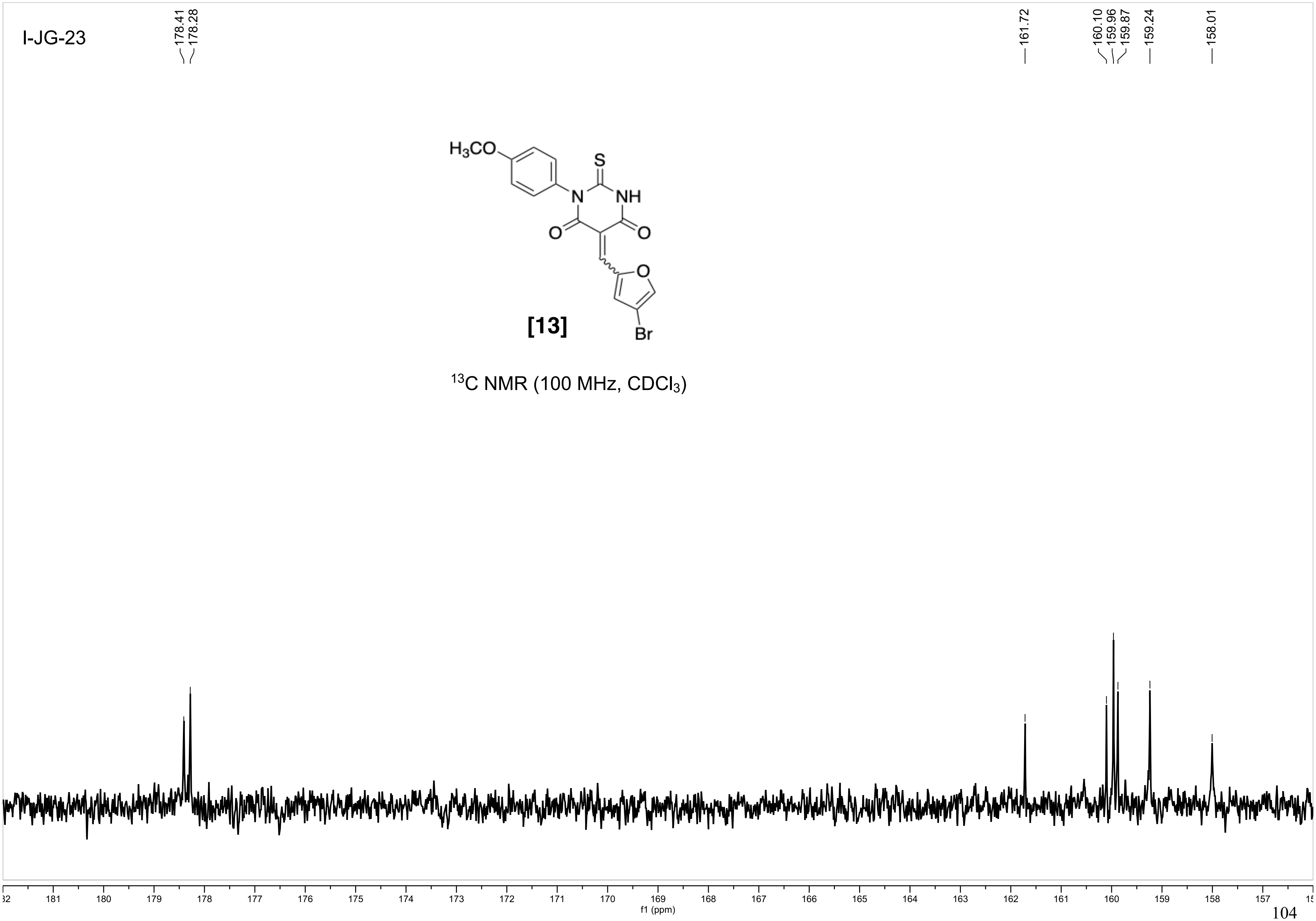

[13]

$^{13}C$ NMR (100 MHz, $CDCl_3$)

I-JG-23

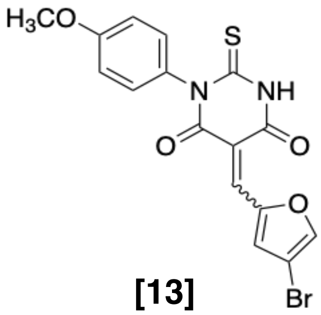

[13]

$^{13}C$ NMR (100 MHz, $CDCl_3$)

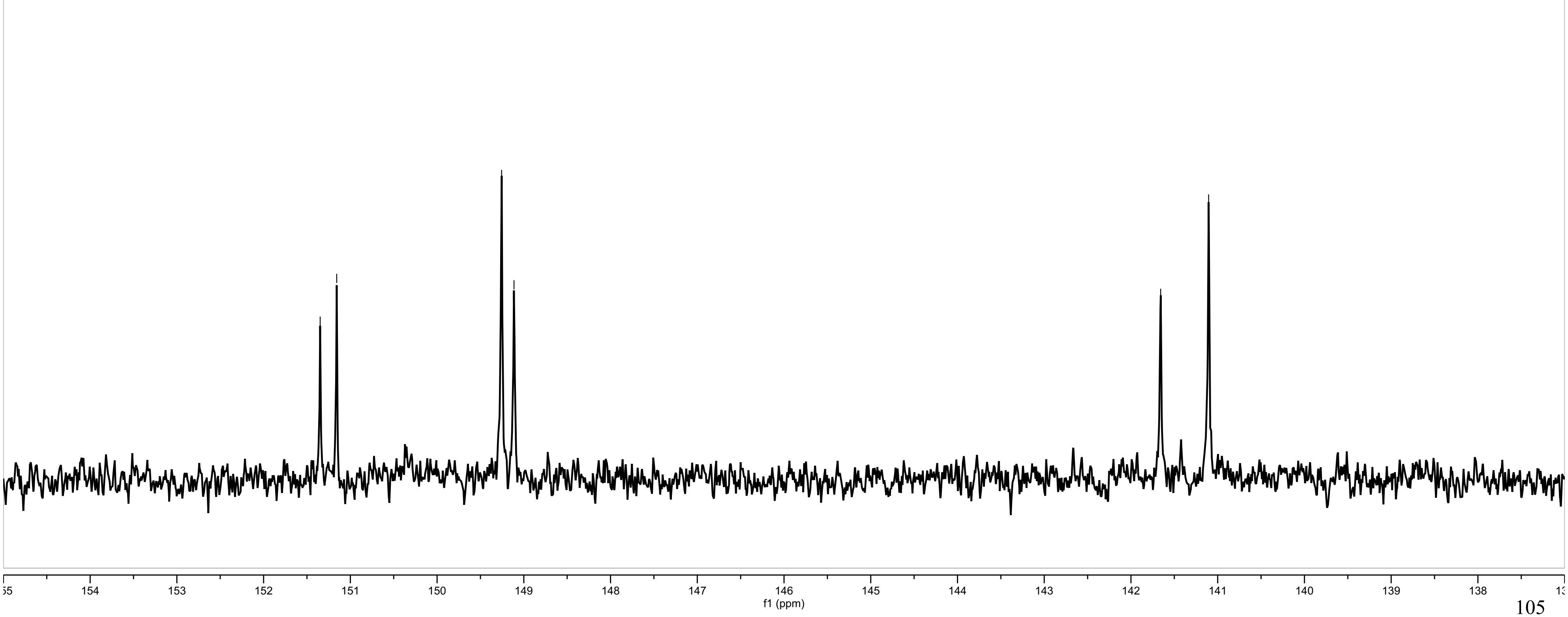

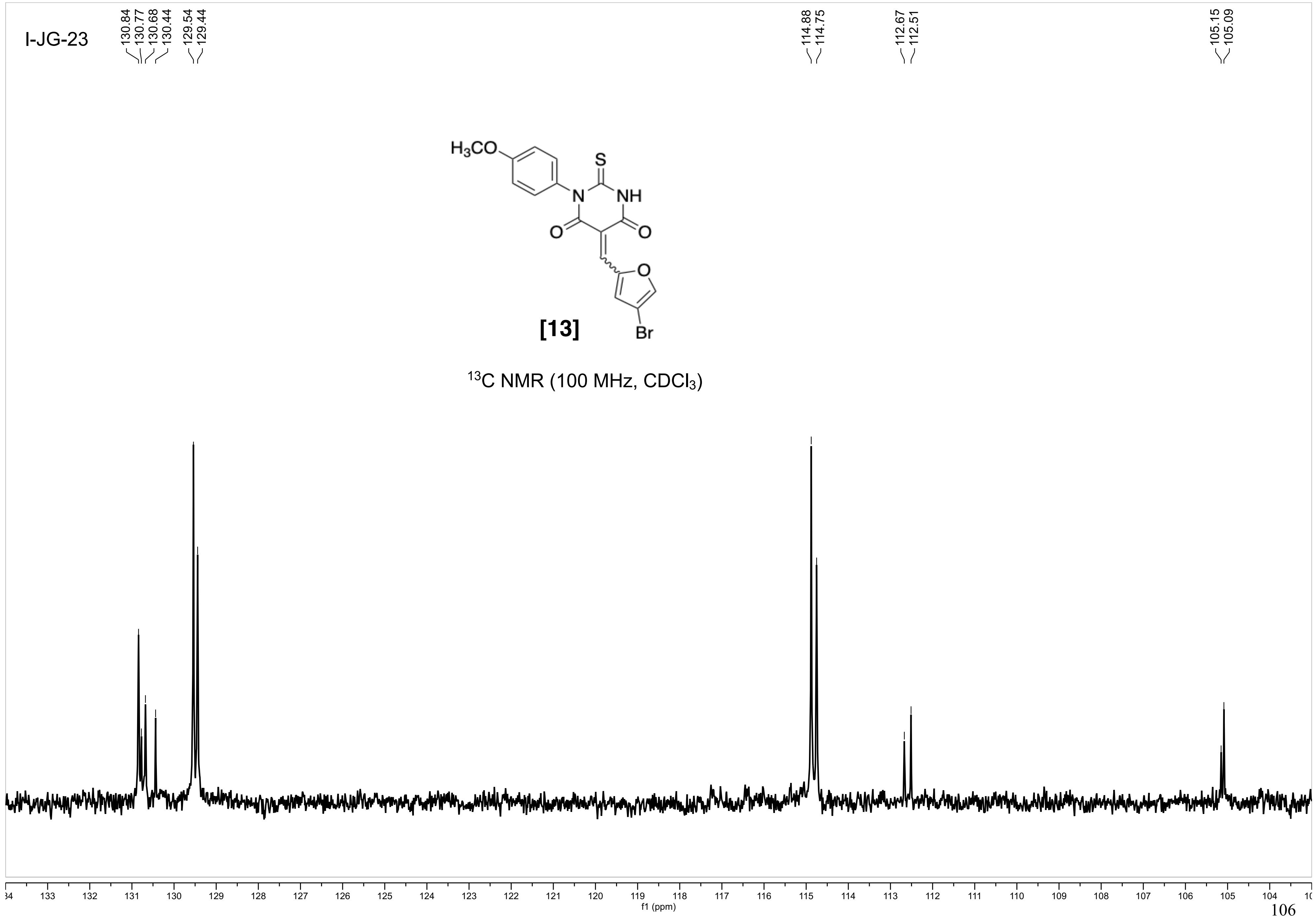
I-JG-23
130.84
130.77
130.68
130.44
129.54
129.44
114.88
114.75
112.67
112.51
105.15
105.09
H3CO
S
N
NH
O
O
O
Br
[13]
13C NMR (100 MHz, CDCl3)
134
133
132
131
130
129
128
127
126
125
124
123
122
121
120
119
118
117
116
115
114
113
112
111
110
109
108
107
106
105
104
f1 (ppm)

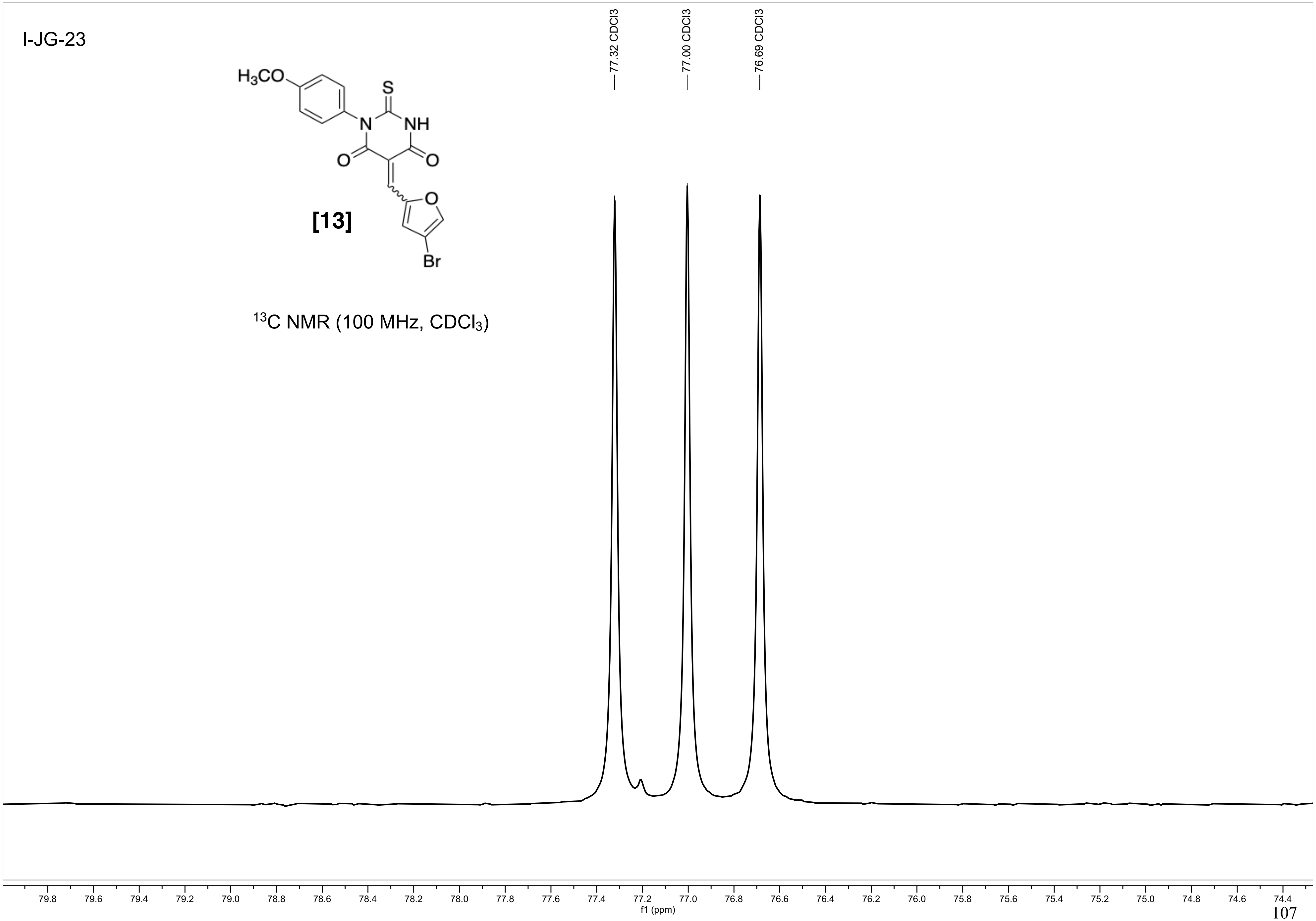
I-JG-23
H3CO
S
N
NH
O
O
O
Br
[13]
13C NMR (100 MHz, CDCl3)
77.32 CDCl3
77.00 CDCl3
76.69 CDCl3
79.8 79.6 79.4 79.2 79.0 78.8 78.6 78.4 78.2 78.0 77.8 77.6 77.4 77.2 77.0 76.8 76.6 76.4 76.2 76.0 75.8 75.6 75.4 75.2 75.0 74.8 74.6 74.4
f1 (ppm)

I-JG-23

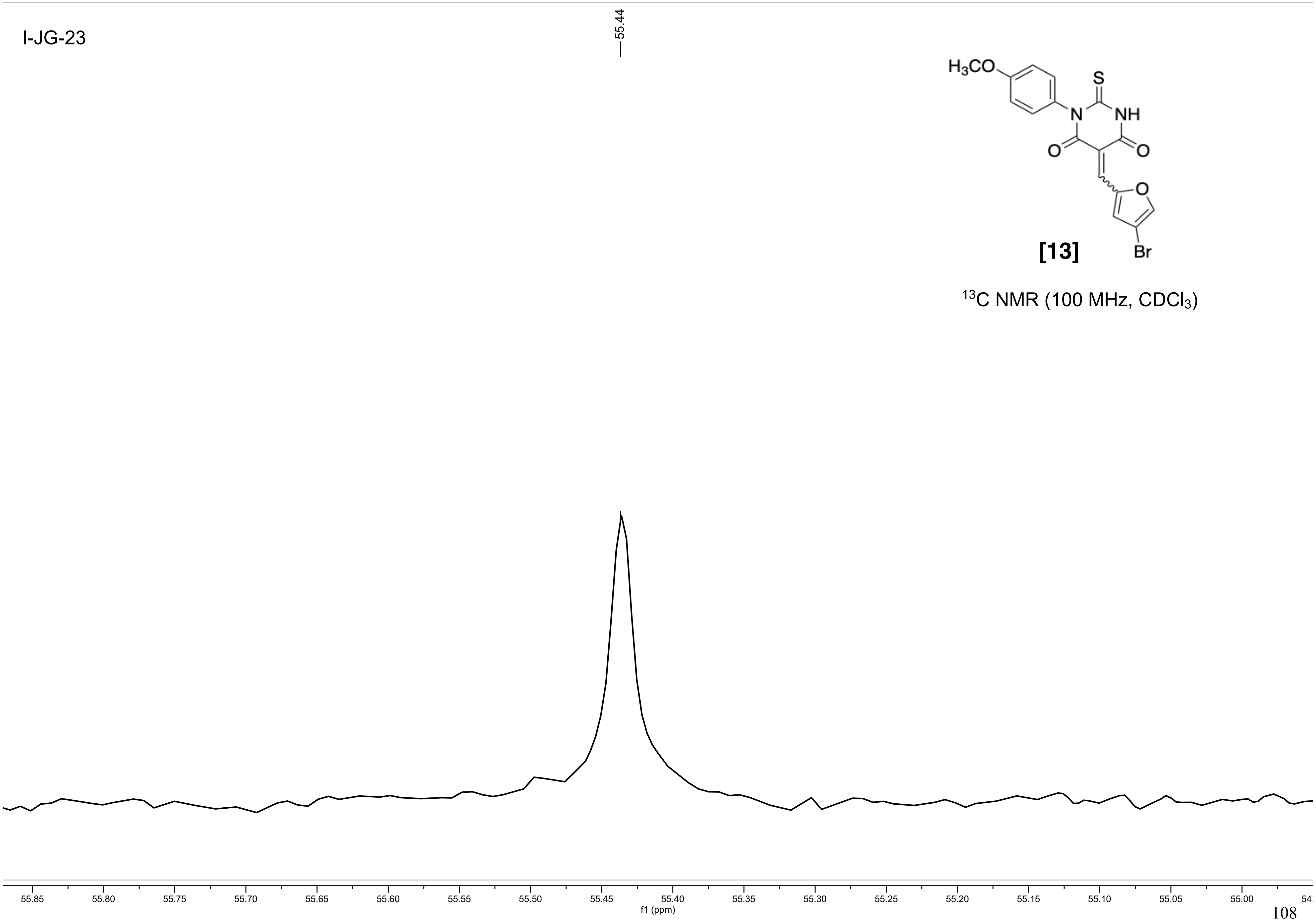
55.44
$H_3CO$
S
N
NH
O
O
O
Br
[13]
$^{13}C$ NMR (100 MHz, $CDCl_3$)
55.85
55.80
55.75
55.70
55.65
55.60
55.55
55.50
55.45
55.40
55.35
55.30
55.25
55.20
55.15
55.10
55.05
55.00
f1 (ppm)

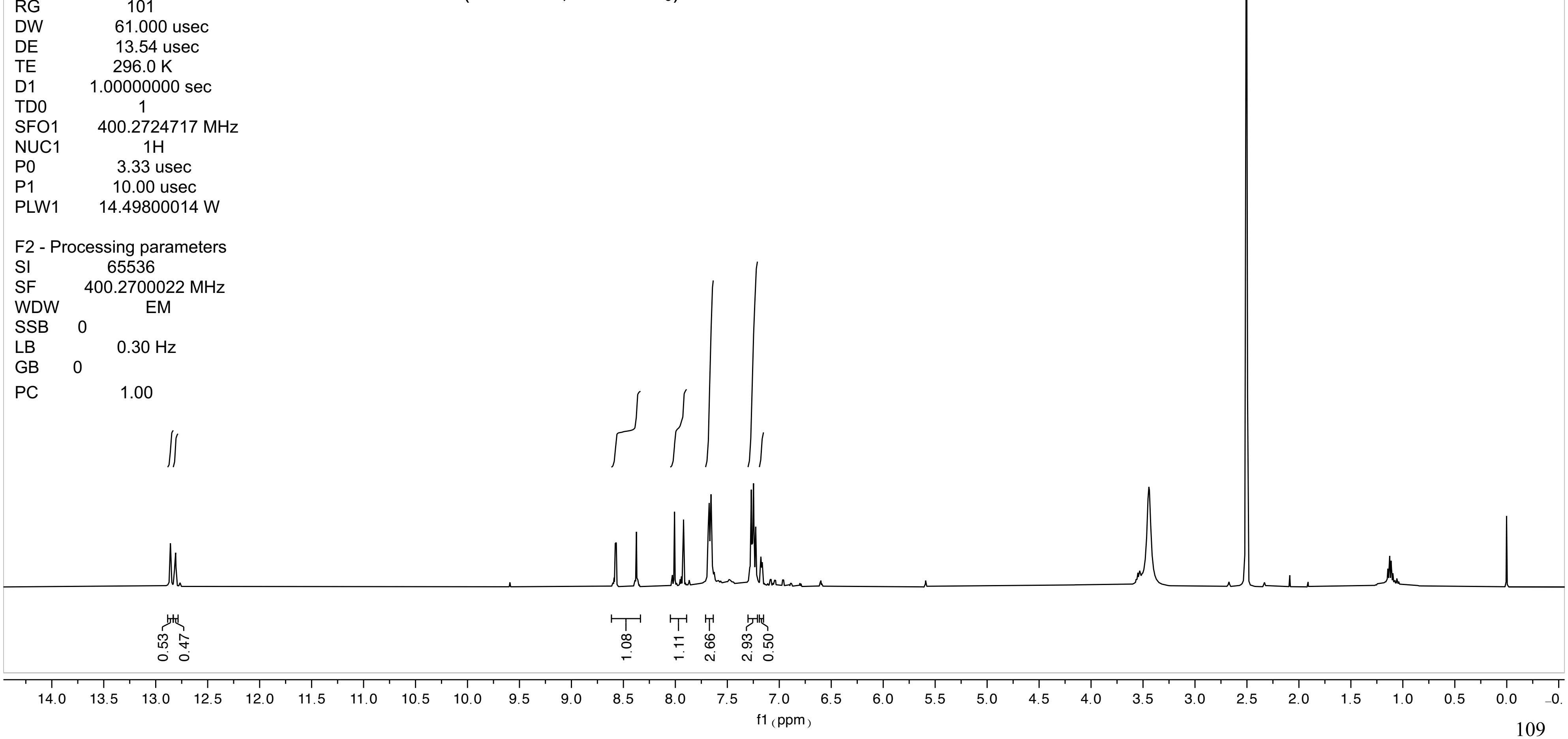

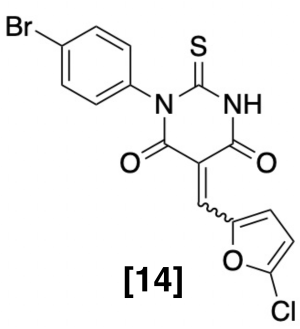

**[14]**

$^1$H NMR (400 MHz, DMSO-$d_6$)

LZ-1-14

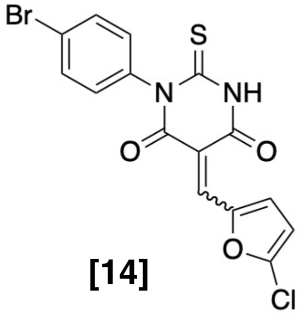

**[14]**

$^{1}$H NMR (400 MHz, DMSO-$d_6$)

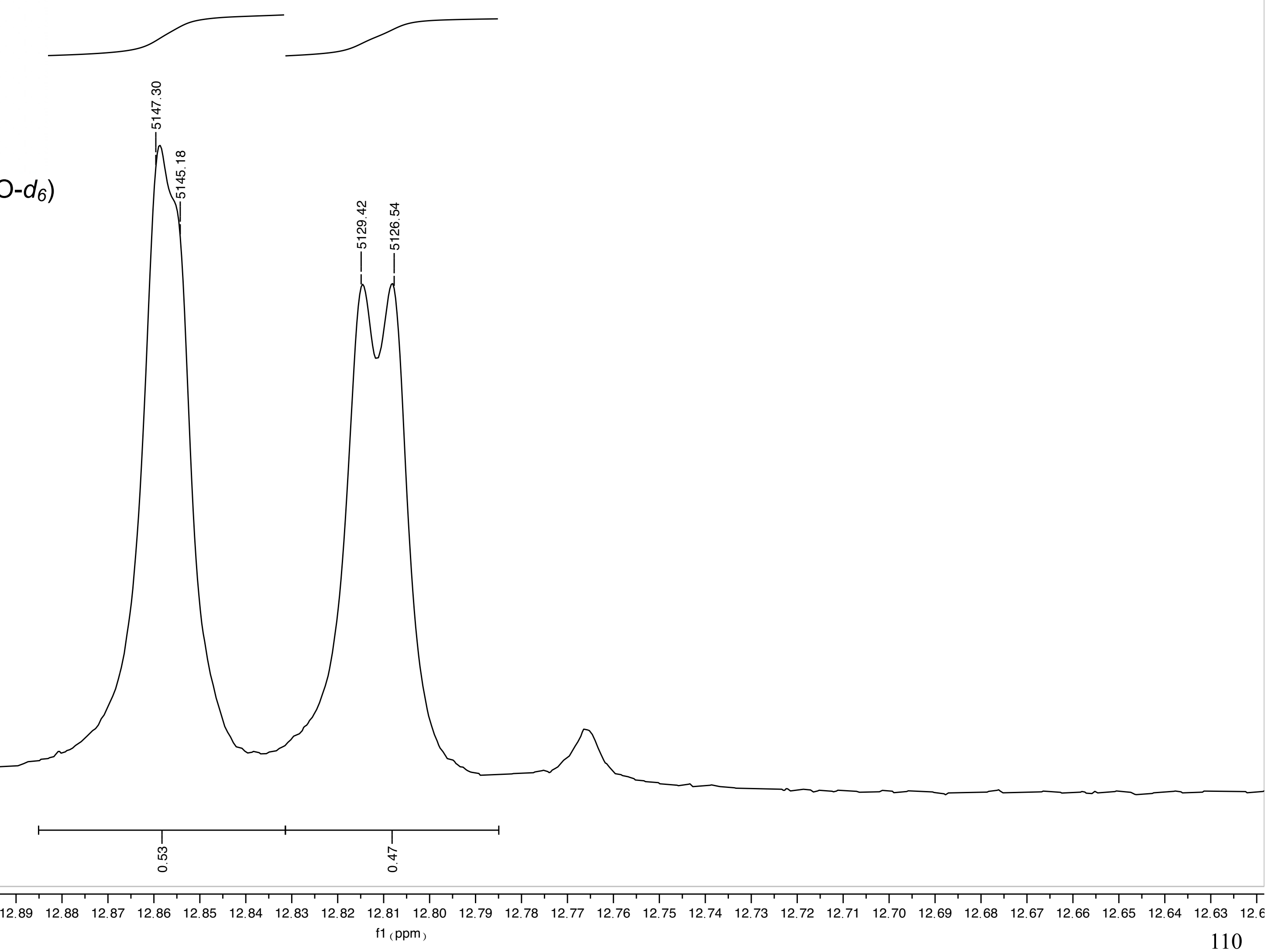

LZ-1-14

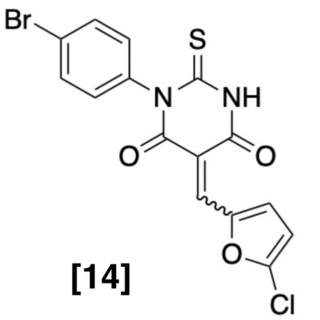

[14]

$^1$H NMR (400 MHz, DMSO-$d_6$)

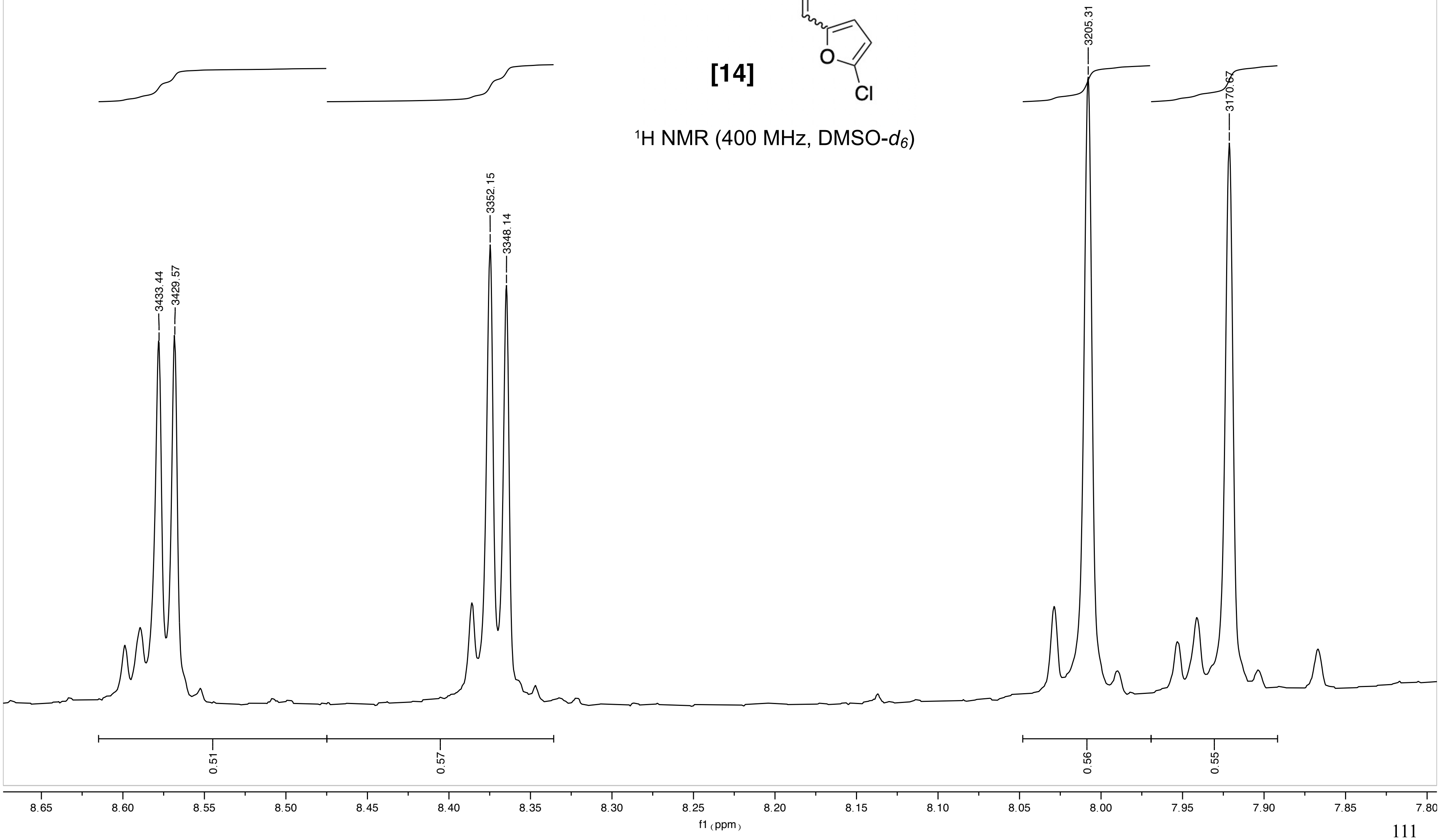

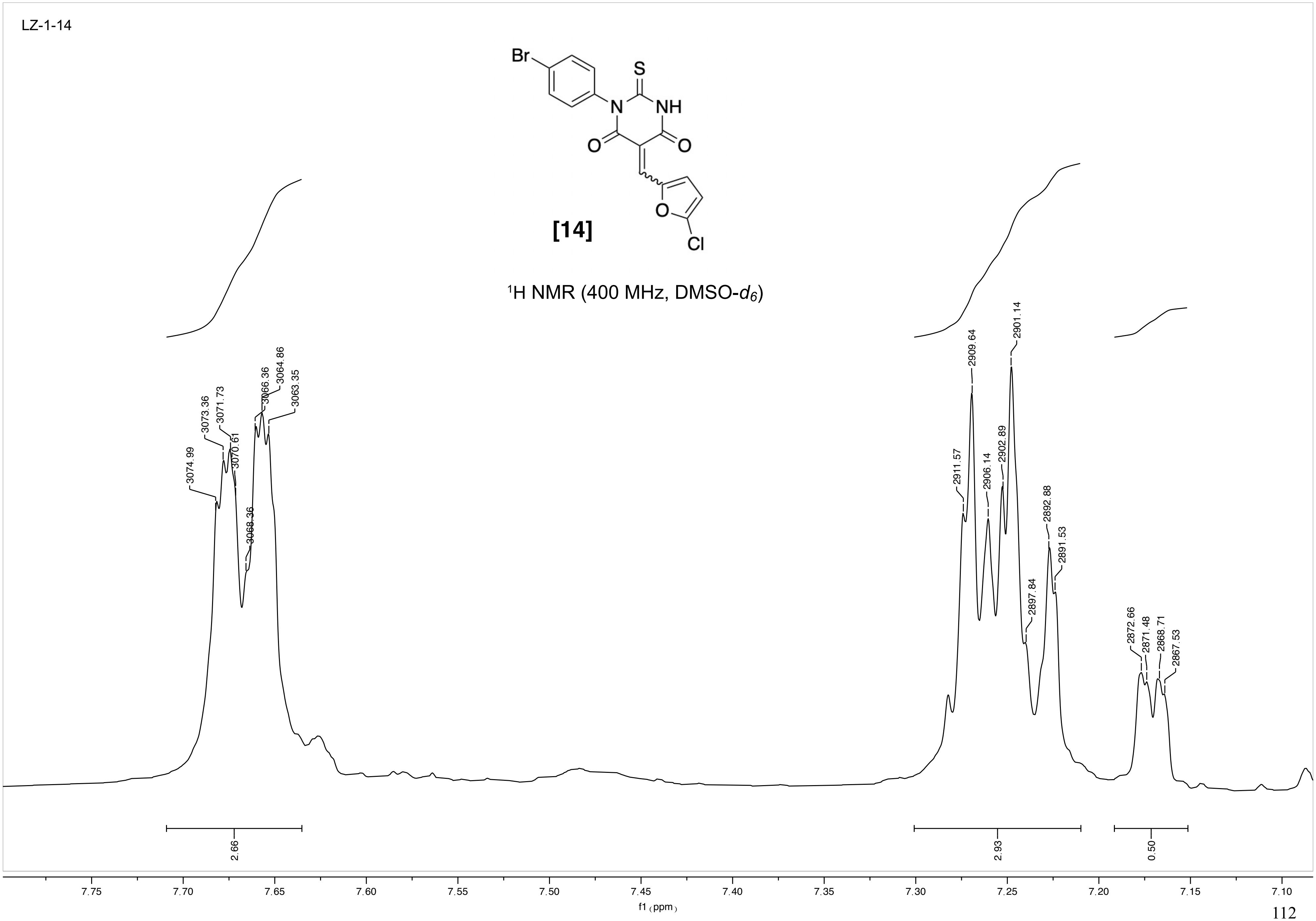
LZ-1-14
[14]
$^1$H NMR (400 MHz, DMSO-$d_6$)
3074.99
3073.36
3071.73
3070.61
3068.36
3066.36
3064.86
3063.35
2911.57
2909.64
2906.14
2902.89
2901.14
2897.84
2892.88
2891.53
2872.66
2871.48
2868.71
2867.53
2.66
2.93
0.50
7.75
7.70
7.65
7.60
7.55
7.50
7.45
7.40
7.35
7.30
7.25
7.20
7.15
7.10
f1 (ppm)

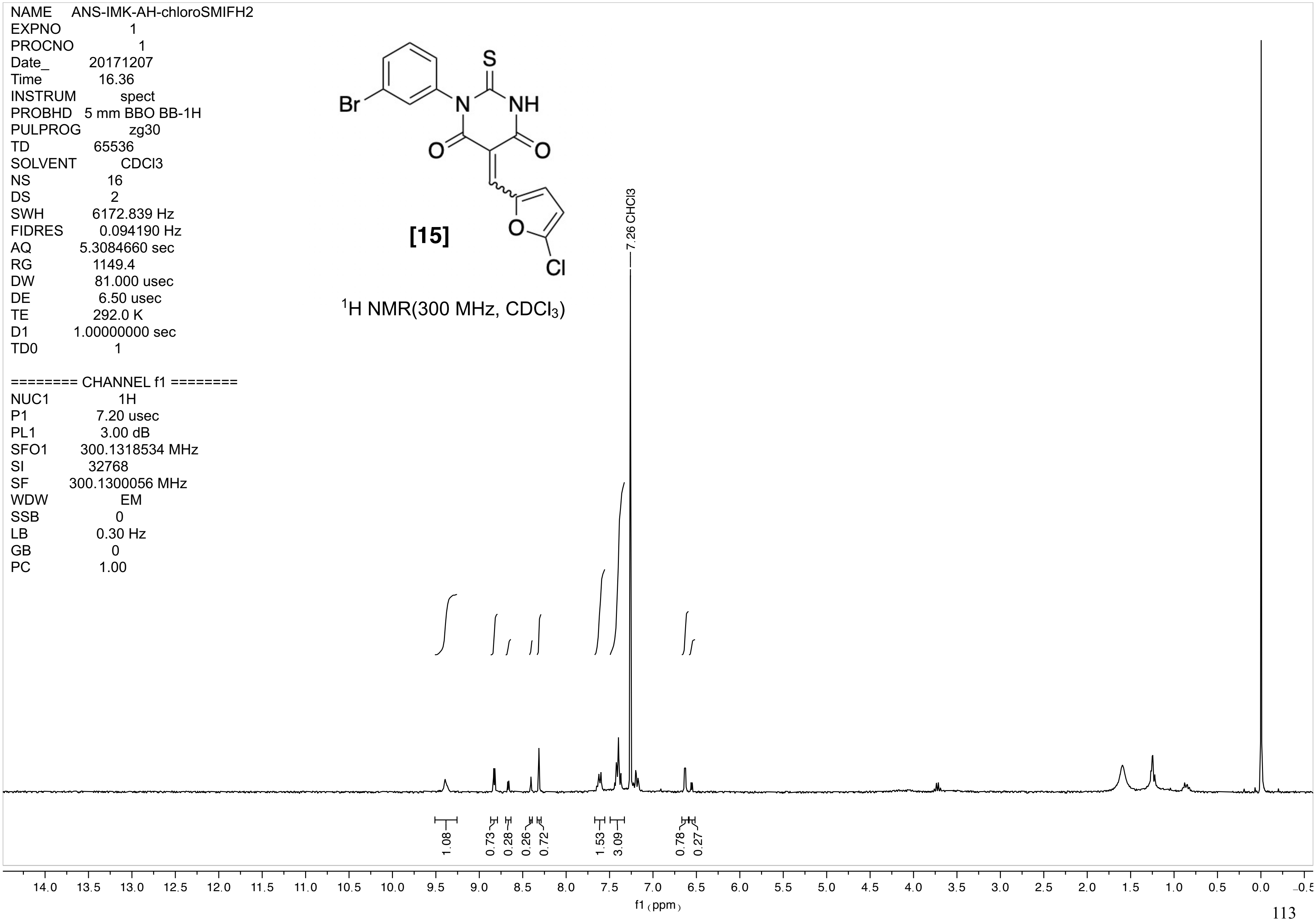
NAME ANS-IMK-AH-chloroSMIFH2
EXPNO 1
PROCNO 1
Date_ 20171207
Time 16.36
INSTRUM spect
PROBHD 5 mm BBO BB-1H
PULPROG zg30
TD 65536
SOLVENT CDCl3
NS 16
DS 2
SWH 6172.839 Hz
FIDRES 0.094190 Hz
AQ 5.3084660 sec
RG 1149.4
DW 81.000 usec
DE 6.50 usec
TE 292.0 K
D1 1.00000000 sec
TD0 1
======== CHANNEL f1 ========
NUC1 1H
P1 7.20 usec
PL1 3.00 dB
SFO1 300.1318534 MHz
SI 32768
SF 300.1300056 MHz
WDW EM
SSB 0
LB 0.30 Hz
GB 0
PC 1.00
S
Br
N
NH
O
O
O
Cl
[15]
$^1$H NMR(300 MHz, $CDCl_3$)
7.26 CHCl3
1.08
0.73
0.28
0.26
0.72
1.53
3.09
0.78
0.27
14.0 13.5 13.0 12.5 12.0 11.5 11.0 10.5 10.0 9.5 9.0 8.5 8.0 7.5 7.0 6.5 6.0 5.5 5.0 4.5 4.0 3.5 3.0 2.5 2.0 1.5 1.0 0.5 0.0 -0.5
f1 (ppm)

ANS-IMK-AH

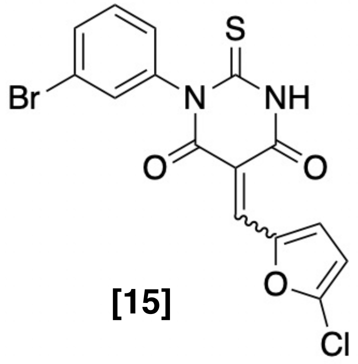

**[15]**

$^1$H NMR(300 MHz, $CDCl_3$)

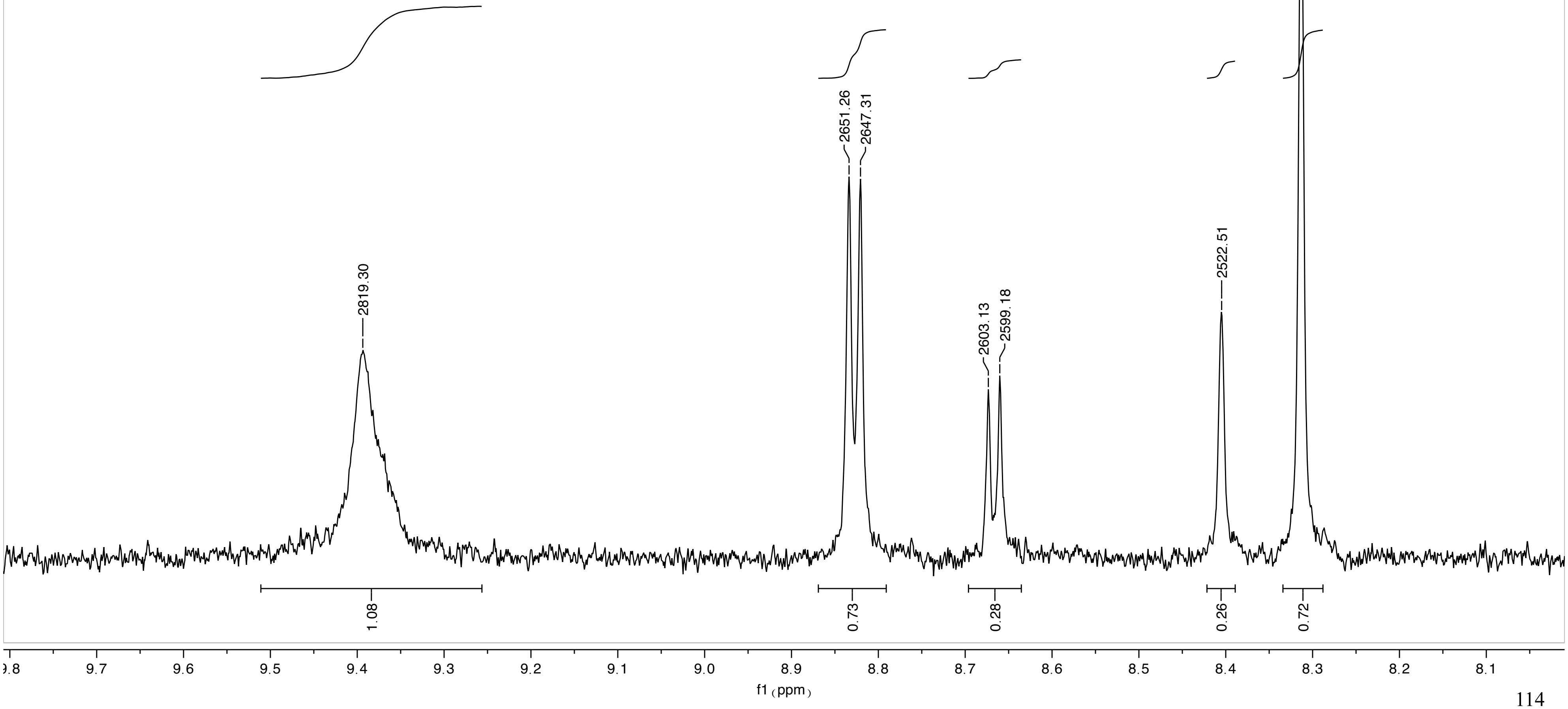

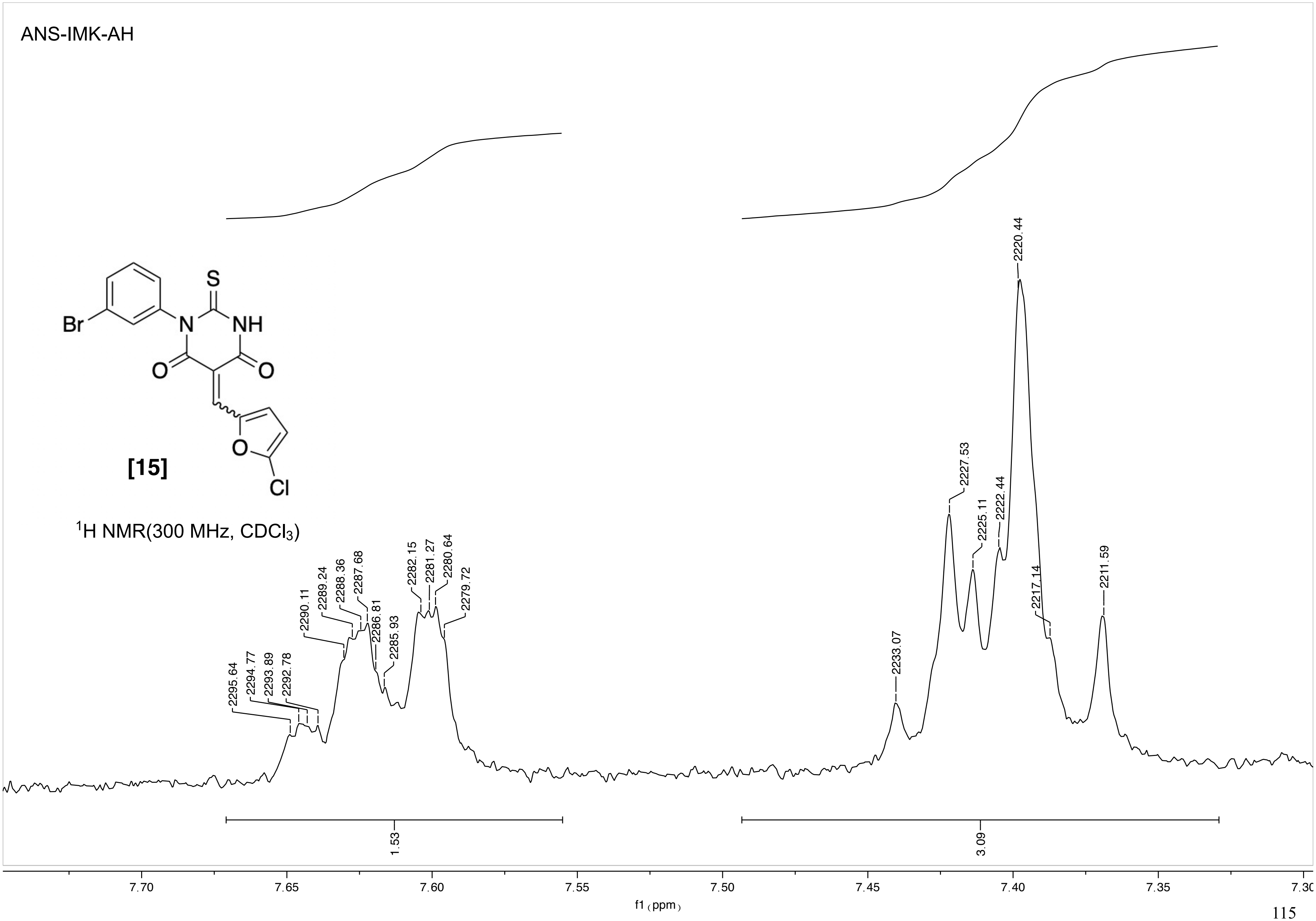
ANS-IMK-AH
Br
N
NH
S
O
O
Cl
[15]
1H NMR(300 MHz, CDCl3)
2295.64
2294.77
2293.89
2292.78
2290.11
2289.24
2288.36
2287.68
2286.81
2285.93
2282.15
2281.27
2280.64
2279.72
2233.07
2227.53
2225.11
2222.44
2220.44
2217.14
2211.59
1.53
3.09
7.70
7.65
7.60
7.55
7.50
7.45
7.40
7.35
7.30
f1 (ppm)

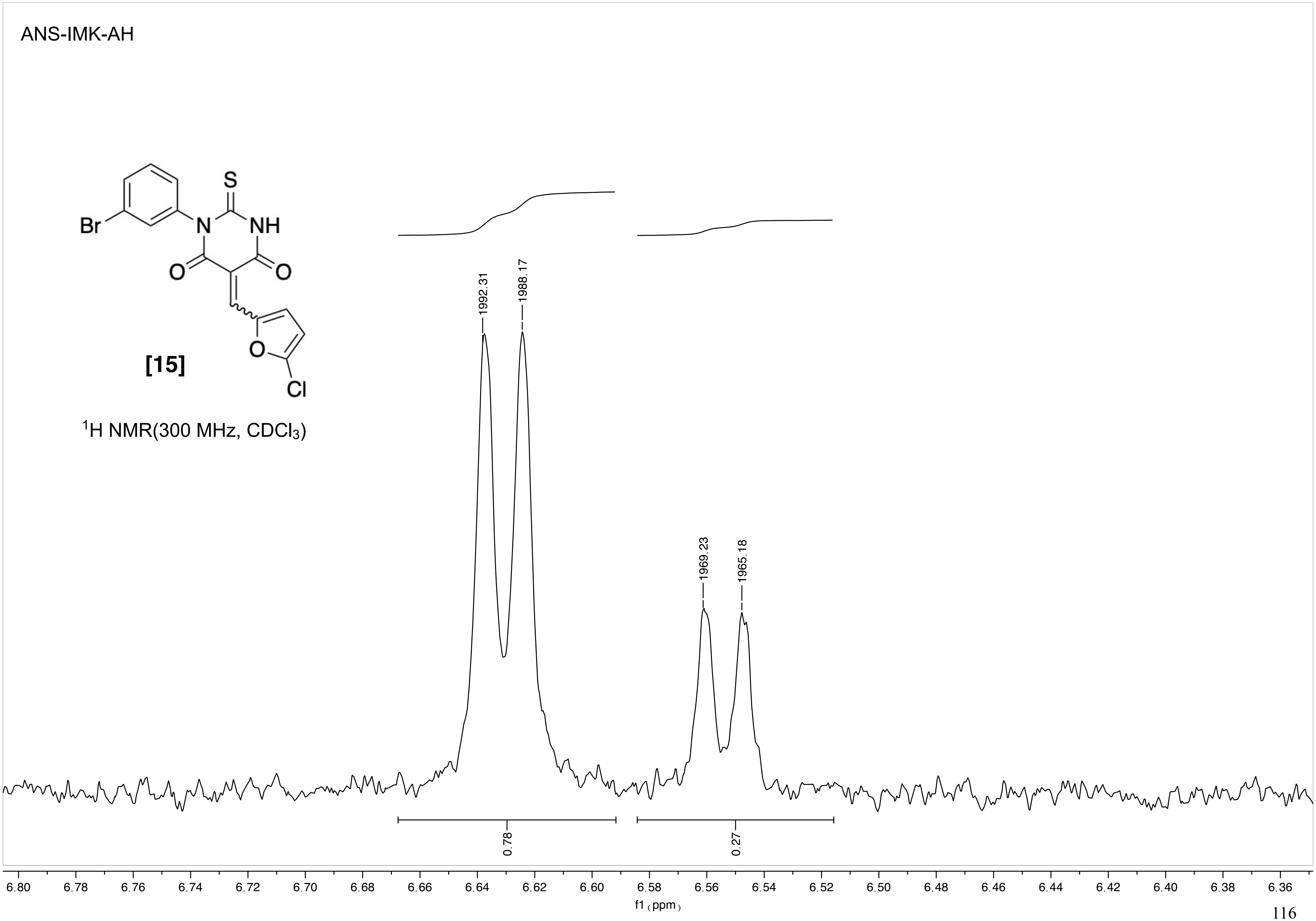

ANS-IMK-AH
[15]
Br
N
NH
S
O
O
O
Cl
1H NMR(300 MHz, CDCl3)
1992.31
1988.17
1969.23
1965.18
0.78
0.27
6.80 6.78 6.76 6.74 6.72 6.70 6.68 6.66 6.64 6.62 6.60 6.58 6.56 6.54 6.52 6.50 6.48 6.46 6.44 6.42 6.40 6.38 6.36
f1 (ppm)

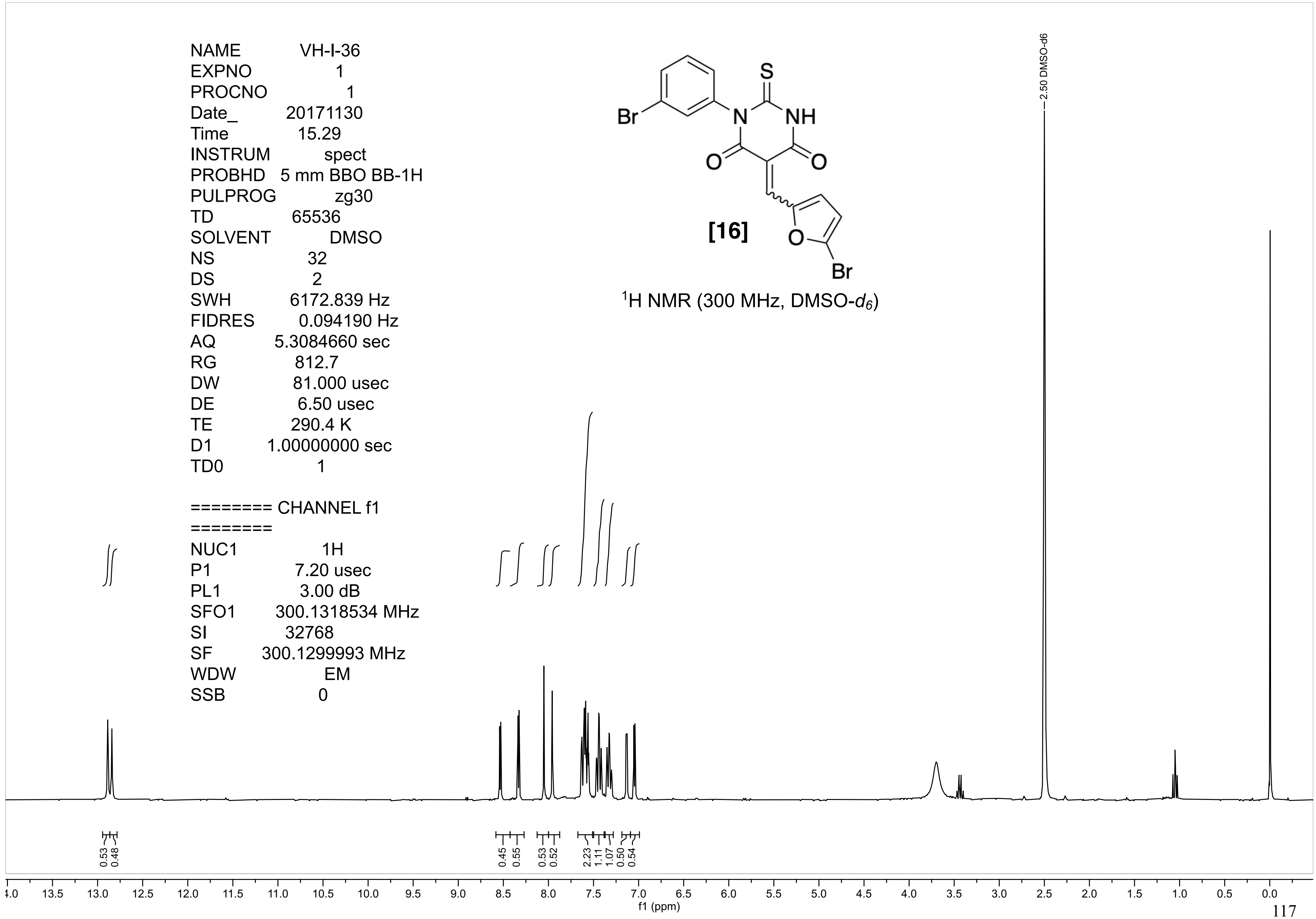
NAME VH-I-36
EXPNO 1
PROCNO 1
Date_ 20171130
Time 15.29
INSTRUM spect
PROBHD 5 mm BBO BB-1H
PULPROG zg30
TD 65536
SOLVENT DMSO
NS 32
DS 2
SWH 6172.839 Hz
FIDRES 0.094190 Hz
AQ 5.3084660 sec
RG 812.7
DW 81.000 usec
DE 6.50 usec
TE 290.4 K
D1 1.00000000 sec
TD0 1
======== CHANNEL f1 ========
NUC1 1H
P1 7.20 usec
PL1 3.00 dB
SFO1 300.1318534 MHz
SI 32768
SF 300.1299993 MHz
WDW EM
SSB 0
S
NH
N
Br
O
O
O
Br
[16]
1H NMR (300 MHz, DMSO-d6)
2.50 DMSO-d6
0.53
0.48
0.45
0.55
0.53
0.52
2.23
1.11
1.07
0.50
0.54
f1 (ppm)

VH-I-36

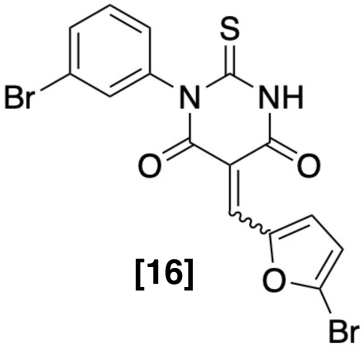

**[16]**

$^1$H NMR (300 MHz, DMSO-$d_6$)

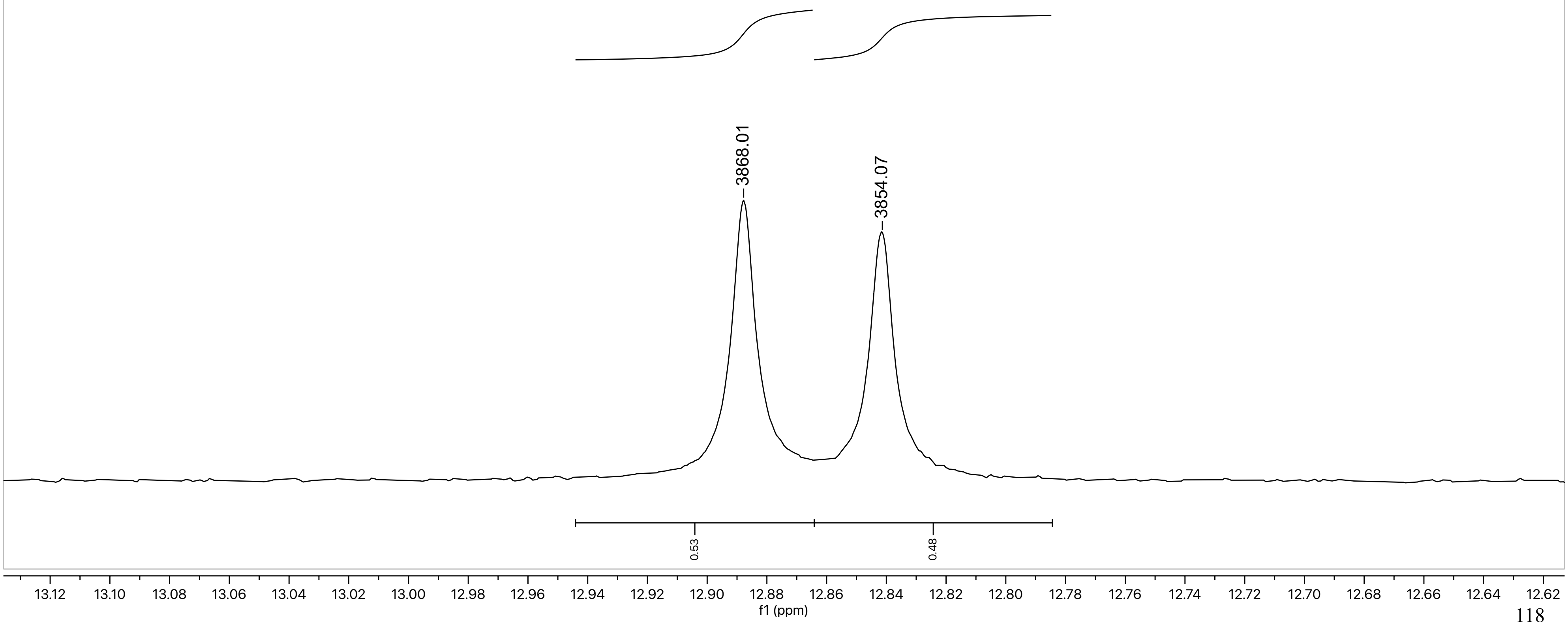

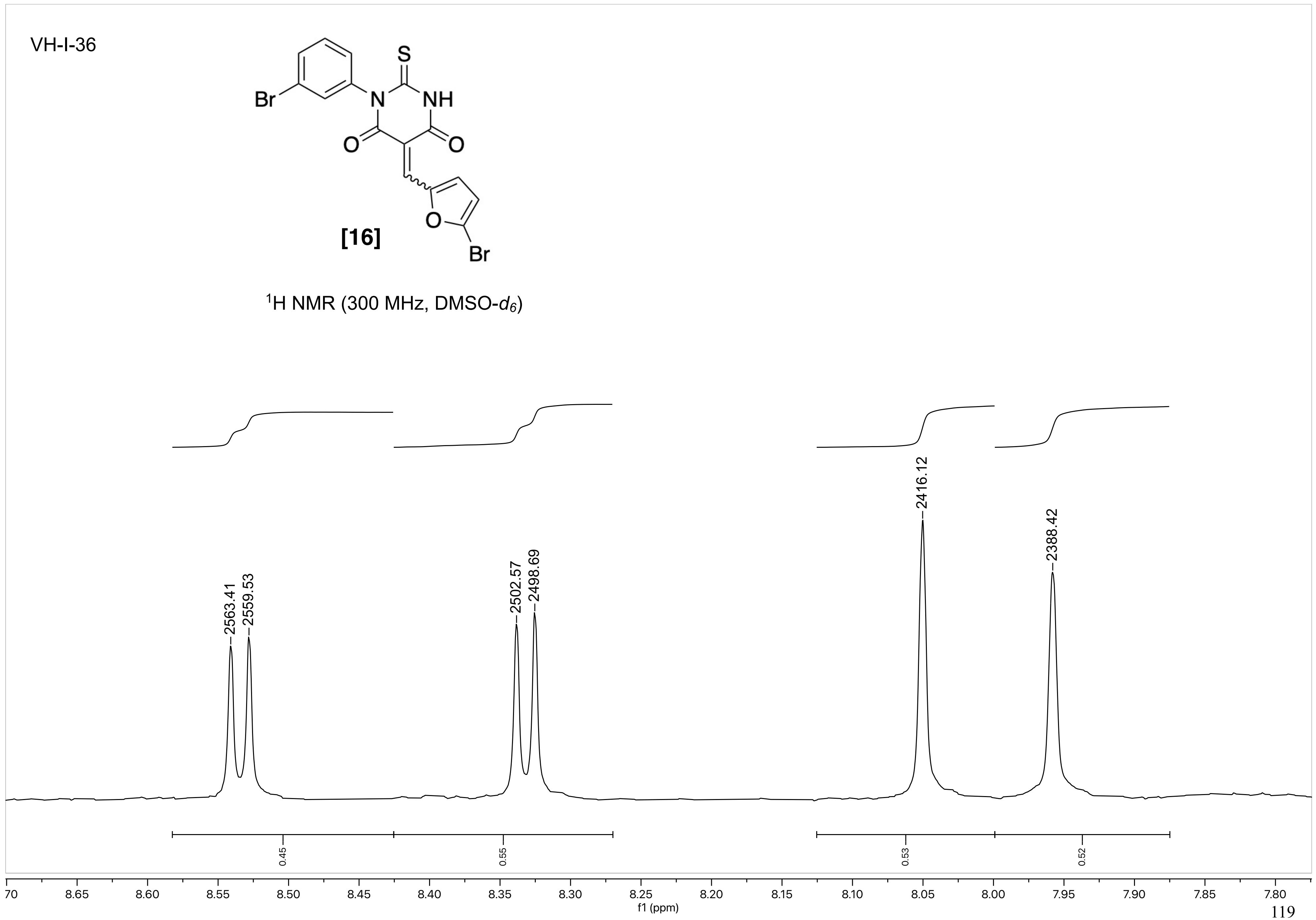
VH-I-36
S
Br
N
NH
O
O
O
Br
[16]
1H NMR (300 MHz, DMSO-d6)
2563.41
2559.53
2502.57
2498.69
2416.12
2388.42
0.45
0.55
0.53
0.52
70
8.65
8.60
8.55
8.50
8.45
8.40
8.35
8.30
8.25
8.20
8.15
8.10
8.05
8.00
7.95
7.90
7.85
7.80
f1 (ppm)

VH-I-36

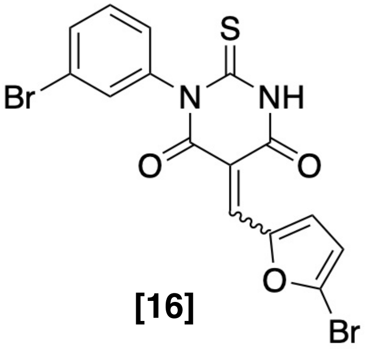

**[16]**

$^1$H NMR (300 MHz, DMSO-$d_6$)

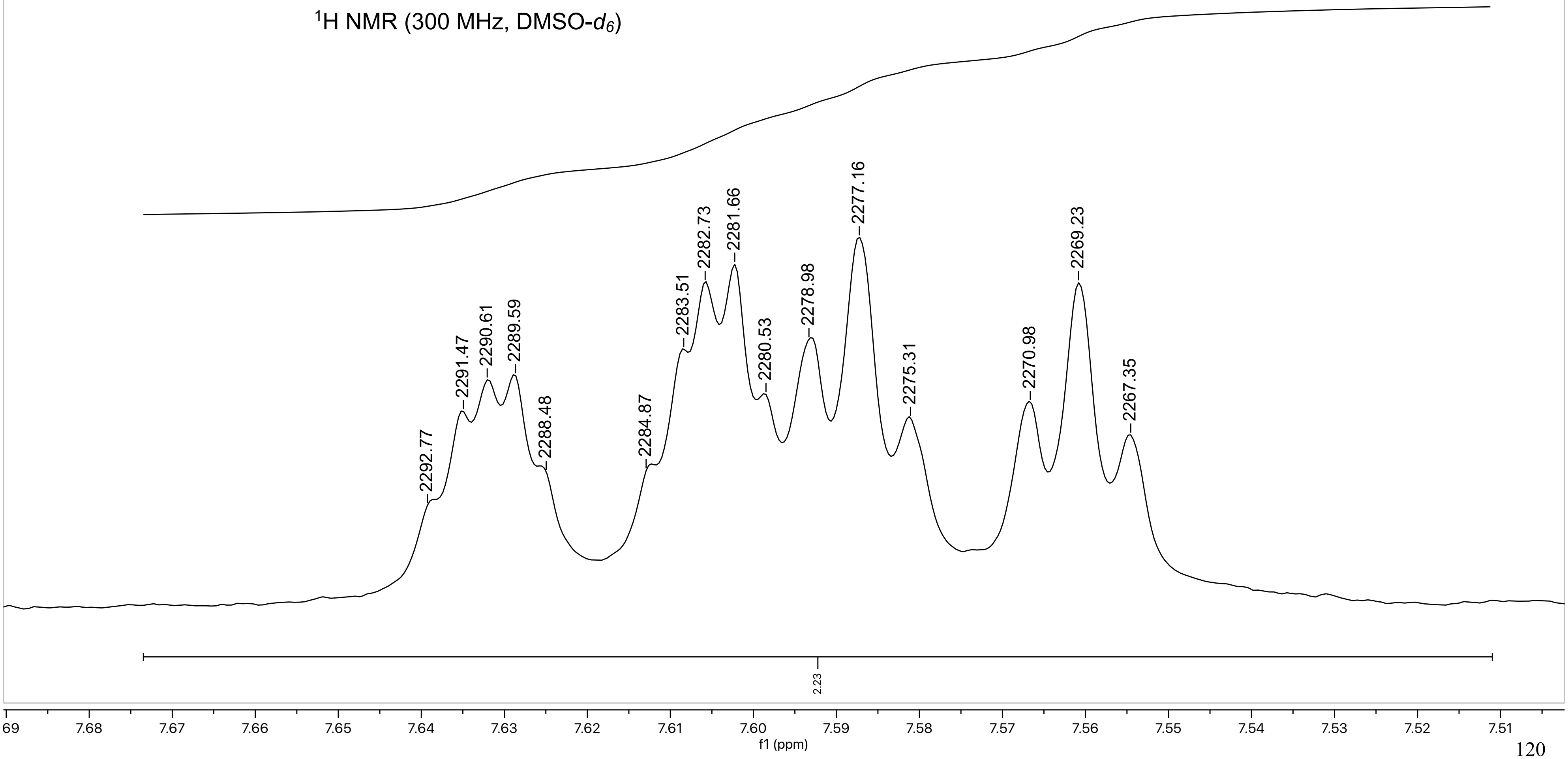

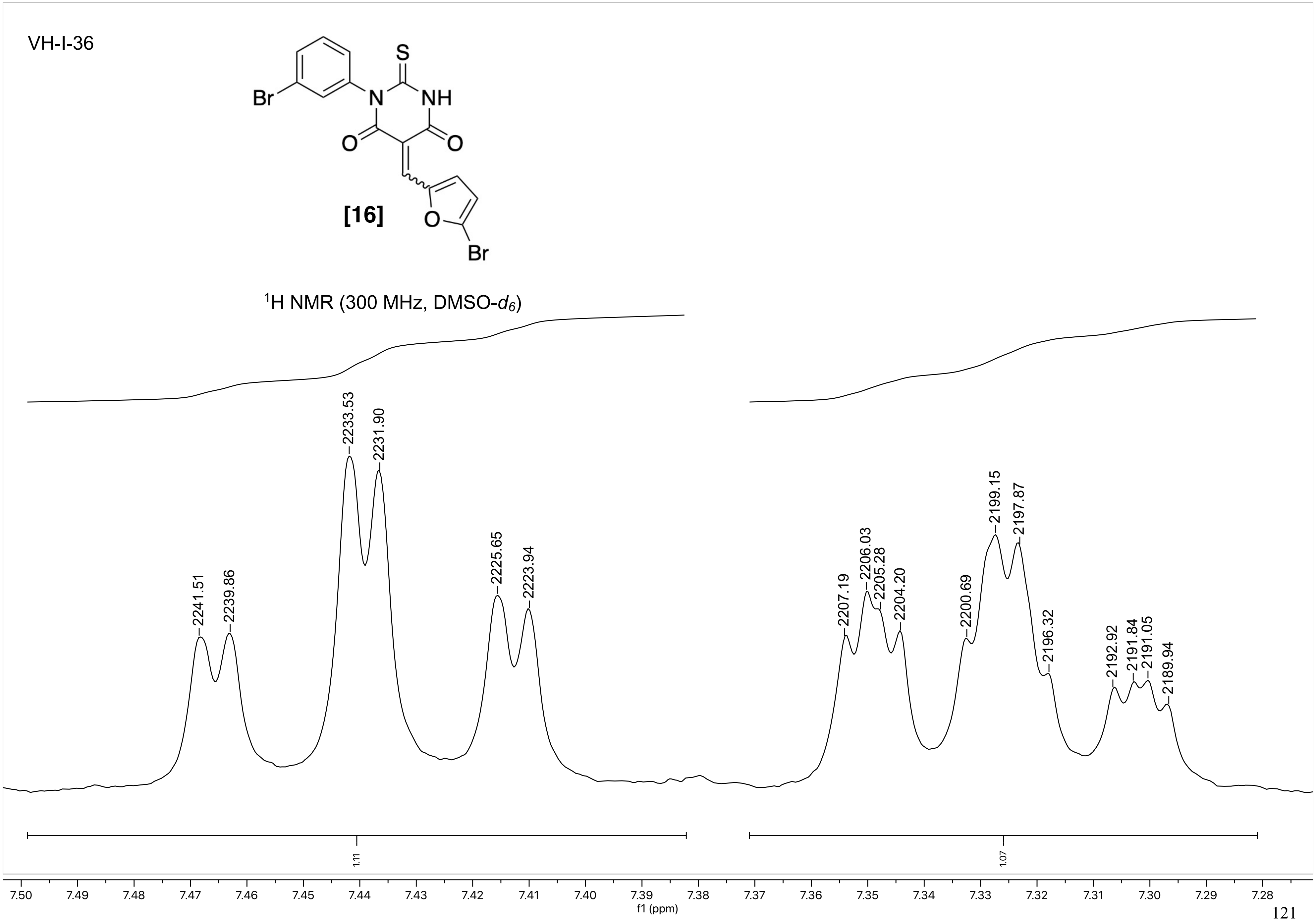
VH-I-36
S
Br
N
NH
O
O
[16]
O
Br
1H NMR (300 MHz, DMSO-d6)
2241.51
2239.86
2233.53
2231.90
2225.65
2223.94
2207.19
2206.03
2205.28
2204.20
2200.69
2199.15
2197.87
2196.32
2192.92
2191.84
2191.05
2189.94
1.11
1.07
7.50 7.49 7.48 7.47 7.46 7.45 7.44 7.43 7.42 7.41 7.40 7.39 7.38 7.37 7.36 7.35 7.34 7.33 7.32 7.31 7.30 7.29 7.28
f1 (ppm)

VH-I-36

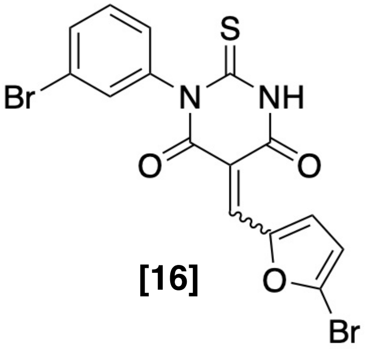

**[16]**

$^{1}H$ NMR (300 MHz, DMSO-$d_6$)

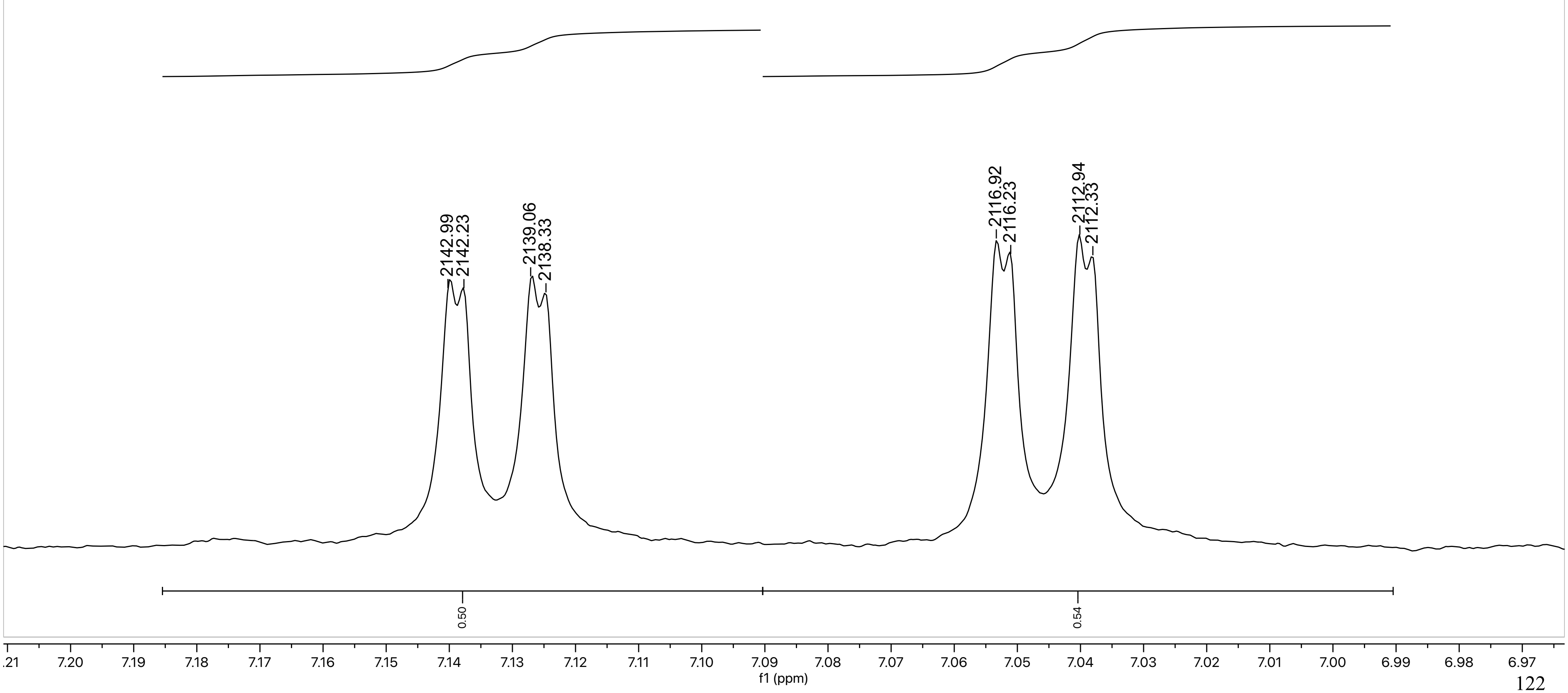

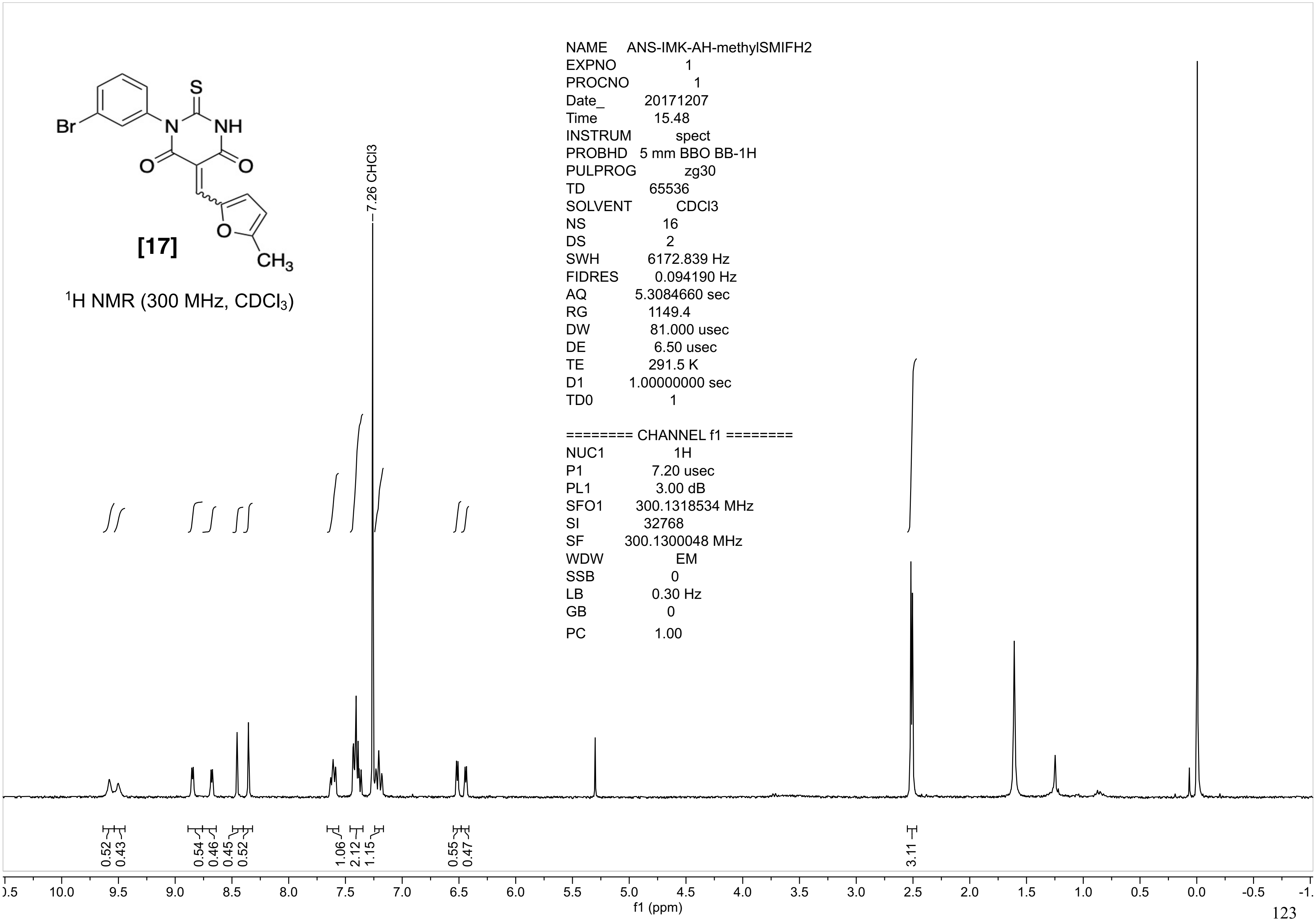

[17]
1H NMR (300 MHz, CDCl3)
7.26 CHCl3
NAME ANS-IMK-AH-methylSMIFH2
EXPNO 1
PROCNO 1
Date_ 20171207
Time 15.48
INSTRUM spect
PROBHD 5 mm BBO BB-1H
PULPROG zg30
TD 65536
SOLVENT CDCl3
NS 16
DS 2
SWH 6172.839 Hz
FIDRES 0.094190 Hz
AQ 5.3084660 sec
RG 1149.4
DW 81.000 usec
DE 6.50 usec
TE 291.5 K
D1 1.00000000 sec
TD0 1
======== CHANNEL f1 ========
NUC1 1H
P1 7.20 usec
PL1 3.00 dB
SFO1 300.1318534 MHz
SI 32768
SF 300.1300048 MHz
WDW EM
SSB 0
LB 0.30 Hz
GB 0
PC 1.00
0.52 0.43 0.54 0.46 0.45 0.52 1.06 2.12 1.15 0.55 0.47 3.11
f1 (ppm)

ANS-IMK-AH-methylSMMIFH2

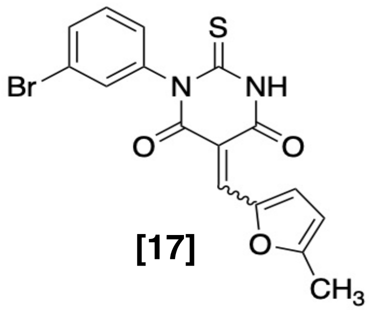

**[17]**

$^{1}H$ NMR (300 MHz, $CDCl_3$)

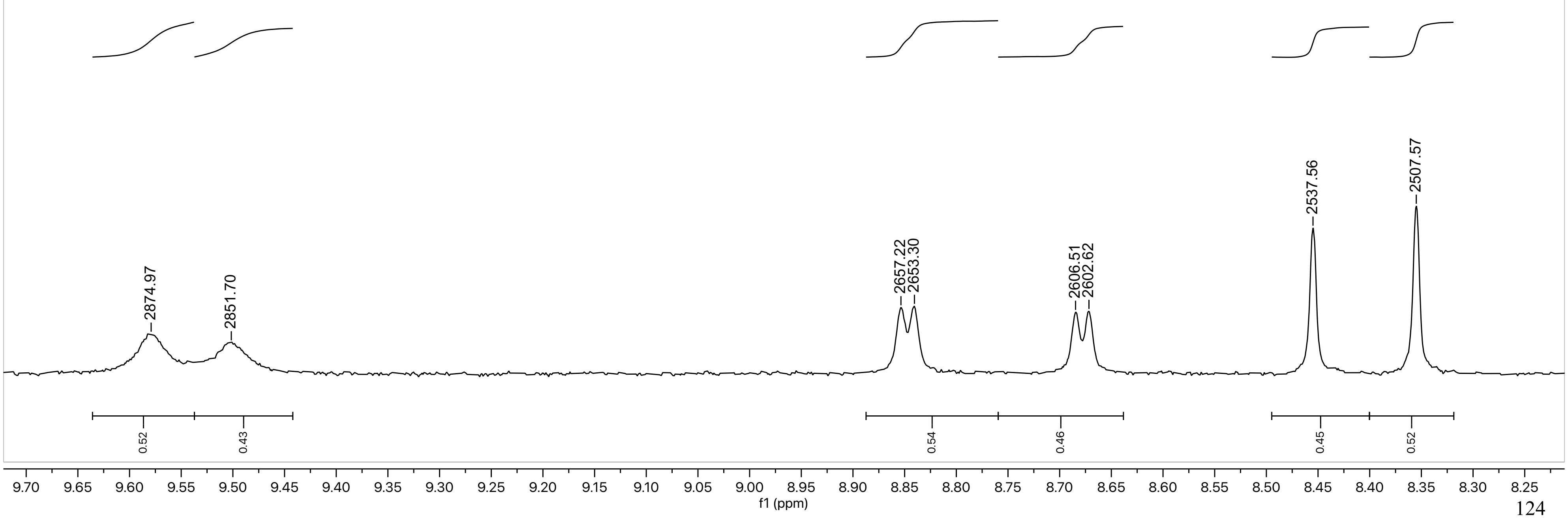

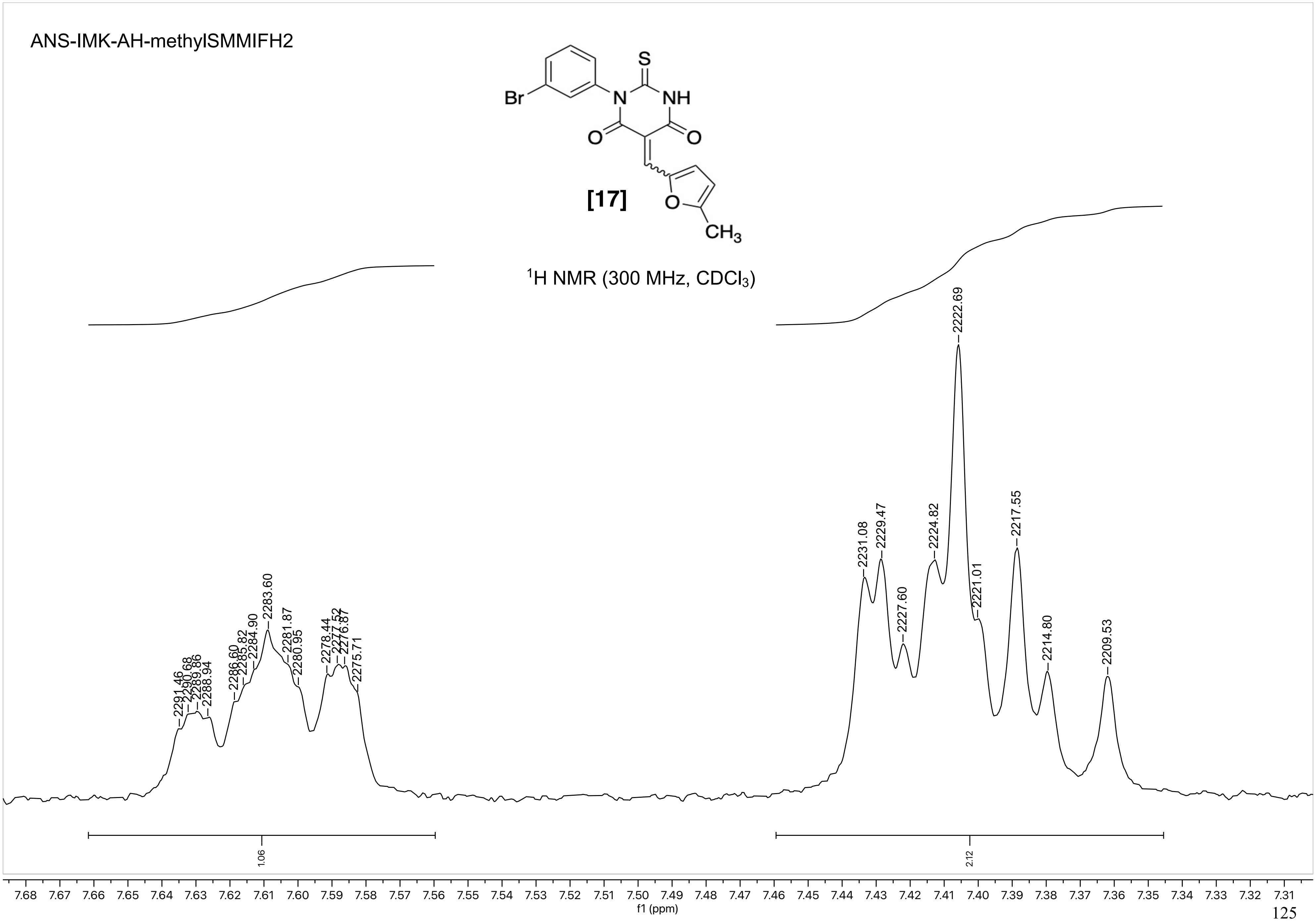
ANS-IMK-AH-methylSMMIFH2
[17]
$^{1}$H NMR (300 MHz, $CDCl_3$)
2291.46
2290.68
2289.86
2288.94
2286.60
2285.82
2284.90
2283.60
2281.87
2280.95
2278.44
2277.52
2276.87
2275.71
2231.08
2229.47
2227.60
2224.82
2222.69
2221.01
2217.55
2214.80
2209.53
1.06
2.12
f1 (ppm)

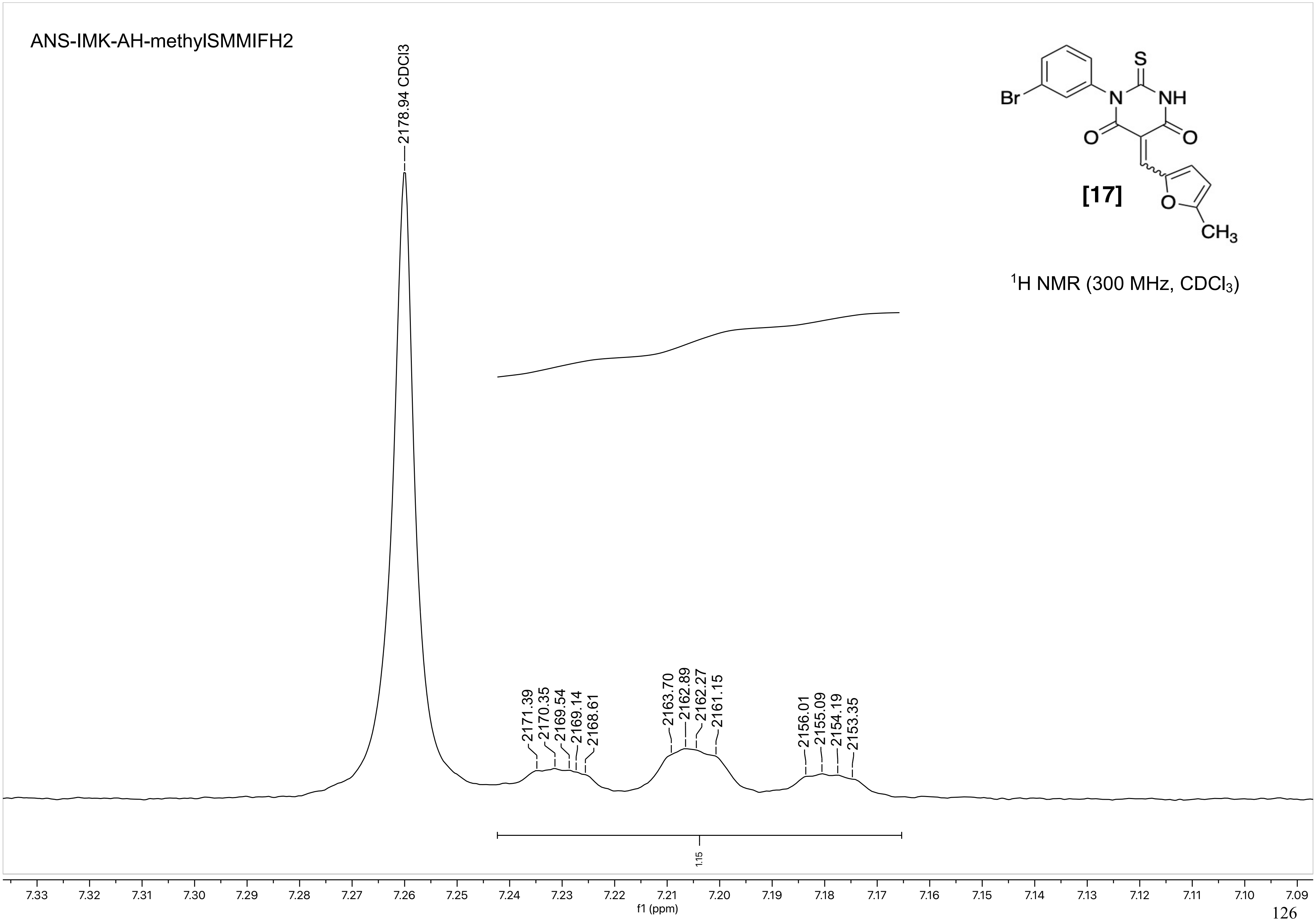
ANS-IMK-AH-methylSMMIFH2
2178.94 CDCl3
2171.39
2170.35
2169.54
2169.14
2168.61
2163.70
2162.89
2162.27
2161.15
2156.01
2155.09
2154.19
2153.35
Br
N
S
NH
O
O
O
CH3
[17]
1H NMR (300 MHz, CDCl3)
1.15
7.33 7.32 7.31 7.30 7.29 7.28 7.27 7.26 7.25 7.24 7.23 7.22 7.21 7.20 7.19 7.18 7.17 7.16 7.15 7.14 7.13 7.12 7.11 7.10 7.09
f1 (ppm)

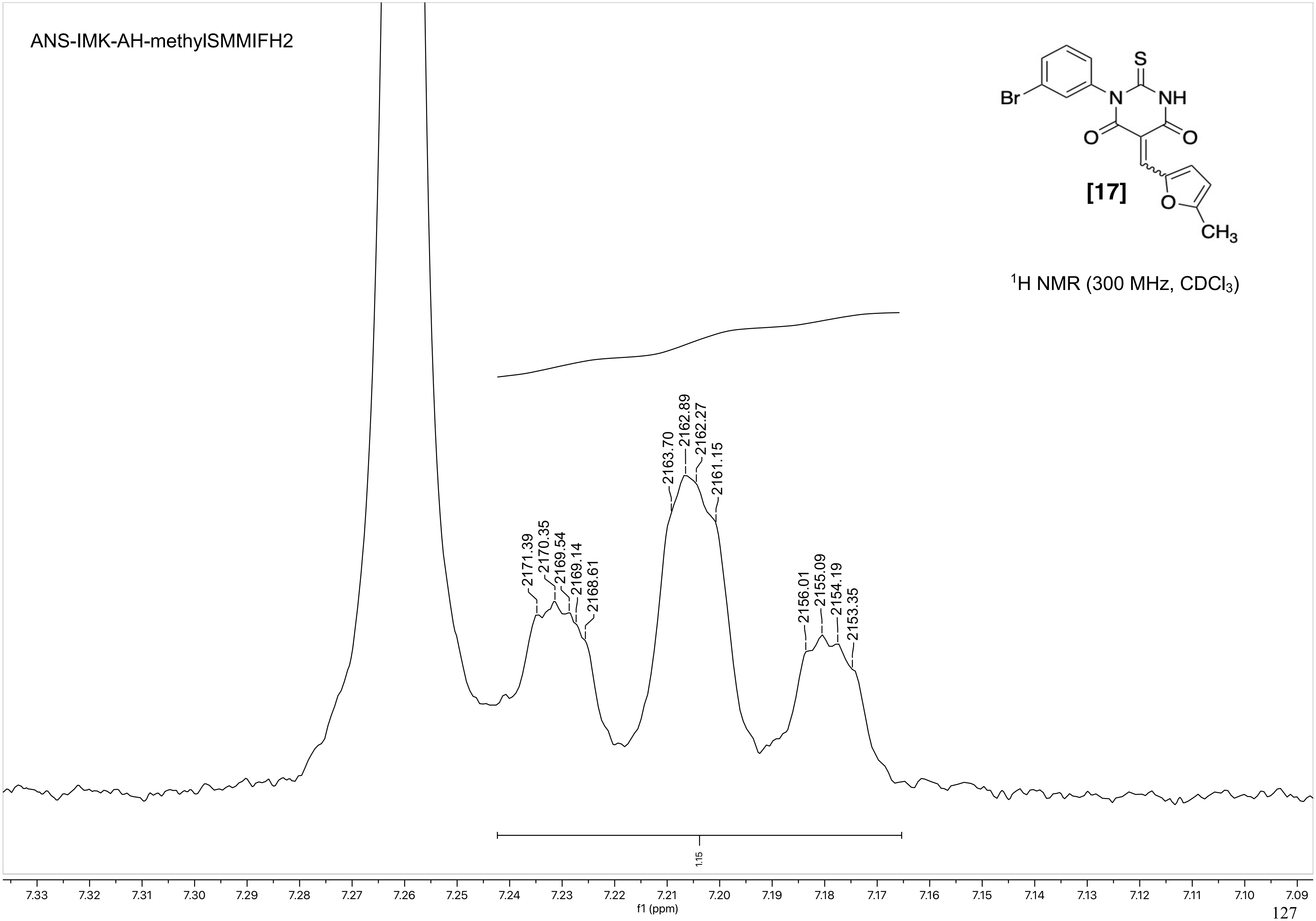
ANS-IMK-AH-methylSMMIFH2
Br
N
S
NH
O
O
[17]
O
CH3
1H NMR (300 MHz, CDCl3)
2171.39
2170.35
2169.54
2169.14
2168.61
2163.70
2162.89
2162.27
2161.15
2156.01
2155.09
2154.19
2153.35
1.15
7.33
7.32
7.31
7.30
7.29
7.28
7.27
7.26
7.25
7.24
7.23
7.22
7.21
7.20
7.19
7.18
7.17
7.16
7.15
7.14
7.13
7.12
7.11
7.10
7.09
f1 (ppm)

ANS-IMK-AH-methylSMMIFH2

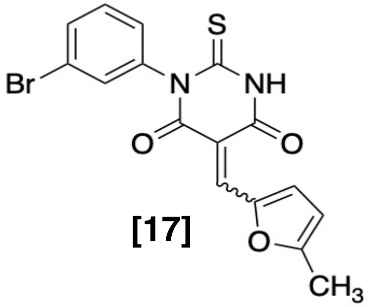

[17]

$^1$H NMR (300 MHz, $CDCl_3$)

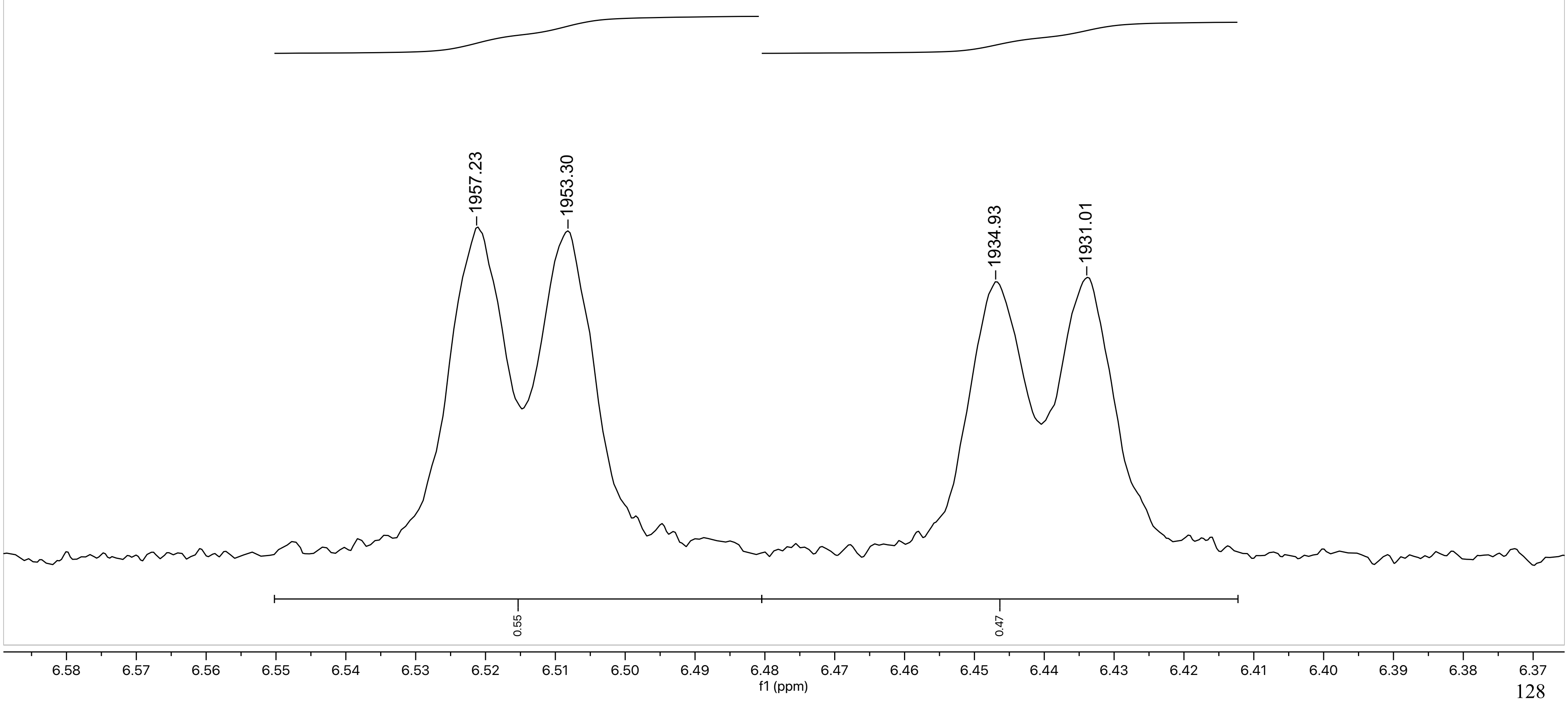

ANS-IMK-AH-methylSMMIFH2

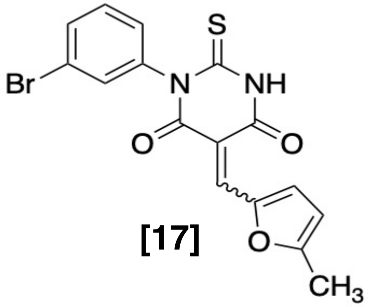

[17]

$^{1}$H NMR (300 MHz, $CDCl_3$)

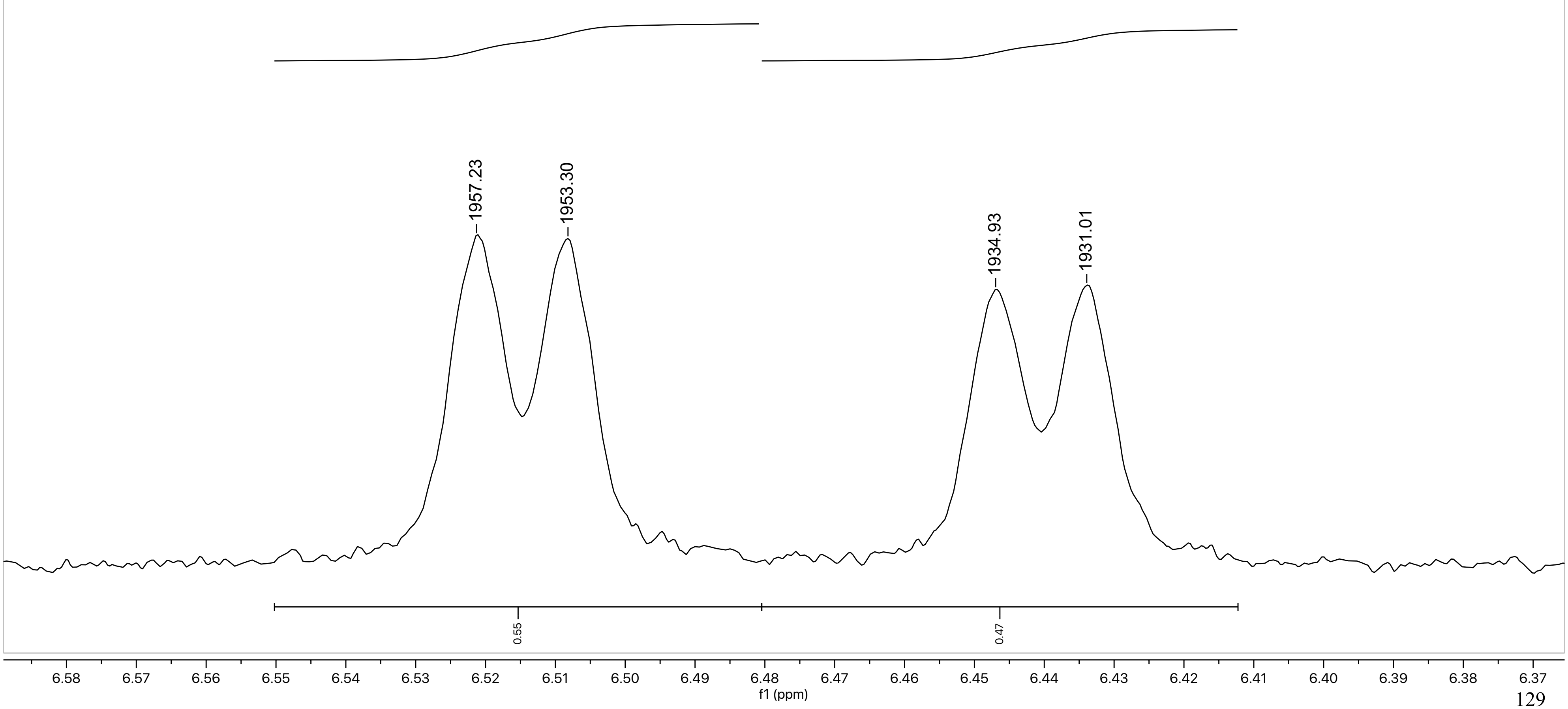

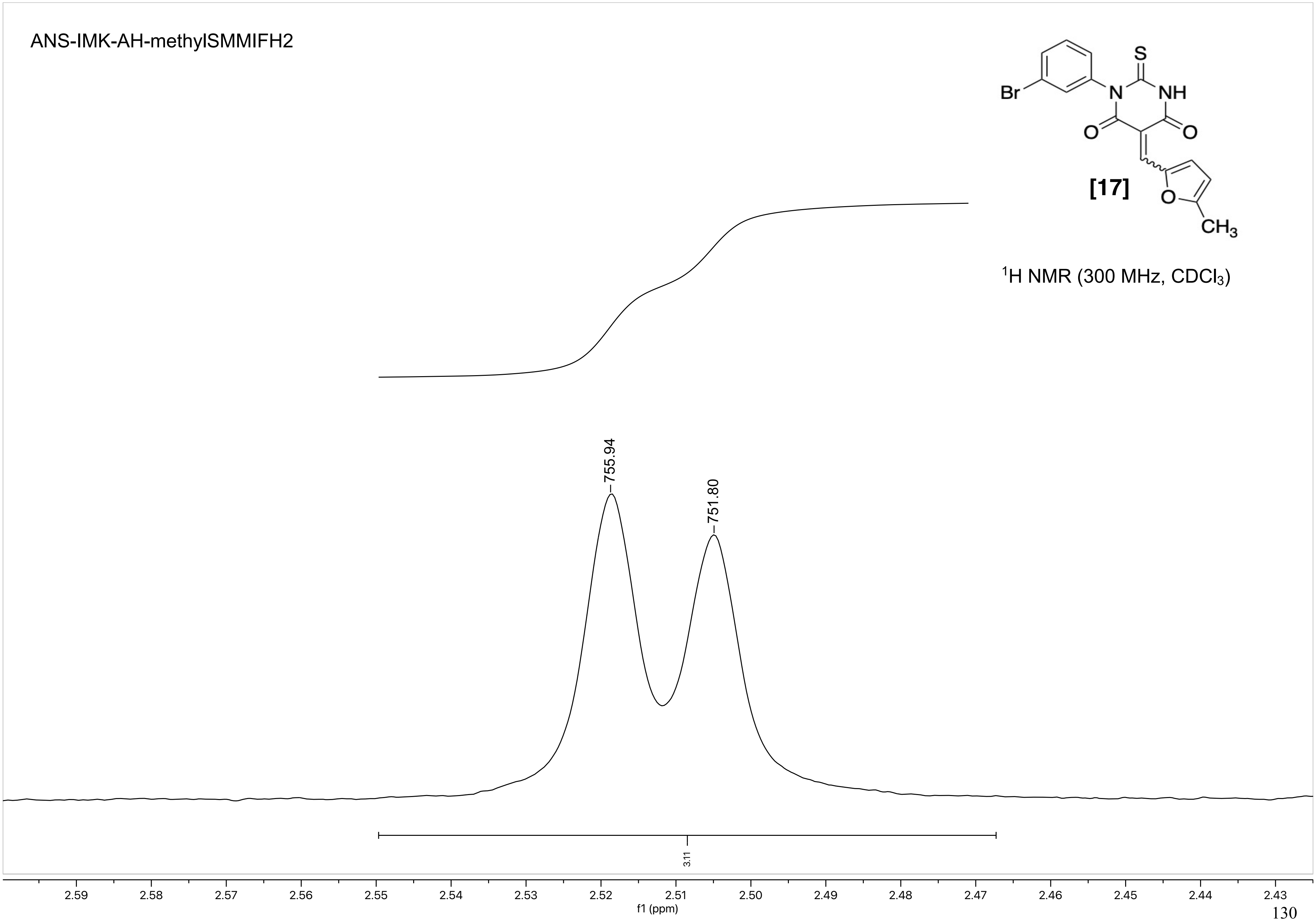

ANS-IMK-AH-methylSMMIFH2
Br
N
S
NH
O
O
[17]
O
CH3
1H NMR (300 MHz, CDCl3)
755.94
751.80
3.11
2.59
2.58
2.57
2.56
2.55
2.54
2.53
2.52
2.51
2.50
2.49
2.48
2.47
2.46
2.45
2.44
2.43
f1 (ppm)

NAME I-MAL-47
EXPNO 1
PROCNO 1

F2 - Acquisition Parameters
Date_ 20210515
Time 18.36 h
INSTRUM Avance Neo Nanobay
PROBHD Z163739_0044 (
PULPROG zg30
TD 65536
SOLVENT DMSO
NS 16
DS 2
SWH 8196.722 Hz
FIDRES 0.250144 Hz
AQ 3.9976959 sec
RG 101
DW 61.000 usec
DE 13.36 usec
TE 296.0 K
D1 1.00000000 sec
TD0 1
SFO1 400.2724717 MHz
NUC1 1H
P0 3.70 usec
P1 11.10 usec
PLW1 14.49800014 W

F2 - Processing parameters
SI 65536
SF 400.2700029 MHz
WDW EM
SSB 0
LB 0.30 Hz
GB 0
PC 1.00

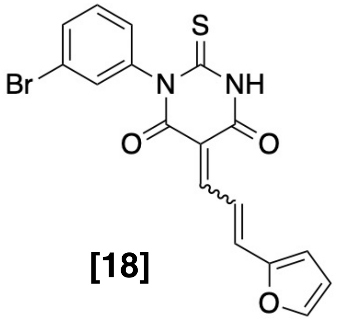

[18]

$^{1}$H NMR (400 MHz, DMSO-$d_6$)

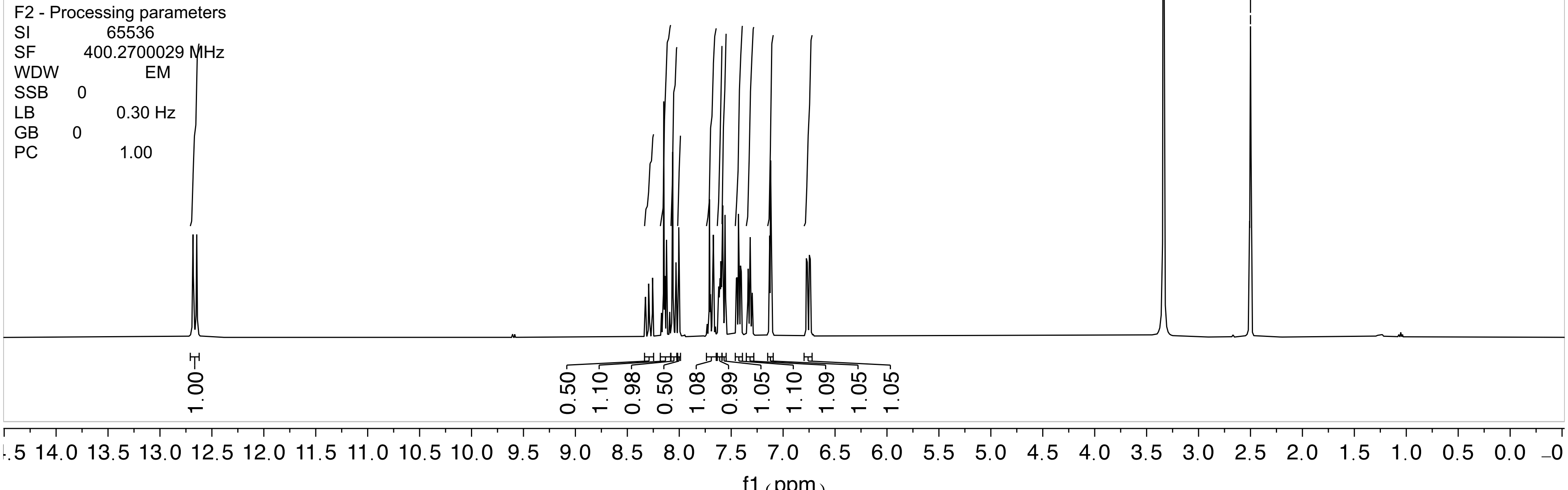

I-MAL-47

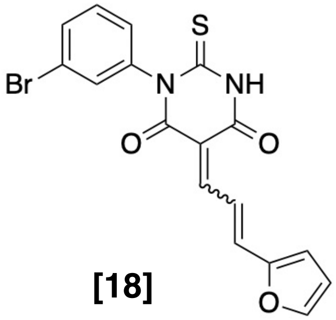

**[18]**

$^{1}H$ NMR (400 MHz, DMSO-$d_6$)

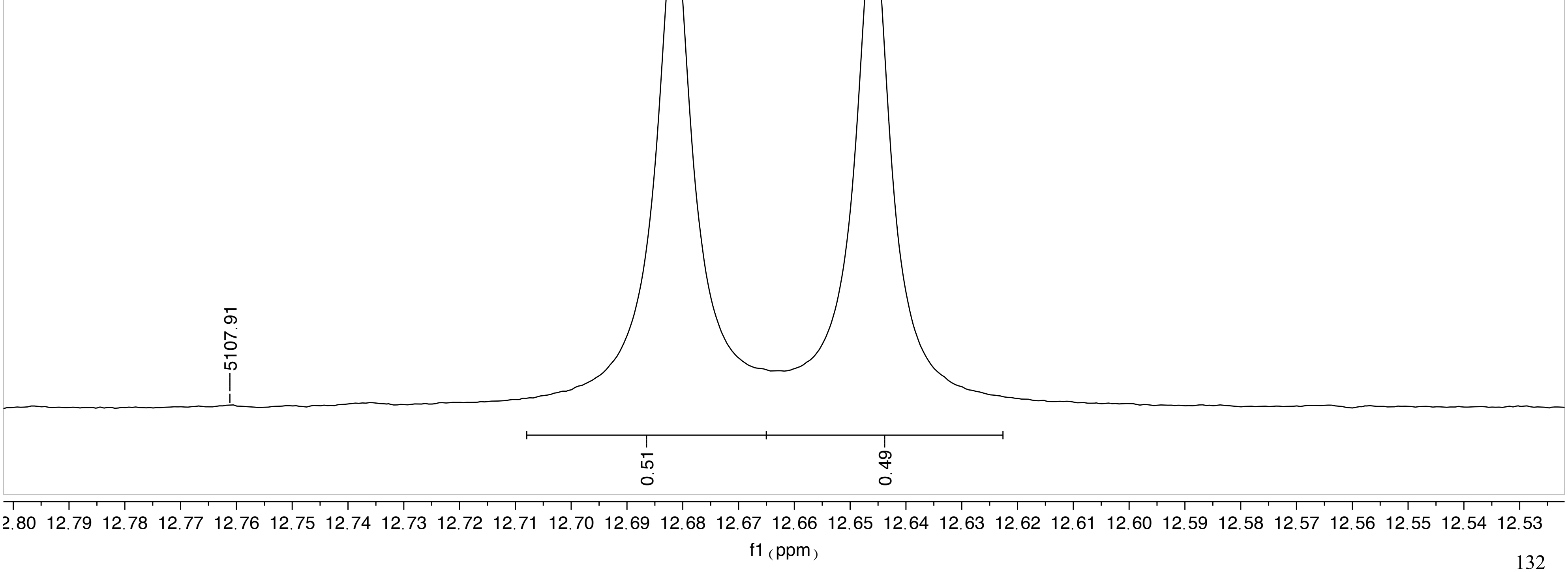

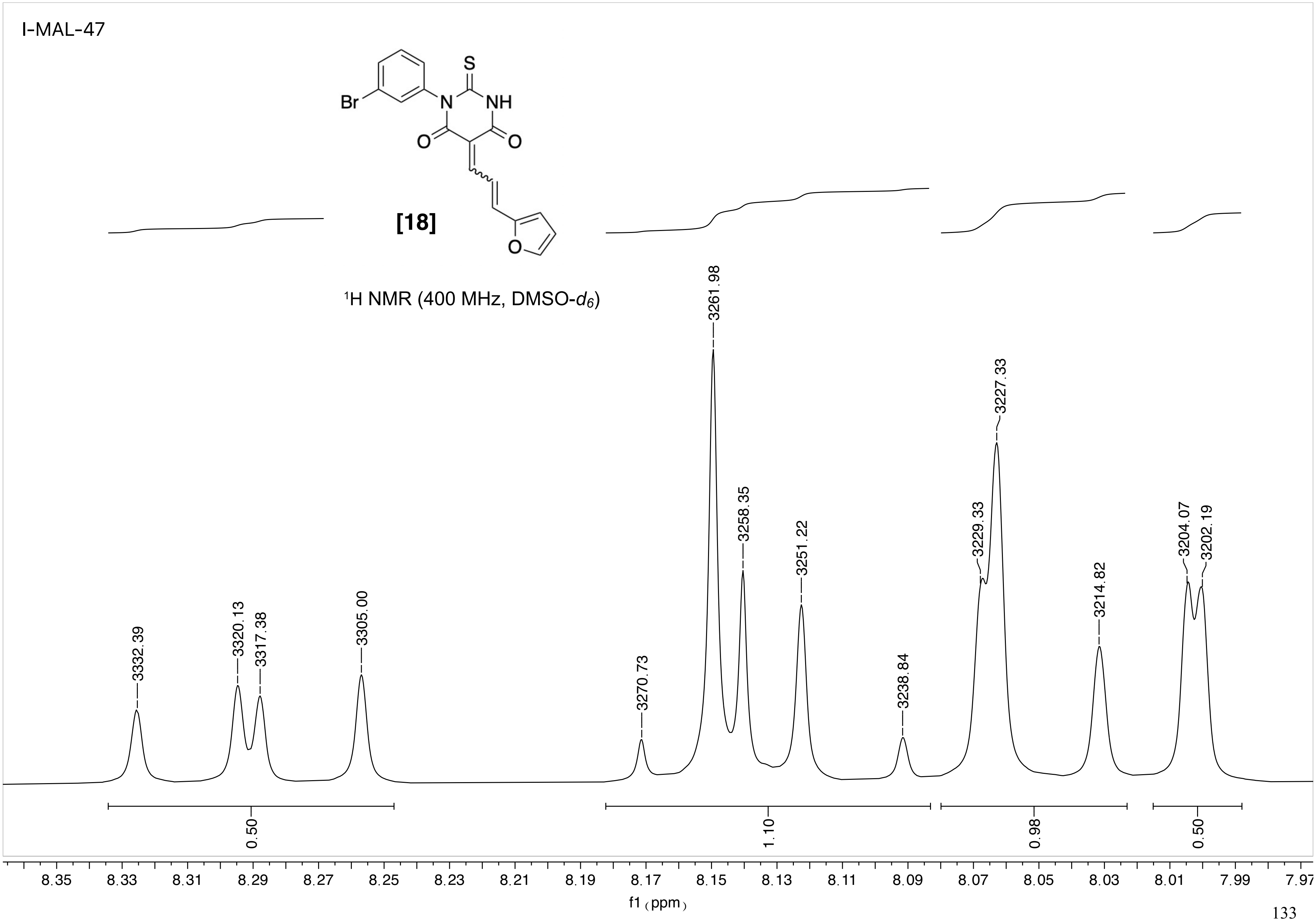

I-MAL-47
Br
N
NH
S
O
O
O
[18]
1H NMR (400 MHz, DMSO-d6)
3332.39
3320.13
3317.38
3305.00
3270.73
3261.98
3258.35
3251.22
3238.84
3229.33
3227.33
3214.82
3204.07
3202.19
0.50
1.10
0.98
0.50
8.35
8.33
8.31
8.29
8.27
8.25
8.23
8.21
8.19
8.17
8.15
8.13
8.11
8.09
8.07
8.05
8.03
8.01
7.99
7.97
f1 (ppm)

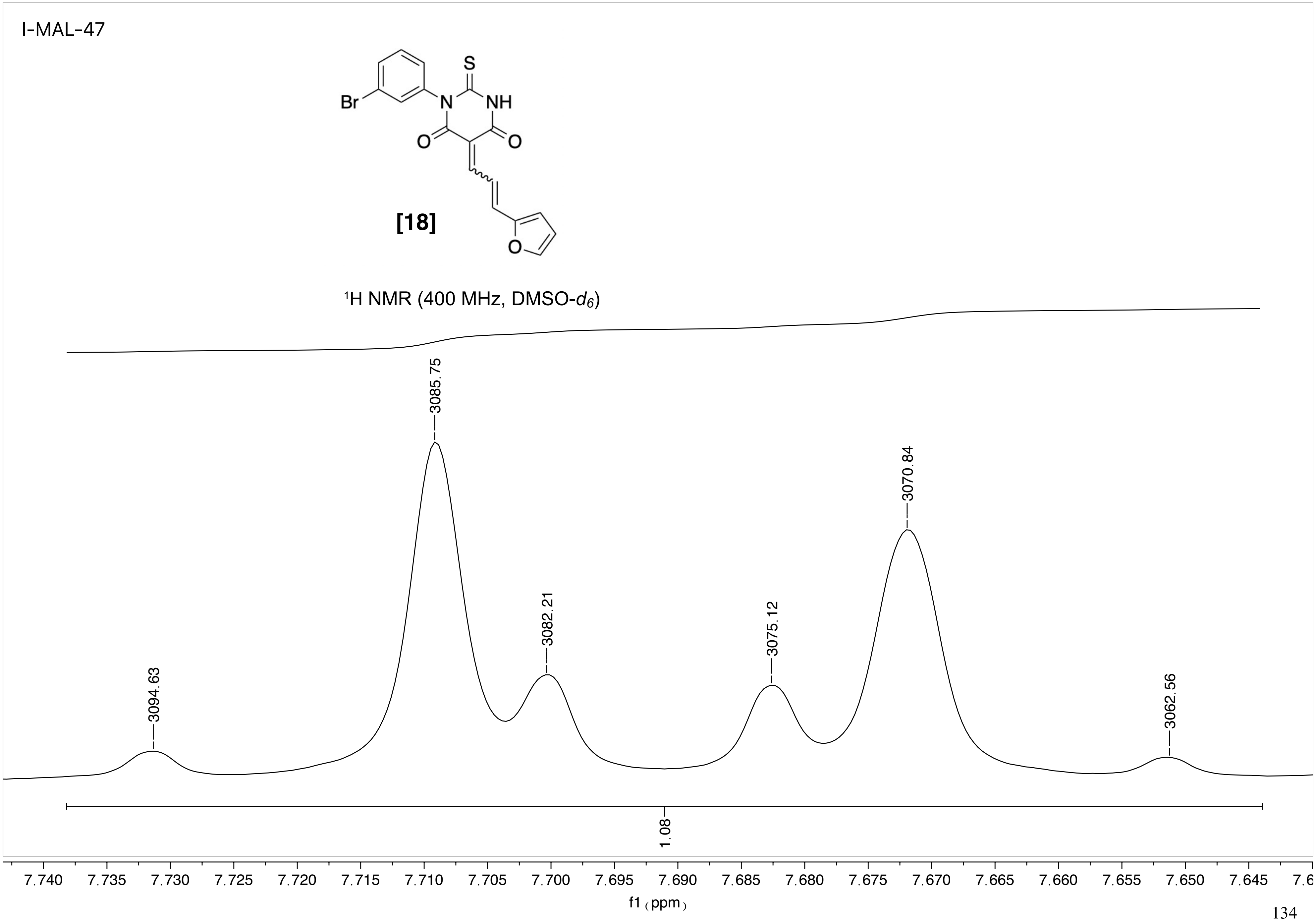
I-MAL-47
Br
S
NH
N
O
O
O
[18]
¹H NMR (400 MHz, DMSO-$d_6$)
3094.63
3085.75
3082.21
3075.12
3070.84
3062.56
1.08
7.740 7.735 7.730 7.725 7.720 7.715 7.710 7.705 7.700 7.695 7.690 7.685 7.680 7.675 7.670 7.665 7.660 7.655 7.650 7.645 7.6
f1 (ppm)

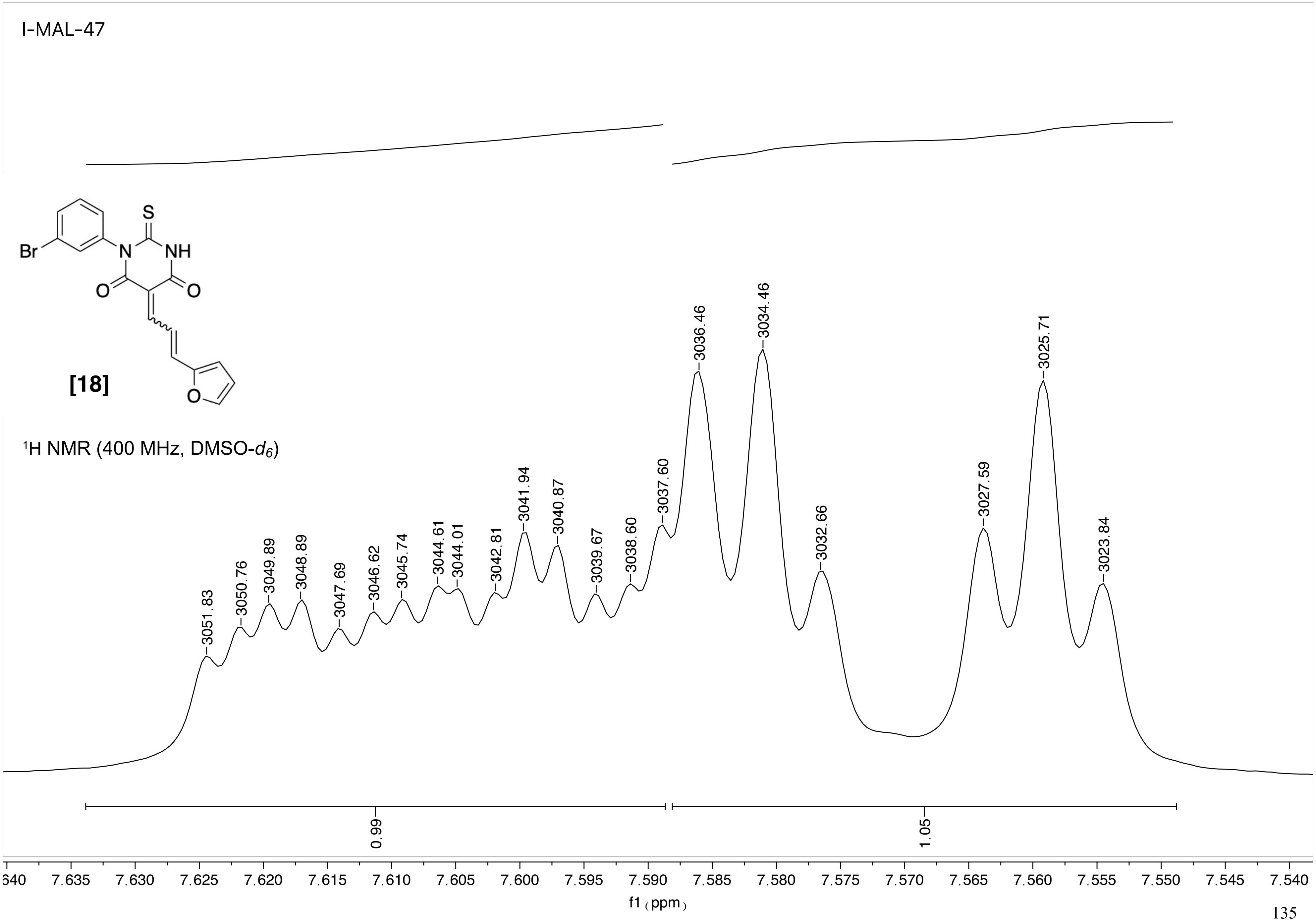

I-MAL-47
S
Br
N
NH
O
O
O
[18]
1H NMR (400 MHz, DMSO-d6)
3051.83
3050.76
3049.89
3048.89
3047.69
3046.62
3045.74
3044.61
3044.01
3042.81
3041.94
3040.87
3039.67
3038.60
3037.60
3036.46
3034.46
3032.66
3027.59
3025.71
3023.84
0.99
1.05
640 7.635 7.630 7.625 7.620 7.615 7.610 7.605 7.600 7.595 7.590 7.585 7.580 7.575 7.570 7.565 7.560 7.555 7.550 7.545 7.540
f1 (ppm)

I-MAL-47

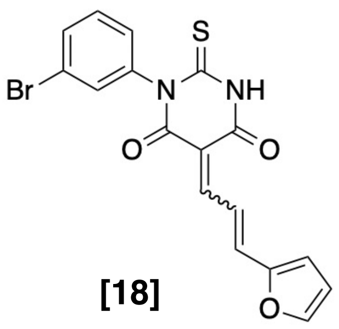

[18]

$^1$H NMR (400 MHz, DMSO-$d_6$)

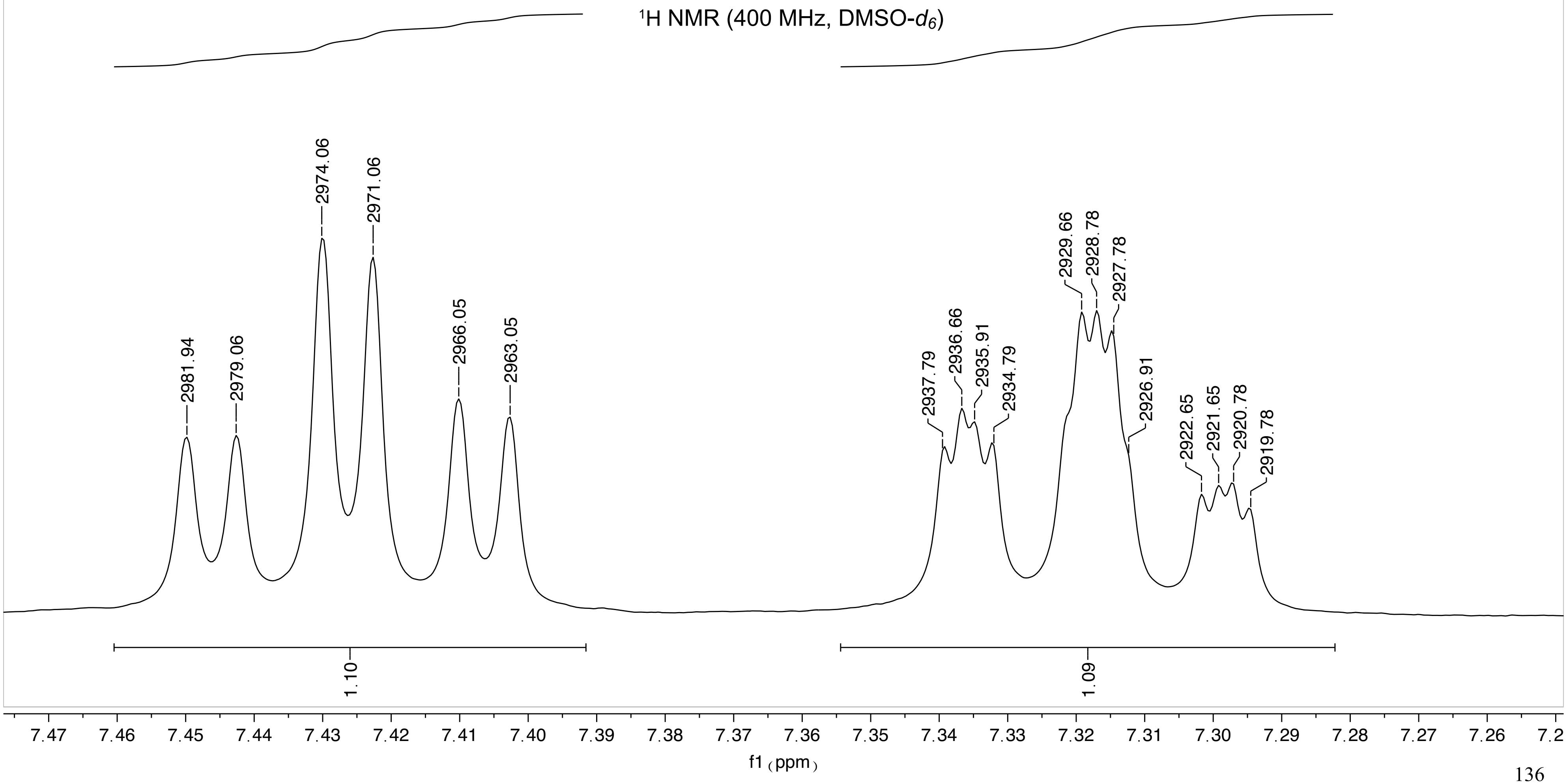

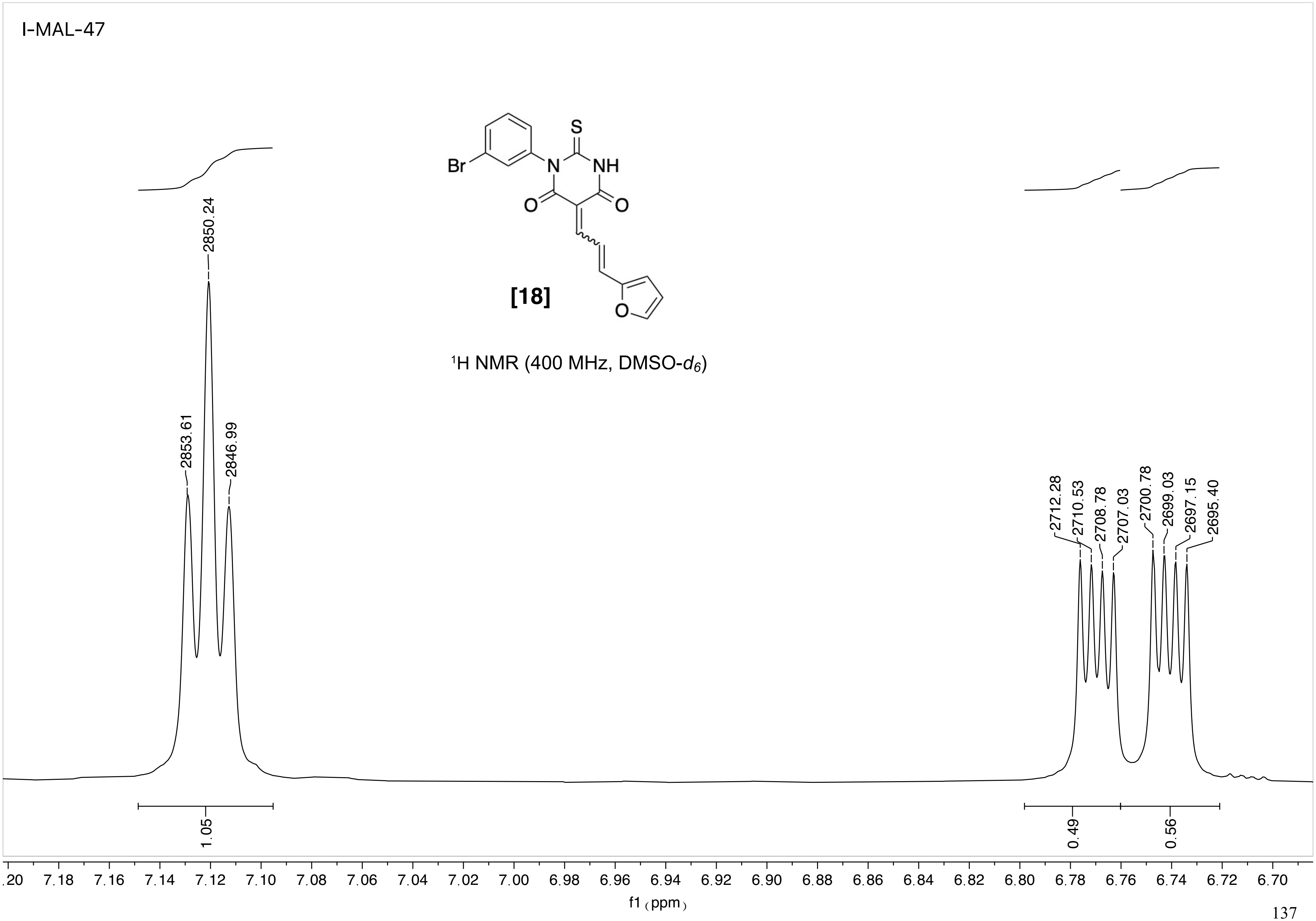
I-MAL-47
S
Br
N
NH
O
O
O
[18]
1H NMR (400 MHz, DMSO-d6)
2853.61
2850.24
2846.99
2712.28
2710.53
2708.78
2707.03
2700.78
2699.03
2697.15
2695.40
1.05
0.49
0.56
.20 7.18 7.16 7.14 7.12 7.10 7.08 7.06 7.04 7.02 7.00 6.98 6.96 6.94 6.92 6.90 6.88 6.86 6.84 6.82 6.80 6.78 6.76 6.74 6.72 6.70
f1 (ppm)

18

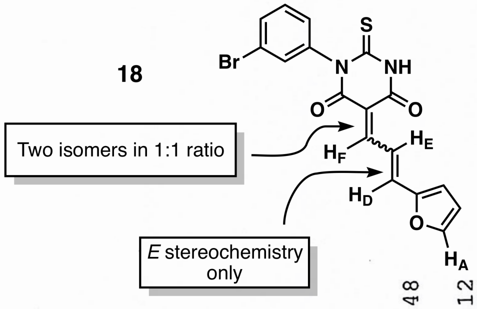

**Figure S6.** Portion of the $^{1}$H NMR spectrum of **18** (400 MHz, DMSO-$d_6$), showing second-order, virtual coupling behavior for the FED 3-spin system in one of the two double bond isomers. The isomer showing second-order behavior is labeled as isomer 2' and the isomer with a first-order spectrum is labeled as isomer 1'. The resonance for one of the furan hydrogens, $H_A$, for each of the isomers also appears in this region, and those peaks are labeled. $H_A$ is spin-isolated from $H_F$, $H_E$, and $H_D$, so we did not determine which $H_A$ signal corresponded to isomer 1' and isomer 2'. Instead, the $H_A$ resonances are labeled with un-primed designations, as isomer 1 and isomer 2.

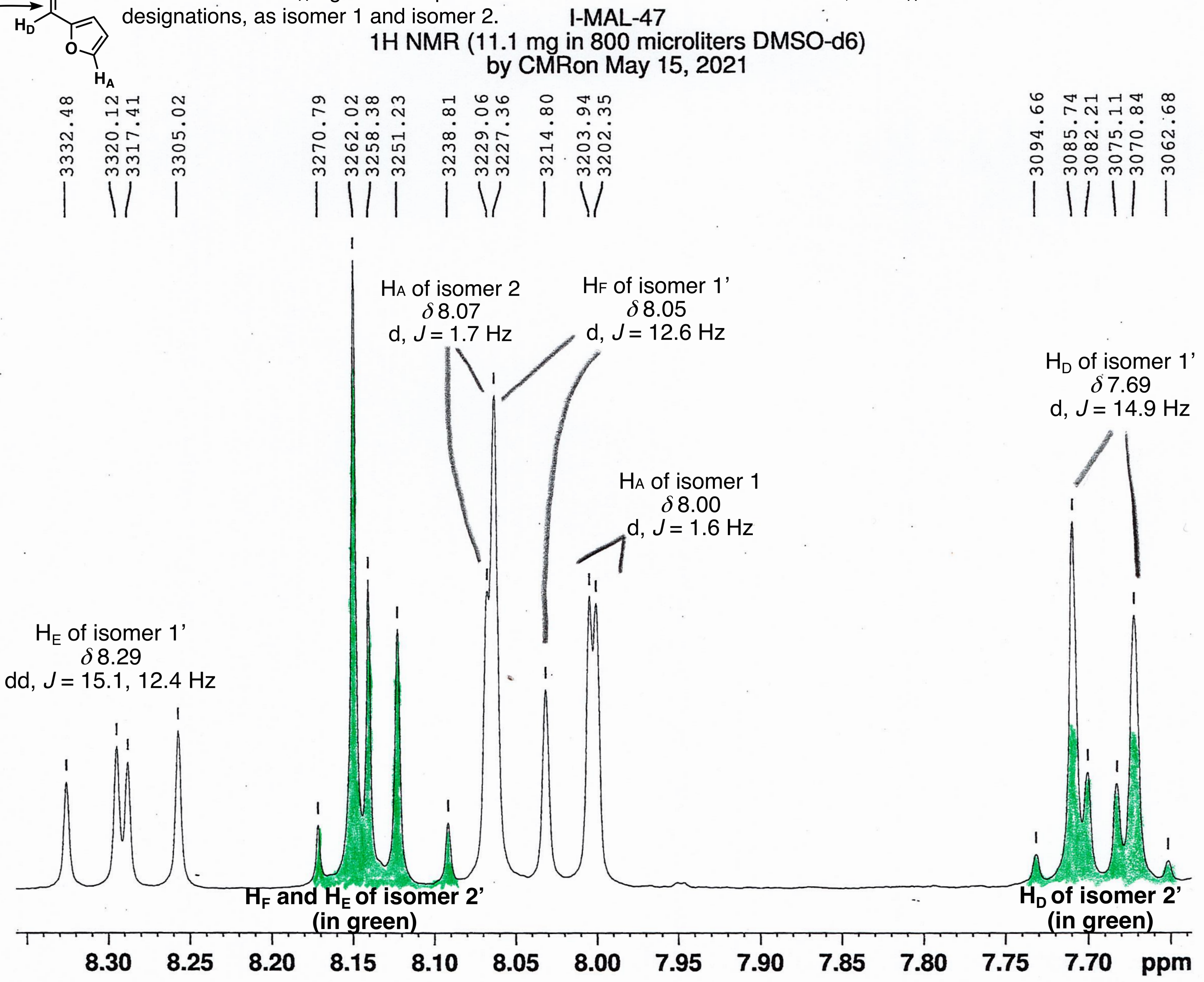

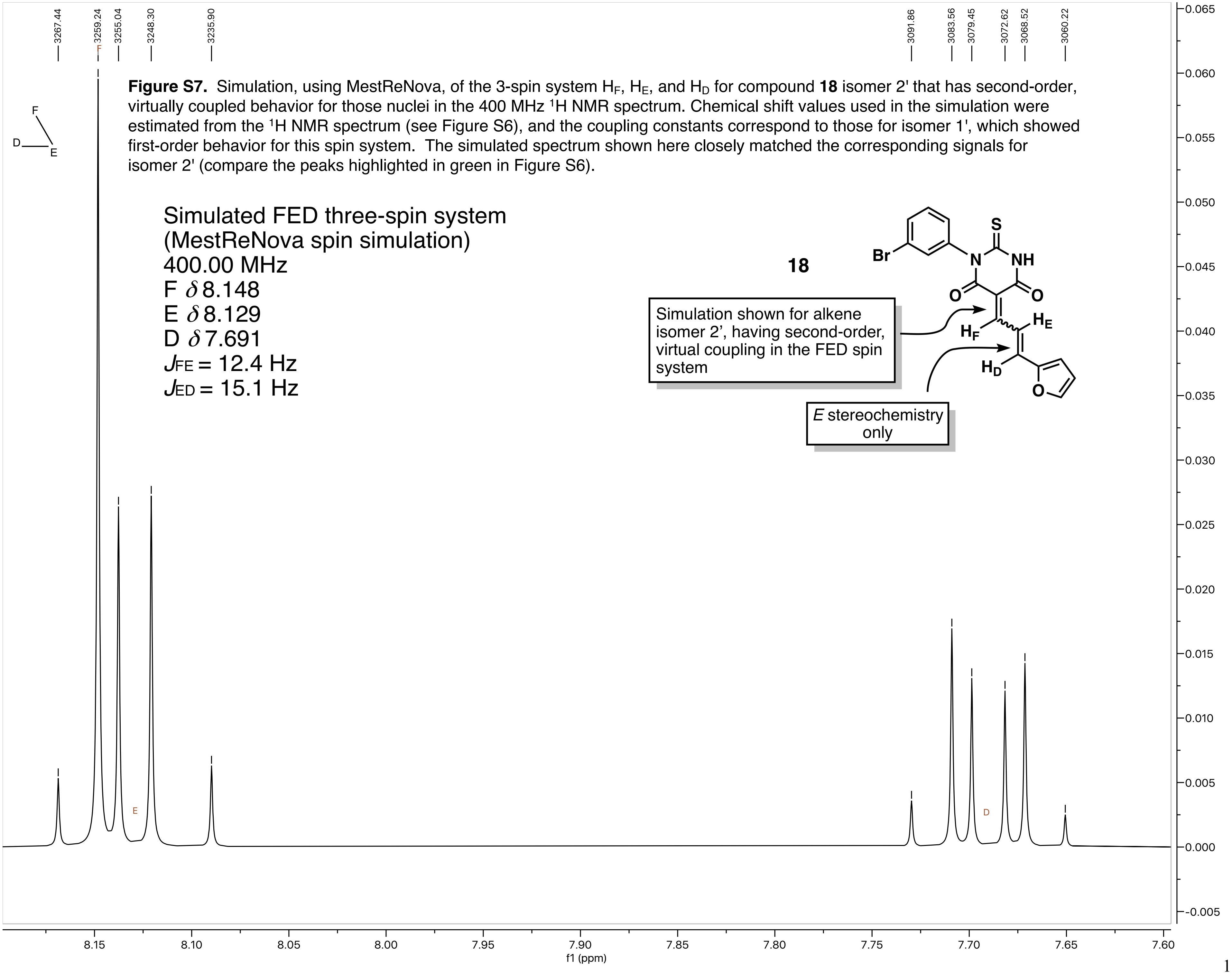

**Figure S7.** Simulation, using MestReNova, of the 3-spin system $H_F$, $H_E$, and $H_D$ for compound **18** isomer 2' that has second-order, virtually coupled behavior for those nuclei in the 400 MHz $^1$H NMR spectrum. Chemical shift values used in the simulation were estimated from the $^1$H NMR spectrum (see Figure S6), and the coupling constants correspond to those for isomer 1', which showed first-order behavior for this spin system. The simulated spectrum shown here closely matched the corresponding signals for isomer 2' (compare the peaks highlighted in green in Figure S6).
